# Bioinformatics-Guided Discovery of Biaryl-Tailored Lasso Peptides

**DOI:** 10.1101/2023.03.06.531328

**Authors:** Hamada Saad, Thomas Majer, Keshab Bhattarai, Sarah Lampe, Dinh T. Nguyen, Markus Kramer, Jan Straetener, Heike Brötz-Oesterhelt, Douglas A. Mitchell, Harald Gross

## Abstract

Lasso peptides are a class of ribosomally synthesized and post-translationally modified peptides (RiPPs) that feature an isopeptide bond and a distinct lariat fold. A growing number of secondary modifications have been described that further decorate lasso peptide scaffolds. Using genome mining, we have discovered a pair of lasso peptide biosynthetic gene clusters (BGCs) that include cytochrome P450 genes. Here, we report the structural characterization of two unique examples of (C-N) biaryl-containing lasso peptides. Nocapeptin A, from Nocardia terpenica, is tailored with Trp-Tyr crosslink while longipepetin A, from Longimycelium tulufanense, features Trp-Trp linkage. Besides the unusual bicyclic frame, longipepetin A receives an S-methylation by a new Met methyltransferase resulting in unprecedented sulfonium-bearing RiPP. Our bioinformatic survey revealed P450(s) and further maturating enzyme(s)-containing lasso BGCs awaiting future characterization.

## Introduction

Ribosomally synthesized and post-translationally modified peptides (RiPPs) represent a structurally and functionally diverse group of natural products. Through the combined effects of improved bioinformatic algorithms, genome sequencing campaigns, and isolation/characterization projects, RiPPs feature an extraordinary array of post-translational modifications (PTMs) and architectural scaffolds that cover a broad range of biological functions.^[1]^

Lasso peptides are one of more than 40 described classes of RiPPs and display a structurally unique lariat conformation as the class-defining feature. The N-terminus of the core peptide (CP) and a side chain carboxylate of an Asp/Glu residing at position 7-9^[2]^ form a macrolactam ring that is threaded by the C-terminal “tail” residues of the CP. Large, steric-locking residues adjacent to the plane of the ring and/or disulfide bridge(s) constrain the conformation of the peptide as a formal rotaxane. Such architectures endow the majority of lasso peptides with extraordinary thermal and proteolytic stability.^[3]^

Lasso peptide biosynthesis starts with the ribosomal synthesis of a bipartite precursor peptide (A) that consists of the N-terminal leader peptide (LP) and C-terminal CP. The LP harbors the recognition sequence, which directs the processing events via interaction with the RiPP precursor recognition element (RRE). Meanwhile, the CP receives all of the PTMs.^[4]^ Upon RRE binding, leader peptidolysis is initiated by the co-occurring (B) protein, releasing the CP as a substrate for the next-acting, ATP-dependent lasso cyclase (C).^[3a,5]^ In some lasso peptide biosynthetic pathways, secondary modifications include disulfide bonds, C-terminal methylation, *N*-acetylation, citrullination, *O*-phosphorylation, glycosylation, epimerization, β-hydroxylation, and aspartimidation.^[2b,6]^

The combination of genome sequencing and enhanced genome-mining algorithms has revealed a large number of RiPP-associated PTMs, including lasso peptides.^[6c,7]^ Within our current discovery program from *Nocardia* isolates,^[7d,8]^ RiPP genome mining applied to the two highly similar *Nocardia terpenica* strains IFM 0406 and 0706^T^ disclosed three putative lasso peptide biosynthetic gene clusters (BGCs) which were so far not linked to any reported natural product. One such BGC contained a member of protein family PF00067,^[9]^ a predicted cytochrome P450 protein, which lack precedent in lasso peptide biosynthesis. Since many oxidative transformations are mediated by cytochrome P450 proteins,^[10]^ we envisioned that the product(s) of the BGC would receive tailoring beyond macrolactam formation. Thus, we sought to characterize the chemical products of this pathway, which were termed the nocapeptins, given the origin was *Nocardia terpenica* (taxonomic order: Corynebacteriales). Further, we sought to characterize homologous products, termed the longipeptins, deriving from a phylogenetically distant actinomycete, *Longimycelium tulufanense* (taxonomic order: Pseudonocardiales).

## Results and Discussion

### Genome Mining Uncovers Lasso Peptide BGCs Associated with Cytochrome P450 Proteins

During our efforts to discover new RiPPs from *Nocardia*, we uncovered an unusual lasso peptide BGC (*nop*, Figure 1). The *nop* BGC is 5.4 kb in length and consists of six open-reading frames (ORFs), designated *nopA-F*. RODEO was used to examine the constituent gene products,^[7a,11]^ which predicted a class II lasso peptide encoded by *nopA*. Functional annotation of the local ORFs identified the requisite lasso cyclase (NopC, PF00733) and leader peptidase (NopB, PF13471) (Figure 1, Table S1). The RRE domain was found to be discretely encoded (NopE, PF05402) and the BGC includes a putative dedicated ABC transporter (NopD, PF00005). Notably, and as the principal criterion for target selection, a cytochrome P450 protein (NopF, PF00067) is locally encoded (Table S1).

**Figure 1.**
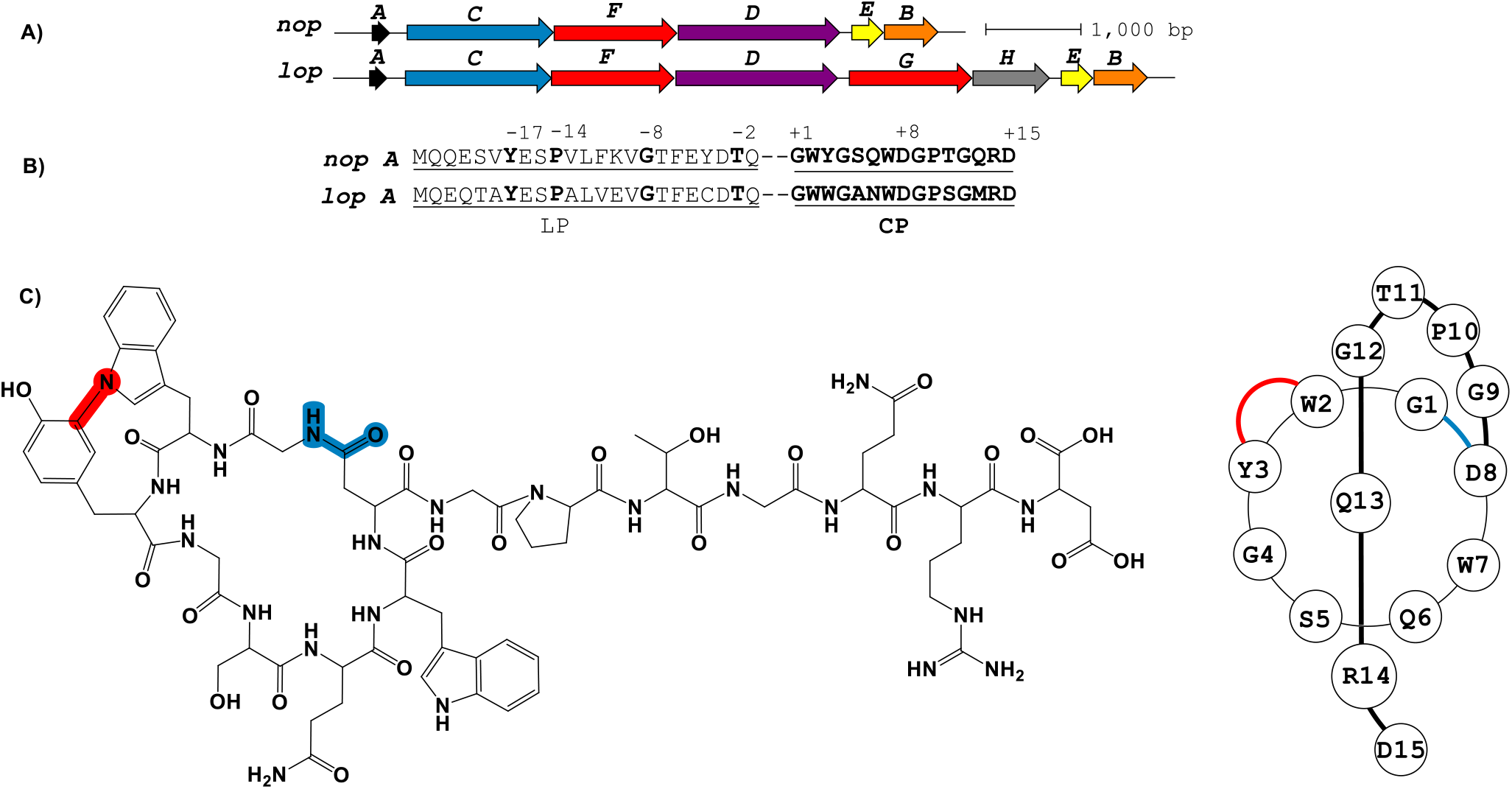
**A**) The putative *nop* and *lop* BGCs that produce nocapeptin and longipeptin, respectively. The *A, C, D, E* and *B* genes encode the precursor peptide, lasso cyclase, ABC transporter, RRE, and leader peptidase, respectively. The *F* and *G* genes encode the cytochrome P450 proteins while *H* encodes a hypothetical unknown protein. **B**) The leader and core regions of the precursor are indicated (LP and CP, respectively). Conserved LP residues are numbered and bolded. **C**) The 2D chemical structure of the bicyclic lasso peptide nocapeptin A (**1**, left); 3D representation of nocapeptin A (**1**, right).

A BLAST-P search of NopA followed by RODEO analysis unveiled a small number of similar BGCs (Figure S2). One such result was from *Longimycelium tulufanense* CGMCC 4.5737 and was termed the longipeptin (*lop*) BGC. The *lop* and *nop* BGCs exhibit considerable protein similarity and genetic synteny (Figure 1, Table S1). However, *L. tulufanense* contains two additional genes *lopG* and *lopH*, encoding a second cytochrome P450 protein and a protein of unknown function, respectively, suggesting that the longipeptin product(s) will receive additional modifications relative to the nocapeptin products.

Pathway analysis predicted leader peptidolysis would occur at Gln/Gly for NopA and LopA, yielding a pair of 15-amino acid linear core peptides (Figure 1). The cleaved CPs would then undergo macrocyclization at Gly1 and Asp8 with C-terminal threading via the action of NopC/LopC. The cytochrome P450 proteins encoded within the *nop* and the *lop* BGCs suggested additional oxidative reactions; however, bioinformatics alone could not reliably predict what specific modification(s) would be installed by these enzymes. Driven by the new CP sequences and the unprecedented co-occurrence with cytochrome P450 proteins, we initiated a media screening campaign to isolate and characterize the nocapeptin and longipeptin pathway products.

### Metabologenomic Identification of Nocapeptins and Longipeptins

Due to our familiarity with *N. terpenica* isolates, we quickly optimized the nocapeptin production conditions of IFM 0406 and subsequently applied the same to *L. tulufanense*. Fortunately, the ability to predict actionable mass values for the pathway products facilitated the analysis of the LC-MS/MS data. However, considering the mass ambiguity of the events mediated by NopF, LopF, and LopG, we considered an array of hypothetical oxidized products during MS data inspection.

From the media screen (n = 40), the MS profiles of IFM 0406 grown in a modified R4 medium at 32 °C exclusively afforded a pair of candidates with *m/z* values of 845.3562 (major) and 861.3517 (minor) as [M+2H]^2+^ species, which were designated as nocapeptin A (**1**) and B (**2**), respectively (Figure S3). The neutral molecular formula (MF) prediction of **1**, C_75_H_96_N_22_O_24_, was consistent with the macrolactam formation on the predicted CP sequence. The MF of **1** and **2** showed an additional degree of unsaturation (RDB) in the form of −2 Da (−2H) suggesting an oxidative modification within the lasso peptide (Figure S3A). Relative to **1**, the MF of **2** was consistent with two additional hydroxylation events (Figure S3A, S4C).

The collision-induced dissociation (CID) fragments of nocapeptin A/B delivered a series of y and b ions confirming the NopA amino acid sequence (Figures S4-S4B). Despite the lower probability of observing internal fragments arising from two CID-based amide bond dissociation events, an array of low-intensity y and b fragments were meticulously retrieved, providing further support of the predicted CP and corroborating the residues that form the macrolactam linkage (Tables S2-S3). Noticeably, a consistent 2 Da deviation in nearly all daughter ions, b_3_-b_13_ and the observation of typical y_1_-y_12_ fragments localized the secondary modifications to the first three CP residues, Gly1-Trp2-Tyr3.

Using similar methods, *L. tulufanense* yielded candidate products of *m/z* 851.3472 (major), 843.3501 (minor), and 844.3395 (trace), designated as longipeptins A-C (**3**-**5**), respectively (Figure S7). The predicted neutral MFs (Figure S7A) were thought to deviate from those of **1-2** primarily owing to the addition of LopG and LopH in the BGC (Figure 1). Using HR-MS/MS and comparative variations of the MFs across the ions set, longipeptin sequence relatedness to the untailored CP was supported (Figures S9-S11, Tables S4-S6). Besides an anticipated crosslink (−2H), longipeptin A (**3**), C_76_H_96_N_22_O_22_S, was envisioned to harbor secondary PTMs, including hydroxylation (+O) and methylation (+CH_2_) (Figures S12-S13). Longipeptins B (**4**) and C (**5**) were hypothesized to be a biosynthetic intermediate and oxidized artifact, respectively, with both containing just one of the additional modifications such that **4** bearing methylation and **5** undergoing oxidation (Figures S12-S13).

Despite the structural similarities between longipeptins by comparative MF analysis, the MS^2^ spectra of **3** and **4** were significantly different from **5** (Figure S8). To explore the discrepancy, we began our analysis with **5**, which yielded the most informative MS^2^ spectrum. The annotated y/b ions agreed with the expected CP displaying a Gly1-Asp8 macrolactam and loss of 2 Da within the Gly1-Trp2-Trp3 region. An exhaustive MS^2^ inspection unveiled a new series of y_n*_ (y_n_-64 Da) ions that demonstrated **5** contained methionine *S*-oxide (Figure S11, Table S6).

Initial trials to assign y ions from the MS^2^ spectra of **3** and **4** were unsuccessful. This was in contrast to the b ion series that supported the sequence and PTM locations. In **3**, the presumed crosslink and the exclusive hydroxylation were localized by the key b_3_ ion. Considering the spectral similarity of **3** and **4** upon methylation (Figure S8) and the annotated b ions, the methylation event was localized to the C-terminal three residues, Met13-Arg14-Asp15. From several possibilities, *S*-methylation of Met13 was the most plausible given such an assignment successfully dereplicated the full MS^2^ spectra of **3** and **4**. The parent ions of **3** and **4** supported *S*-methylation of Met13 by affording a characteristic M-62 fragment ion, arising from a presumed beta-elimination of dimethylsulfide (Figures 2A, 2B). In addition, the complete annotation of a new series of y fragments y_n*_, (y_n_-62) including the most intense ions y_4* - 7*_, verified such an unusual modification in the presumed structures of **3** and **4** (Figures 2C, 2D, S9-S10A, Tables S4-S5).

**Figure 2.**
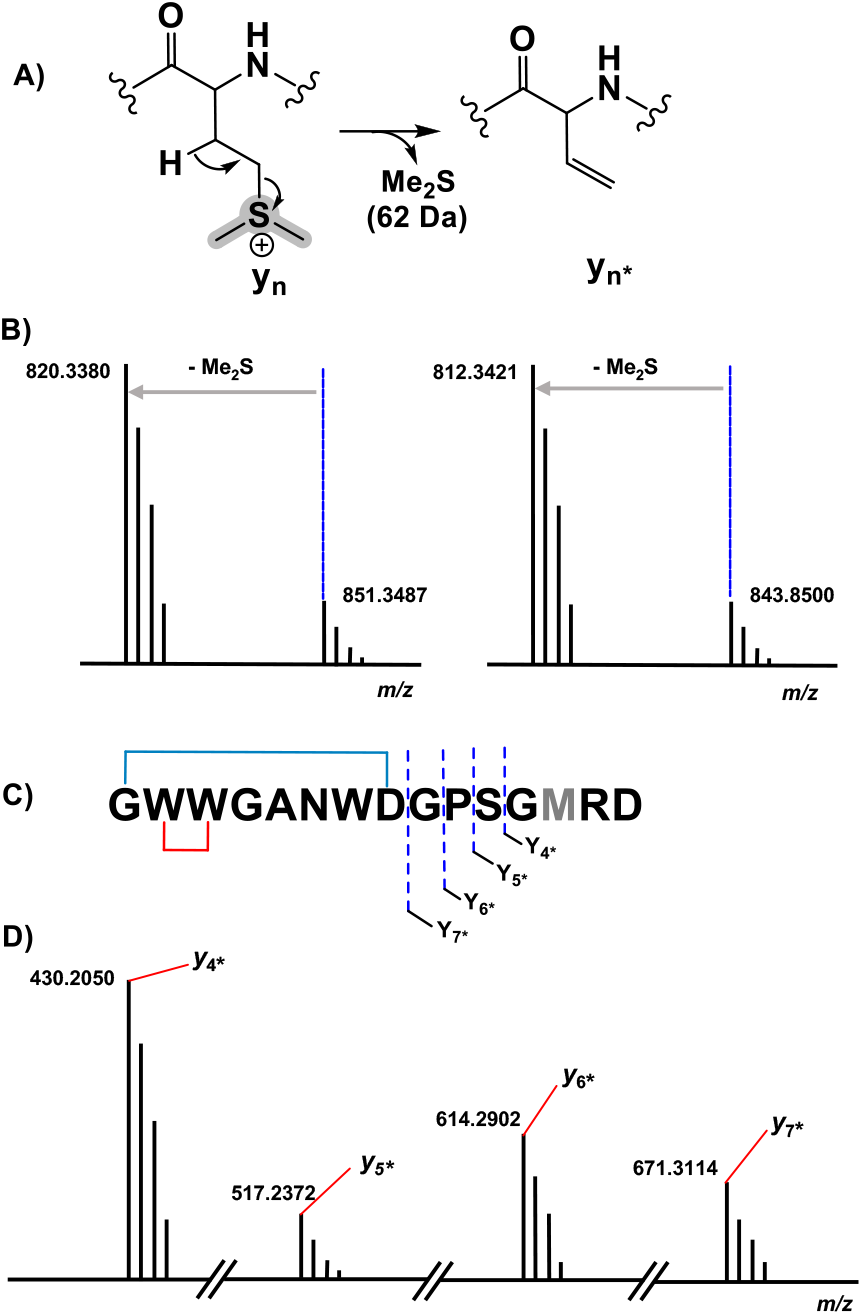
**A**) Proposed sulfonium fragmentation resulting in Me_2_S loss. **B**) Tandem mass spectra of the [M+2H]^2+^ of longipeptin A (**3**, left) and longipeptin B (**4**, right) exhibiting loss of Me_2_S. **C**) Longipeptins A (**3**) and B (**4**) sequences detailing the y_n_* ions. **D**) Annotated MS^2^ spectrum of longipeptins A (**3**) and B (**4**), [M+2H]^2+^ highlighting a new set of y ions (y_n_-62 => y_n_*).

### Stable Isotope Labelling of Nocapeptin A

To evaluate the chemical nature of the −2 Da mass deviation localized to the first three residues of nocapeptin and longipeptin, feeding studies were conducted with [^2^H_7_] L-tyrosine and [^2^H_8_] L-tryptophan. For **1**, the LC-MS profiles of IFM 0406 cultures supplemented with [^2^H_7_] L-Tyr and [^2^H_8_] L-Trp (C-D replacing all C-H) separately showed the incorporation of +6 and +16 Da, respectively (Figures 3, S5, S6). The [^2^H_7_] L-Tyr feeding data provided unambiguous proof that the modification could not be dehydrogenation and that a single C-H bond of Tyr3 was involved in the PTM. The [^2^H_8_] L-Trp data showed the proper shift of two Trp and concluded that no C-H bonds of Trp participated in the TM.

**Figure 3.**
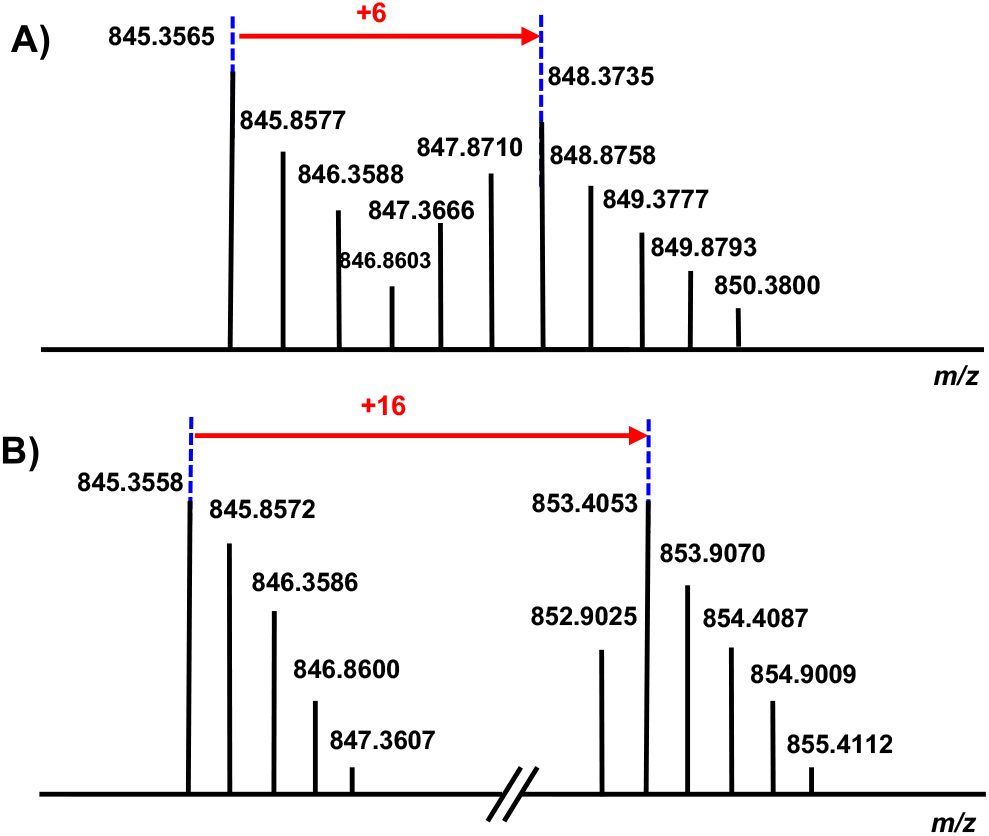
**A**) The MS profile of nocapeptin A (**1**), [M+2H]^2+^ upon the culture supplementation with [^2^H_7_] L-tyrosine. **B**) The MS profile of nocapeptin A (**1**), [M+2H]^2+^ upon the culture supplementation with [^2^H_8_] L-tryptophan.

### Isolation and Structure Elucidation of Nocapeptin A and Longipeptin A

Larger scale fermentations of IFM 0406 and *L. tulufanense* under the optimized expression conditions were next performed to isolate the quantity of material required for structure determination. Subjecting *n*-butanol extracts of the supernatants to C_18_-vacuum liquid chromatography (RP-VLC)- and RP-HPLC-guided with LC-MS enabled the isolation of nocapeptin A (**1**) and traces of longipeptin A (**3**). The elemental composition of **1** was determined as C_75_H_96_N_22_O_24_ with 39 RDB. An extensive combination of one- and two-dimensional NMR analysis was then performed to elucidate the structure of nocapeptin A.

^1^H-^1^H COSY, ^1^H-^1^H TOCSY, ^1^H-^13^C HSQC, and ^1^H-^13^C HSQC-TOCSY of the exchangeable NH protons (*δ*_H_ 6.59.0) enabled assignment of nearly all spin systems (Figures 4D, S26A, S26B, S27A, S29B). These structural fragments were complemented with ^1^H-^13^C HMBC correlations between their side chain hydrogens and carbon atoms to deliver a complete set of amino acid moieties, 4xGly, 1xSer, 2xGln, 2xAsp, 1xPro, 1xThr and 1xArg (Figures 4D, S30). Furthermore, three aromatic residues, 2xTrp and 1xTyr, were mainly disclosed with the aid of ^1^H-^13^C HMBC and ^1^H-^1^H NOESY experiments (Figures 4D, S38-S39B, S42-S42B). In contrast to the typical AA’XX’ spin pattern for Tyr, an alternative ABX/AMX coupling system with a characteristic upfield signal (*δ*_H_ 5.46, d ≈ 2 Hz)^[12]^ was observed, signifying a *meta* and *para*-disubstituted phenyl substructure. Making use of the ^1^H-^13^C HMBC, and ^1^H-^1^H NOESY couplings, such a signal was found to be the 2H-Tyr3 (Figures 4C, S39-S39B).

**Figure 4.**
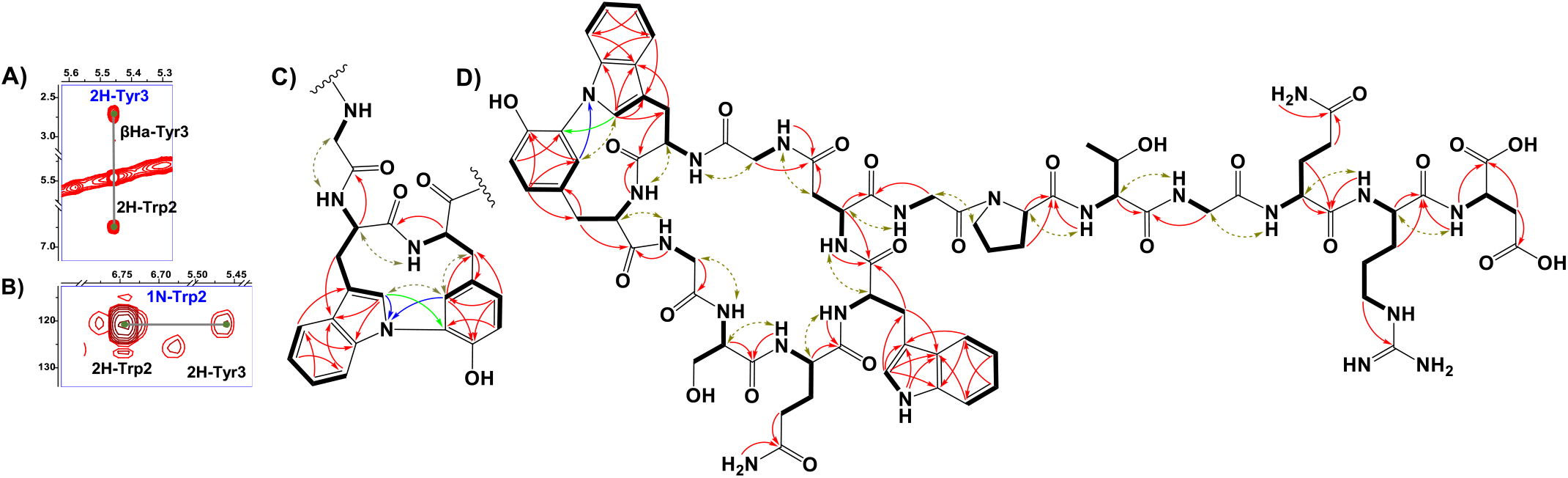
**A**) ^1^H-^1^H NOESY spectrum highlighting the correlations between 2H-Trp2 and 2H-Tyr3 centers in nocapeptin A (**1**). **B**) ^1^H-^15^N HMBC spectrum showing the correlations between the 1N-Trp2 and 2H-Tyr3 in nocapeptin A (**1**). **C**) The key NMR correlations proving the elucidated biaryl fragment (^1^H-^1^H COSY and ^1^H-^1^H TOCSY: bold lines, ^1^H-^1^H NOESY: brown arrows, ^1^H-^13^C HMBC: red arrows, ^1^H-^15^N HMBC: blue arrows and ^1^H-^13^C LR-HSQMBC: green arrows). **D**) The complete 2D-chemical structure of nocapeptin A (**1**) with the key NMR correlations.

Notably, ^1^H-NMR analysis highlighted a discriminative resonance of a single aromatic NH of one Trp residue (*δ*_H_ 10.47 ppm, broad singlet), despite two Trp present in the peptide (Figure S24). ^1^H-^13^C HMBC and ^1^H-^1^H NOESY correlations assigned the unmodified Trp as Trp7 (Figures S42-S42B), and hence, Trp2 was expected to be modified at the indolic N1 position. The observable couplings obtained from a ^1^H-^15^N HMBC spectrum, between H2-Trp2 and H2-Tyr3 with N1-Trp2 (*δ*_N_ 120.80) in addition to ^1^H-^13^C LRHSQMBC experiment,^[13]^ ^1^H-^1^H NOESY relationships, and the isotope labelling experiments permitted the assignment of a biaryl connection between the N1 of Trp2 and C3 of Tyr3 (Figures 4A-4C, S31, S33, S39A).

The polypeptide backbone of **1** was assigned via the sequential connectivity of the delineated fragments using ^1^H-^13^C HMBC cross-peaks from α-H resonances to the amide carbonyls of the neighboring amino acid. In addition, the detected H_α,β_(i)→HN(i+1) correlations in the 2D NOESY spectrum supported the complete sequence. The macrolactam between Gly1 and Asp8 was validated similarly (Figures 4D, S43). Ultimately, a threaded conformation of **1** was evident with Gln13 and Arg14 assigned as upper and lower plugs, respectively, via ^1^H-^1^H NOESY correlations (Figures S33A, S33B).

Unlike nocapeptin A (**1**), the production titer of longipeptin A (**3**) was too low to conduct a complete NMR-based structural assignment. However, keeping in mind the anticipated PTMs of **3**, the available NMR data permitted partial structural elucidation, including the PTM-containing substructures. The annotated spin systems from ^1^H-^1^H COSY, ^1^H-^1^H TOCSY, and ^1^H-^13^C HSQC-TOCSY enabled the assignment of 1xAla, 1xPro, 1xSer, 1xArg, 1xAsn, 2xAsp, and 2xGly (Figures S57, S55A). Expectedly, the NMR data showed a significant downfield shift of Met γ-CH_2_ (*δ*_C_ 43.13 instead of the typical ~30 ppm). Thus, in agreement with the MS data, we proposed this signal originated from *S*-methylation of Met13, forming a deshielded sulfonium (Figures 5A-5C, S57). In addition, three candidate spin systems including the backbone amidic NH, *α*-H, and *β*-H_a+b_ were assembled in tandem with three aromatic systems to constitute the 3xTrp units (Figures S57A, S57B).

**Figure 5.**
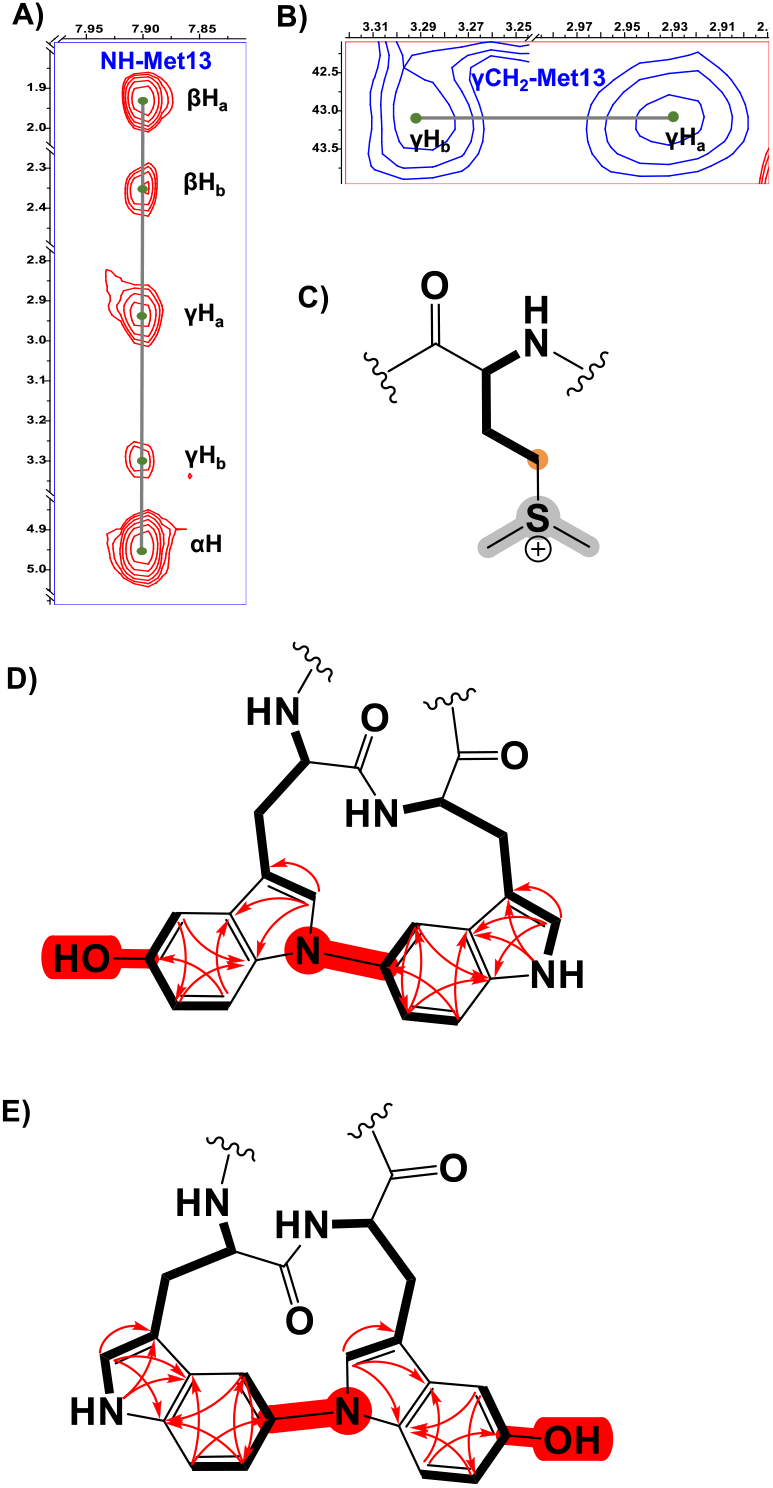
**A**) ^1^H-^1^H TOCSY spectrum showing the Met13 spin system of longipeptin A (**3**) **B**) ^1^H-^13^C HSQC spectrum highlighting the downfield crosspeaks of the γCH_2_ of S-methylated Met13 residue in longipeptin A (**3**) **C**) Structure of the Met13 sulfonium elucidated by MS/NMR. **D** and **E**) The proposed positional isomers of the (C-N) biaryl fragment of longipeptin A (**3**) with the key NMR couplings (^1^H-^1^H TOCSY, bold lines and ^1^H-^13^C HMBC, red arrows) that assembled the constituting structural units, Trp2 and Trp3.

Unfortunately, an adequate ^1^H-^1^H NOESY spectrum could not be obtained. This prevented a confident connection of the Trp separate substructures even though weaker W-couplings, ^4^*J*_2H, βHa+b_ from the ^1^H-^1^H TOCSY spectrum suggested possible connectivities (Figures S55A, S55B, S57C). The characteristic indolic NHs of 2xTrp as a pair of singlets (*δ*_H_ 10.18 and 10.55) (Figure S51A) and the ^1^H-^13^C HMBC correlations identified high-confidence locations of a crosslink and hydroxylation event in the biaryl-containing substructure (Figures 5D-E, S56A, S57C). Building on the MS data and comparable NMR shifts of **1**, two possible isomers were proposed in which the (N1-C5) biaryl crosslink was adopted alternatively between Trp2 and Trp3 (Figures 5D, 5E, S58).

### Evaluation of the Biological Activity of Nocapeptin A

Nocapeptin A (**1**) displayed no activity in the National Cancer Institute’s (NCI) cell-line cytotoxicity screen for antitumor agents. However, when assessed against a panel of bacteria, **1** exhibited moderate growth suppression of *Micrococcus luteus* (Tables S10-11). Molecular target identification will require future investigation.

### Nocapeptins and Longipeptins are Uniquely Tailored Bicyclic Lasso Peptides

The dual usage of high-resolution MS and isotopic labelling enabled the expedited discovery of new lasso peptide PTMs. Importantly, the in-depth analysis of the peptide fragments upon CID assisted in uncovering the locations of the structural modifications mediated by the cytochrome P450s in both architectures.

While the experimental strategy elucidated the final product structures of interest, the selected labelled precursor peptides were only limited to inferring a single residue of those involved in the oxidative modification of nocapeptin A (**1**). An epimerization event was considered to rationalize the mass shift upon supplementation with [^2^H_7_] L-tyrosine; however, this possibility was eliminated for two reasons: (*i*) the absence of b_2_/y_13_ ions vs the prominent presence of b_3_/y_12_ ions in **1** and **2** suggested a higher likelihood of a biaryl-linkage, and (*ii*) the absence of a local candidate epimerase in the *nop* BGC, yet having a distantly encoded one is still valid. Even though isotope-labelling studies of **1** were not definitive in assigning Trp2 as being engaged in biaryl coupling, 2D-NMR supported a linkage between N1 of Trp2 and C3 of Tyr3 to afford a bicyclic lasso peptide framework.

The homologous longipeptin BGC was expected to encode a lasso peptide with related chemical features. Unusually, longipeptin A (**3**) contained the oxidative biaryl linkage of interest in addition to indole hydroxylation and Met *S*-methylation. Despite the low production level, two positional isomers were presented for the (C-N) biaryl fragment (Figures 5D-E). The current genomic and biosynthetic setting is likely to be in favor of the substructure in which N1 of Trp2 is coupled with C5 of Trp3 (Figure 5D).

Considering the content of the *nop/lop* BGCs, the cytochrome P450 proteins Nop/LopF are suspected to form the biaryl linkages in **1** and **3**, respectively. The second cytochrome P450 protein encoded in the *lop* BGC, LopG, is predicted to be a Trp hydroxylase. In general, cytochrome P450-catalyzed crosslinks are present in several distinct RiPPs. Crocagin A^[14]^ is a tripeptide RiPP, in which the two most C-terminal residues (Tyr-Trp) undergo indole-backbone (C-N) cyclization by a dioxygenase. Atropeptides^[15]^ represent another P450-modified RiPP class that contains C-C and C-N linkages between Trp and Tyr residues. Finally, biarylitides^[16]^, also contain P450-dependent C-C or C-N linkages between Tyr and His residues.

While the biarylitides display crosslinks between the first and third residues of the CP, **1** and **3** possess biaryl crosslinks at contiguous aromatic residues. To shed light on the sequence-function relatedness of Nop/LopF versus known P450-modified RiPPs, a sequence similarity network (SSN) was constructed using the top 1000 non-redundant BLAST-P hits of NopF, predicted atropeptide- and biaryltide-associated cytochrome P450 proteins, and 883 cytochrome P450 proteins encoded within 10 ORFs of an RRE domain.^[15,17]^ The SSN and similarity/identity analysis (Figures S59, S60) suggested that NopF/LopF are rare examples of cytochrome P450 proteins within lasso peptide BGCs. A comparative analysis of sequence space also demonstrates that NopF/LopF have significantly diverged from other RiPP-associated cytochrome P450 proteins.

Perhaps the most unusual PTM described in this study is the rare *S*-methylation of Met that affords a trivalent sulfonium. Given the other functional assignments, LopH is the most probable candidate Met *S*-methyltransferase. LopH shares modest sequence similarity (45%) shared with an uncharacterized, hypothetical methyltransferase (WP_051757134.1, PF00145, Figure S61). To support or refute LopH as a methyltransferase, we obtained an AlphaFold-predicted structure and used DALI to identify tertiary fold matches from the Protein DataBank.^[19]^ The top hit was a methyltransferase domain-containing subunit of human 5,10-methylenetetrahydrofolate reductase (PDB code: 6fcx). This structure was crystallized with *S*-adenosylhomocysteine (SAH) bound (Figures S62, S63, S65).^[16]^ A comparison of 6fcx and LopH found considerable similarity in the *S*-adenosylmethionine (SAM)-binding sites (specifically, 6fcx residues Glu463, Thr464, Thr481, Ser484, Thr560, and Thr573; which are equivalent to LopH Glu45, Thr46, Thr63, Ser66, Thr126, and Thr134) (Figures S64, S66). As SAM is a common methyl donor for methyltransferases, the predicted structures and the sequence comparison support the conclusion of LopH as a novel methyltransferase that forms the sulfonium moiety in **3**.^[20]^

Sulfoniums are known to readily react with nucleophiles if dealkylation or substitution is possible.^[21]^ The *S*-methylated Met residue of longipeptin A was stable to extraction and purification. The enhanced stability may be attributed to the position of Met within the lasso peptide structure as this position is equivalent to a steric plug residue established in nocapeptin A.

*S*-methylated RiPPs are quite rare with only two cases of thiopeptides, Sch40832^[22]^ and the structurally related thioxamycin^[23]^ and thioactin.^[24]^ *S*-methylation has also been described for a proteusin that is proposed to contain a single *S*-methylated Cys.^[25]^ Thiol methylation was also illustrated in a few NRPS cases. Echinomycin and thiocoraline represent NRPS-derived products where *S*-methylation is catalyzed by a SAM-dependent methyltransferase (Ecm18) or a bifunctional enzyme (TioN) with *S*-methyltransferase and amino acid adenylation domains.^[26]^ Maremycin A/B/G and FR900452 provide further examples of *S*-methylation from BGCs containing homologs of the methyltransferase MarQ.^[27]^

These examples highlight *S*-methylation of Cys, thereby differing from the current report on *S*-methylmethionine.^[25]^ Charged sulfonium units such as dimethylsulfoniopropionate (DMSP) are well known metabolites found in marine environments. Related compounds have been described, such as gonyol and gonydiol, which are DMSP-derived biosynthetic intermediates of malleicyprols.^[29]^ Generally, DMSP-related metabolites are part of the sulfur cycle, and some marine microorganisms, they protect against osmotic, oxidative, and thermal stresses. In this context, it is notable that the longipeptins producer, *L. tulufanense*, was isolated from a high-salinity lake^[30]^ which may connect the molecular structure of **3** with DMSP ecology.

## Conclusion

In summary, we describe two lasso peptides, **1** and **3**, tailored with four novel PTMs (C-N biaryl linkages between Trp-Trp and Trp-Tyr, *S*-methyl-Met, and 5-hydroxy-Trp) which are installed by a unique combination of RiPP biosynthetic enzymes. The current findings illustrate the value of targeted genome mining in prioritizing novel BGCs from underexplored/rare actinomycetes. As shown earlier, the usage of tandem MS/isotopic incorporation facilitated the gene-to-molecule connection and also compensated for bioinformatic limitations in predicting the chemistry of the secondary PTM enzymes under investigation. Lasso peptides **1** and **3** represent highly tailored entities with 12/13 cyclic C-N biaryl systems fused with an 8-mer macrolactam cycle, respectively.

The BGCs of **1** and **3** encode unique structural features, and the bioinformatic efforts in this study expand the range of PTMs associated with lasso peptide biosynthesis, suggesting that additional oxidative tailoring steps may yet be uncovered (Figure S67). To our surprise, the co-occurrence of cytochrome P450 proteins with further maturation enzymes in some retrieved candidate BGCs presents a roadmap to discover additional lasso peptides (Figure S67). Given the structural constraint installed by NopF/LopF, future work is warranted to reconstitute the enzymatic activity and substrate scope.^[31]^ Lastly, the prediction that LopH is a founding member of a new Met *S*-methyltransferase family suggests that biochemical characterization of LopH may result in diversifying the existing methylation panel with a sulfonium PTM that can be harnessed in different bioconjugation contexts.^[32]^

## Supporting information

LopH predicted structire

## Acknowledgments

We thank Dr. D. Wistuba and her team (Mass Spectrometry Department, Institute for Organic Chemistry, University of Tübingen, Germany) for HR-MS measurements. H.S. gratefully acknowledges the Ministry of Higher Education of Egypt (MOHE) for funding. K.B. gratefully acknowledges the funding for a Ph.D scholarship from the German Academic Exchange Service (DAAD). D.A.A. acknowledges funding from the U.S. National Institute of General Medical Sciences (GM123998) while H. B.-O. acknowledges funding from the German Center for Infection Research (DZIF, TTU 09-818). Infrastructural support is from the Cluster of Excellence EXC 2124 funded by the Deutsche Forschungsgemeinschaft (DFG).

