## Supplementary material for "Bioinformatics-Guided Discovery of Biaryl-Tailored Lasso Peptides": SI document

**Author Contributions**

Project conceptualization (HS), performed experiments (HS, TM, KB, SL, DTN, MK, JS), performed data analysis (HS, TM, SL, DTN, DAM, HG), wrote the first draft (HS), edited the paper (HS, HBO, DAM, HG). All authors approved the final version of the manuscript.

**Table of Contents**

| **Experimental Procedures** | |
| --- | --- |
| 1.1 | General experimental procedures**…………………………………………………………………………………………………………………….….** |
| 1.2 | NMR Spectroscopy**……………………………………………………………………………………………………………………………………...…** |
| 1.3 | Bacterial strains**………………………………………………………………………………………………………………………………………….…** |
| 1.4 | Bioinformatics**………………………………………………………………………………………………………………………………………….…...** |
| 1.5 | Isotopic Labeling Experiments**……………………………………………………………………………………………………………………………** |
| 1.6 | Large Scale Fermentation, Extraction Scheme, Fractionation, and Isolation**……………………………………………………………………….** |
| 1.7 | Biological Assays**…………………………………………………………………………………………………………………………………………..** |
| **Bioinformatic Analysis of *nop* and *lop* BGCs** | |
| Figure S1 | Putative biosynthetic gene clusters of nocapeptin and longipeptin**……………………………………………………………………………….…...** |
| Table S1 | Putative functions of proteins from the *nop* and *lop* BGCs using the web tool RODEO**……………………………………………………………** |
| Figure S2 List of lasso peptides BGCs bearing NopA homologue using the BLAST-P search**………….……………………….……………….…….....….** | |
| **OSMAC-MSMS-Isotopes** | |
| Figure S3 | LCMS profile of modified R4 medium-based cultivation highlighting the production of nocapeptins A (**1**) and B (**2**)**………….…….……..........** |
| Figure S3A | Molecular formula predictions of nocapeptins A (**1**) and B (**2**)**……………………………………..……...……...……...……...……………….......** |
| Figure S4 | Comparative MS^2^ spectra of nocapeptins A (**1**) and B (**2**)**……………………………………………………....…...……......................................** |
| Figures S4A-4B | Annotated MS^2^ spectra of nocapeptins A (**1**) and B (**2**)**……………………………………..……………………....…...…………………................** |
| Figure S14C | Cartoon structures of nocapeptins A (**1**) and B (**2**) illustrating their annotated fragments**…………..…………………......…...…......................** |
| Figures S5-6 | Comparative MS^1^ spectra of nocapeptin A (**1**) and its [^2^H_7_] L-tyrosine/tryptophan-labeled version**………..……………………....…...……......** |
| Figure S7 | LCMS profile of modified R4 medium-based cultivation highlighting the production of longipeptins A (**3**), B (**4**) and C (**5**)**.................................** |
| Figure S7A | Molecular formula predictions of longipeptins A (**3**), (**4**) and C (**5**)**………………………...…………………………....…...……............................** |
| Figure S8 | Comparative MS^2^ spectra of longipeptins A (**3**), B (**4**) and C (**5**)**……………………...……………………………....…...……..............................** |
| Figures S9-11 | Annotated MS^2^ spectra of longipeptins A (**3**), B (**4**) and C (**5**)**……………........................................................................................................** |
| Figure S12 | Cartoon structures of longipeptins A (**3**), B (**4**) and C (**5**) illustrating their annotated fragments**…………………………..….…………………….** |
| Figure S13 | Detailed structures of longipeptins A (**3**), B (**4**) and C (**5**) illustrating the different PTMs localities**……………………………………………..…...** |
| Tables S2-3 | The assigned fragments of nocapeptins A (**1**) and B (**2**)**……............................................................................................................................** |
| Tables S4-6 | The assigned fragments of longipeptins A (**3**), B (**4**) and C (**5**)**……………........................................................................................................** |
| **400 MHz NMR dataset of 1 in *d*_3_-MeOH** | |
| Figures S14-16 | 400 MHz 1D-NMR spectra of **1…………..…..………………..…….….……………..……………………………………..…………………………..** |
| Figures S17-18 | 400 MHz ^1^H-^13^C edited HSQC and HQSC-TOCSY spectra of **1..…..………………..………………………..…….………………………………** |
| Figure S19 | 400 MHz ^1^H-^1^H COSY spectrum of **1..…..………………..………….…………….……………..…………………………………………………..…** |
| Figures S20-20A | 400 MHz ^1^H-^1^H TOCSY (with WET solvent suppression) spectra of **1….……….……………..……………………………………………………** |
| Figure S21 | 400 MHz ^1^H-^13^C HMBC spectrum of **1..…..………………..………….…………….……………..………………………………………………….…** |
| Figures S22-22A | 400 MHz ^1^H-^1^H NOESY, 300 msec (with WET solvent suppression) spectra of **1…….……….………….…………………………………………** |
| Figures S23-23A | 400 MHz ^1^H-^1^H NOESY, 500 msec (with WET solvent suppression) spectra of **1………….….………….…………………………………………** |
| Table S7 | 400 MHz ^1^H and 100 MHz ^13^C NMR data **1** in *d*_3_-MeOH**………………..…………………………….………………………………………………** |
| **700 MHz NMR dataset of 1 in *d*_3_-MeOH/H_2_O (96:4)** | |
| Figures S24-25 | 700 MHz 1D-NMR spectra of **1…………..…..………………..…….….……………..……………………………………..…………………………..** |
| Figures S26-26B | 700 MHz (Annotated) ^1^H-^13^C edited HSQC spectra of **1..…..……………………………………………….……………..………….………………** |
| Figures S27-27A | 700 MHz (Annotated) ^1^H-^13^C HQSC-TOCSY spectra of **1………...………………………………………......………………………………………** |
| Figure S28 | 700 MHz ^1^H-^1^H COSY spectrum of **1..…..………………..………….…………….……………..…………………………………………………..…** |
| Figures S29-29B | 700 MHz (Annotated) ^1^H-^1^H TOCSY spectra of **1…...……….………………………..…………..……………………………………………………** |
| Figures S30-30A | 700 MHz ^1^H-^13^C (band-selective) HMBC spectrum of **1..…..………….………….…….………..………………………………………………….…** |
| Figure S31 | 700 MHz ^1^H-^13^C LR-HSQMBC spectrum of **1……………………………………………………………………………………………………………** |
| Figure S32-33 | 700 MHz ^1^H-^1^H NOESY (300, 500 msec) spectra of **1…..…..………………..………….…………….………..……………………………………..** |
| Figure S33A | 700 MHz NOE correlations, highlighting the threading property of **1…………………………………………………………………………………..** |
| Figure S33B | Schematic representation of the lasso conformation of **1** based on NOE correlations**………………………………………………………………** |
| Figures S34-35 | 700 MHz (Annotated) ^1^H-^15^N HSQC and HMBC spectra of **1.………...………….…………………..………………………………………..………** |
| Table S8 | 700 MHz ^1^H, ^13^C and ^15^N NMR data of **1** in *d*_3_*-*MeOH/H_2_O (96:4) **…………………………..…..…………….…………….……………..…………..** |
| Figure S36 | NMR key correlations of **1………………...………….………………………………..………...…………………………………………………….….** |
| Figures S37-49 | Detailed annotation of ^1^H-^1^H TOCSY, ^1^H-^13^C HSQC-TOCSY and ^1^H-^1^H NOESY spectra of **1** residues**……………..…………………………..** |
| Figure S50 | UV, FT-IR and CD spectra of **1…………..…………….…………….……………...……………………………………………………………………** |
| **700 MHz NMR dataset of 3 in *d*_3_-MeOH/H_2_O (96:4)** | |
| Figures S51-51A | 700 MHz ^1^H NMR spectrum of **3…………………………………….……...….…………….…………….……………………………………………..** |
| Figures S52-52A | 700 MHz (Annotated indolic systems) ^1^H-^13^C edited HSQC spectra of **3..…..…...…………………………………………………………..………** |
| Figure S53 | ^1^H-^13^C HSQC-TOCSY spectrum of **3…………………………………………………………………………………………………………………….** |
| Figure S54 | 700 MHz ^1^H-^1^H COSY spectrum of **3..…..………………….…………..………….……………..…………………………………………………..…** |
| Figures S55-55B | 700 MHz (Annotated) ^1^H-^1^H TOCSY spectra **3…..………………………………………………………………..………………………..…………..** |
| Figures S56-56A | 700 MHz (Annotated indolic systems) ^1^H-^13^C HMBC spectrum of **3..…..……………………..………………………………………………….…** |
| Table S9 | 700 MHz ^1^H and ^13^C NMR data of longipeptin A (**3**) in *d*_3_*-*MeOH/H_2_O (96:4) **…………..…………….…………….……………..………….……..** |
| Figures S57-57C | Schematic representation of the assembled spin systems of **3** using 2D NMR spectra**…….…………….……………..……….…………….….** |
| Partial Structure Elucidation of **3……….…………………………………………….………………………………………………………………………………..………...…….** | |
| **Biological Assays of 1** | |
| Tables S10-11 Results of the antimicrobial and cytotoxicity (one dose NCI-60 panel) assays of **1………………………………...……………..…………………** | |
| **Bioinformatic Survey of NopF, LopF, LopG and LopH** | |
| Figure S59 | Sequence similarity network (SSN) of RiPPs-based P450s**……………………………………………………………………………………………** |
| Figure S60 | A percent/global similarity and identity matrices of NopF, LopF and LopG**…………………………………………………………………………..** |
| Figure S61 | Genomic neighborhoods of representative BLAST-P hits of LopH show co-occurrence with DNA polymerase**…………………………………** |
| Figure S62 | Tertiary structure comparison between AlphaFold-predicted LopH with SAH and its closest PDB protein structure**…………………………….** |
| Figure S63 | The pAE and pLDDT plots of the highest-ranked structure of LopH generated by AlphaFold**……………………………………………………....** |
| Figure S64 | Comparative ligand interaction for SAH docked with LopH vs the crystal structure of human 5,10-methylenetetrahydrofolate reductase.**…** |
| Figure S65 | The secondary-structure alignment generated by DALI between LopH and human 5,10-methylenetetrahydrofolate reductase**……………….** |
| Figure S66 | Comparative structures depicting SAH interaction with amino acid residues in LopH structure predicted by AlphaFold, and the crystallized human 5,10-methylenetetrahydrofolate reductase structure**…………………………….…………………………………………………………….** |
| Figure S67 | Expanded list of putative lasso peptides BGCs associated with P450 enzyme(s)**……………………………………………………………...……** |
| Table S12 | The prediction of the corresponding core peptide of p450(s)-associated lasso peptide gene cluster**……………………………………………** |
| Relevant known scaffolds containing the PTMs under investigation**………………………………………………………….……………………………………………………** | |
| **Supplemental References** | |

1. **Experimental Procedures**
   1.1 **General Experimental Procedures**

Solvents were all HPLC grade. Chemical reagents and standards were purchased from Sigma Aldrich unless indicated otherwise. The isotopically labeled substrates [L-tyrosine (D7, 98%), and L-tryptophan (D8, 97-98%)] were purchased from Cambridge Isotope Laboratories. Optical rotation values were measured on a Jasco P-2000 polarimeter, using a 3.5 mm × 10 mm cylindrical quartz cell. UV spectra were recorded on a PerkinElmer Lambda 25 UV/vis spectrometer. Infrared spectra were obtained by employing a Jasco FT/IR 4200 spectrometer, interfaced with a MIRacle ATR device (ZnSe crystal).

For Liquid Chromatography/High-resolution Electron Spray Ionization Mass Spectrometry (LC/HRESI-MSMS) measurements, an Ultimate 3000 HPLC (Thermo Fisher Scientific) system united with MaXis-4G instrument (Bruker Daltonics, Bremen, Germany) was used. The developed HPLC-method was (0.1% FA in H_2_O as solvent A and CH_3_CN as solvent B), a gradient of 10% B to 100% B in 30 min ending with 100% B for an additional 10 min, with a flow rate of 0.3 ml/min, 5 μl injection volume and UV detector (UV/VIS) wavelength monitoring at 210, 254, 280 and 360 nm. Integrating Phenomenex Luna Omega polar C18 (3 µm, 150 x 3 mm) column enabled the separation with MS acquisition range of *m/z* 50-1800. A capillary voltage of 4500 V, nebulizer gas pressure (nitrogen) of 2 (1.6) bar, ion source temperature of 200 °C, the dry gas flow of 9 l/min source temperature, and spectral rates of 3 Hz for MS^1^ and 10 Hz for MSMS were used. For MS/MS fragmentation, the 10 most intense ions per MS^1^ were chosen for subsequent collision-induced dissociation (CID) with the stepped recommended CID energies.^[1]^ For the mass calibration, sodium formate was directly infused before each sample measurement.

Vacuum liquid chromatography (VLC) was accomplished using the reversed-phase (RP) C18 column (dimensions: 10×5 cm; material: Macherey-Nagel Polygoprep 50–60 C18 RP silica gel). HPLC profiling was carried out using a system consisting of Waters 1525 Binary Pump with a 7725i Rheodyne injection port, a Kromega Solvent Degasser, Waters 2998 Photodiode Array Detector, and a Luna Omega polar C18 (5 µm, 250 × 4.6 mm, Phenomenex). ACN (solvent A) and H_2_O + 0.1% TFA (solvent B) were used for the gradient elution of the analytes with a steady flow rate of 0.5 ml/min with an injection volume of 7 μl. For the main separation and purification, the same previous RP-HPLC setup was recalled using a Phenomenex Kinetex PFP column (5 µm, 4.6×250 mm); 1 ml/min flow rate, and UV monitoring at 211, 250 and 280 nm.

1.2 **NMR Spectroscopy**

For nocapeptin A (**1**), 1D and 2D NMR spectra were measured on a Bruker Avance III HD spectrometer (400 and 100 MHz for ^1^H, and ^13^C NMR, respectively) at 297 K using a 5 mm SMART probe head. The NMR spectra were collected in (*d*_3_-CH_3_OH) processed with TopSpin 3.5 and MestReNova 12.0.4 and calibrated to the residual solvent signals (*δ*_H/C_ 3.31/49.15). Mixing times were 80 ms for ^1^H-^1^H TOCSY and 300/500 ms for ^1^H-^1^H NOESY, respectively.

Further NMR datasets for nocapeptin A (**1**) and longipeptin A (**3**), were attained from Bruker Avance III HDX spectrometer (700, 176 and 71 MHz for ^1^H, ^13^C and ^15^N NMR, respectively) in (*d*_3_-CH_3_OH/H_2_O, 96:4) equipped with a 5 mm Prodigy TCI CryoProbe head. Mixing times were 80 ms for ^1^H-^1^H TOCSY and 300/500 ms for ^1^H-^1^H NOESY. Band-Selective constant time ^1^H-^13^C HMBC spectra were recorded to dissect the peptide carbonyl region.^15^N unreferenced chemical shifts were reported in ppm (spectrometer default values). All ^13^C-^1^H LR-HSQMBC experiments were performed according to Williamson *et al*.^[2]^ with the exception that, instead of the originally proposed composite-pulse decoupling, an adiabatic pulse decoupling was employed. For each sample, we recorded four LR-HSQMBC spectra with an optimized delay Δ for a long-range coupling constant of 1, 2, 4 and 8 Hz.

**1.3 Bacterial Strains**

*Nocardia terpenica* IFM 0406 was obtained from the Medical Mycology Research Center (MMRC) culture collection, Chiba University, Chiba, Japan, while *N. terpenica* IFM 0706 (DSM 44935), and *Longimycelium tulufanense* CGMCC 4.5737 (DSM 46696) were purchased from the DSMZ (German collection of microorganisms and cell cultures).

**1.4 Bioinformatics**

The initial bioinformatic analysis was carried out using antiSMASH 5.1^[3]^ and RODEO 2.0^[4]^ to detect the putative biosynthetic gene clusters (BGCs) of nocapeptins from *Nocardia terpenica* and longipeptins from *Longimycelium tulufanense*. Using RODEO (<https://rodeo.igb.illinois.edu>) annotation, the assignment of the possible functions of each biosynthetic gene was achievable. The retrieval of further homologous nocapeptins was also facilitated by a manual BLAST-P query of the nocapeptin precursor peptide (NopA) against the non-redundant NCBI protein database in tandem with the RODEO web tool.

**AlphaFold Structural Prediction of LopH**

AlphaFold was used to predict the structure of LopH with each of the five trained model parameters.^[5]^ The multiple sequence alignment (MSA) generation and the AlphaFold predictions were enabled by ColabFold, a publicly available Jupyter notebook,^[6]^ on a Google Colab GPU cluster.

**Docking S-adenosyl-L-homocysteine (SAH) to AlphaFold-predicted LopH**

The highest ranking AlphaFold structure of LopH was aligned using PyMOL Align using the top DALI^[7]^ match as the template: human 5,10-methylenetetrahydrofolate reductase (PDB code: 6fcx, chain A), which was co-crystallized with SAH.^[8]^ The resulting coordinates from the alignment of LopH and SAH were then utilized for molecular docking using the Molecular Operating Environment 2020.0901 (MOE) software (Chemical Computing Group Inc.). This initial LopH-SAH structure was protonated with the Protonate 3D module of MOE.^[9]^ Specifically, the generalized Born/volume integral (GB/VI) approach^[10]^ was utilized to model hydration with electrostatic interactions described with Coloumb’s law with a 15 Å cutoff and dielectric of 2 inside of the protein. A dielectric constant of 80 was used to model the implicit solvent. The utilized temperature was 300K, pH was 7.0, and salt concentration was 0.1 mM. The van der Waals interactions were set with the 800R3 potential and a cutoff of 10 Å. After protonation, partial charges were assigned using the CHARMM27 forcefield.^[11]^ Energy minimization was then performed with the protonated ligand structure with convergence criteria of RMS 0.00001 kcal/mol/ Å2, with a constraint of rigid water molecules. All protein structures were visualized with either PyMOL or UCSF Chimera.^[12]^

**Generating Ligand Interaction Network using LigPlot**

The ligand interaction networks for SAH in AlphaFold-predicted LopH docked with SAH, and crystallized human 5,10-methylenetetrahydrofolate reductase (PDB code: 6fcx), were generated using LigPlot plus using standard parameters.^[13]^

**Retrieval of cytochrome P450 proteins associated with RiPP precursor recognition elements (RRE)**

Using the genome-mining tool RRE-Finder operating in precision mode^[14]^, we first compiled an updated list of all identifiable RRE domains using the most recent release of UniProt (2022_04; released on 12-Oct-2022). Members of protein family PF00067 (cytochrome P450)^[15]^ that co-occur within 10 open-reading frames of any detected RRE domain were collected (*n* = 932). Removal of identical sequences (including cases were two distinct strains produce identical proteins) yielded a set of 883 RRE-associated cytochrome P450 proteins.

**Sequence Similarity Network (SSN), Similarity and Identity Matrices Generation**

The cytochrome P450 SSN (Figure S10) was generated using EFI-EST (http://efi.igb.illinois.edu/efi-est) using an alignment score of 99 and repnode 100 parameters to represent any sequences that are 100% identical as a single node in the network.^[16]^ The sequences used were from the above-described RRE-associated cytochrome P450 dataset (n = 883 proteins), UniProt BLAST-P hits to NopF (n = 1000 proteins), a previously reported set of atropopeptide- and biaryltide-associated cytochrome P450 proteins,^[17]^ and cytochrome P450 proteins in lasso peptide BGCs predicted in this study (Figure S9). Accession identifiers of cytochrome P450 proteins annotated in this network are provided in Supplementary Dataset 1.

NopF, LopF, LopG, annotated cytochrome P450 proteins in the SSN (Figure S10), and the five top BLAST-P hits to NopF (UniProt) were aligned using the E-INS-i algorithm in MAFFT version 7 (https://mafft.cbrc.jp/alignment/server/index.html).^[18]^ The alignment was then submitted to SIAS (http://imed.med.ucm.es/Tools/sias.html) to generate similarity and identity matrices using the BLOSUM62 matrix.

**1.5 Isotopic Labeling Experiments**

*N. terpenica* IFM 0406 was revived on Brain Heart Infusion (BHI) broth agar plates (2%) at 37 °C. Colony growth was detected after 3 d of cultivation. Using fresh spores of IFM 0406, triplicates of seed cultures were prepared in Brain heart infusion (BHI) media, BHI broth 3.7%, (80 ml) in 250 ml baffled Erlenmeyer flasks at 37 °C with 150 rpm for 4 d. Starter cultures (0.4 ml) were used to inoculate 20 ml of the production medium, consisting of a modified R4 medium [glucose 0.5%, yeast extract 0.1%, MgCl_2_•6H_2_O 0.5%, CaCl_2_•2H_2_O 0.2%, K_2_SO_4_ 0.1%, casamino acids 0.05%, L-proline 0.07%, L-valine 0.12%, TES (N-Tris(hydroxymethyl)methyl-2-aminoethanesulfonic acid) 0.28%, and 50 µl trace elements solution (ZnCl_2_ 40 mg/l, FeCl_3_•6H_2_O 20 mg/l, CuCl_2_•2H_2_O 10 mg/l, MnCl_2_•4H_2_O 10 mg/l, Na_2_B_4_O_7_•10H_2_O 10 mg/l and (NH_4_)_6_Mo_7_O_24_•4H_2_O 10 mg/l)] in a 50 ml Erlenmeyer baffled flask at 32 °C with 150 rpm. The supplementation of the labeled amino acids, [^2^H_7_] L-tyrosine and [^2^H_8_] L-tryptophan into to the production medium was adjusted to a 2 mM as a final concentration of each. After 6-7 d, the cell-free supernatants were prepared by centrifugation and extracted twice with 50 ml of *n*-BuOH. The organic phases were combined, dried *in vacuo*, dissolved in MeOH, and submitted to LC/HRESI-MSMS.

**1.6 Large Scale Fermentation, Extraction Scheme, Fractionation, and Isolation**

*N. terpenica* IFM 0406 and *Longimycelium tulufanense* were grown using the nutrients recipe and the growth parameters as previously described in the above section, except the 250 ml Erlenmeyer baffled flasks were filled with 120 ml of the production medium. A 25 L cultivation was done in the case of nocapeptin A (**1**) and around 40 L for longipeptin A (**3**). To remove the cells, the liquid cultures were centrifuged twice in a Thermo Scientific Heraeus Multifuge 4KR centrifuge at 4000 g at 4 °C for 30 min. Subsequently and using n-BuOH (1:1), the cell free supernatants (SN) were extracted twice. Under reduced pressure, the *n*-BuOH extracts were evaporated affording the crude extracts (Bu SN extracts), which were resuspended in methanol followed by centrifugation to get rid of debris prior to LC/MS analysis, HPLC profiling, and VLC. Fractionation of the Bu SN extracts was accomplished through a VLC system by stepwise elution of H_2_O mixed with methanol controlled by vacuum with a decreased polarity fashion, shifting from 100% H_2_O to pure MeOH in ten fractions (750 ml per fraction). Guided by LCMS, the prioritized VLC fractions, 60% MeOH VLC in *N. terpenica* and 80 plus 90% MeOH VLC in *L. tulufanense*, were redissolved in a MeOH/H_2_O and filtered prior to their injection into an RP-HPLC system with an optimized polar gradient for 28 min using the formerly described HPLC setup and integrated with a Phenomenex Kinetex PFP column (5µm, 4.6×250 mm) to isolate the pure lasso peptides, nocapeptin A (**1**) (≈17 mg) and longipeptin A (**3**) (≈2 mg).

**1.7 Biological Assays**

**Antibacterial assay**

The minimal inhibitory concentration (MIC) was determined as described previously^[19]^ in cation-adjusted Mueller–Hinton medium according to the standards and guidelines of the Clinical and Laboratory Standards Institute.^[20]^ A 2-fold serial dilution of the test compound was prepared in microtiter plates and seeded with a final test bacterial inoculum 5 × 10^5^ colony-forming units (CFU)/ml. After an overnight incubation at 37 °C, the MIC was read as the lowest compound concentration preventing visible bacterial growth. The strain panel included representative “**ESKAPE**” human pathogenic bacteria. Specifically, the following strains were used: ***E****nterococcus faecium* BM 4147-1, ***S****taphylococcus aureus* ATCC 29213, ***K****lebsiella pneumoniae* ATCC 12657, ***A****cinetobacter baumannii* 09987, ***P****seudomonas aeruginosa* ATCC 27853, and ***E****nterobacter aerogenes* ATCC 13048. Additional MIC testing was performed on: *Bacillus subtilis* 168, *Enterococcus faecalis* ATCC 29212, *Escherichia coli* ATCC 25922, *Escherichia coli* HN 818, *Micrococcus luteus* ATCC 4698, *Neisseria gonorrhoeae* ATCC 19424, *Neisseria gonorrhoeae* S1441, and *Mycobacterium smegmatis* mc^2^ 155 ATCC 700084.

**Cytotoxicity assay (one dose NCI-60 panel)**

**1** was selected for the anticancer drug screening service as a part of the Developmental Therapeutics Program at the National Cancer Institute (NCI). *In vitro* tumor growth inhibitory effects were explored using a standard protocol with a single high dose test against a panel comprising 60 human cancer cell lines.^[21]^ Results for cell line NCI-H23 are omitted based on NCI staff informing that authentication was not performed during the screening time frame.

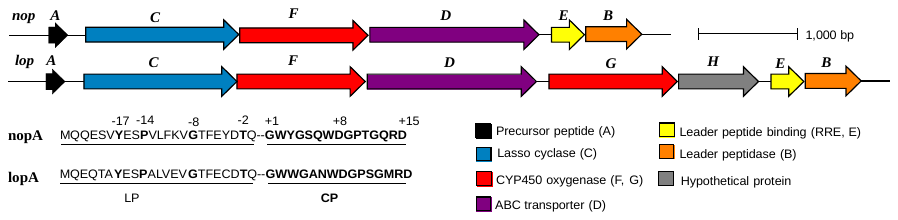

**Figure S1**. (Top) Putative biosynthetic gene clusters (BGCs) of nocapeptin, *nop* and longipeptin, *lop* from *Nocardia terpenica* and *Longimycelium tulufanense*, respectively. (Bottom left) Products of *nop*A and *lop*A, respectively, including prediction of the cleavage site between the leader peptide (LP) and core peptide (CP).

**Table S1.** Putative functions of proteins from the *nop* and *lop* BGCs using RODEO

| **RODEO Analysis** | | | | | | |
| --- | --- | --- | --- | --- | --- | --- |
| *nop* BGC | Protein | NCBI Accession ID | Length [aa] | PFAM | Description | E-value |
|  | NopA | WP_195116738.1 | 43 | ------------ | ------------ | ---------- |
|  | NopC | WP_171983240.1 | 547 | PF00733 | Asparagine synthase | 4.20E^-36^ |
|  | NopF | WP_156674500.1 | 404 | PF00067 | Cytochrome P450 | 4.00E^-50^ |
|  | NopD | WP_171983239.1 | 575 | PF00005 | ABC transporter | 9.40E^-96^ |
|  | NopE | WP_067588831.1 | 86 | PF05402 | Stand alone lasso RRE | 2.10E^-27^ |
|  | NopB | WP_082871451.1 | 157 | PF13471 | Transglutaminase | 7.60E^-24^ |
| *lop* BGC | LopA | WP_189061731.1 | 38 | ------------ | ------------ | ---------- |
|  | LopC | WP_194500064.1 | 536 | PF00733 | Asparagine synthase | 1.40E^-38^ |
|  | LopF | WP_189061729.1 | 404 | PF00067 | Cytochrome P450 | 1.10E^-50^ |
|  | LopD | WP_189061728.1 | 596 | PF00005 | ABC transporter | 1.20E^-94^ |
|  | LopG | WP_189061727.1 | 406 | PF00067 | Cytochrome P450 | 1.00E^-46^ |
|  | LopH | WP_189061726.1 | 191 | ------------ | ------------ | ---------- |
|  | LopE | WP_189061725.1 | 85 | PF05402 | Stand alone lasso RRE | 3.50E^-27^ |
|  | LopB | WP_189061724.1 | 137 | PF13471 | Transglutaminase | 1.80E^-26^ |

**A**

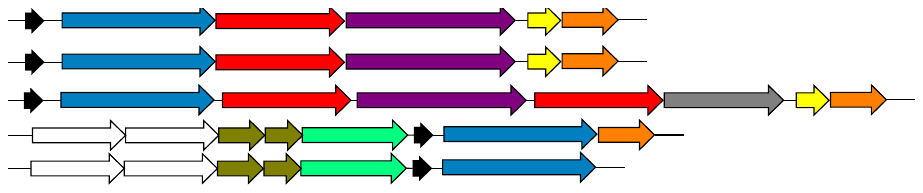

**B**

*S. roseoverticillatius*

*L. tulufanense*

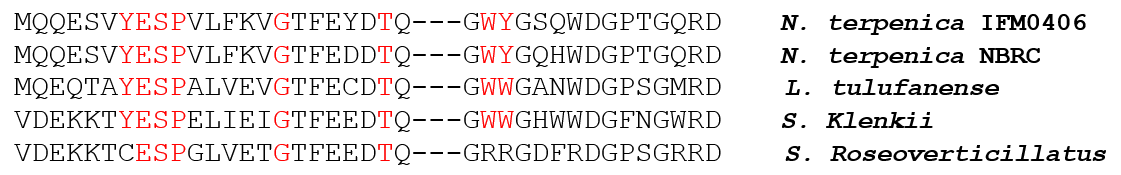

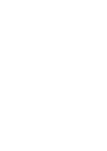

*N. terpenica* IFM 0406

*N. terpenica* NBRC

*Nocardia terpenica* NBRC

*Longimycelium tulufanense*

*Streptomyces klenkii*

*Nocardia terpenica* IFM 0406

*Streptomyces roseoverticillatius*

*S. klenkii*

**Figure S2**. Putative list of lasso peptides BGCs bearing *nop A* homologue using BLAST-P retrieval. (**A**) Predicted biosynthetic gene clusters and bacterial strains (**B**) sequence alignment of their lasso peptide precursors. The red-colored residues in the LP portion are predicted to be either the recognition sequence (YxxP) or the conserved residues in lasso peptide precursors (G and T). The red-colored residues in the CP portion (WY and WW) are predicted to be the motifs of the biaryl crosslink.

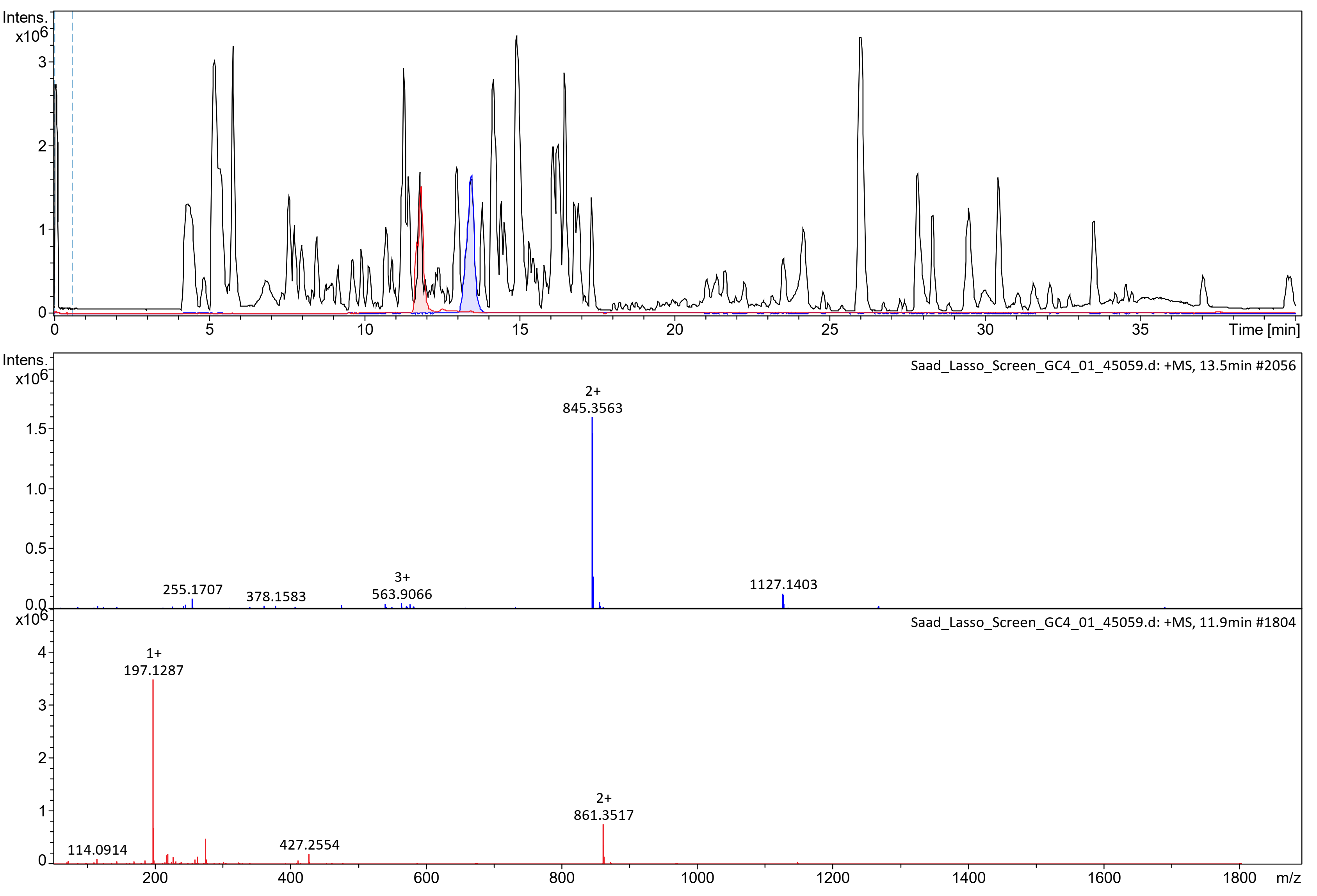

**Figure S3.** LCMS profile of modified R4 medium-based cultivation highlighting the production of nocapeptin A (**1**) [845 Da] and nocapeptin B (**2**) [861 Da].

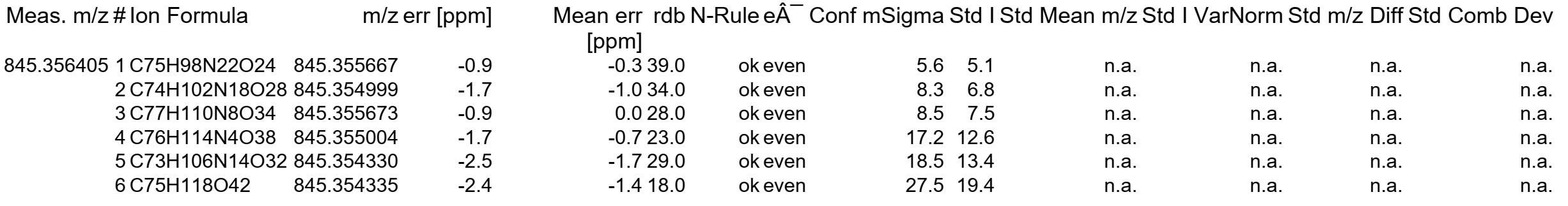

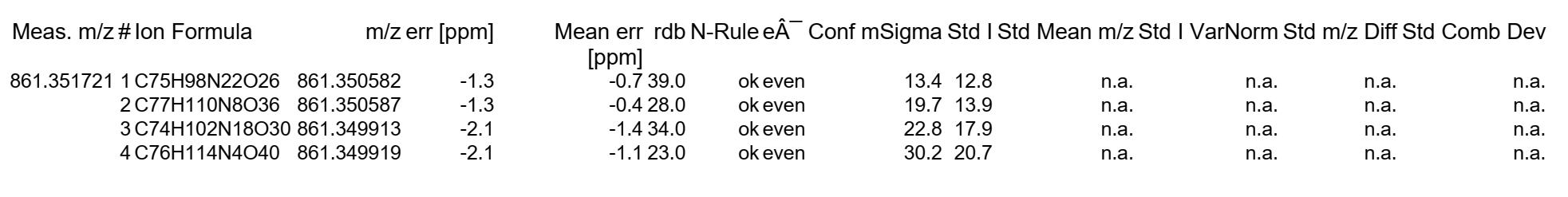

**Figure S3A**. Molecular formula predictions of nocapeptin A (**1**) [845 Da] and nocapeptin B (**2**) [861 Da]

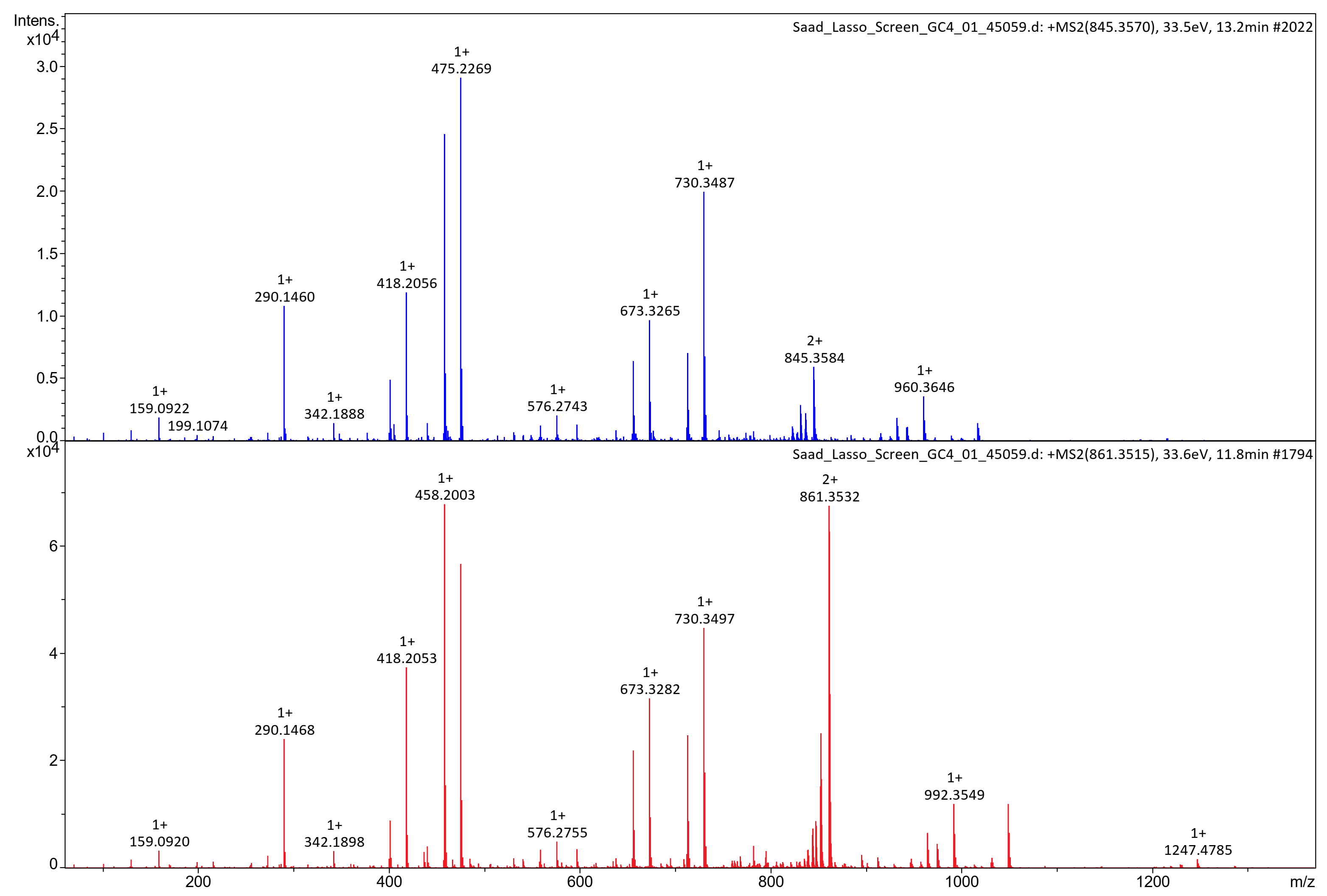

**Figure S4.** Comparative MS^2^ spectra of nocapeptin A (**1**) [845 Da] and nocapeptin B (**2**) [861 Da]

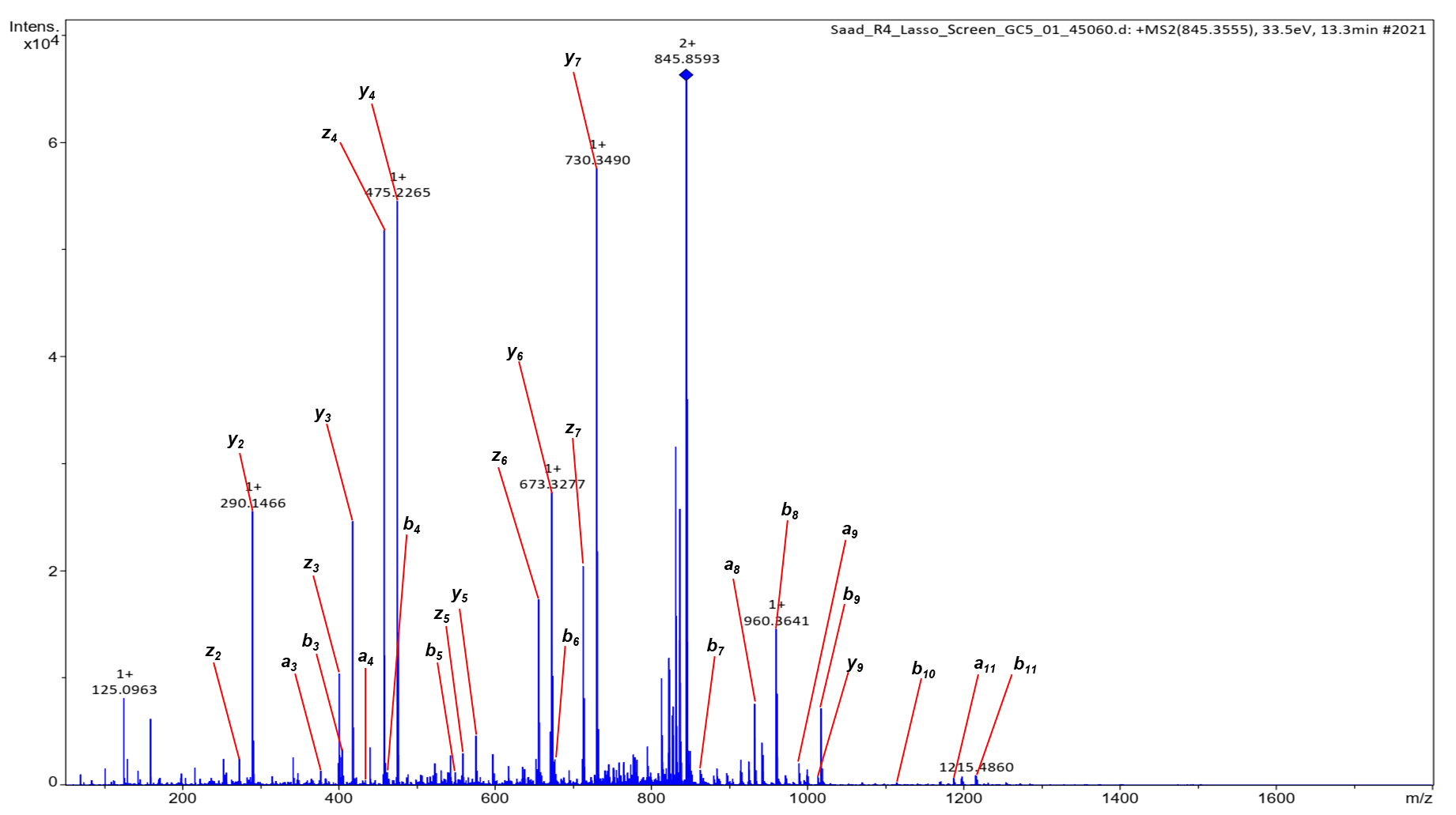

**Figure S4A.** Annotated MS^2^ spectrum of nocapeptin A (**1**) [845 Da]

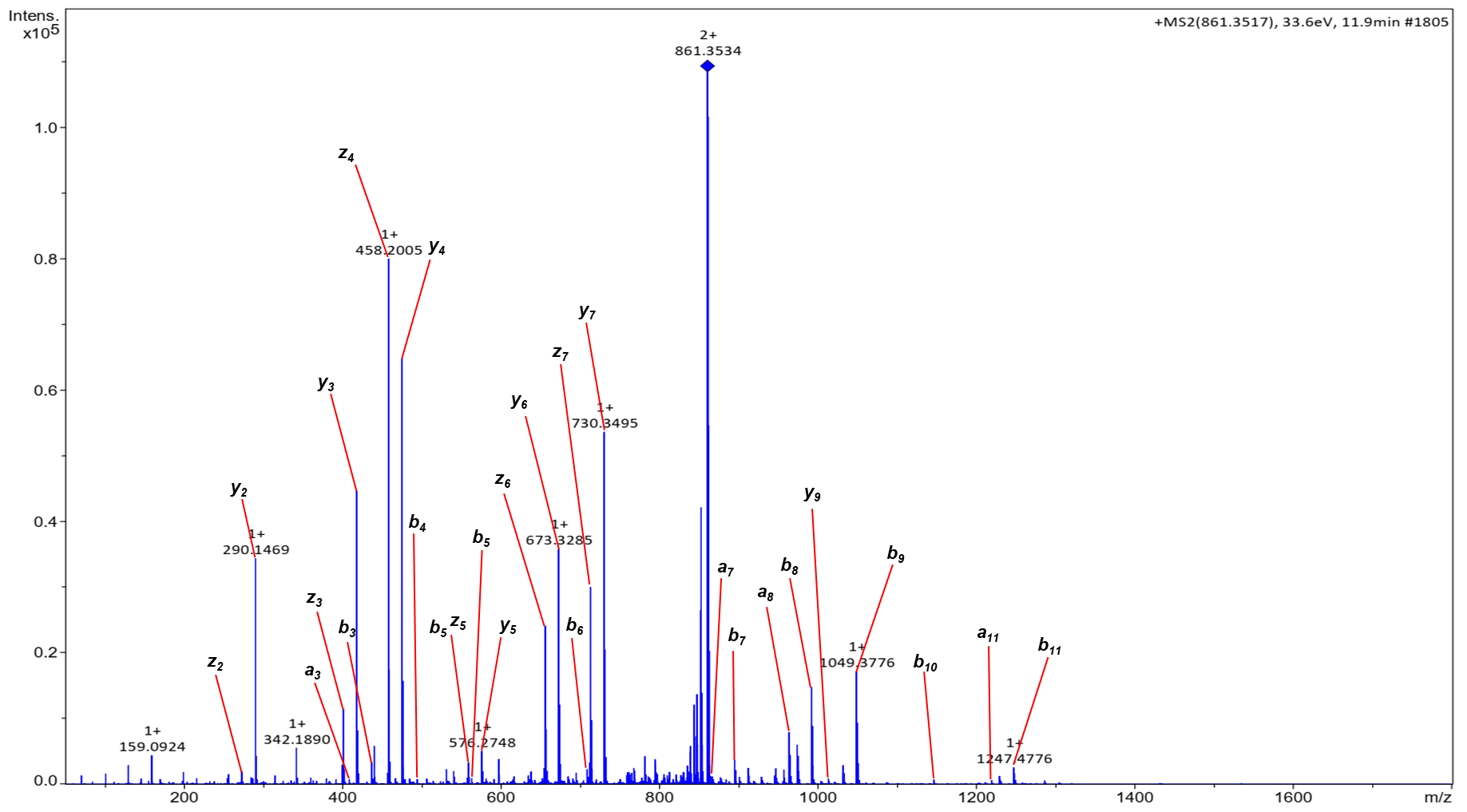

**Figure S4B**. Annotated MS^2^ spectrum of nocapeptin B (**2**) [861 Da]

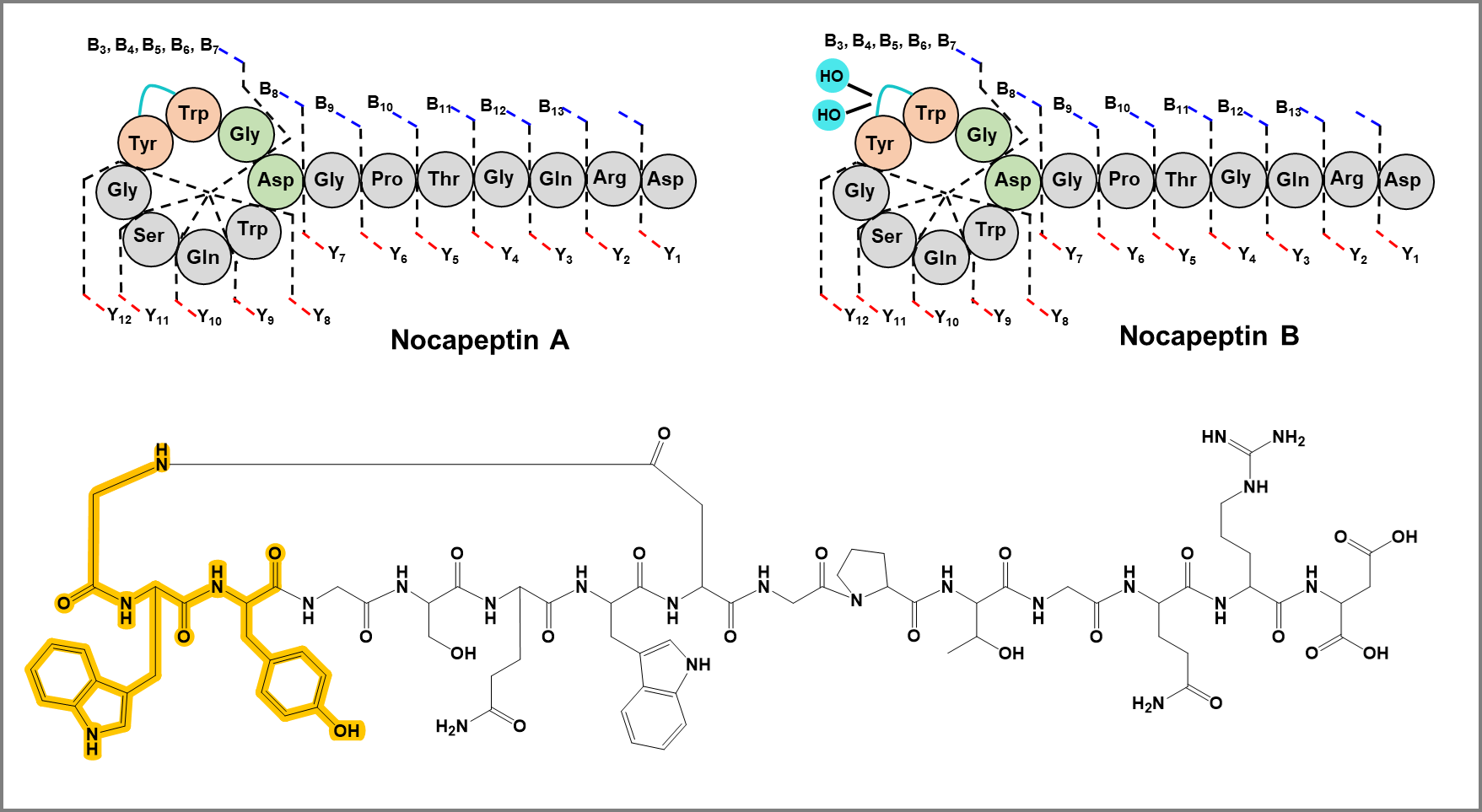

**Figure S4C**. Cartoon structures of nocapeptins A (**1**) and B (**2**) illustrating their annotated fragments. The substrcture highlighted in yellow was the presumed fragment to receive the tailoring.

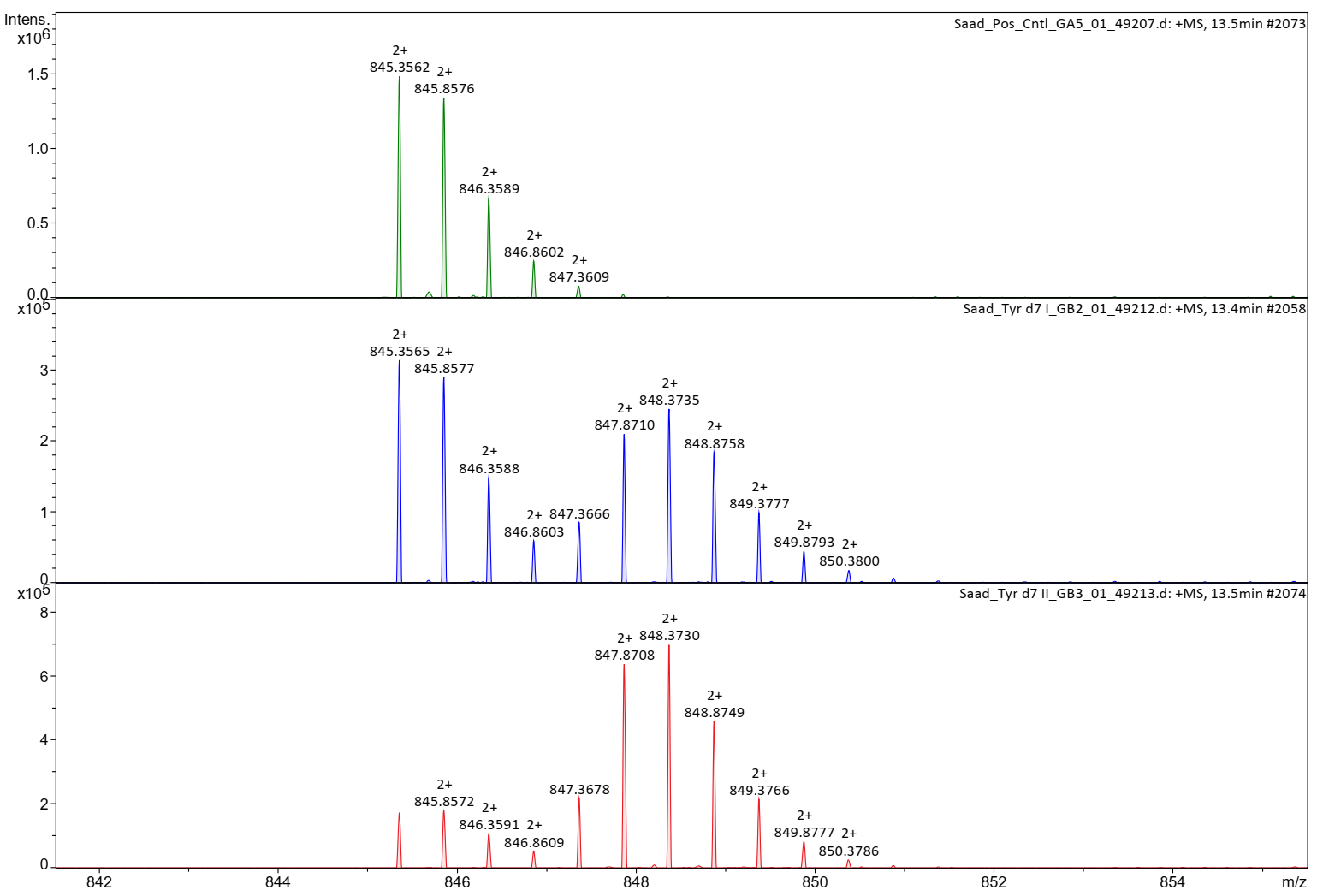

**Positive control**

**6**

**[^2^H_7_] Tyrosine, replicate 1**

**[^2^H_7_] Tyrosine, replicate 2**

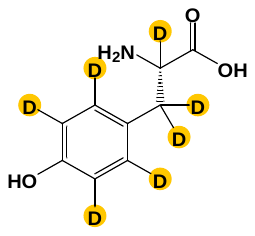

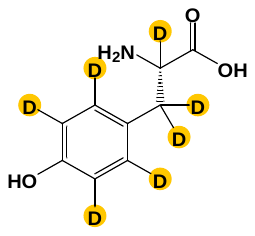

**Figure S5.** Comparative MS^1^ spectra of nocapeptin A (**1**) and its [^2^H_7_] L-tyrosine-labeled version

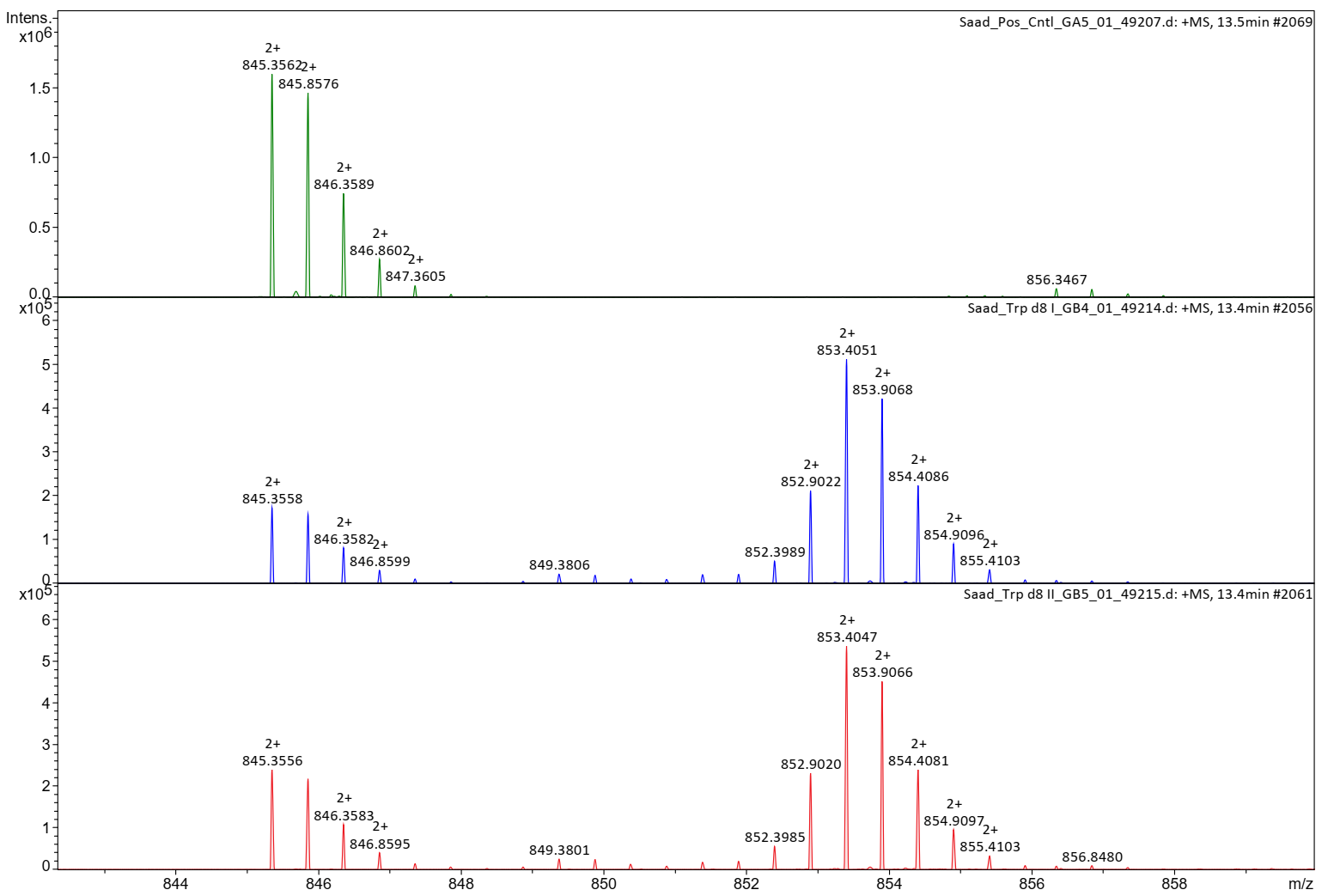

**Positive control**

**[^2^H_8_] Tryptophan, replicate 1**

**16**

**[^2^H_8_] Tryptophan, replicate 2**

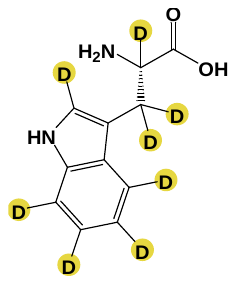

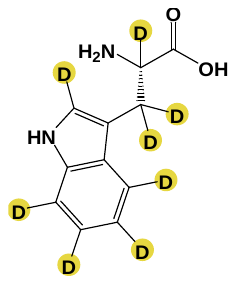

**Figure S6.** Comparative MS^1^ spectra of nocapeptin A (**1**) and its [^2^H_8_] L-tryptophan-labeled version

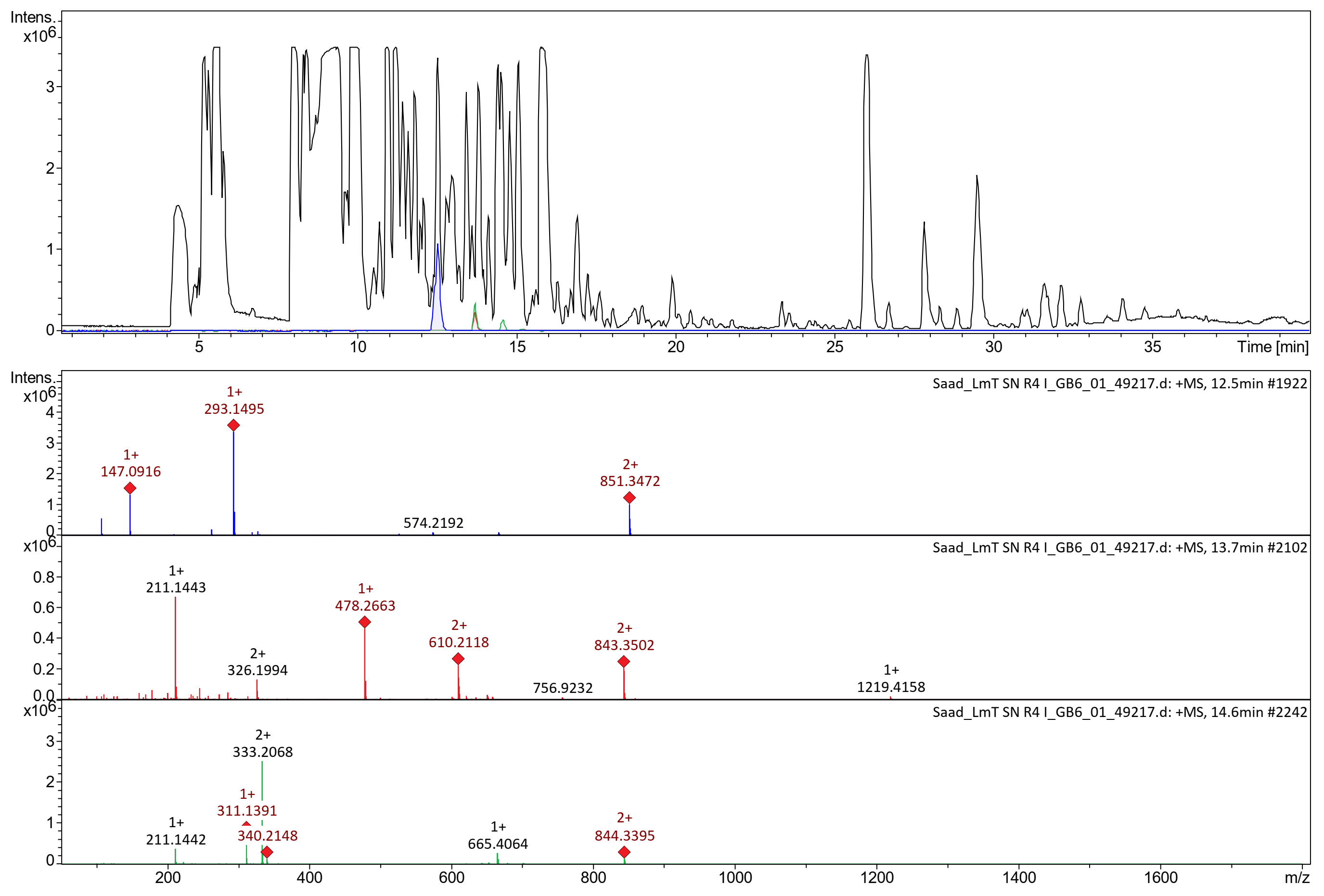

**Longipeptin A**

**Longipeptin B**

**Longipeptin C**

**Figure S7.** LCMS profile of modified R4 medium-based cultivation highlighting the production of longipeptin A (**3**), longipeptin B (**4**) and longipeptin C (**5**)

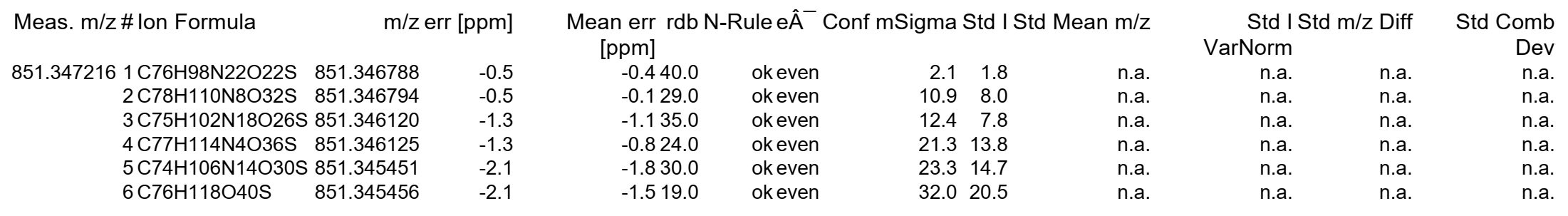

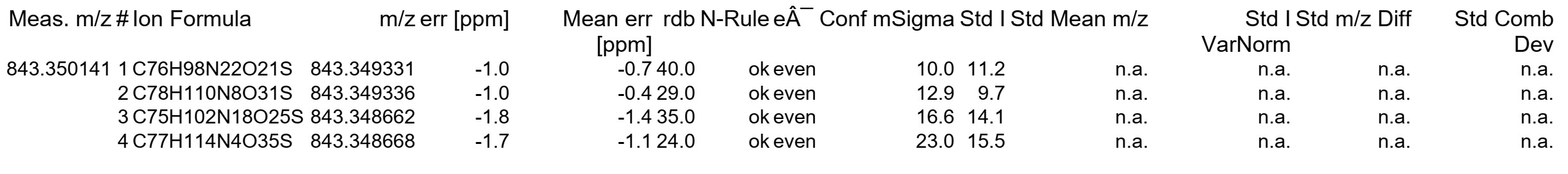

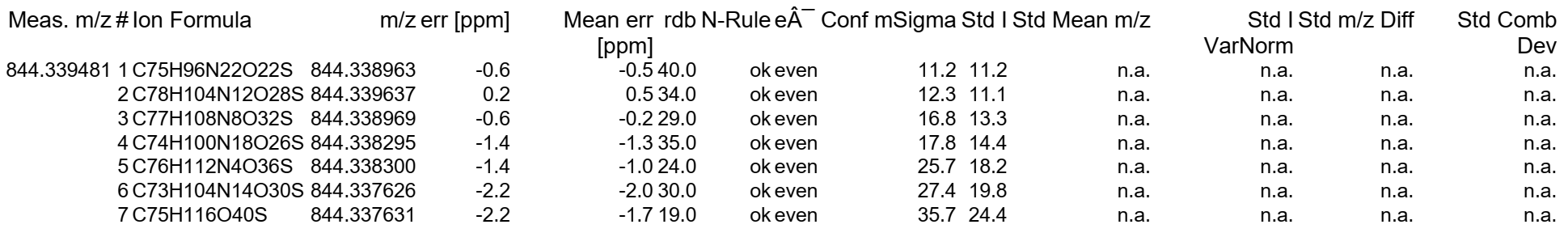

**Figure S7A.** Molecular formula predictions of longipeptin A (**3**) [851 Da], longipeptin B (**4**) [843 Da] and longipeptin C (**5**) [844 Da]

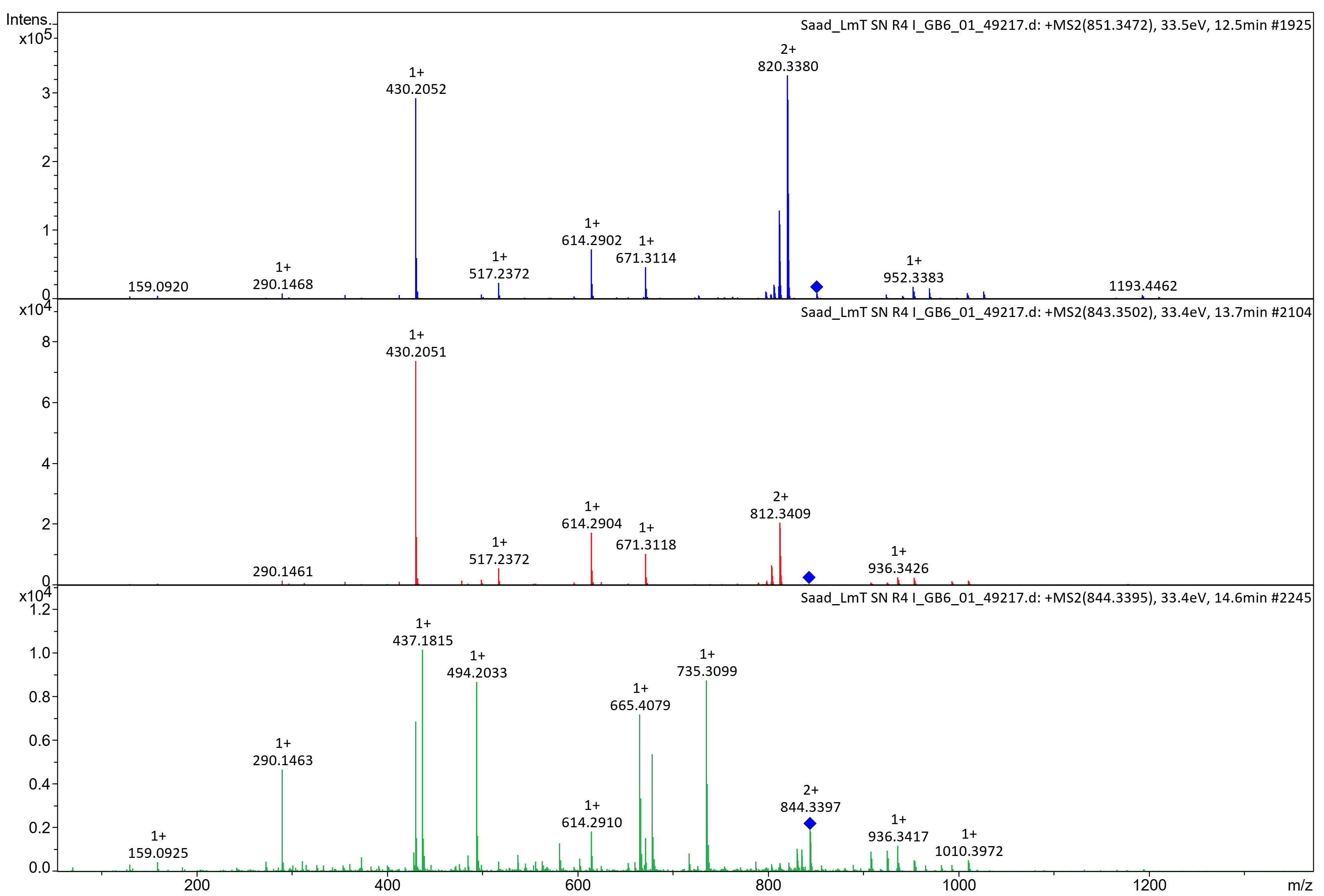

**Longipeptin C**

**Longipeptin B**

**Longipeptin A**

**Figure S8.** Comparative MS^2^ spectra of longipeptins A (**3**) [851 Da], B (**4**) [843 Da] and C (**5**) [844 Da]

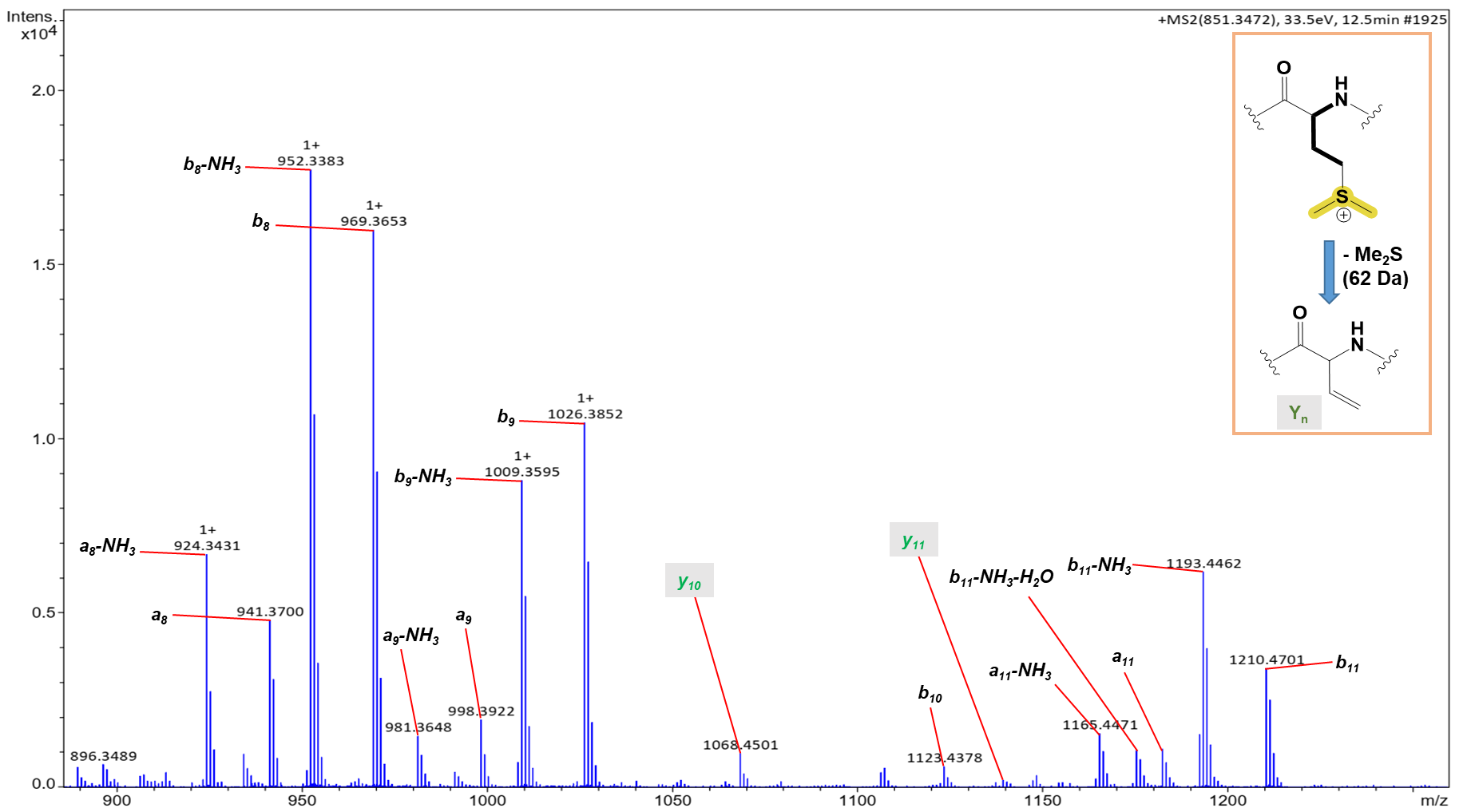

**Figure S9.** Annotated MS^2^ spectrum of longipeptin A (**3**) [851 Da]

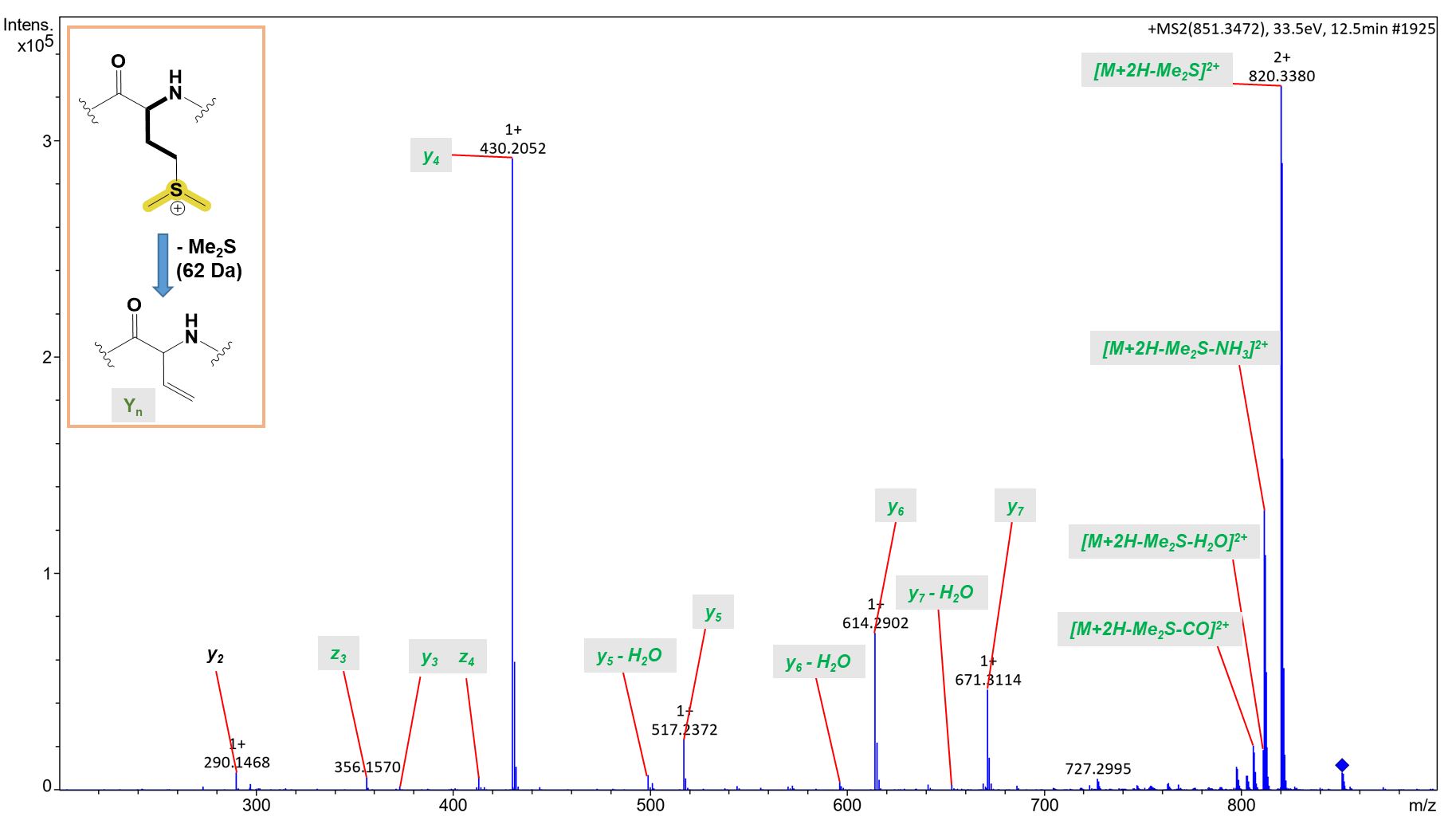

**Figure S9A.** Annotated MS^2^ spectrum of longipeptin A (**3**) [851 Da]

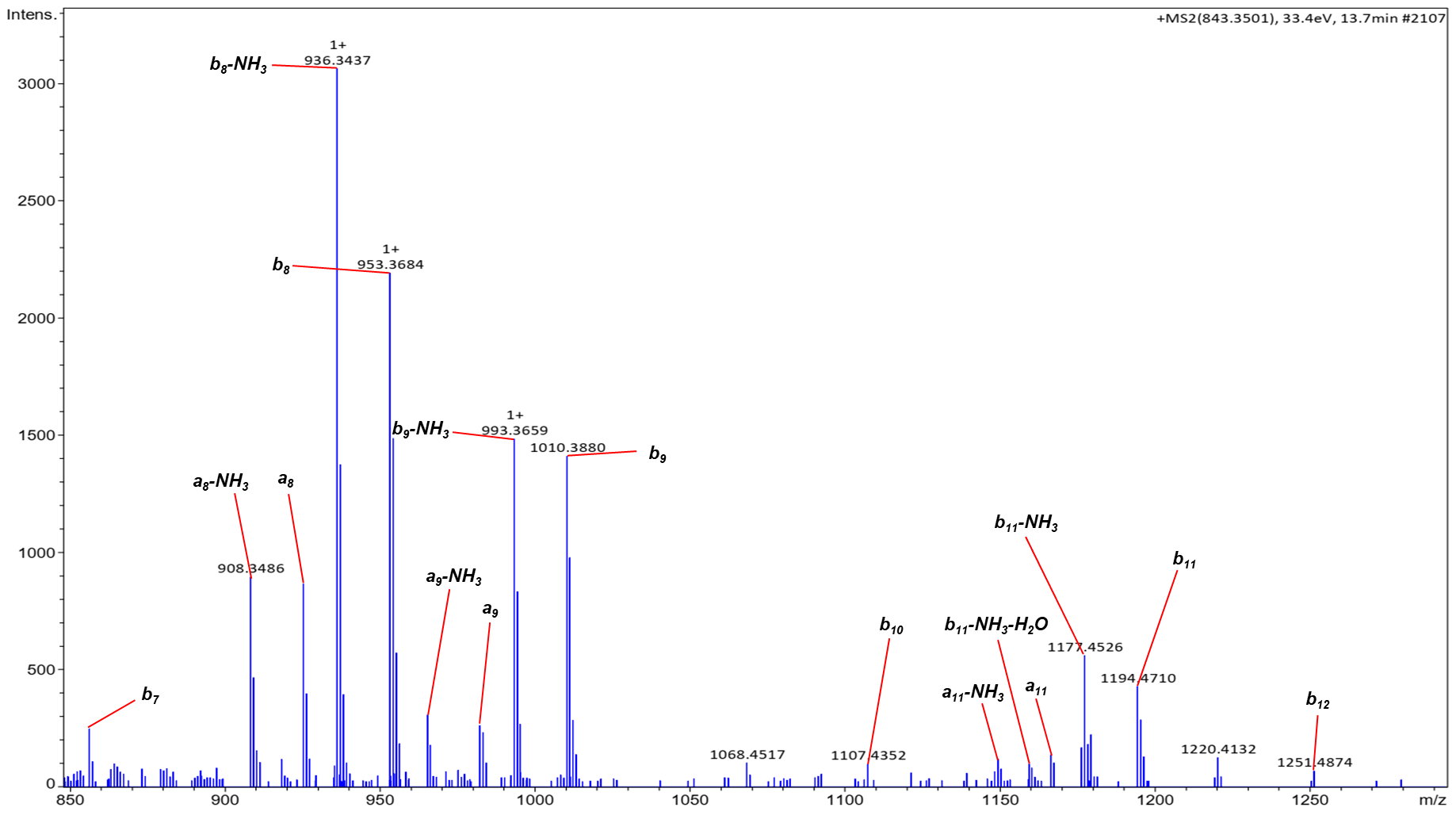

**Figure S10.** Annotated MS^2^ spectrum of longipeptin B (**4**) [843 Da]

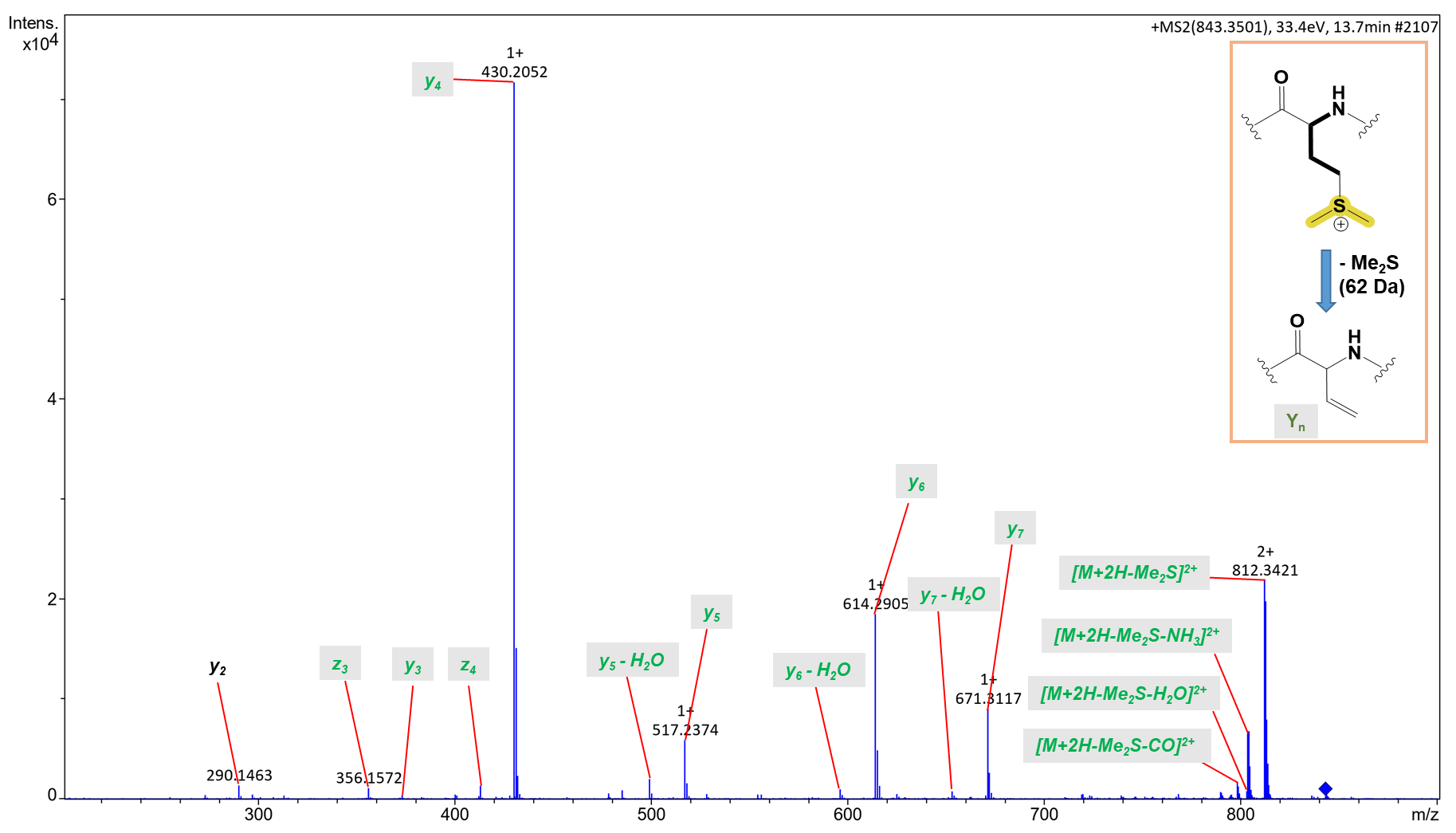

**Figure S10A.** Annotated MS^2^ spectrum of longipeptin B (**4**) [843 Da]

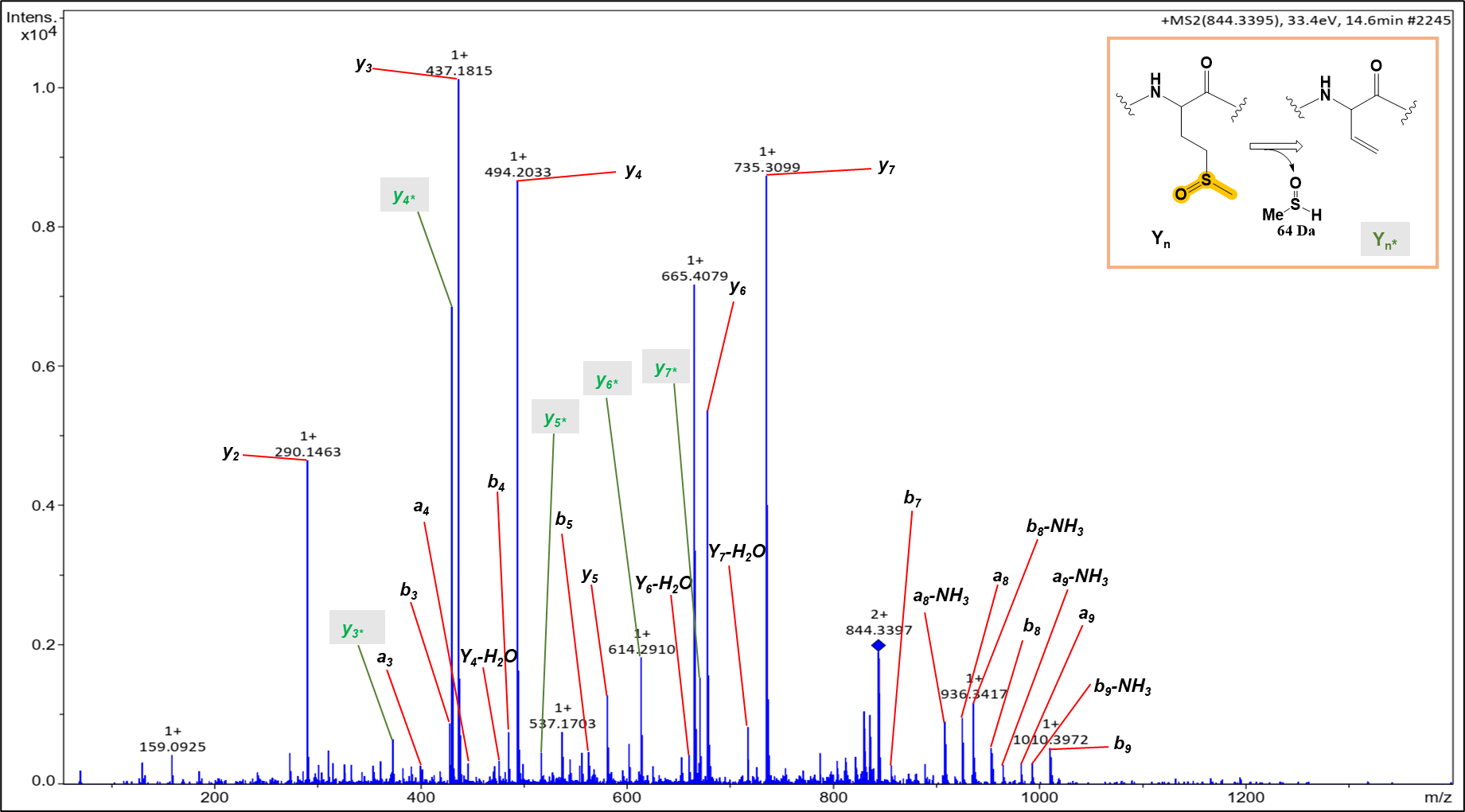

**Figure S11.** Annotated MS^2^ spectrum of longipeptin C (**5**) [844 Da]

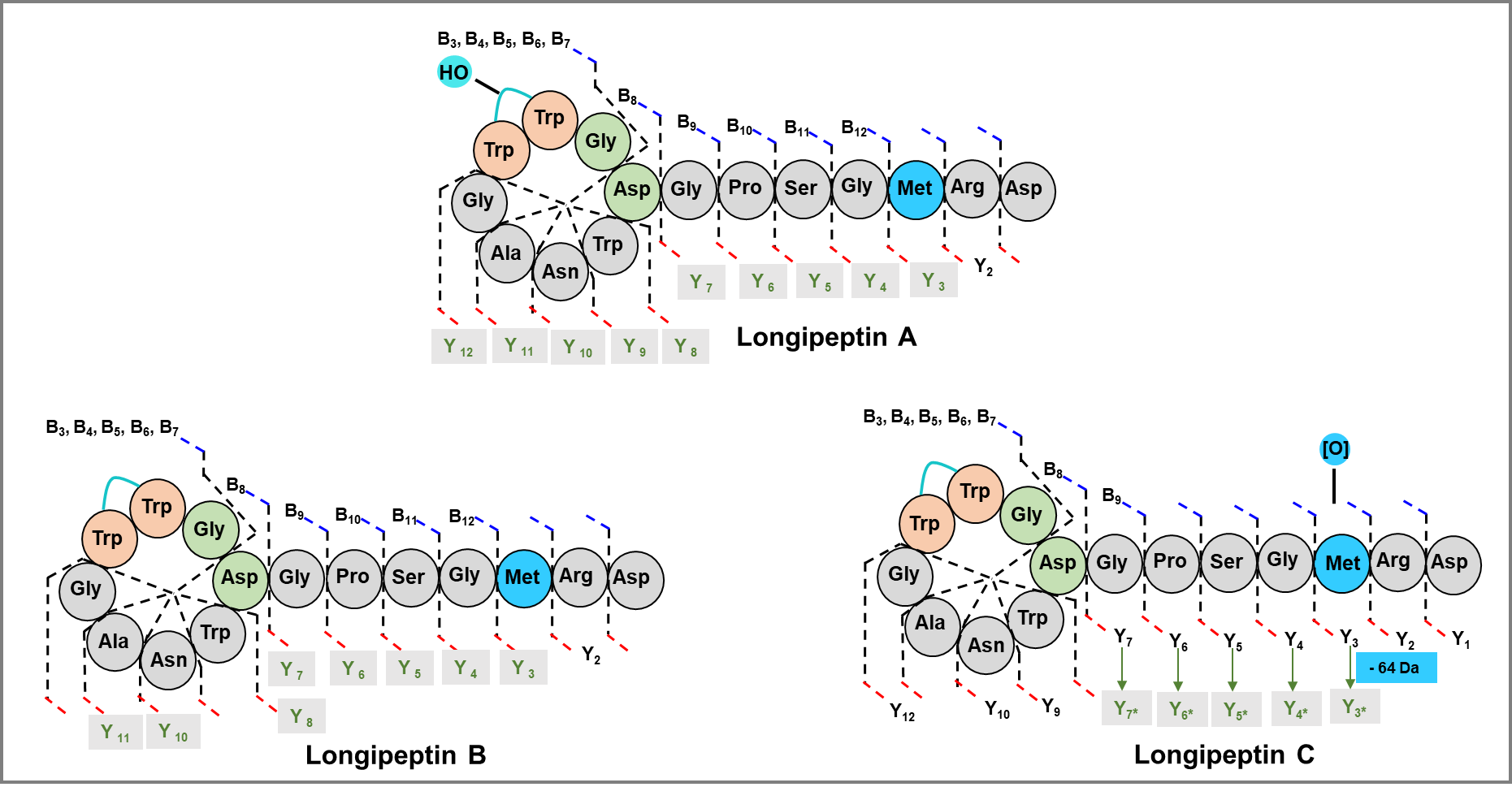

**Figure S12**. Cartoon structures of longipeptins A, B and C (**3**-**5**) illustrating their annotated fragments

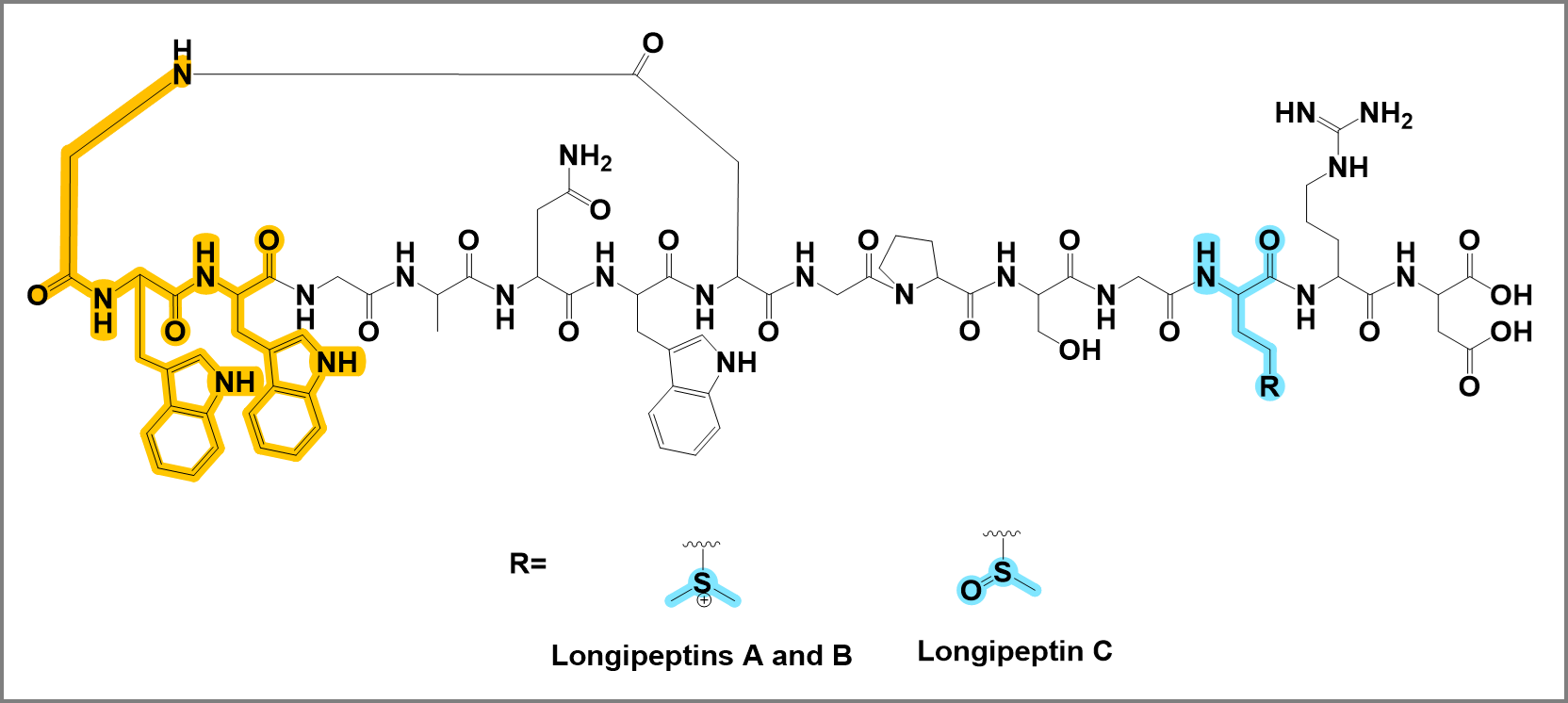

**Figure S13.** Detailed structures of longipeptins A, B and C (**3**-**5**) illustrating the different PTMs localities

**Table S2.** The assigned fragments of nocapeptin A (**1**) [845 Da], degree of unsaturation (RDB)

| Fragment | Ion Formula | RDB | Calc. *m/z* | Meas. *m/z* | Error in ppm | Sequence |
| --- | --- | --- | --- | --- | --- | --- |
| *y_1_* | C_4_H_8_NO_4_ | 2 | 134.0453 | 134.0450 | 2.24 | GSQ W DGPTGQRD |
| *y_2_* | C_10_H_20_N_5_O_5_ | 4 | 290.1464 | 290.1466 | 0.69 | GSQW DGPTGQRD |
| *z_2_* | C_10_H_17_N_4_O_5_ | 5 | 273.1199 | 273.1198 | 0.37 |  |
| *y_3_* | C_15_H_28_N_7_O_7_ | 6 | 418.2050 | 418.2053 | 0.72 | G SQWD GPTGQRD |
| *z_3_* | C_15_H_25_N_6_O_7_ | 7 | 401.1785 | 401.1784 | 0.25 |  |
| *y_4_* | C_17_H_31_N_8_O_8_ | 7 | 475.2265 | 475.2265 | 0.00 | GSQWD GPTGQRD |
| *z_4_* | C_17_H_28_N_7_O_8_ | 8 | 458.1999 | 458.2002 | 0.65 |  |
| *y_5_* | C_21_H_38_N_9_O_10_ | 8 | 576.2742 | 576.2741 | 0.17 | G SQW DGPTGQRD |
| *z_5_* | C_21_H_35_N_8_O_10_ | 9 | 559.2476 | 559.2482 | 1.07 |  |
| *y_6_* | C_26_H_45_N_10_O_11_ | 10 | 673.3269 | 673.3277 | 1.19 | G SQW DGPTGQRD |
| *z_6_* | C_26_H_42_N_9_O_11_ | 11 | 656.3004 | 656.3009 | 0.76 |  |
| *y_7_* | C_28_H_48_N_11_O_12_ | 11 | 730.3484 | 730.3490 | 0.82 | G SQ WDGPTGQRD |
| *z_7_* | C_28_H_45_N_10_O_12_ | 12 | 713.3218 | 713.3223 | 0.70 |  |
| *y_8_* | C_32_H_51_N_12_O_14_ | 14 | 827.3648 | --------- | --------- | GS QW**D**GPTGQRD |
| *y_9_* | C_43_H_61_N_14_O_15_ | 21 | 1013.4441 | 1013.4434 | 0.69 | G S QW**D**GPTGQRD |
| *z_9_* | C_43_H_58_N_13_O_15_ | 22 | 996.4175 | 996.4169 | 0.60 |  |
| *y_10_* | C_48_H_69_N_16_O_17_ | 23 | 1141.5027 | 1141.5036 | 0.79 | G SQW**D**GPTGQRD |
| *y_11_* | C_51_H_74_N_17_O_19_ | 24 | 1228.5347 | --------- | --------- |  |
| *y_12_* | C_53_H_77_N_18_O_20_ | 25 | 1285.5562 | 1285.5639 | 5.98 | GSQW**D**GPTGQRD |
| *z_12_* | C_53_H_74_N_17_O_20_ | 26 | 1268.5296 | 1268.5147 | 11.74 |  |
| *y_13_* | --------- | --- | --------- | --------- | --------- |  |
| *y_14_* | --------- | --- | --------- | --------- | --------- |  |
| *b_1_* | --------- | --- | --------- | --------- | --------- |  |
| *b_2_* | --------- | --- | --------- | --------- | --------- |  |
| *b_3_* | C_22_H_21_N_4_O_4_ | 15 | 405.1563 | 405.1562 | 0.25 | **G**WY |
| *a_3_* | C_21_H_21_N_4_O_3_ | 14 | 377.1614 | 377.1610 | 1.06 |  |
| *b_4_* | C_24_H_24_N_5_O_5_ | 16 | 462.1777 | 462.1778 | 0.22 | **G**WYG |
| *a_4_* | C_23_H_24_N_5_O_4_ | 15 | 434.1828 | 434.1842 | 3.22 |  |
| *b_5_* | C_27_H_29_N_6_O_7_ | 17 | 549.2098 | 549.2099 | 0.18 | **G**WYGS |
| *a_5_* | C_26_H_29_N_6_O_6_ | 16 | 521.2149 | 521.2142 | 1.34 |  |
| *b_6_* | C_32_H_37_N_8_O_9_ | 19 | 677.2683 | 677.2675 | 1.18 | **G**WYGSQ |
| *a_6_* | C_31_H_37_N_8_O_8_ | 18 | 649.2734 | 649.2746 | 1.84 |  |
| *b_7_* | C_43_H_47_N_10_O_10_ | 26 | 863.3477 | 863.3494 | 1.97 | **G**WYGSQW |
| *b_8_* | C_47_H_50_N_11_O_12_ | 29 | 960.3640 | 960.3641 | 0.10 | **G**WYGSQW**D** |
| *a_8_* | C_46_H_50_N_11_O_11_ | 28 | 932.3691 | 932.3705 | 1.50 |  |
| *b_9_* | C_49_H_53_N_12_O_13_ | 30 | 1017.3855 | 1017.3865 | 0.98 | **G**WYGSQW**D**G |
| *a_9_* | C_48_H_53_N_12_O_12_ | 29 | 989.3906 | 989.3904 | 0.20 |  |
| *b_10_* | C_54_H_60_N_13_O_14_ | 32 | 1114.4383 | 1114.4405 | 1.97 | **G**WYGSQW**D**GP |
| *a_10_* | C_53_H_60_N_13_O_13_ | 31 | 1086.4434 | 1086.4480 | 4.23 |  |
| *b_11_* | C_58_H_67_N_14_O_16_ | 33 | 1215.4859 | 1215.4860 | 0.08 | **G**WYGSQW**D**GPT |
| *a_11_* | C_57_H_67_N_14_O_15_ | 32 | 1187.4910 | 1187.4883 | 2.27 |  |
| *b_12_* | C_60_H_70_N_15_O_17_ | 34 | 1272.5074 | 1272.5056 | 1.41 | **G**WYGSQW**D**GPTG |
| *a_12_* | C_59_H_70_N_15_O_16_ | 33 | 1244.5125 | 1244.5106 | 1.53 |  |
| *b_13_* | C_65_H_78_N_17_O_19_ | 36 | 1400.5660 | 1400.5628 | 2.28 | **G**WYGSQW**D**GPTGQ |
| *a_13_* | C_64_H_78_N_17_O_18_ | 35 | 1372.5711 | 1372.5808 | 7.07 |  |
| *b_14_* | C_71_H_90_N_21_O_20_ | 38 | 1556.6671 | --------- | --------- |  |

**Table S3**. The assigned fragments of nocapeptin B (**2**) [861 Da], degree of unsaturation (RDB)

| Fragment | Ion Formula | RDB | Calc. *m/z* | Meas. *m/z* | error in ppm | Sequence |
| --- | --- | --- | --- | --- | --- | --- |
| *y_1_* | C_10_H_20_N_5_O_5_ | 2 | 134.0453 | 134.0450 | 2.24 | GSQW D GPTGQRD |
| *y_2_* | C_10_H_20_N_5_O_5_ | 4 | 290.1464 | 290.1469 | 1.72 | GSQ W DGPTGQRD |
| *z_2_* | C_10_H_17_N_4_O_5_ | 5 | 273.1199 | 273.1207 | 2.93 |  |
| *y_3_* | C_15_H_28_N_7_O_7_ | 6 | 418.2050 | 418.2057 | 1.67 | GSQ W DGPTGQRD |
| *z_3_* | C_15_H_25_N_6_O_7_ | 7 | 401.1785 | 401.1795 | 2.49 |  |
| *y_4_* | C_17_H_31_N_8_O_8_ | 7 | 475.2265 | 475.2273 | 1.68 | GSQ W DGPTGQRD |
| *z_4_* | C_17_H_28_N_7_O_8_ | 8 | 458.1999 | 458.2005 | 1.31 |  |
| *y_5_* | C_21_H_38_N_9_O_10_ | 8 | 576.2742 | 576.2748 | 1.04 | GSQ W DGPTGQRD |
| *z_5_* | C_21_H_35_N_8_O_10_ | 9 | 559.2476 | 559.2488 | 2.15 |  |
| *y_6_* | C_26_H_45_N_10_O_11_ | 10 | 673.3269 | 673.3285 | 2.38 | GSQ W DGPTGQRD |
| *z_6_* | C_26_H_42_N_9_O_11_ | 11 | 656.3004 | 656.3018 | 2.13 |  |
| *y_7_* | C_28_H_48_N_11_O_12_ | 11 | 730.3484 | 730.3495 | 1.50 | GS Q WDGPTGQRD |
| *z_7_* | C_28_H_45_N_10_O_12_ | 12 | 713.3218 | 713.3232 | 1.96 |  |
| *y_8_* | C_32_H_51_N_12_O_14_ | 14 | 827.3648 | 827.3652 | 0.48 | GS Q W**D**GPTGQRD |
| *y_9_* | C_43_H_61_N_14_O_15_ | 21 | 1013.4441 | 1013.4437 | 0.39 | GS QW**D**GPTGQRD |
| *z_9_* | C_43_H_58_N_13_O_15_ | 22 | 996.4175 | 996.4167 | 0.80 |  |
| *y_10_* | C_48_H_69_N_16_O_17_ | 23 | 1141.5027 | 1141.4966 | 5.34 | G SQW**D**GPTGQRD |
| *y_11_* | C_51_H_74_N_17_O_19_ | 24 | 1228.5347 | --------- | --------- | G |
| *y_12_* | C_53_H_77_N_18_O_20_ | 25 | 1285.5562 | 1285.5593 | 2.41 | GSQW**D**GPTGQRD |
| *y_13_* | --------- | --- | --------- | --------- | --------- |  |
| *y_14_* | --------- | --- | --------- | --------- | --------- |  |
| *b_1_* | --------- | --- | --------- | --------- | --------- |  |
| *b_2_* | --------- | --- | --------- | --------- | --------- |  |
| *b_3_* | C_22_H_21_N_4_O_6_ | 15 | 437.1461 | 437.1465 | 0.92 | **G**WYGSQWDGPTGQR |
| *a_3_* | C_21_H_21_N_4_O_5_ | 14 | 409.1512 | 409.1508 | 0.98 |  |
| *b_4_* | C_24_H_24_N_5_O_7_ | 16 | 494.1676 | 494.1684 | 1.62 | **G**WYGSQWDGPTGQR |
| *b_5_* | C_27_H_29_N_6_O_9_ | 17 | 581.1996 | 581.2012 | 2.75 | **G**WYGSQWDGPTGQR |
| *a_5_* | C_26_H_29_N_6_O_8_ | 16 | 553.2047 | 553.2033 | 2.53 |  |
| *b_6_* | C_32_H_37_N_8_O_11_ | 19 | 709.2582 | 709.2588 | 0.85 | **G**WYGSQWDGPTGQR |
| *a_6_* | C_31_H_37_N_8_O_10_ | 18 | 681.2633 | 681.2683 | 7.34 |  |
| *b_7_* | C_43_H_47_N_10_O_12_ | 26 | 895.3375 | 895.3382 | 0.78 | **G**WYGSQWDGPTGQR |
| *a_7_* | C_42_H_47_N_10_O_11_ | 25 | 867.3426 | 867.3448 | 2.54 |  |
| *b_8_* | C_47_H_50_N_11_O_14_ | 29 | 992.3539 | 992.3546 | 0.71 | **G**WYGSQW**D**GPTGQR |
| *a_8_* | C_46_H_50_N_11_O_13_ | 28 | 964.3590 | 964.3606 | 1.66 |  |
| *b_9_* | C_49_H_53_N_12_O_15_ | 30 | 1049.3753 | 1049.3776 | 2.19 | **G**WYGSQW**D**GPTGQR |
| *a_9_* | C_48_H_53_N_12_O_14_ | 29 | 1021.3804 | 1021.3828 | 2.35 |  |
| *b_10_* | C_54_H_60_N_13_O_16_ | 32 | 1146.4281 | 1146.4279 | 0.17 | **G**WYGSQW**D**GPTGQR |
| *a_10_* | C_53_H_60_N_13_O_15_ | 31 | 1118.4332 | 1118.4271 | 5.45 |  |
| *b_11_* | C_58_H_67_N_14_O_18_ | 33 | 1247.4758 | 1247.4776 | 1.44 | **G**WYGSQW**D**GPTGQR |
| *a_11_* | C_57_H_67_N_14_O_17_ | 32 | 1219.4809 | 1219.4809 | 0.00 |  |
| *b_12_* | C_60_H_70_N_15_O_19_ | 34 | 1304.4972 | 1304.4935 | 2.84 | **G**WYGSQW**D**GPTGQR |
| *a_12_* | C_59_H_70_N_15_O_18_ | 33 | 1276.5023 | 1276.4985 | 2.98 |  |
| *b_13_* | C_65_H_78_N_17_O_21_ | 36 | 1432.5558 | 1432.5660 | 7.12 | **G**WYGSQW**D**GPTGQR |
| *b_14_* | C_71_H_90_N_21_O_22_ | 38 | 1588.6569 | --------- | --------- |  |

**Table S4**. The assigned fragments of longipeptin A (**3**) [851 Da]; *Y*, *Z*, *Y-H_2_O* indicate ions upon Me_2_S (62 Da) loss, degree of unsaturation (RDB)

| Fragment | Ion Formula | RDB | Calc. *m/z* | Meas. *m/z* | error in ppm | Sequence |
| --- | --- | --- | --- | --- | --- | --- |
| *y_1_* | C_4_H_8_NO_4_ | 2 | 134.0453 | --------- | --------- | GAN WDGPSGMR |
| *y_2_* | C_10_H_20_N_5_O_5_ | 4 | 290.1464 | 290.1468 | 1.37 | GA NWDGPSGMRD |
| *y_3_* | C_14_H_25_N_6_O_6_ | 6 | 373.1836 | 373.1837 | 0.27 | GA NWDGPSGMRD |
| *z_3_* | C_14_H_22_N_5_O_6_ | 7 | 356.1570 | 356.1570 | 0.00 |  |
| *y_4_* | C_16_H_28_N_7_O_7_ | 7 | 430.2050 | 430.2052 | 0.46 | GA NWDGPSGMRD |
| *z_4_* | C_16_H_25_N_6_O_7_ | 8 | 413.1785 | 413.1783 | 0.48 |  |
| *y_5_* | C_19_H_33_N_8_O_9_ | 8 | 517.2370 | 517.2372 | 0.39 | GA NWDGPSGMRD |
| *y_5_-H_2_O* | C_19_H_31_N_8_O_8_ | 9 | 499.2265 | 499.2265 | 0.00 |  |
| *y_6_* | C_24_H_40_N_9_O_10_ | 10 | 614.2898 | 614.2902 | 0.65 | GA NWDGPSGMRD |
| *y_6_-H_2_O* | C_24_H_38_N_9_O_9_ | 11 | 596.2792 | 596.2786 | 1.00 |  |
| *y_7_* | C_26_H_43_N_10_O_11_ | 11 | 671.3113 | 671.3114 | 0.15 | GA NWDGPSGMRD |
| *y_7_-H_2_O* | C_26_H_41_N_10_O_10_ | 12 | 653.3007 | 653.3015 | 1.22 |  |
| *y_8_* | C_30_H_46_N_11_O_13_ | 14 | 768.3277 | 768.3240 | 4.81 | G ANW**D**GPSGMRD |
| *y_9_* | C_41_H_56_N_13_O_14_ | 21 | 954.4070 | 954.4048 | 2.30 | GANW**D**GPSGMRD |
| *y_10_* | C_45_H_62_N_15_O_16_ | 23 | 1068.4499 | 1068.4501 | 0.19 | G ANW**D**GPSGMRD |
| *y_11_* | C_48_H_67_N_16_O_17_ | 24 | 1139.4870 | 1139.4801 | 6.05 | G ANW**D**GPSGMRD |
| *y_12_* | C_50_H_70_N_17_O_18_ | 25 | 1196.5085 | 1196.5090 | 0.42 | GANW**D**GPSGMRD |
| *y_13_* | --------- | --- | --------- | --------- | --------- |  |
| *y_14_* | --------- | --- | --------- | --------- | --------- |  |
| *b_1_* | --------- | --- | --------- | --------- | --------- |  |
| *b_2_* | --------- | --- | --------- | --------- | --------- |  |
| *b_3_* | C_24_H_22_N_5_O_4_ | 17 | 444.1672 | 444.1681 | 2.03 | **G**WWGANWDGPSG |
| *a_3_* | C_23_H_22_N_5_O_3_ | 16 | 416.1723 | 416.1728 | 1.20 |  |
| *b_4_* | C_26_H_25_N_6_O_5_ | 18 | 501.1886 | 501.1892 | 1.20 | **G**WWGANWDGPSG |
| *a_4_* | C_25_H_25_N_6_O_4_ | 17 | 473.1937 | 473.1945 | 1.70 |  |
| *b_5_* | C_29_H_30_N_7_O_6_ | 19 | 572.2258 | 572.2252 | 1.05 | **G**WWGANWDGPSG |
| *a_5_* | C_28_H_30_N_7_O_5_ | 18 | 544.2308 | 544.2306 | 0.37 |  |
| *b_6_* | C_33_H_36_N_9_O_8_ | 21 | 686.2687 | 686.2701 | 2.04 | **G**WWGANWDGPSG |
| *b_6_-NH_3_* | C_33_H_33_N_8_O_8_ | 22 | 669.2421 | 669.2424 | 0.45 |  |
| *a_6_* | C_32_H_36_N_9_O_7_ | 20 | 658.2738 | 658.2715 | 3.50 |  |
| *a_6_-NH_3_* | C_32_H_33_N_8_O_7_ | 21 | 641.2472 | 641.2470 | 0.31 |  |
| *b_7_* | C_44_H_46_N_11_O_9_ | 28 | 872.3480 | 872.3483 | 0.34 | **G**WWGANWDGPSG |
| *b_8_* | C_48_H_49_N_12_O_11_ | 31 | 969.3644 | 969.3653 | 0.93 | **G**WWGANW**D**GPSG |
| *b_8_-NH_3_* | C_48_H_46_N_11_O_11_ | 32 | 952.3378 | 952.3383 | 0.53 |  |
| *a_8_* | C_47_H_49_N_12_O_10_ | 30 | 941.3695 | 941.3700 | 0.53 |  |
| *a_8_-NH_3_* | C_47_H_46_N_11_O_10_ | 31 | 924.3429 | 924.3431 | 0.22 |  |
| *b_9_* | C_50_H_52_N_13_O_12_ | 32 | 1026.3858 | 1026.3852 | 0.58 | **G**WWGANW**D**GPSG |
| *b_9_-NH_3_* | C_50_H_49_N_12_O_12_ | 33 | 1009.3593 | 1009.3595 | 0.20 |  |
| *a_9_* | C_49_H_52_N_13_O_11_ | 31 | 998.3909 | 998.3922 | 1.30 |  |
| *a_9_-NH_3_* | C_49_H_49_N_12_O_11_ | 32 | 981.3644 | 981.3648 | 0.41 |  |
| *b_10_* | C_55_H_59_N_14_O_13_ | 34 | 1123.4386 | 1123.4378 | 0.71 | **G**WWGANW**D**GPSG |
| *b_11_* | C_58_H_64_N_15_O_15_ | 35 | 1210.4706 | 1210.4701 | 0.41 | **G**WWGANW**D**GPSG |
| *b_11_-NH_3_* | C_58_H_61_N_14_O_15_ | 36 | 1193.4441 | 1193.4462 | 1.76 |  |
| *b_11_-NH_3_-H_2_O* | C_58_H_59_N_14_O_14_ | 37 | 1175.4335 | 1175.4315 | 1.70 |  |
| *a_11_* | C_57_H_64_N_15_O_14_ | 34 | 1182.4757 | 1182.4743 | 1.18 |  |
| *a_11_-NH_3_* | C_57_H_61_N_14_O_14_ | 35 | 1165.4492 | 1165.4471 | 1.80 |  |
| *b_12_* | C_60_H_67_N_16_O_16_ | 36 | 1267.4921 | 1267.5161 | 18.94 | **G**WWGANW**D**GPSG |
| *a_12_* | C_59_H_67_N_16_O_15_ | 35 | 1239.4972 | 1239.5103 | 10.56 |  |
| *[M +H – Me_2_S]^2+^* | C_74_H_92_N_22_O_22_ | 40 | 820.3379 | 820.3380 | 0.12 | **G**WWGANW**D**GPSGMRD |
| *[M +H – Me_2_S – NH_3_]^2+^* | C_74_H_89_N_21_O_22_ | 41 | 811.8246 | 811.8268 | 2.70 | **G**WWGANW**D**GPSGMRD |
| *[M +H – Me_2_S– H_2_O]^2+^* | C_74_H_90_N_22_O_21_ | 41 | 811.3326 | 811.3340 | 1.70 | **G**WWGANW**D**GPSGMRD |
| *[M +H – Me_2_S – CO]^2+^* | C_73_H_92_N_22_O_21_ | 39 | 806.3404 | 806.3415 | 1.36 | **G**WWGANW**D**GPSGMRD |

**Table S5**. The assigned fragments of longipeptin B (**4**) [843 Da]; *Y*, *Z*, *Y-H_2_O* indicate ions upon Me_2_S (62 Da) loss, degree of unsaturation (RDB)

| Fragment | Ion Formula | RDB | Calc. *m/z* | Meas. *m/z* | error in ppm | Sequence |
| --- | --- | --- | --- | --- | --- | --- |
| *y_1_* | C_4_H_8_NO_4_ | 2 | 134.0453 | --------- | --------- | GAN WDGPSGM |
| *y_2_* | C_10_H_20_N_5_O_5_ | 4 | 290.1464 | 290.1463 | 0.34 | GAN WDGPSGMRD |
| *y_3_* | C_14_H_25_N_6_O_6_ | 6 | 373.1836 | 373.1830 | 1.61 | GAN WDGPSGMRD |
| *z_3_* | C_14_H_22_N_5_O_6_ | 7 | 356.1570 | 356.1572 | 0.56 |  |
| *y_4_* | C_16_H_28_N_7_O_7_ | 7 | 430.2050 | 430.2052 | 0.46 | GANW DGPSGMRD |
| *z_4_* | C_16_H_25_N_6_O_7_ | 8 | 413.1785 | 413.1782 | 0.73 |  |
| *y_5_* | C_19_H_33_N_8_O_9_ | 8 | 517.2370 | 517.2374 | 0.77 | GANW DGPSGMRD |
| *y_5_-H_2_O* | C_19_H_31_N_8_O_8_ | 9 | 499.2265 | 499.2267 | 0.40 |  |
| *y_6_* | C_24_H_40_N_9_O_10_ | 10 | 614.2898 | 614.2905 | 1.14 | GAN WDGPSGMRD |
| *y_6_-H_2_O* | C_24_H_38_N_9_O_9_ | 11 | 596.2792 | 596.2788 | 0.67 |  |
| *y_7_* | C_26_H_43_N_10_O_11_ | 11 | 671.3113 | 671.3117 | 0.60 | GAN WDGPSGMRD |
| *y_7_-H_2_O* | C_26_H_41_N_10_O_10_ | 12 | 653.3007 | 653.3009 | 0.31 |  |
| *y_8_* | C_30_H_46_N_11_O_13_ | 14 | 768.3277 | 768.3253 | 3.12 | G ANW**D**GPSGMRD |
| *y_9_* | C_41_H_56_N_13_O_14_ | 21 | 954.4070 | --------- | --------- | G ANW**D**GPSGMRD |
| *y_10_* | C_45_H_62_N_15_O_16_ | 23 | 1068.4499 | 1068.4517 | 1.68 | G ANW**D**GPSGMRD |
| *y_11_* | C_48_H_67_N_16_O_17_ | 24 | 1139.4870 | 1139.4790 | 7.02 | GANW**D**GPSGMRD |
| *y_12_* | C_50_H_70_N_17_O_18_ | 25 | 1196.5085 | --------- | --------- |  |
| *y_13_* | --------- | --- | --------- | --------- | --------- |  |
| *y_14_* | --------- | --- | --------- | --------- | --------- |  |
| *b_1_* | --------- | --- | --------- | --------- | --------- |  |
| *b_2_* | --------- | --- | --------- | --------- | --------- |  |
| *b_3_* | C_24_H_22_N_5_O_3_ | 17 | 428.1723 | 428.1730 | 1.63 | **G**WWGANWDGPSG |
| *a_3_* | C_23_H_22_N_5_O_2_ | 16 | 400.1773 | 400.1775 | 0.50 |  |
| *b_4_* | C_26_H_25_N_6_O_4_ | 18 | 485.1937 | 485.1936 | 0.21 | **G**WWGANWDGPSG |
| *a_4_* | C_25_H_25_N_6_O_3_ | 17 | 457.1988 | 457.1994 | 1.31 |  |
| *b_5_* | C_29_H_30_N_7_O_5_ | 19 | 556.2308 | 556.2309 | 0.18 | **G**WWGANWDGPSG |
| *a_5_* | C_28_H_30_N_7_O_4_ | 18 | 528.2359 | 528.2360 | 0.19 |  |
| *b_6_* | C_33_H_36_N_9_O_7_ | 21 | 670.2738 | 670.2753 | 2.24 | **G**WWGANWDGPSG |
| *b_7_* | C_44_H_46_N_11_O_8_ | 28 | 856.3531 | 856.3499 | 3.73 | **G**WWGANWDGPSG |
| *b_8_* | C_48_H_49_N_12_O_10_ | 31 | 953.3695 | 953.3684 | 1.15 | **G**WWGANW**D**GPSG |
| *b_8_-NH_3_* | C_48_H_46_N_11_O_10_ | 32 | 936.3429 | 936.3437 | 0.85 |  |
| *a_8_* | C_47_H_49_N_12_O_9_ | 30 | 925.3745 | 925.3723 | 2.38 |  |
| *a_8_-NH_3_* | C_47_H_46_N_11_O_9_ | 31 | 908.3480 | 908.3486 | 0.67 |  |
| *b_9_* | C_50_H_52_N_13_O_11_ | 32 | 1010.3909 | 1010.3880 | 2.87 | **G**WWGANW**D**GPSG |
| *b_9_-NH_3_* | C_50_H_49_N_12_O_11_ | 33 | 993.3644 | 993.3659 | 1.51 |  |
| *a_9_* | C_49_H_52_N_13_O_10_ | 31 | 982.3960 | 982.4099 | 14.15 |  |
| *a_9_-NH_3_* | C_49_H_49_N_12_O_10_ | 32 | 965.3695 | 965.3748 | 5.49 |  |
| *b_10_* | C_55_H_59_N_14_O_12_ | 34 | 1107.4437 | 1107.4352 | 7.68 | **G**WWGANW**D**GPSG |
| *b_11_* | C_58_H_64_N_15_O_14_ | 35 | 1194.4757 | 1194.4710 | 3.93 | **G**WWGANW**D**GPSG |
| *b_11_-NH_3_* | C_58_H_61_N_14_O_14_ | 36 | 1177.4492 | 1177.4526 | 2.89 |  |
| *b_11_-NH_3_-H_2_O* | C_58_H_59_N_14_O_13_ | 37 | 1159.4386 | 1159.4420 | 2.93 |  |
| *a_11_* | C_57_H_64_N_15_O_13_ | 34 | 1166.4808 | 1166.4720 | 7.54 |  |
| *a_11_-NH_3_* | C_57_H_61_N_14_O_13_ | 35 | 1149.4543 | 1149.4795 | 21.92 |  |
| *b_12_* | C_60_H_67_N_16_O_15_ | 36 | 1251.4972 | 1251.4874 | 7.83 | **G**WWGANW**D**GPSG |
| *[M +H – Me_2_S]^2+^* | C_74_H_92_N_22_O_21_ | 40 | 812.3404 | 812.3421 | 2.09 | **G**WWGANW**D**GPSGMRD |
| *[M+H – Me_2_S – NH_3_]^2+^* | C_74_H_89_N_21_O_21_ | 41 | 803.8271 | 803.8285 | 1.74 | **G**WWGANW**D**GPSGMRD |
| *[M +H – Me_2_S– H_2_O]^2+^* | C_74_H_90_N_22_O_20_ | 41 | 803.3351 | 803.3379 | 3.48 | **G**WWGANW**D**GPSGMRD |
| *[M +H – Me_2_S – CO]^2+^* | C_73_H_92_N_22_O_20_ | 39 | 798.3429 | 798.3429 | 0.00 | **G**WWGANW**D**GPSGMRD |

**Table S6**. The assigned fragments of longipeptin C (**5**) [844 Da]; *Y_n*_* indicates *Y* ions upon MeSOH (64 Da) loss, degree of unsaturation (RDB)

| Fragment | Ion Formula | RDB | Calc. *m/z* | Meas. *m/z* | error in ppm | Sequence |
| --- | --- | --- | --- | --- | --- | --- |
| *y_1_* | C_4_H_8_NO_4_ | 2 | 134.0453 | --------- | --------- | GANWDGPSGMR |
| *y_2_* | C_10_H_20_N_5_O_5_ | 4 | 290.1464 | 290.1463 | 0.34 | GANWDGPSGMRD |
| *z_2_* | C_10_H_17_N_4_O_5_ | 5 | 273.1199 | 273.1191 | 2.93 |  |
| *y_3_* | C_15_H_29_N_6_O_7_S | 5 | 437.1818 | 437.1821 | 0.69 | GANWDGPSGMRD |
| *y_3*_* | C_14_H_25_N_6_O_6_ | 6 | 373.1836 | 373.1831 | 1.34 |  |
| *y_4_* | C_17_H_32_N_7_O_8_S | 6 | 494.2033 | 494.2035 | 0.40 | GANWDGPSGMRD |
| *y_4*_* | C_16_H_28_N_7_O_7_ | 7 | 430.2050 | 430.2056 | 1.39 |  |
| *y_5_* | C_20_H_37_N_8_O_10_S | 7 | 581.2353 | 581.2343 | 1.72 | GANWDGPSGMRD |
| *y_5*_* | C_19_H_33_N_8_O_9_ | 8 | 517.2370 | 517.2372 | 0.39 |  |
| *y_5_-H_2_O* | C_20_H_35_N_8_O_9_S | 8 | 563.2248 | 563.2241 | 1.24 |  |
| *y_6_* | C_25_H_44_N_9_O_11_S | 9 | 678.2881 | 678.2879 | 0.30 | G PSGMRD |
| *y_6*_* | C_24_H_40_N_9_O_10_ | 10 | 614.2898 | 614.2910 | 1.95 |  |
| *y_6_-H_2_O* | C_25_H_42_N_9_O_10_S | 10 | 660.2775 | 660.2776 | 0.15 |  |
| *y_7_* | C_27_H_47_N_10_O_12_S | 10 | 735.3096 | 735.3099 | 0.41 | GANWDGPSGMRD |
| *y_7*_* | C_26_H_43_N_10_O_11_ | 11 | 671.3113 | 671.3112 | 0.15 |  |
| *y_7_-H_2_O* | C_27_H_45_N_10_O_11_S | 11 | 717.2990 | 717.3002 | 1.67 |  |
| *y_8_* | C_31_H_50_N_11_O_14_S | 13 | 832.3259 | --------- | --------- | GANW**D**GPSGMRD |
| *y_9_* | C_42_H_60_N_13_O_15_S | 20 | 1018.4053 | 1018.4202 | 14.63 | GANW**D**GPSGMRD |
| *y_10_* | C_46_H_66_N_15_O_17_S | 22 | 1132.4482 | 1132.4430 | 4.60 | GANW**D**GPSGMRD |
| *y_11_* | C_49_H_71_N_16_O_18_S | 23 | 1203.4853 | --------- | --------- |  |
| *y_12_* | C_51_H_74_N_17_O_19_S | 24 | 1260.5068 | --------- | --------- |  |
| *y_13_* | --------- | --- | --------- | --------- | --------- |  |
| *y_14_* | --------- | --- | --------- | --------- | --------- |  |
| *b_1_* | --------- | --- | --------- | --------- | --------- |  |
| *b_2_* | --------- | --- | --------- | --------- | --------- |  |
| *b_3_* | C_24_H_22_N_5_O_3_ | 17 | 428.1723 | 428.1737 | 3.26 | **G**WWGANWDGPSG |
| *a_3_* | C_23_H_22_N_5_O_2_ | 16 | 400.1773 | 400.1754 | 4.74 |  |
| *b_4_* | C_26_H_25_N_6_O_4_ | 18 | 485.1937 | 485.1947 | 2.06 | **G**WWGANWDGPSG |
| *a_4_* | C_25_H_25_N_6_O_3_ | 17 | 457.1988 | --------- | --------- |  |
| *b_5_* | C_29_H_30_N_7_O_5_ | 19 | 556.2308 | 556.2293 | 2.70 | **G**WWGANWDGPSG |
| *a_5_* | C_28_H_30_N_7_O_4_ | 18 | 528.2359 | --------- | --------- |  |
| *b_6_* | C_33_H_36_N_9_O_7_ | 21 | 670.2738 | 670.2753 | 2.23 | **G**WWGANWDGPSG |
| *a_6_* | C_32_H_36_N_9_O_6_ | 20 | 642.2789 | --------- | --------- |  |
| *b_7_* | C_44_H_46_N_11_O_8_ | 28 | 856.3531 | 856.3567 | 4.20 | **G**WWGANWDGPSG |
| *b_8_* | C_48_H_49_N_12_O_10_ | 31 | 953.3695 | 953.3727 | 3.35 | **G**WWGANW**D**GPSG |
| *b_8_-NH_3_* | C_48_H_46_N_11_O_10_ | 32 | 936.3429 | 936.3417 | 1.28 |  |
| *a_8_* | C_47_H_49_N_12_O_9_ | 30 | 925.3745 | 925.3743 | 0.21 |  |
| *a_8_-NH_3_* | C_47_H_46_N_11_O_9_ | 31 | 908.3480 | 908.3497 | 1.87 |  |
| *b_9_* | C_50_H_52_N_13_O_11_ | 32 | 1010.3909 | 1010.3972 | 6.23 | **G**WWGANW**D**GPSG |
| *b_9_-NH_3_* | C_50_H_49_N_12_O_11_ | 33 | 993.3644 | 993.3627 | 1.71 |  |
| *a_9_* | C_49_H_52_N_13_O_10_ | 31 | 982.3960 | 982.3906 | 5.49 |  |
| *a_9_-NH_3_* | C_49_H_49_N_12_O_10_ | 32 | 965.3695 | 965.3679 | 1.66 |  |

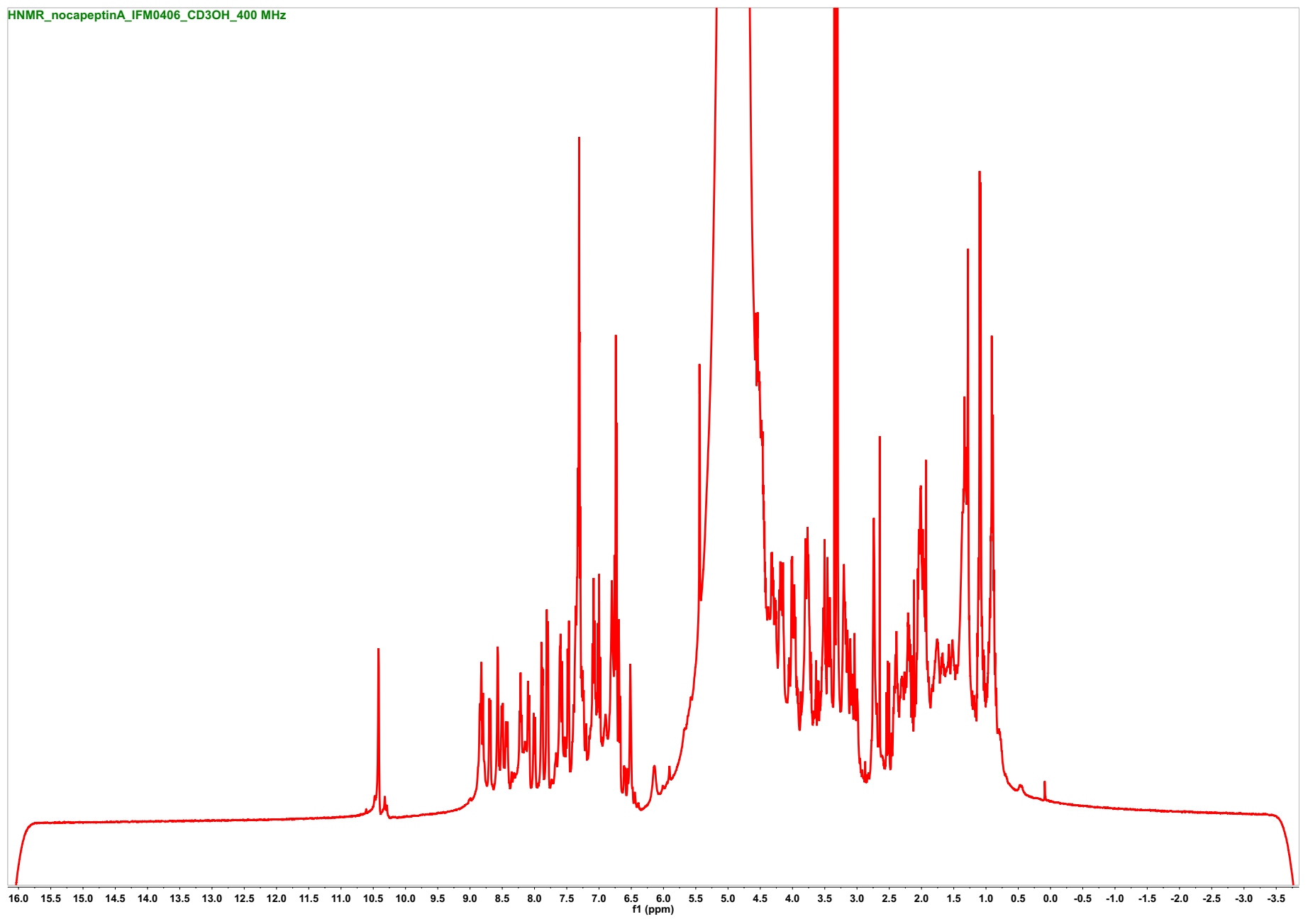

**Figure S14**. ^1^H- NMR spectrum of nocapeptin A (**1**) (*d*_3_-CH_3_OH, 400 MHz)

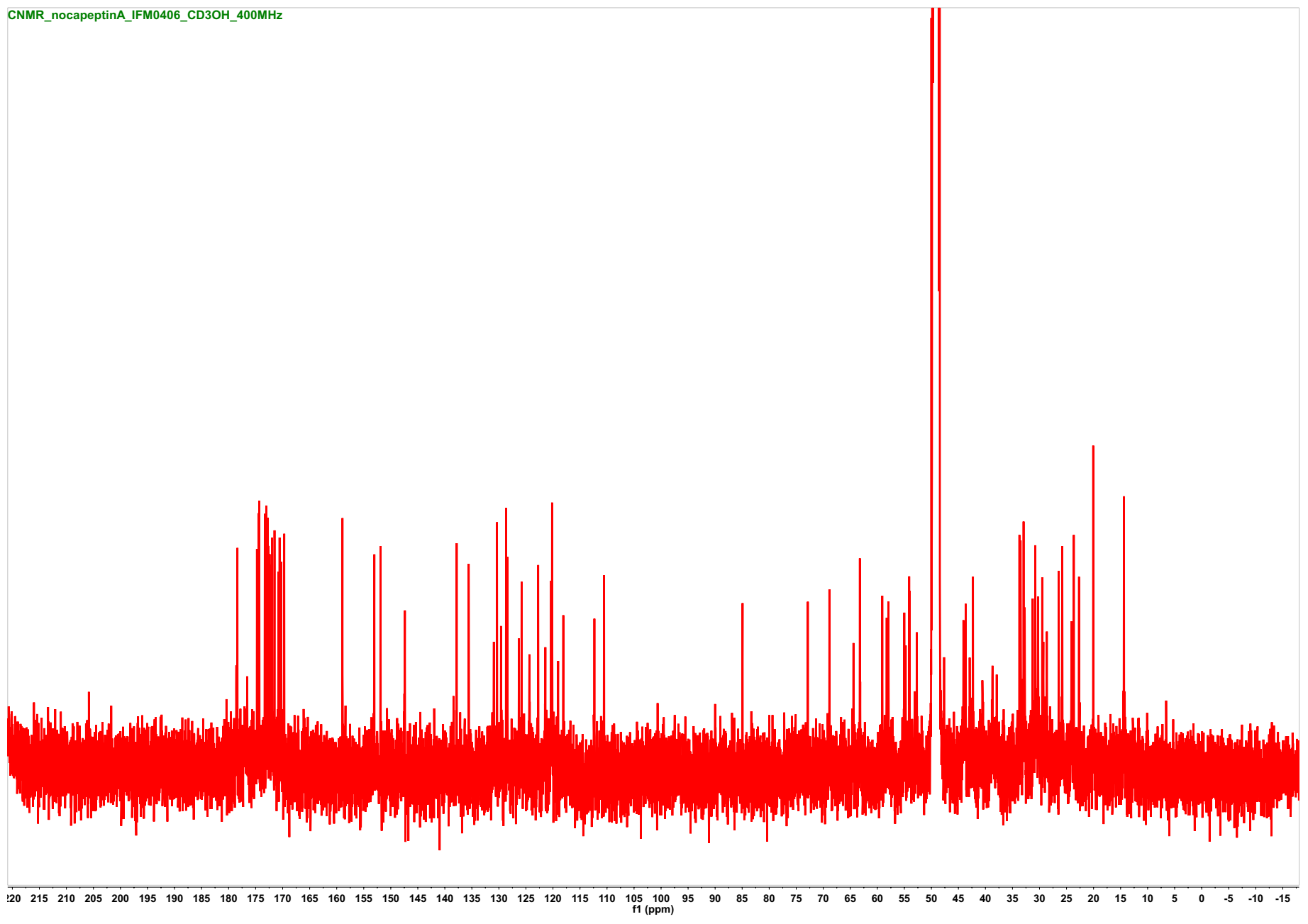

**Figure S15**. ^13^C- NMR spectrum of nocapeptin A (**1**) (*d*_3_-CH_3_OH, 100 MHz)

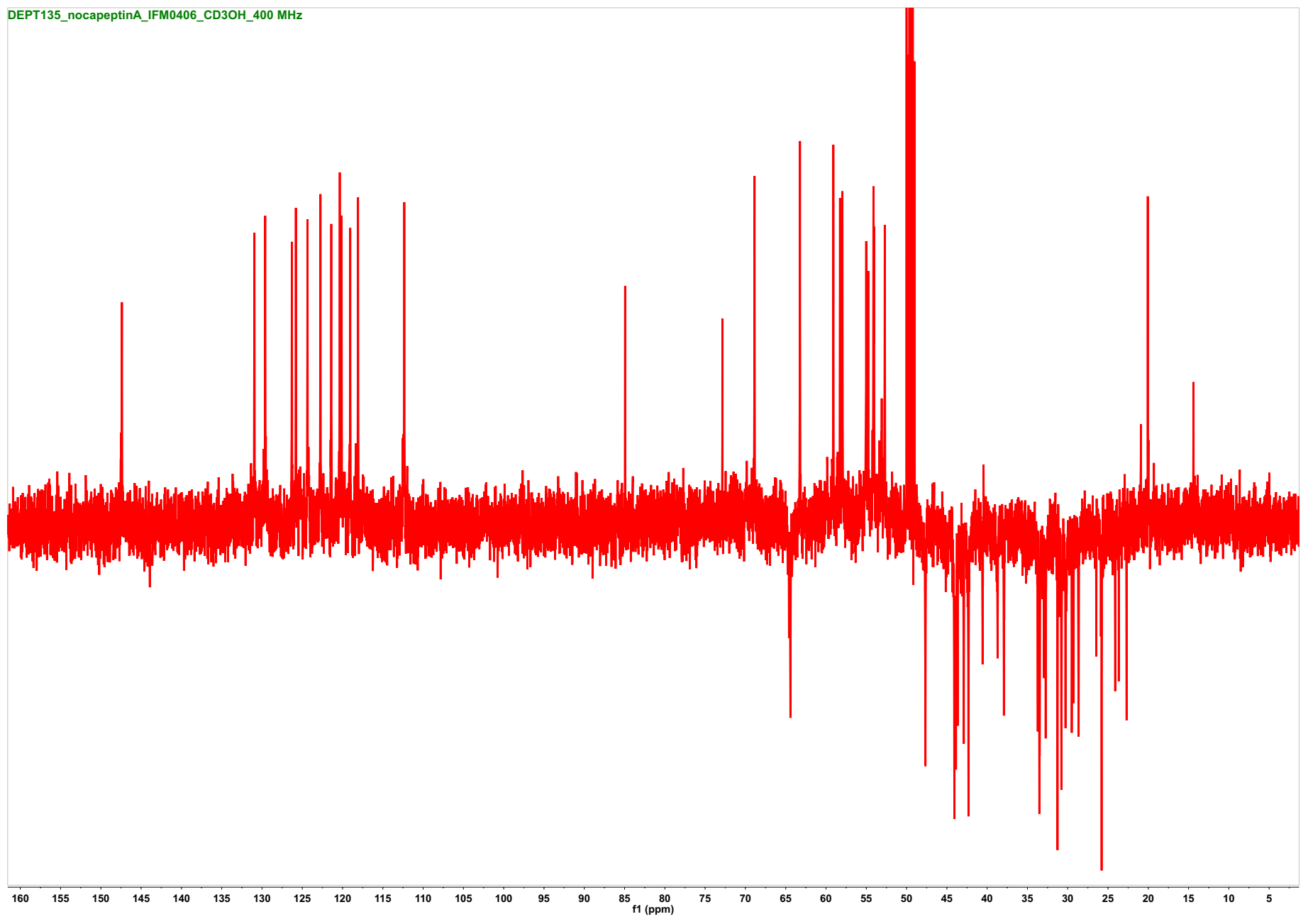
**Figure S16.** DEPT-135 spectrum of nocapeptin A (**1**) (*d*_3_-CH_3_OH, 100 MHz)

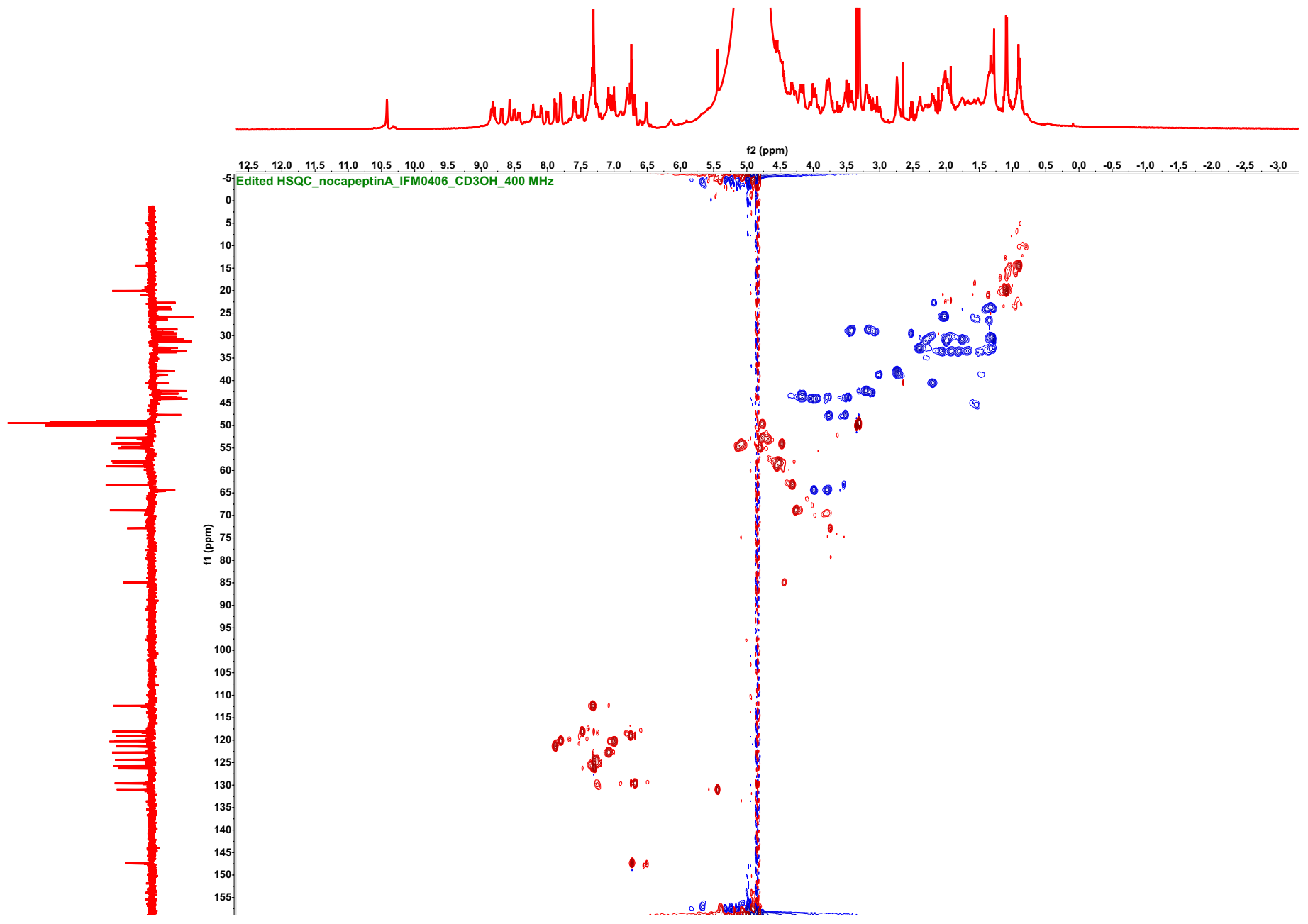

**Figure S17**. ^1^H-^13^C Edited HSQC spectrum of nocapeptin A (**1**) (CH/CH_3_ are in red and CH_2_ are in blue, *d*_3_-CH_3_OH, 400/100 MHz)

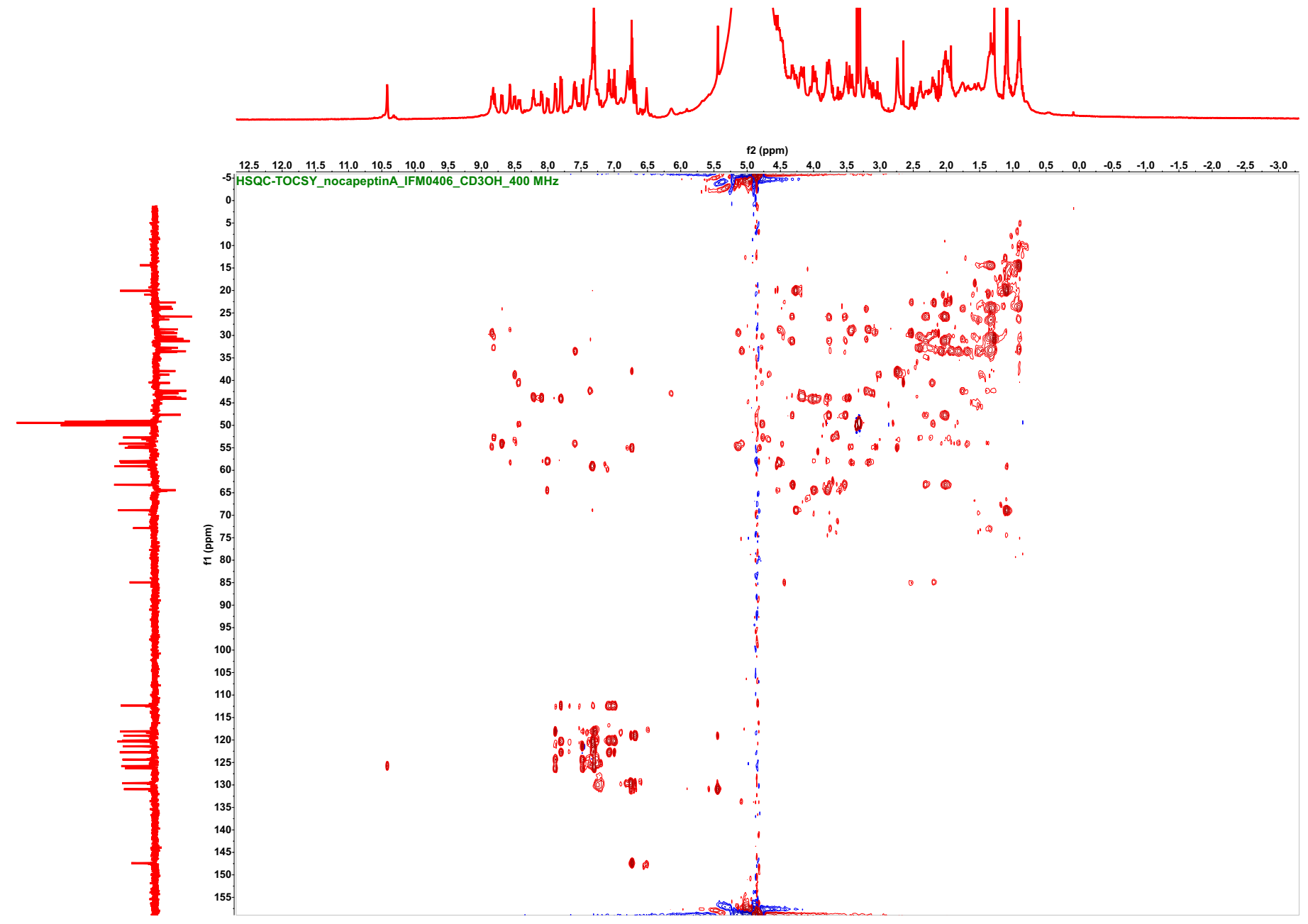

**Figure S18**. ^1^H-^13^C HSQC-TOCSY spectrum of nocapeptin A (**1**) (*d*_3_-CH_3_OH, 400/100 MHz)

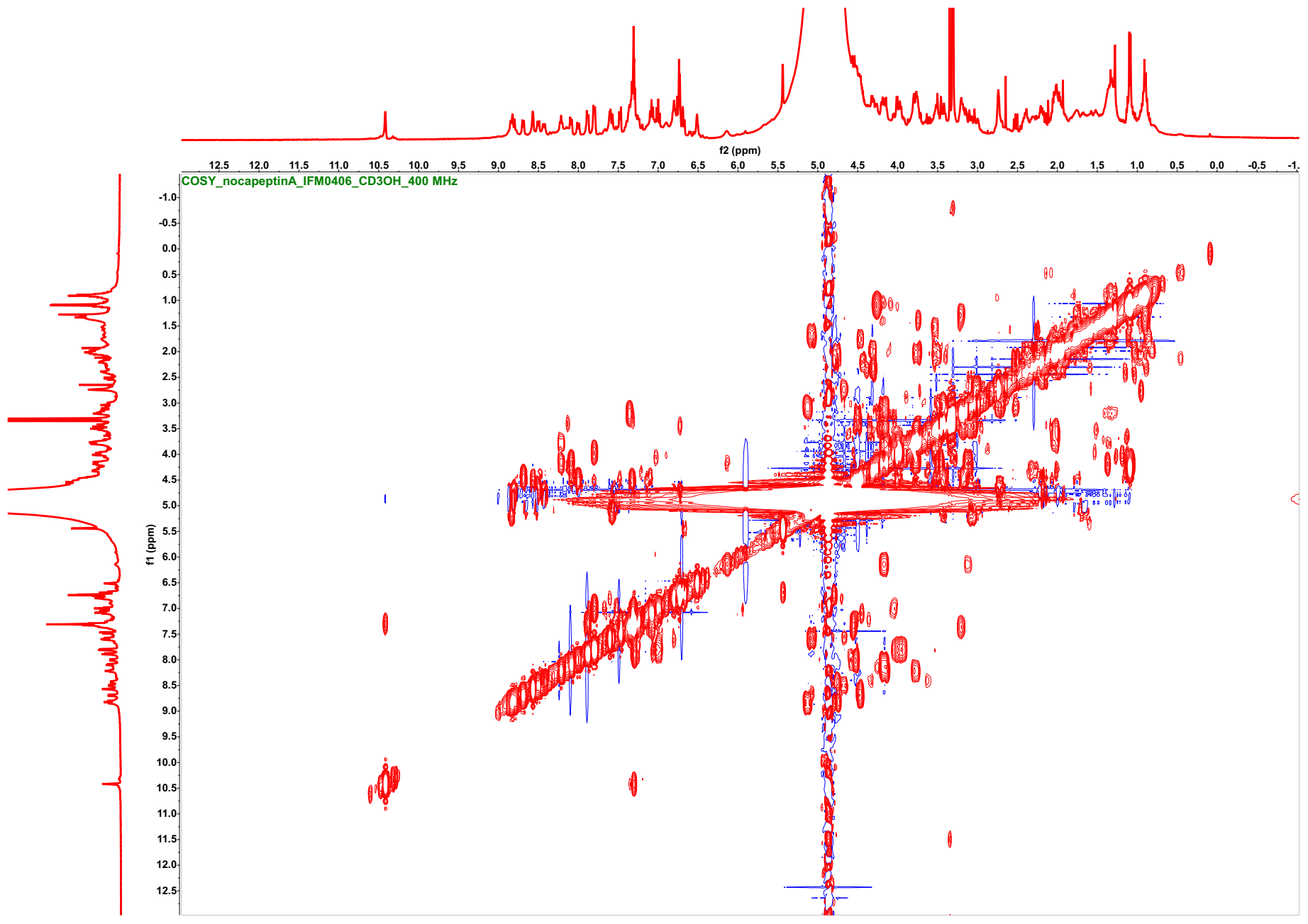

**Figure S19**. ^1^H-^1^H COSY spectrum of nocapeptin A (**1**) (*d*_3_-CH_3_OH, 400 MHz)

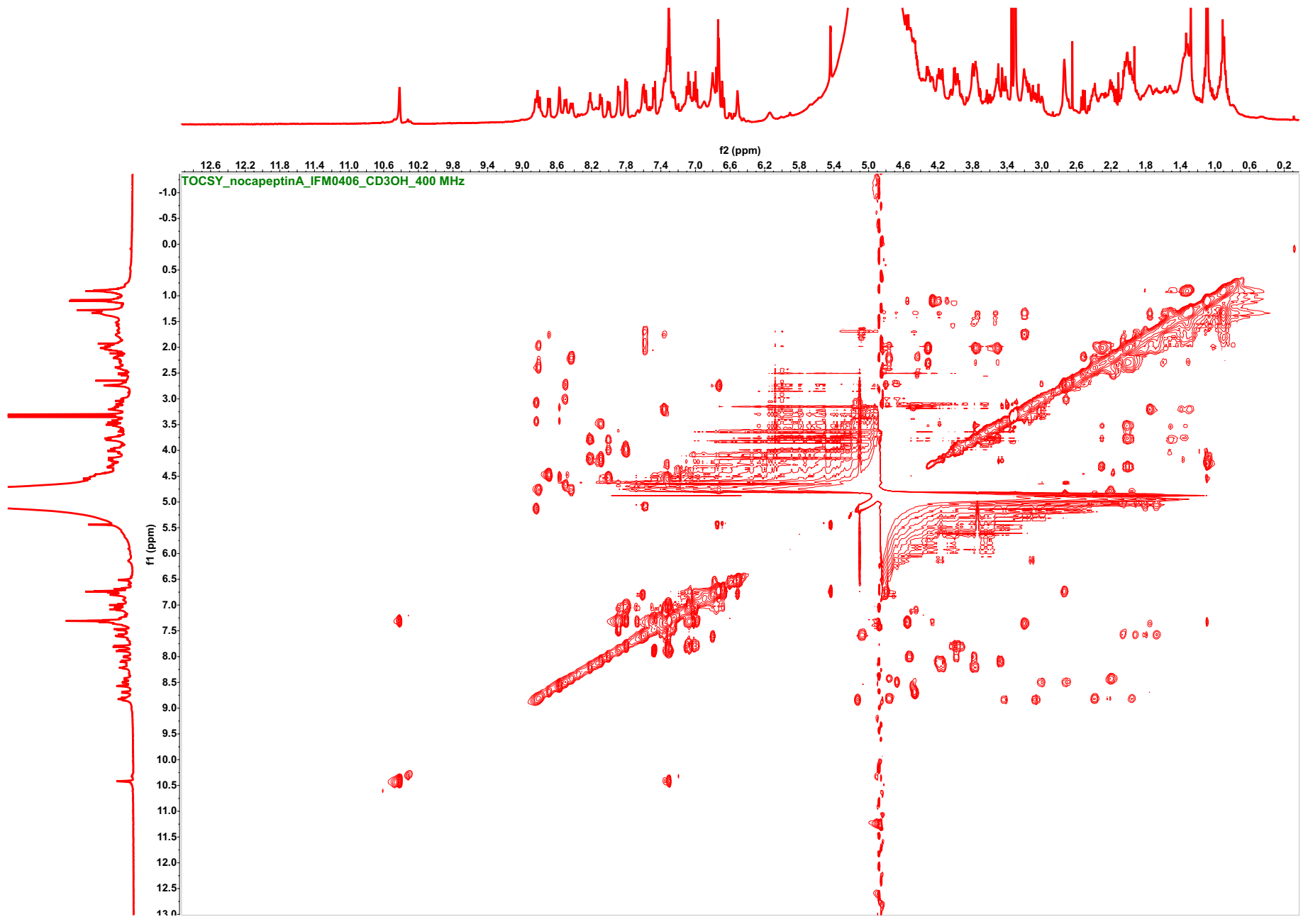

**Figure S20**. ^1^H-^1^H TOCSY spectrum of nocapeptin A (**1**) (*d*_3_-CH_3_OH, 400 MHz)

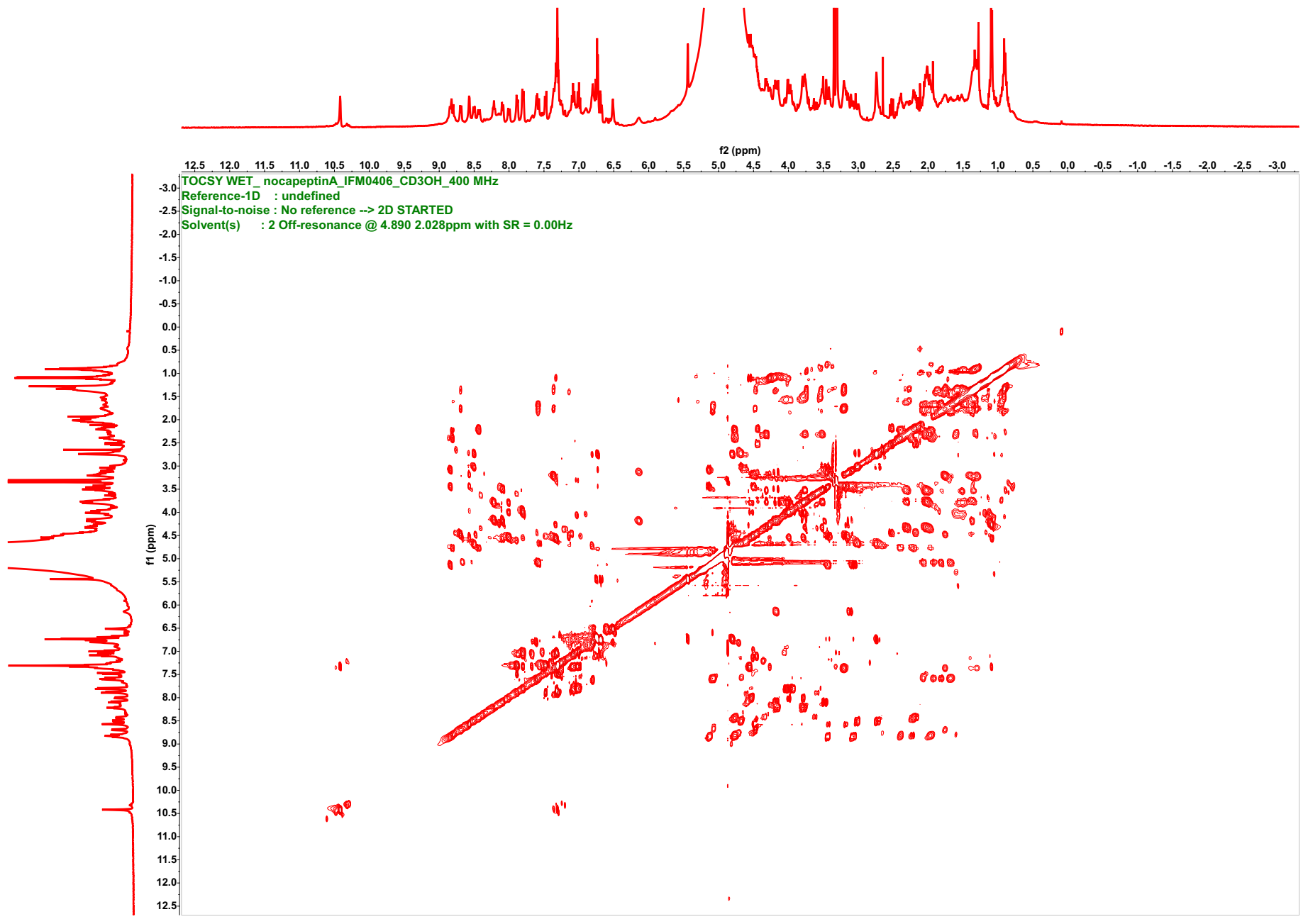

**Figure S20A**. ^1^H-^1^H TOCSY spectrum of nocapeptin A (**1**) with WET solvent suppression (*d*_3_-CH_3_OH, 400 MHz)

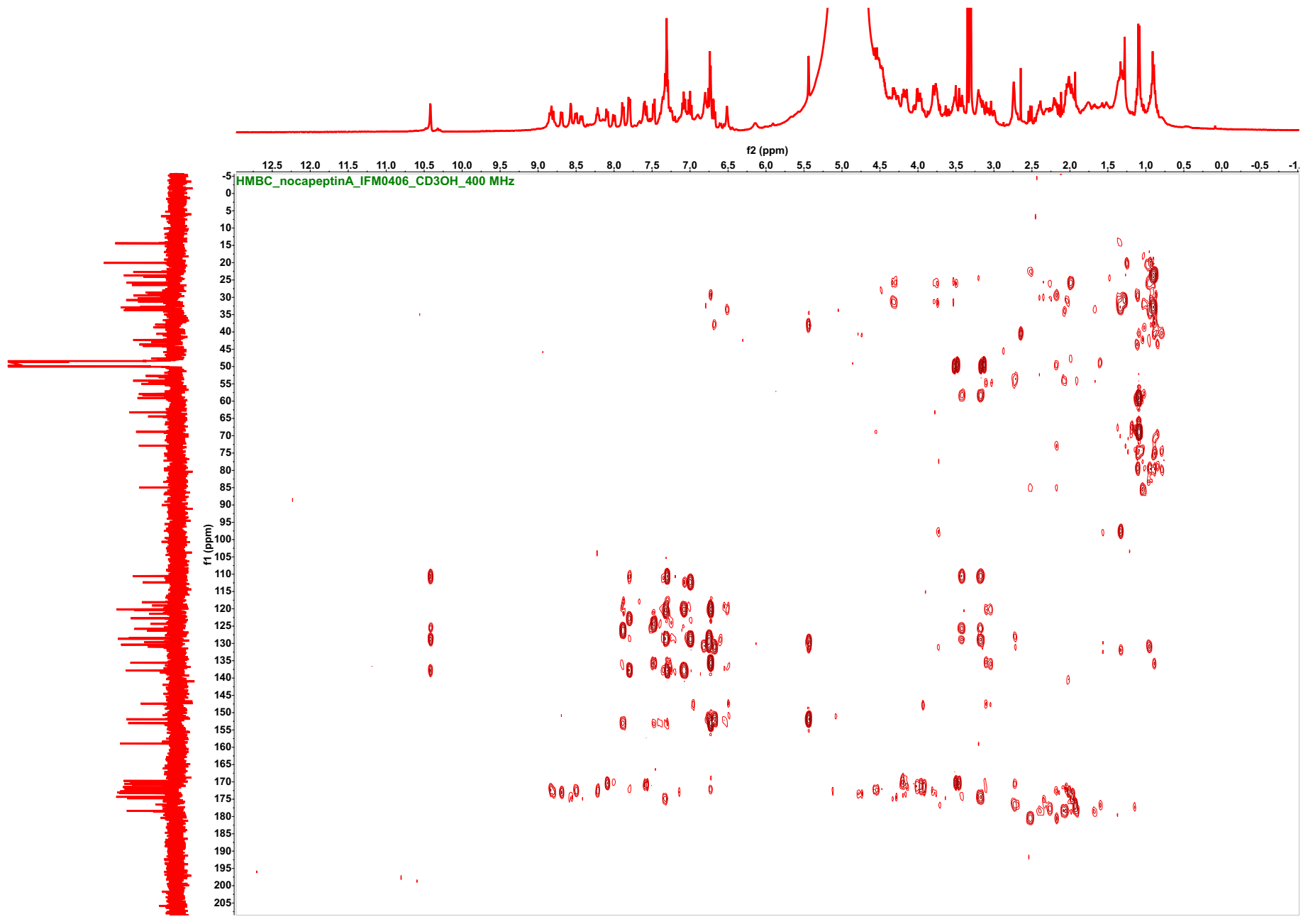

**Figure S21**. ^1^H-^13^C HMBC spectrum of nocapeptin A (**1**) (*d*_3_-CH_3_OH, 400/100 MHz)

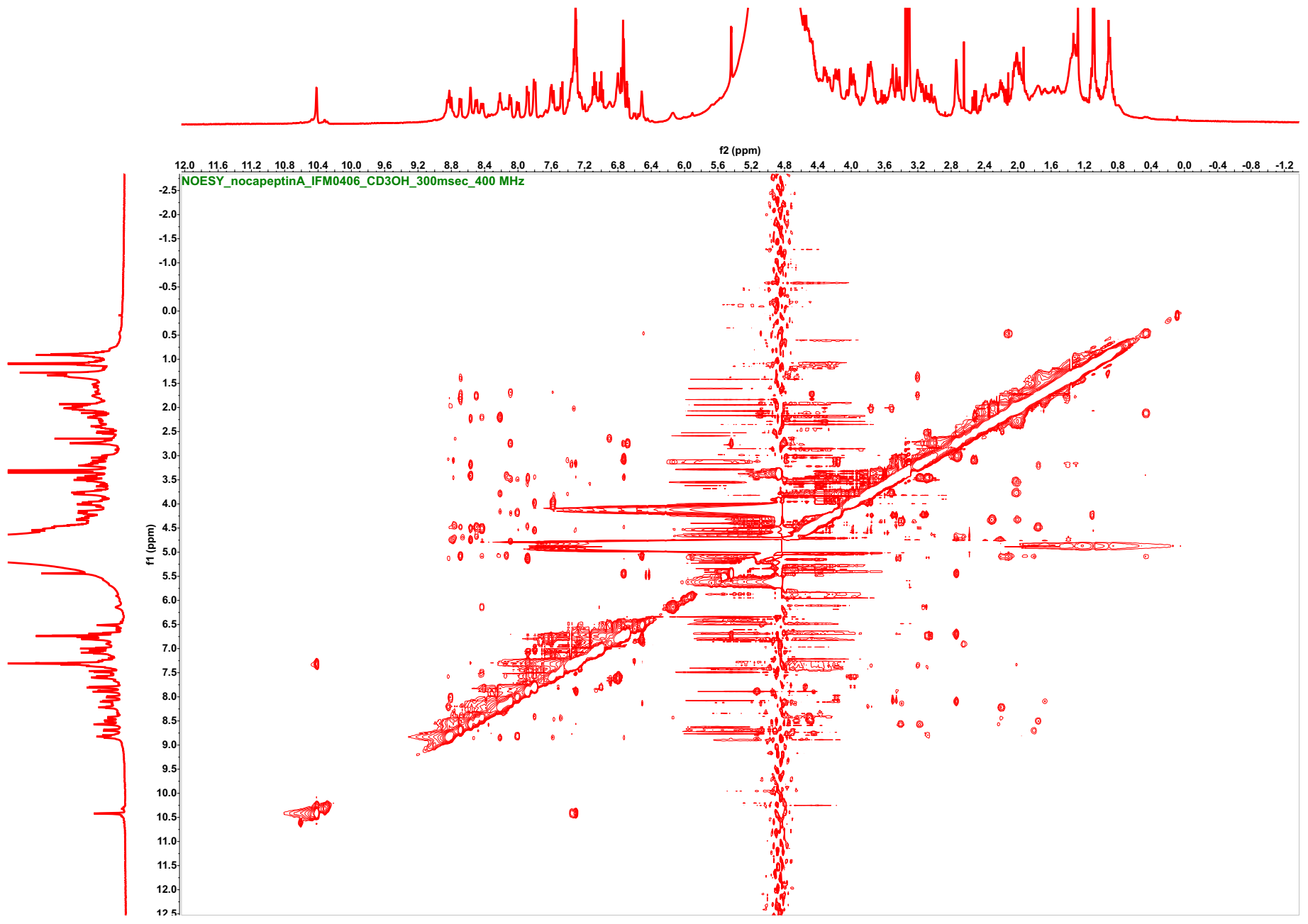

**Figure S22**. ^1^H-^1^H NOESY spectrum of nocapeptin A (**1**) (*d*_3_-CH_3_OH, 300 msec, 400 MHz)

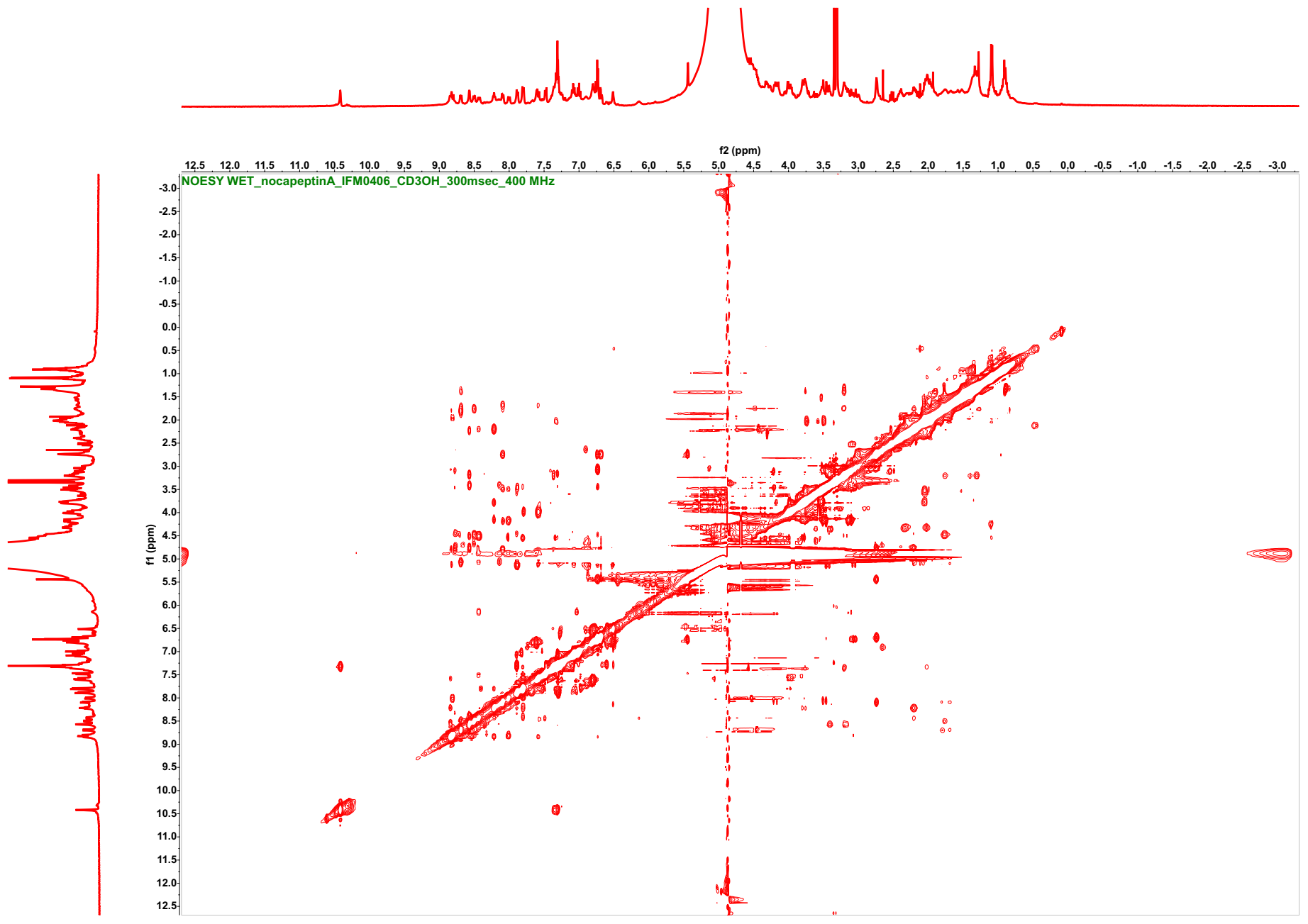

**Figure S22A**. ^1^H-^1^H NOESY spectrum of nocapeptin A (**1**) with WET solvent suppression (*d*_3_-CH_3_OH, 300 msec, 400 MHz)

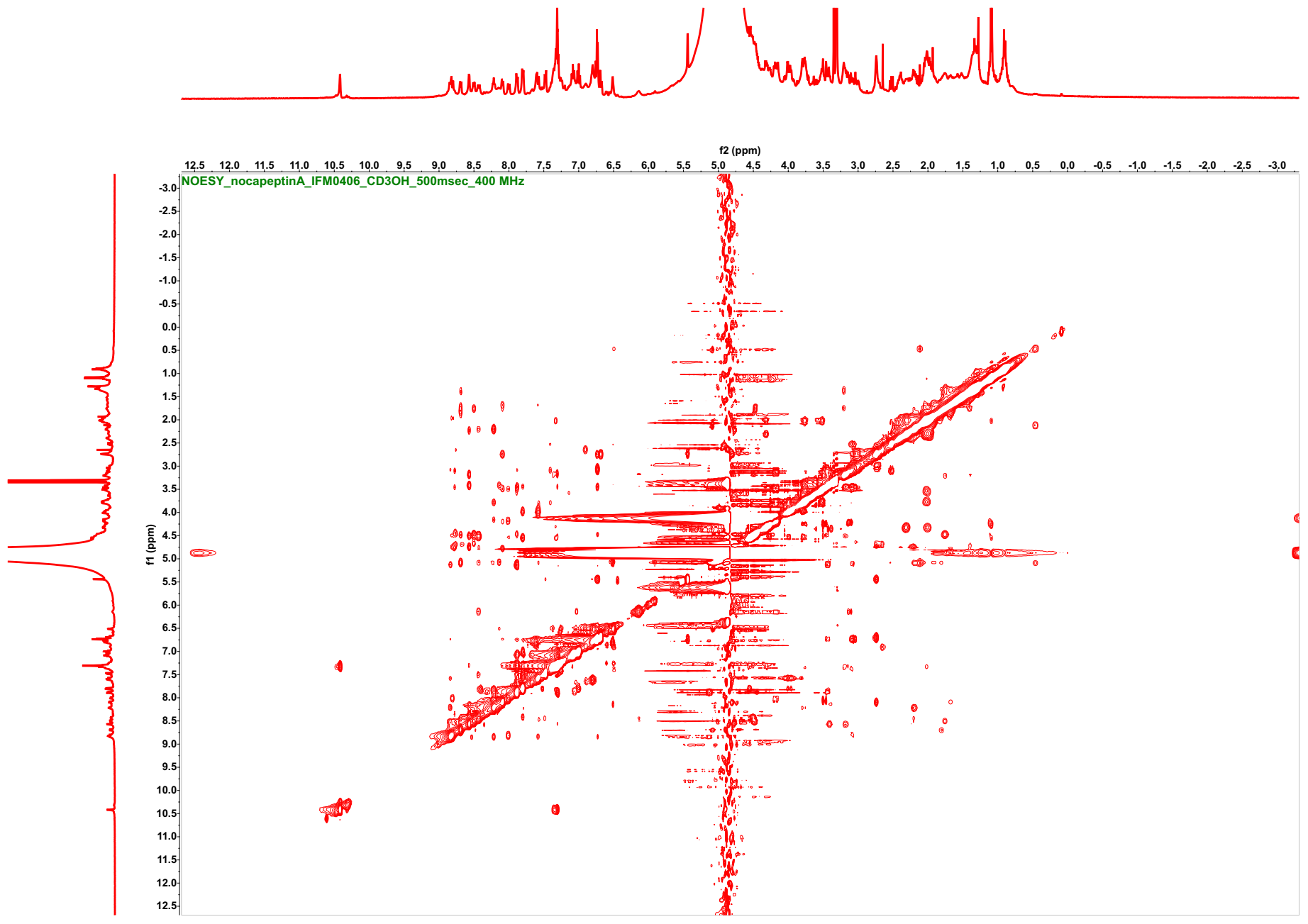

**Figure S23**. ^1^H-^1^H NOESY spectrum of nocapeptin A (**1**) (*d*_3_-CH_3_OH, 500 msec, 400 MHz)

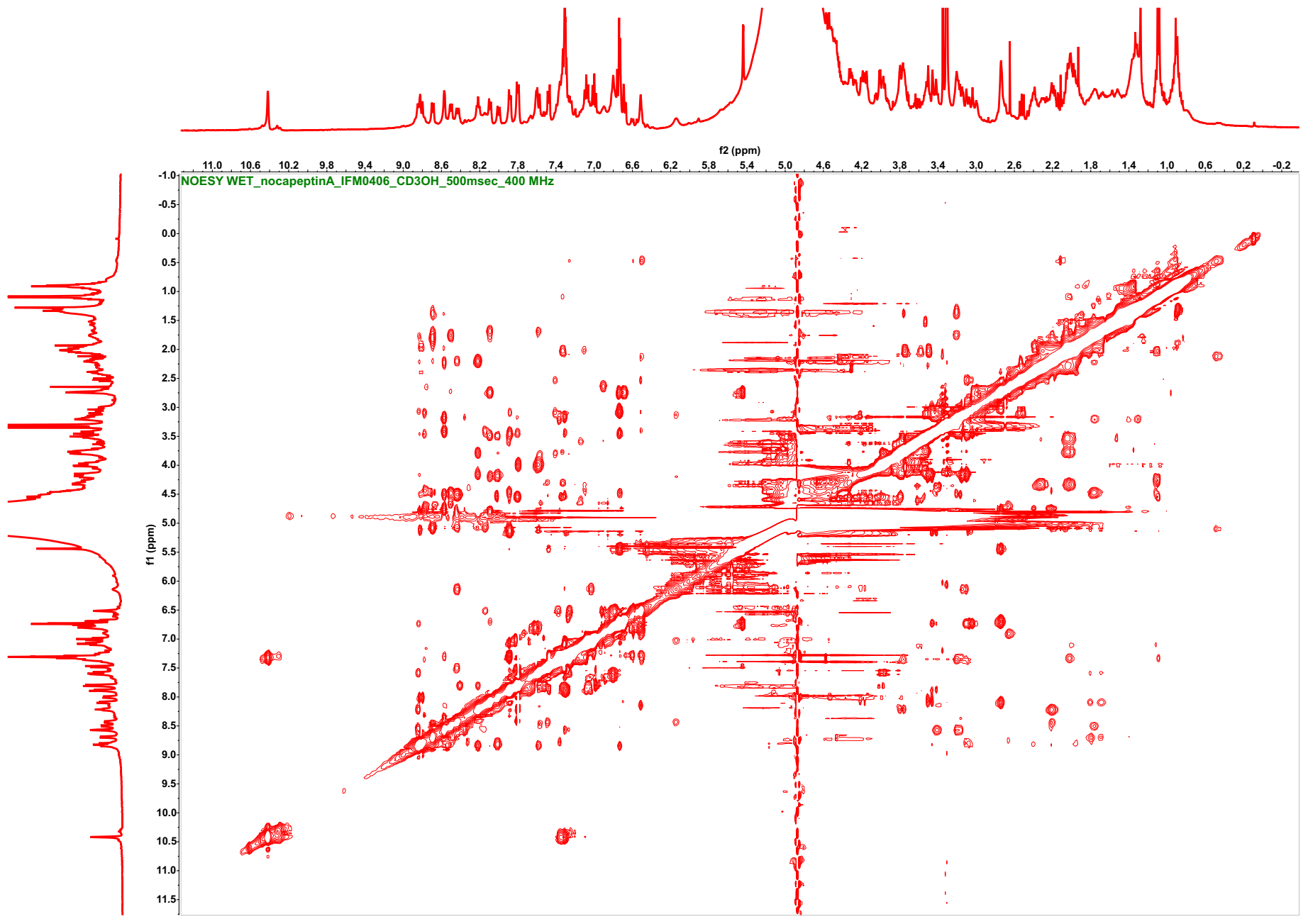

**Figure S23A**. ^1^H-^1^H NOESY spectrum of nocapeptin A (**1**) with WET solvent suppression (*d*_3_-CH_3_OH, 500 msec, 400 MHz)

**Table S7**. ^1^H, and ^13^C-NMR data of nocapeptin A (**1**) (*d*_3_-CH_3_OH; 400/100 MHz)

| **Residue** | **Position** | **δ_C_** | **δ_H_, mult (*J* in Hz)** | **Residue** | **Position** | **δ_C_** | **δ_H_, mult (*J* in Hz)** |
| --- | --- | --- | --- | --- | --- | --- | --- |
| **Gly1** | CO | n.d. | --------- | **Trp7** | CO | n.d. | --------- |
|  | α | 43.65, CH_2_ | 3.78, m + 4.17, m |  | α | 58.25, CH | 4.51, overlapped |
|  | -NH- | --------- | 8.22, dd (6.2, 5.9) |  | β | 28.65, CH_2_ | 3.17, m + 3.42, m |
| **Trp2*** | CO | n.d. | --------- |  | 1 (NH) | --------- | 10.42, s |
|  | α | 54.75, CH | 5.14, masked |  | 2 | 125.80, CH | 7.30, overlapped |
|  | β | 29.27, CH_2_ | 3.07, m + 3.44, dm (14.9) |  | 3 | 110.55, C | --------- |
|  | 1 (N) | --------- | --------- |  | 3a | 128.42/128.66, C | --------- |
|  | 2 | 147.39, CH | 6.73, s |  | 4 | 120.15, CH | 7.80, overlapped |
|  | 3 | 120.34, C | --------- |  | 5 | 120.34, CH | 7.00, dd (7.8, 7.5) |
|  | 3a | 135.59, C | --------- |  | 6 | 122.76, CH | 7.08, dd (7.6, 7.3) |
|  | 4 | 121.40, CH | 7.88, dd (6.0, 3.5) |  | 7 | 112.34, CH | 7.32, overlapped |
|  | 5 | 124.33, CH | 7.29, overlapped |  | 7a | 137.81, C | --------- |
|  | 6 | 126.29, CH | 7.30, overlapped |  | -NH- | --------- | 8.57, br d |
|  | 7 | 118.08, CH | 7.47, d (7) | **Asp8** | CO | n.d. | --------- |
|  | 7a | 153.04, C | --------- |  | α | 49.66, CH | 4.77, masked |
|  | -NH- | --------- | 8.83, d (8) |  | β | 40.56, CH_2_ | 2.20, m |
| **Tyr3*** | CO | n.d. | --------- |  | γ | n.d. | --------- |
|  | α | 55.00, CH | 4.80, masked |  | -NH- | --------- | 8.43, d (9.5) |
|  | β | 37.92, CH_2_ | 2.74, m | **Gly9** | CO | n.d. | --------- |
|  | 1 | 128.43/128.66, C | --------- |  | α | n.d. | n.d. |
|  | 2 | 130.94, CH | 5.44, d (1.9) |  | -NH- | --------- | n.d. |
|  | 3 | 130.36, C | --------- | **Pro10** | CO | n.d. | --------- |
|  | 4 | 151.89, C | --------- |  | α | 63.24, CH | 4.31, dd (9.6, 4.7) |
|  | 5 | 119.05, CH | 6.75, overlapped |  | β | 31.25, CH_2_ | 2.00, m + 2.30, m |
|  | 6 | 129.60, CH | 6.68, dd (8.4, 1.9) |  | γ | 25.80, CH_2_ | 2.03, m |
|  | OH | --------- | n.d. |  | δ | 47.66, CH_2_ | 3.52, m + 3.76, m |
|  | -NH- | --------- | 6.72, overlapped |  | =N- | --------- | --------- |
| **Gly4** | CO | n.d. | --------- | **Thr11** | CO | n.d. | --------- |
|  | α | 43.85, CH_2_ | 3.48, m + 4.18, m |  | α | 59.08, CH | 4.55, masked |
|  | -NH- | --------- | 8.09, m |  | β | 68.86, CH | 4.26, m |
| **Ser5** | CO | n.d. | --------- |  | γ | 20.06, CH_3_ | 1.09, d (6.5) |
|  | α | 57.96/58.26, CH | 4.53, m |  | OH | --------- | n.d. |
|  | β | 64.41, CH_2_ | 3.79, m + 3.99, m |  | -NH- | --------- | 7.32, overlapped |
|  | OH | --------- | n.d. | **Gly12** | CO | n.d. | --------- |
|  | -NH- | --------- | 8.00, d (8.9) |  | α | 44.07, CH_2_ | 3.98, m |
| **Gln6** | CO | n.d. | --------- |  | -NH- | --------- | 7.80, m |
|  | α | 52.69, CH | 4.76, m | **Gln13** | CO | n.d. | --------- |
|  | β | 30.26, CH_2_ | 1.96, m + 2.23, m |  | α | 54.03/54.10, CH | 5.09, masked |
|  | γ | 32.73, CH_2_ | 2.39, m |  | β | 33.47/33.51, CH_2_ | 1.68, m + 1.81, m |
|  | δ | n.d. | --------- |  | γ | 33.47/33.51, CH_2_ | 1.93, m + 2.08, m |
|  | -NH_2_ | --------- | n.d. |  | δ | n.d. | --------- |
|  | -NH- | --------- | 8.81, d (9.7) |  | -NH_2_ | --------- | n.d. |
|  |  |  |  |  | -NH- | --------- | 7.60, overlapped |
| **Arg14** | CO | n.d. | --------- |  | | | |
|  | α | 54.10, CH | 4.47, m |  | | | |
|  | β | 30.78, CH_2_ | 1.75, m |  | | | |
|  | γ | 24.09, CH_2_ | 1.29, m + 1.39, m |  | | | |
|  | δ | 42.31, CH_2_ | 3.21, m |  | | | |
|  | εNH | --------- | 7.36, overlapped |  |  |  |  |
|  | ζ | 158.94**, C | --------- |  |  |  |  |
|  | NHη1 | --------- | n.d. |  |  |  |  |
|  | NH_2_η1 | --------- | n.d. |  |  |  |  |
|  | -NH- | --------- | 8.60, overlapped |  |  |  |  |
| **Asp15** | CO_2_H | n.d. | --------- |  |  |  |  |
|  | α | 53.08, CH | 4.68, masked |  |  |  |  |
|  | β | 38.70, CH_2_ | 2.71, m + 3.01, m |  |  |  |  |
|  | γ | n.d. | --------- |  |  |  |  |
|  | -NH- | --------- | 8.50, d (7.7) |  |  |  |  |
| * Residues involved in the C-N biaryl crosslink.  ** The chemical shift was assigned based on literature values. | | | |  |  |  |  |

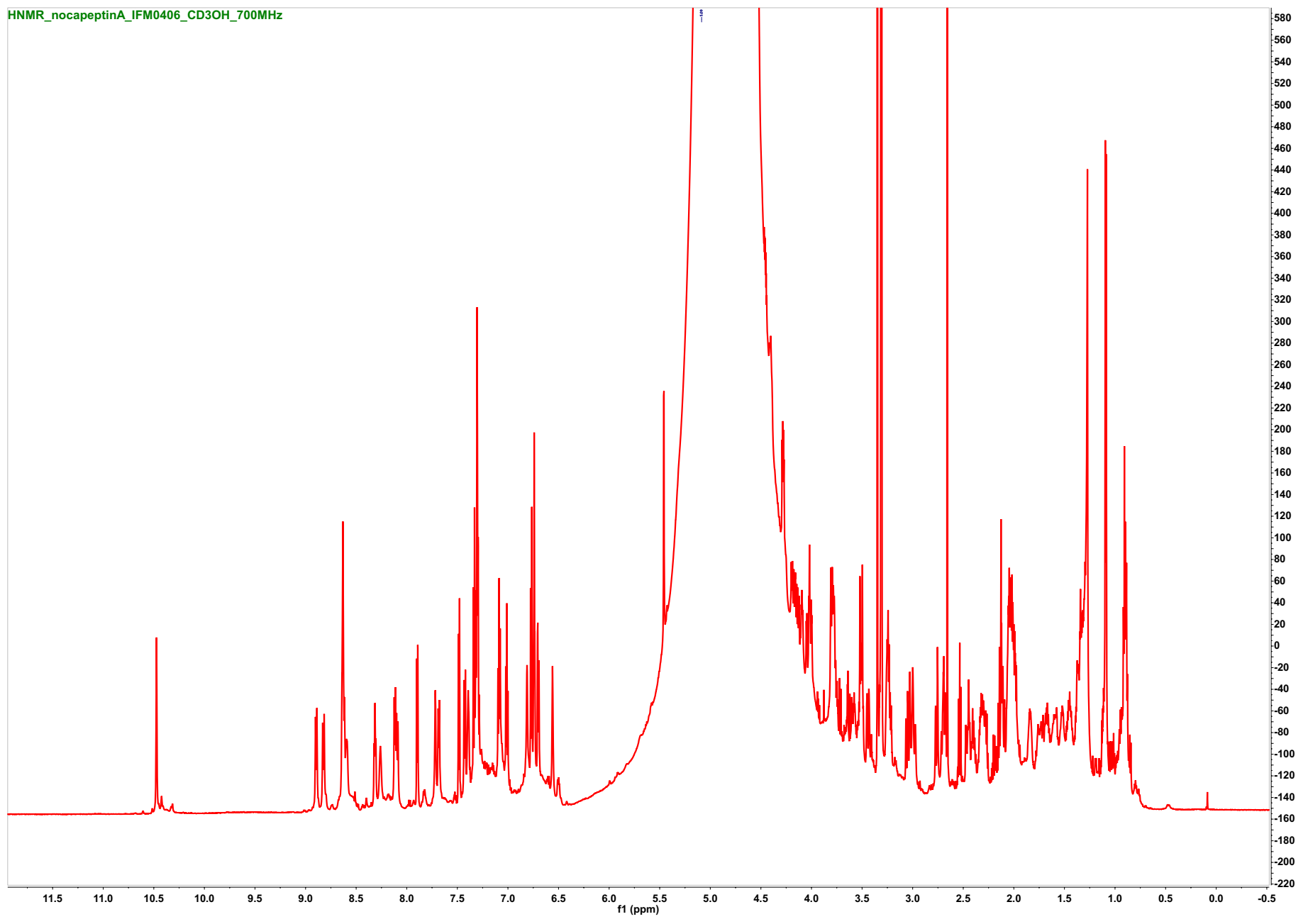

**Figure S24**. ^1^H- NMR spectrum of nocapeptin A (**1**) (*d*_3_-CH_3_OH/H_2_O (96:4), 700 MHz)

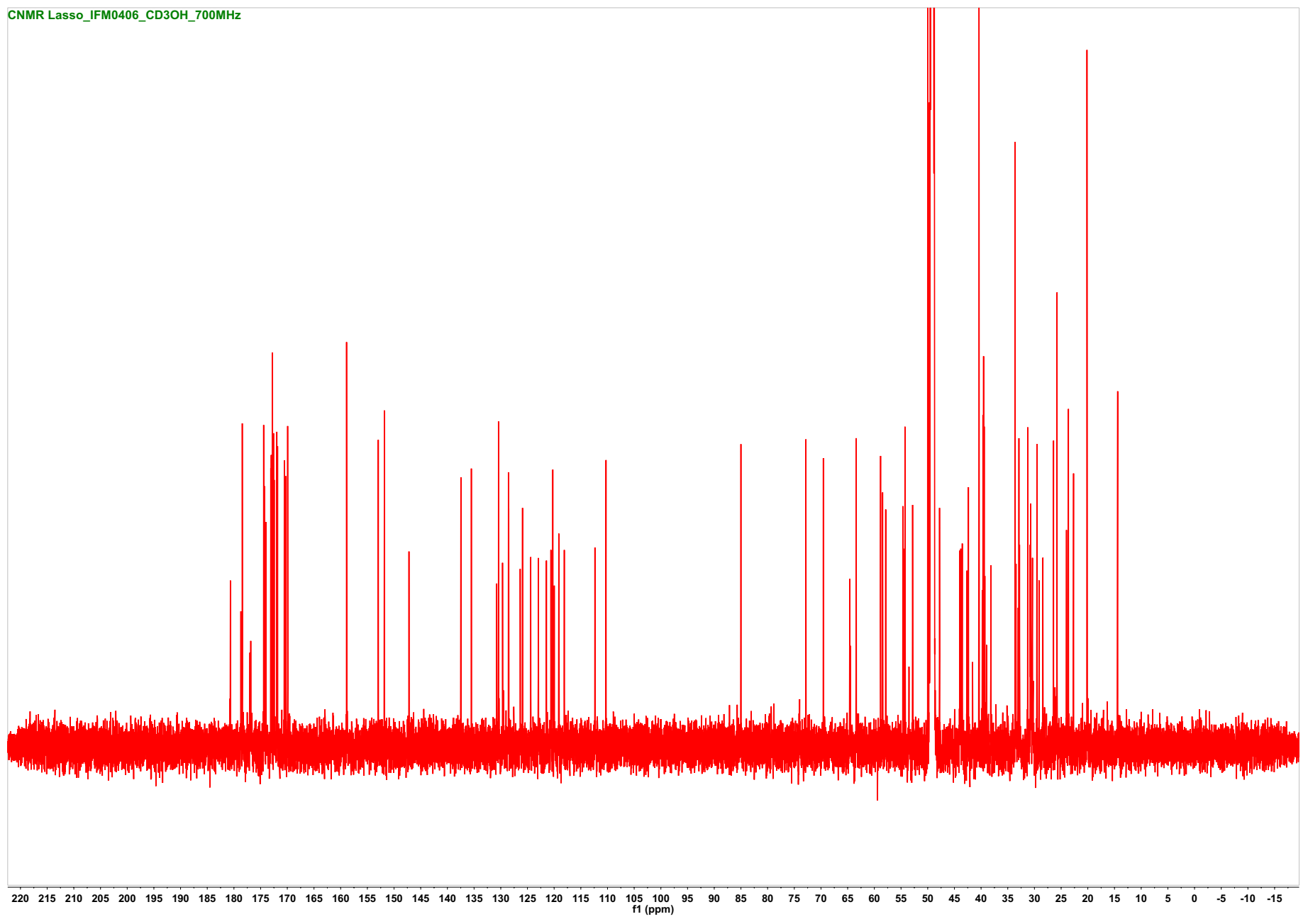

**Figure S25**. ^13^C- NMR spectrum of nocapeptin A (**1**) (*d*_3_-CH_3_OH/H_2_O (96:4), 176 MHz)

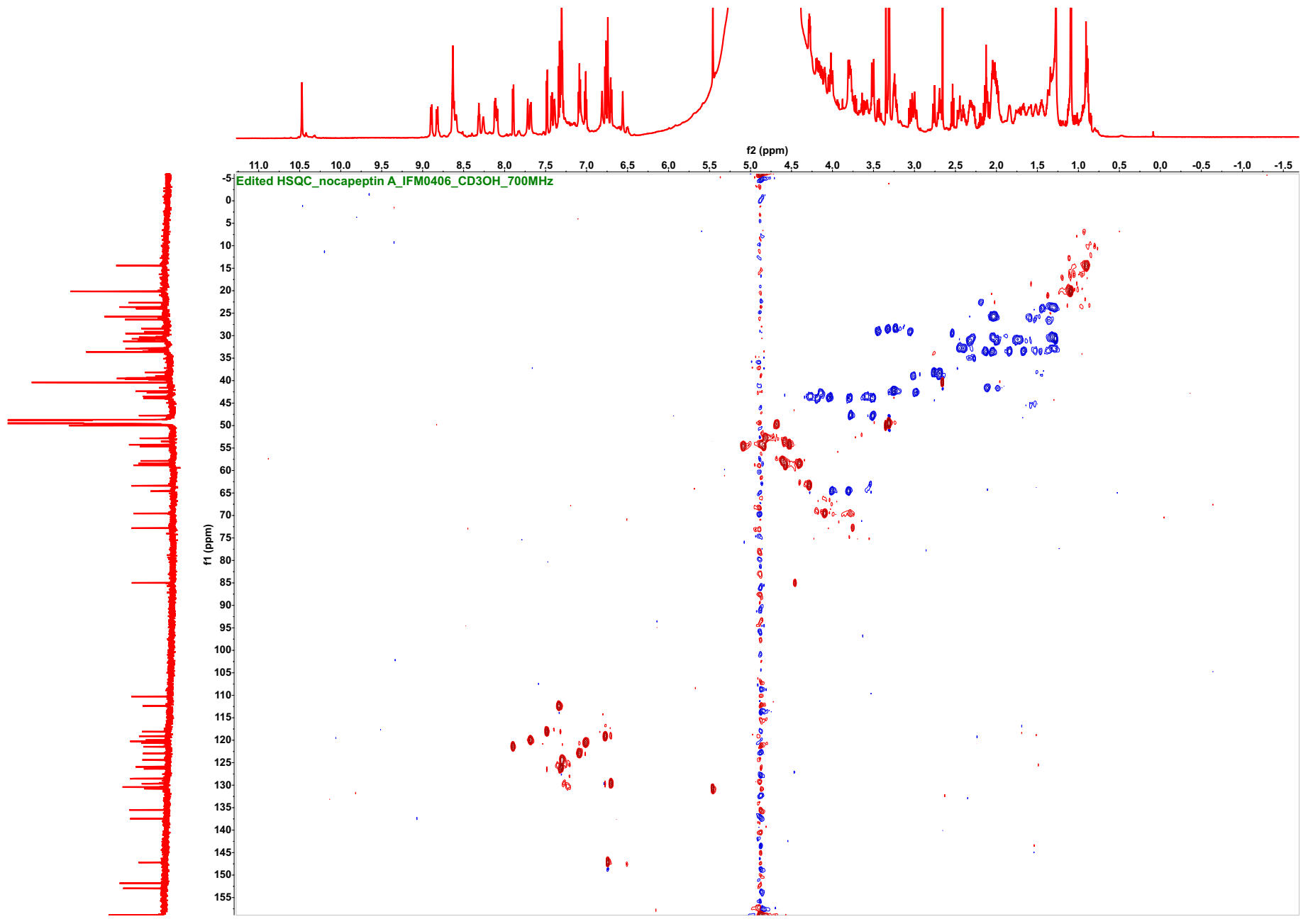

**Figure S26**. ^1^H-^13^C Edited HSQC spectrum of nocapeptin A (**1**) (CH/CH_3_ are in red and CH_2_ are in blue, *d*_3_-CH_3_OH/H_2_O (96:4), 700/176 MHz)

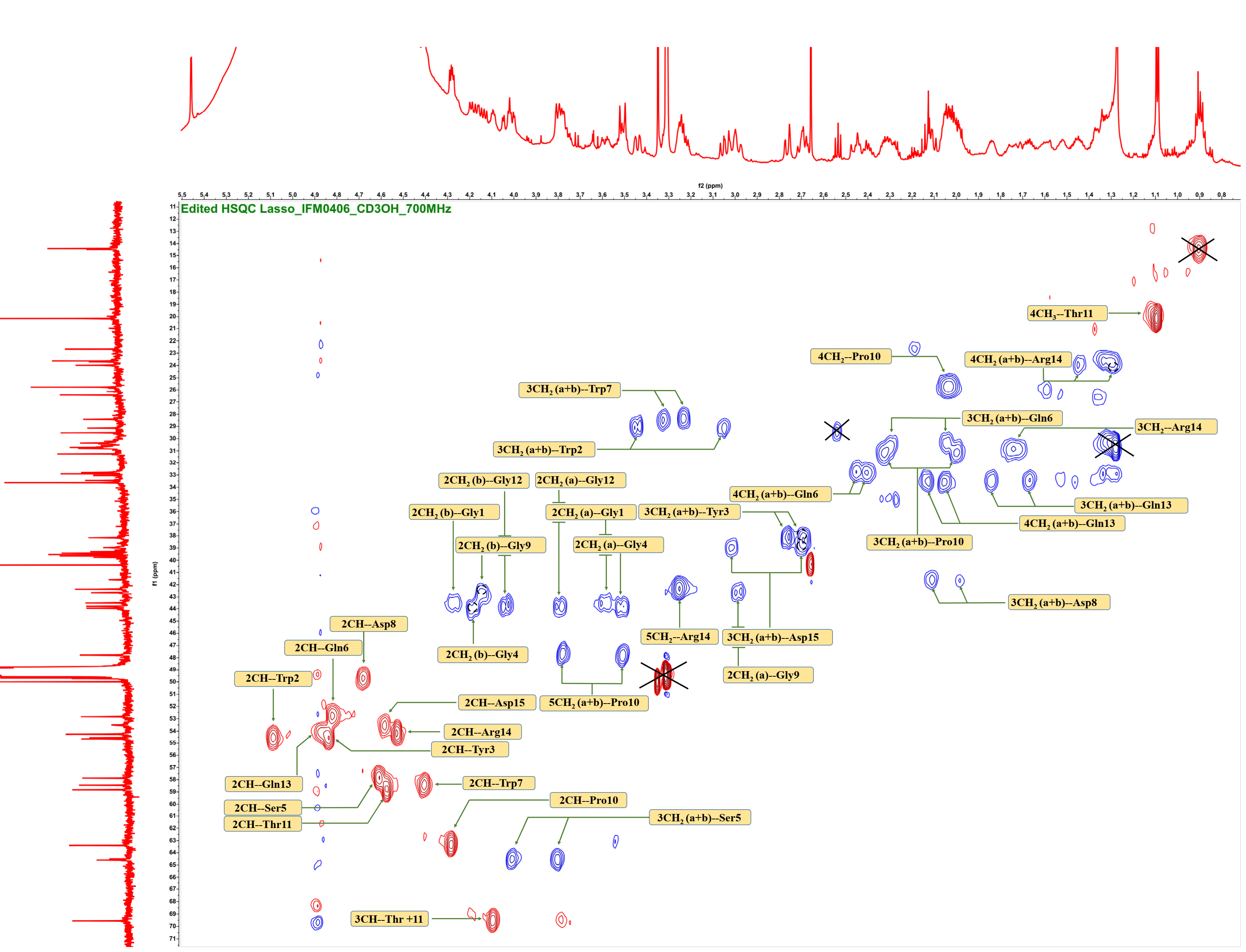

**Figure S26A**. Annotated ^1^H-^13^C edited HSQC spectrum of nocapeptin A (**1**)

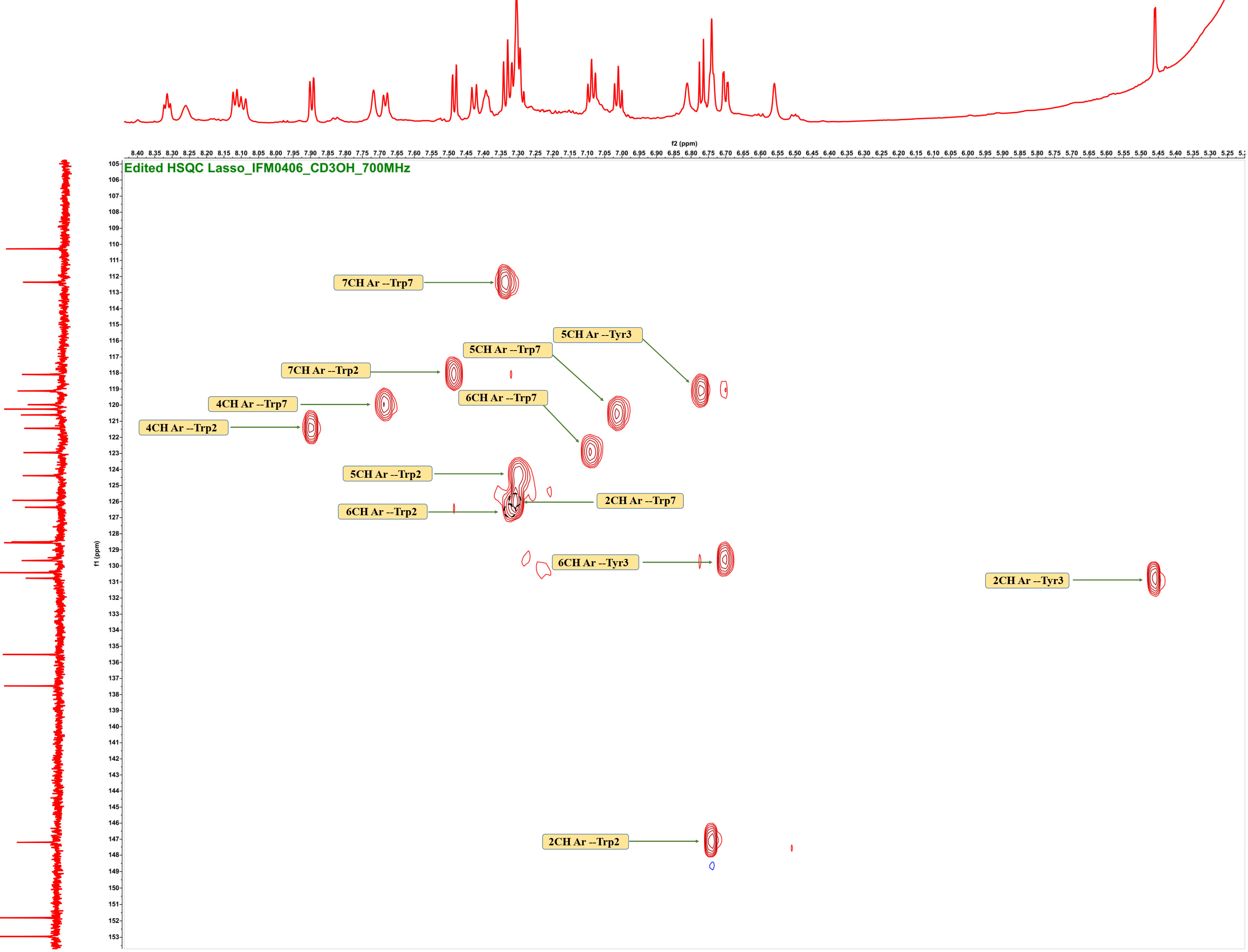

**Figure S26B**. Annotated ^1^H-^13^C edited HSQC spectrum of nocapeptin A (**1**)

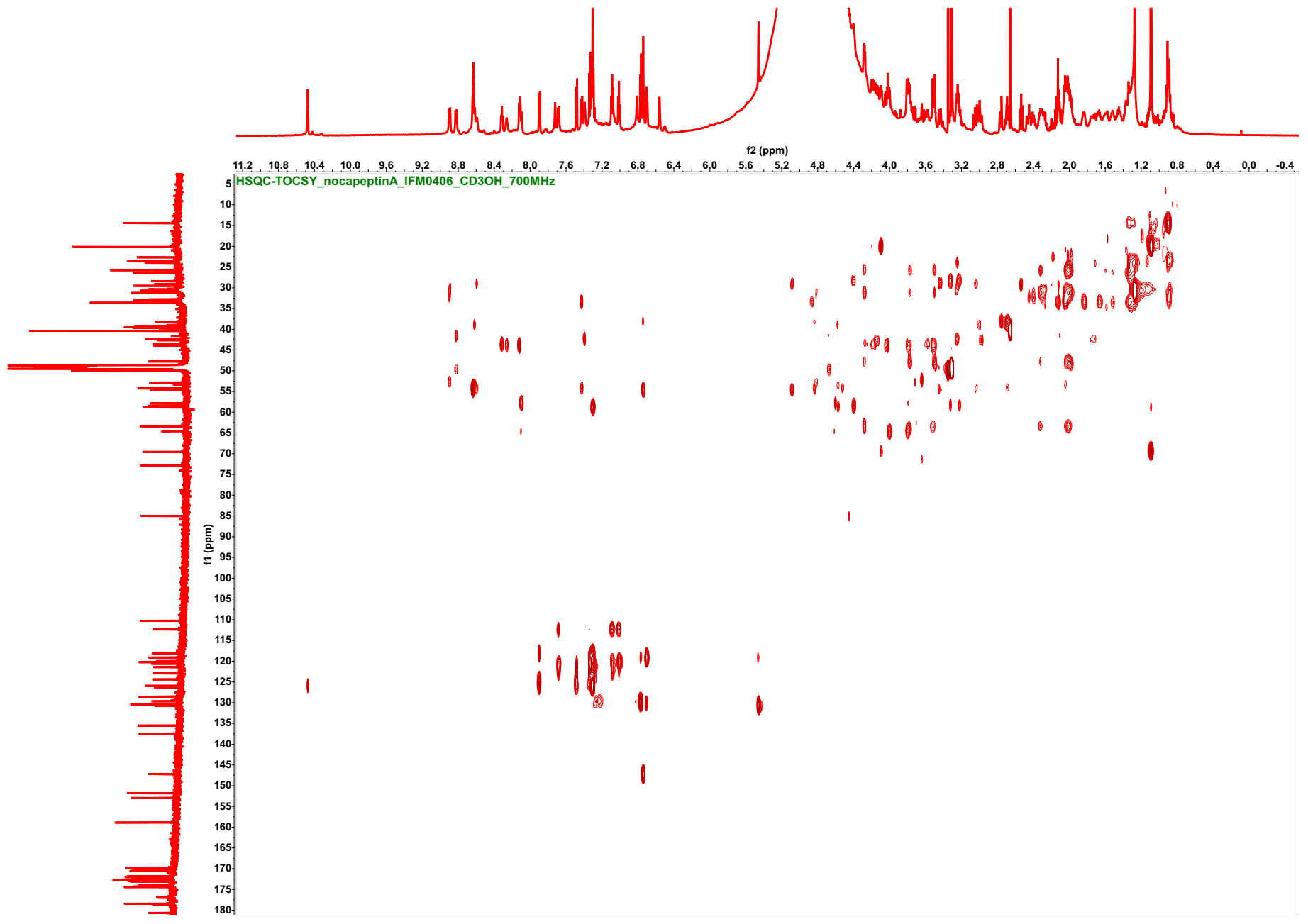

**Figure S27**. ^1^H-^13^C HSQC-TOCSY spectrum of nocapeptin A (**1**) (*d*_3_-CH_3_OH/H_2_O (96:4), 700/176 MHz)

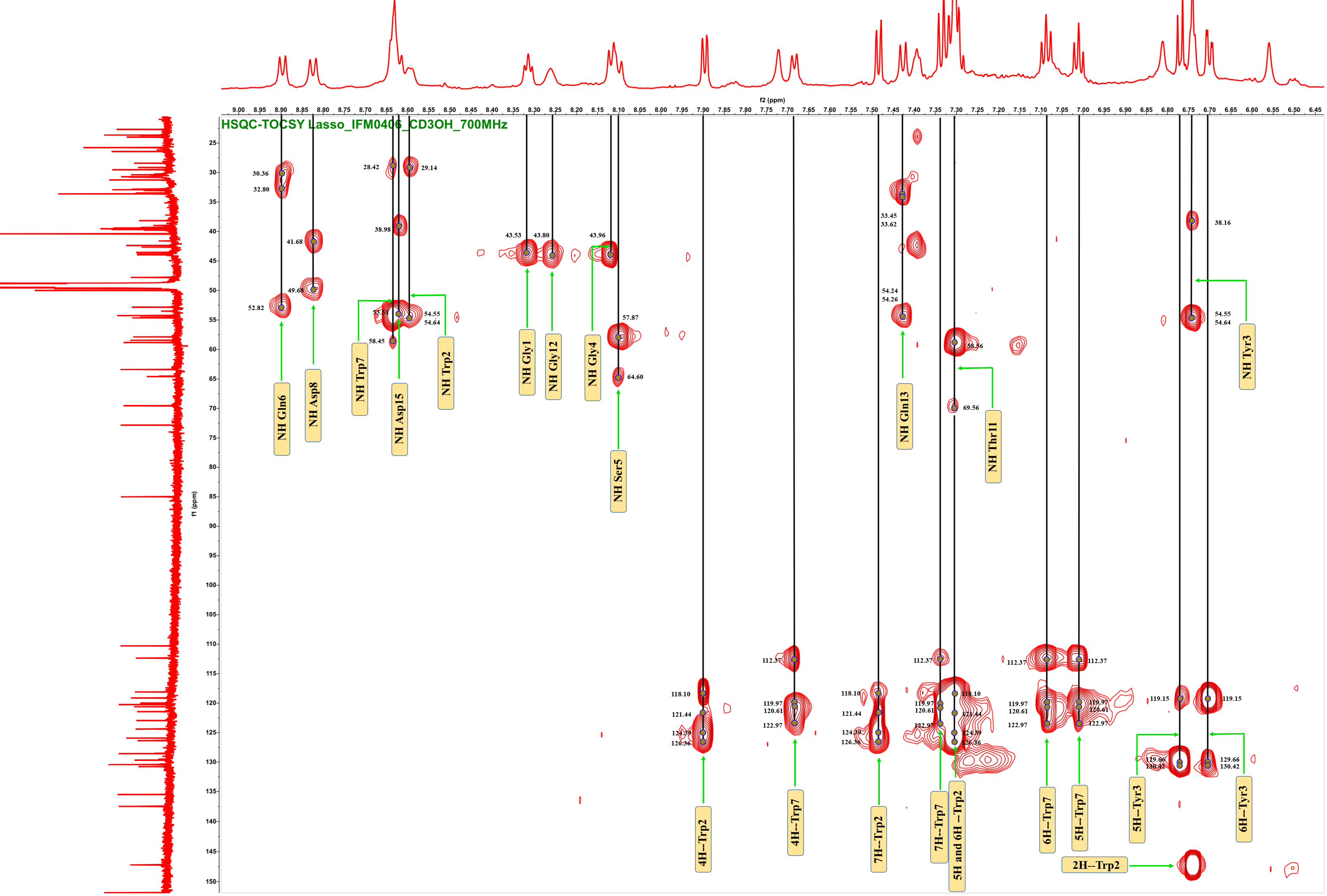

**Figure S27A**. Annotated ^1^H-^13^C HSQC-TOCSY spectrum of nocapeptin A (**1**)

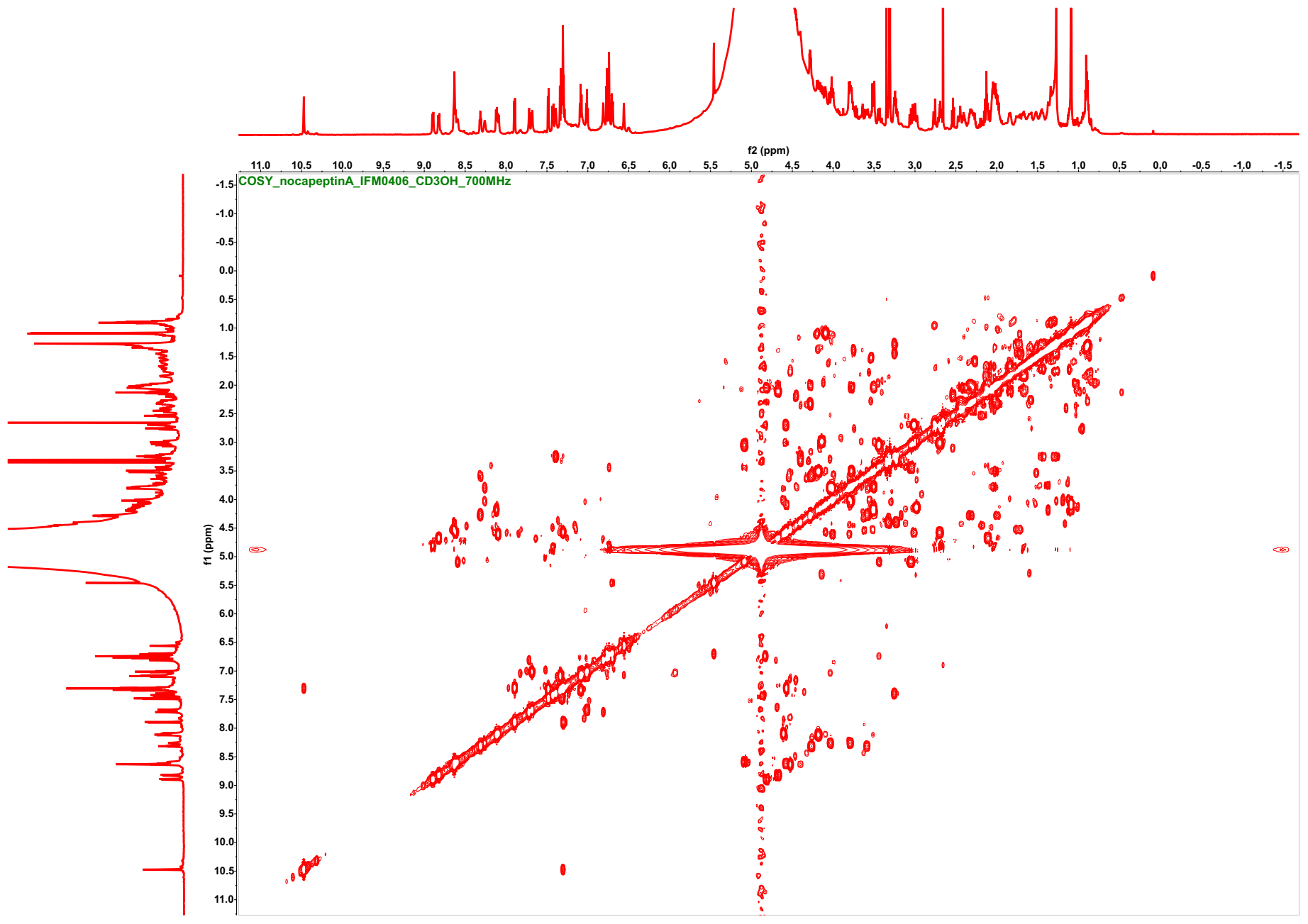

**Figure S28**. ^1^H-^1^H COSY spectrum of nocapeptin A (**1**) (*d*_3_-CH_3_OH/H_2_O (96:4), 700 MHz)

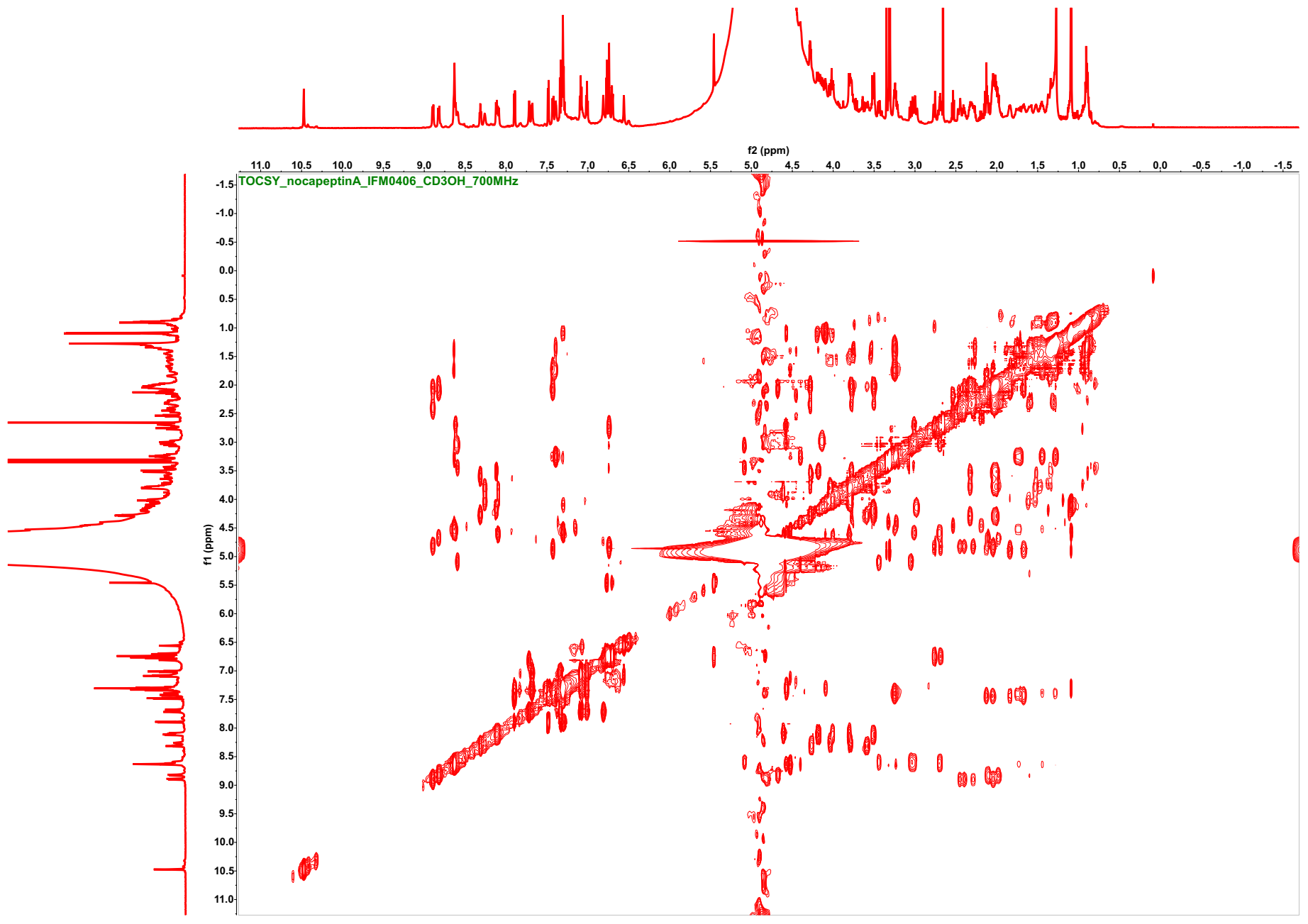

**Figure S29**. ^1^H-^1^H TOCSY spectrum of nocapeptin A (**1**) (*d*_3_-CH_3_OH/H_2_O (96:4), 700 MHz)

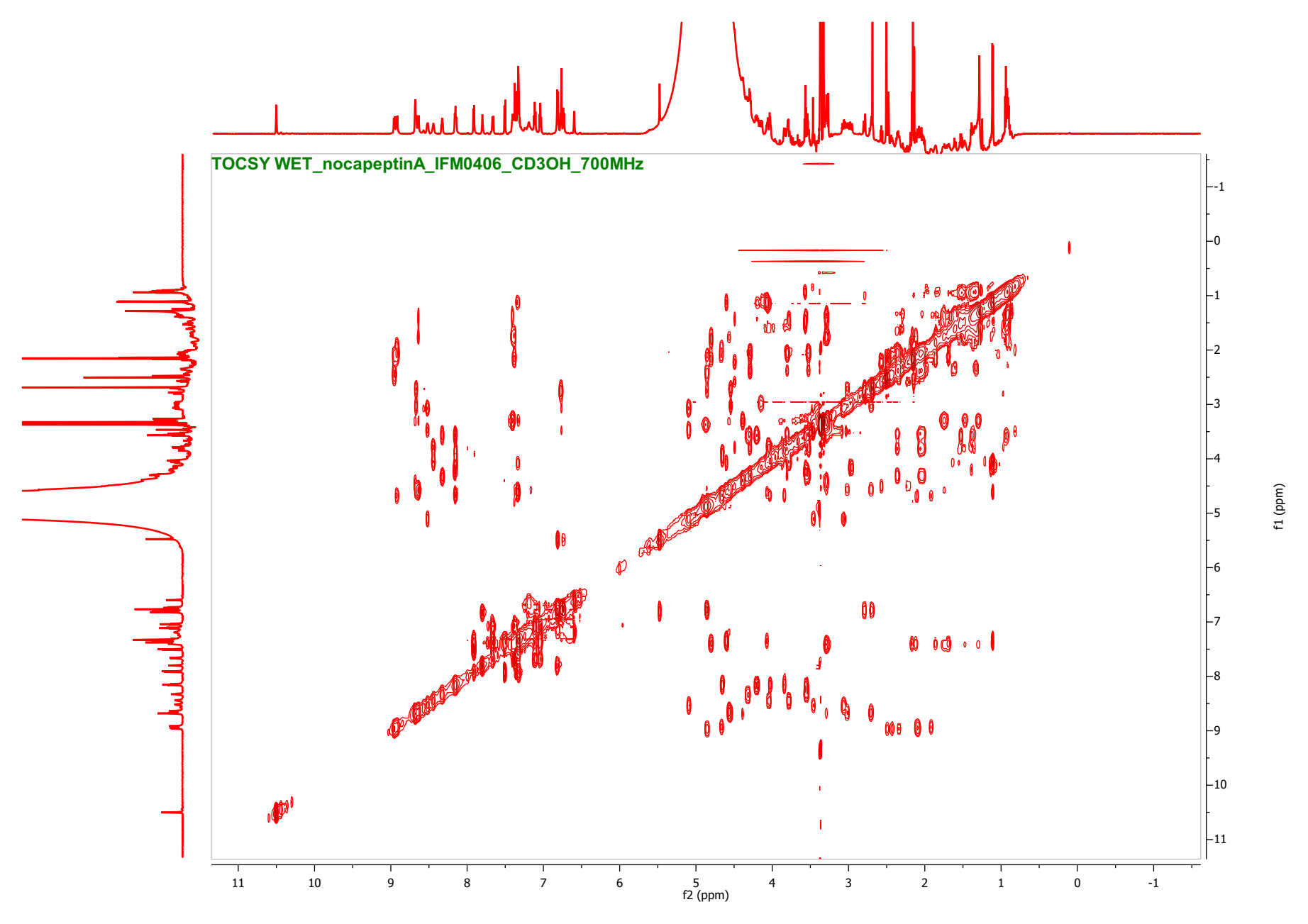

**Figure S29A**. ^1^H-^1^H TOCSY spectrum of nocapeptin A (**1**) with WET solvent suppression (d_3_-CH_3_OH/H_2_O (96:4), 700 MHz)

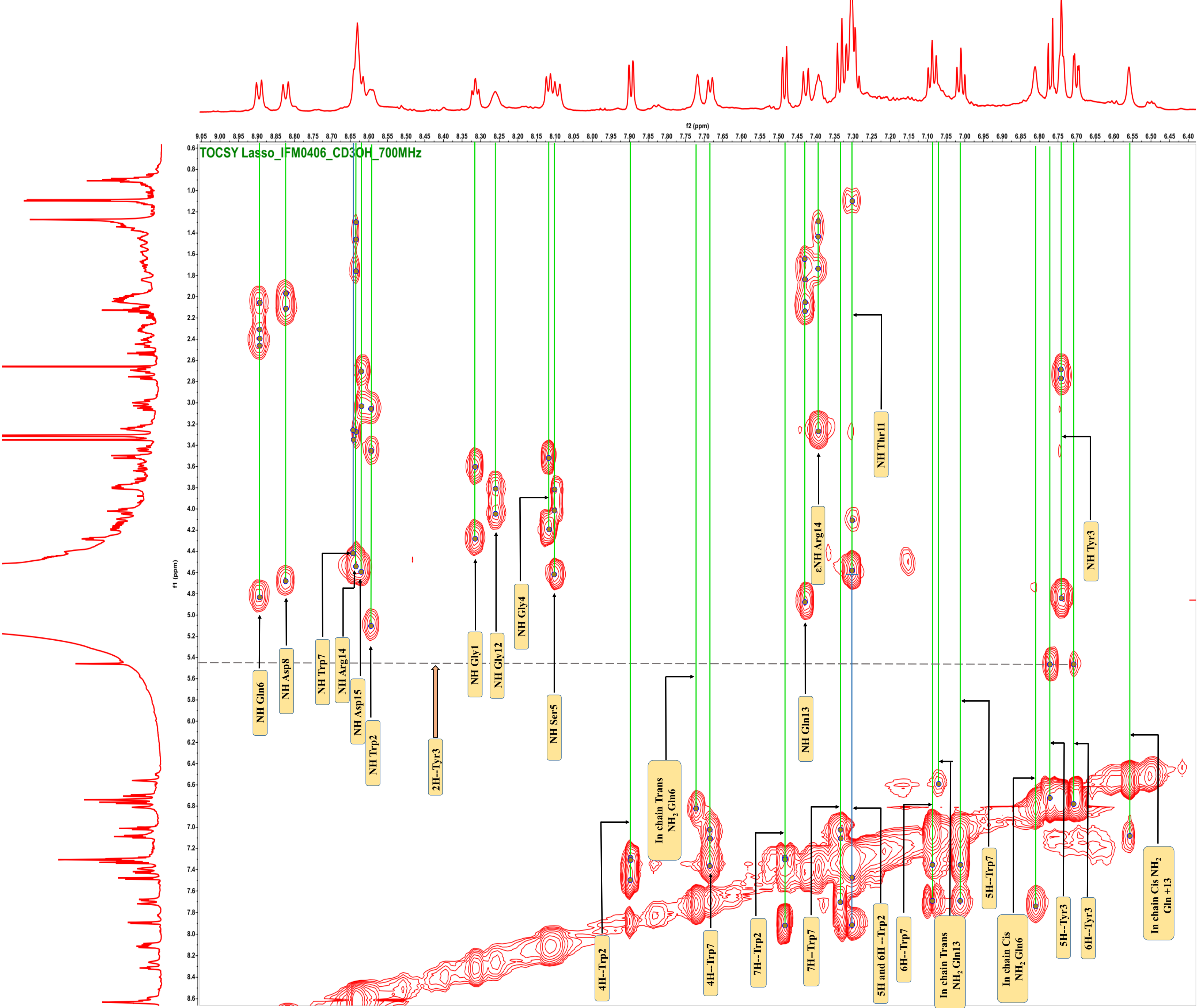

**Figure S29B**. Annotated ^1^H-^1^H TOCSY spectrum of nocapeptin A (**1**)

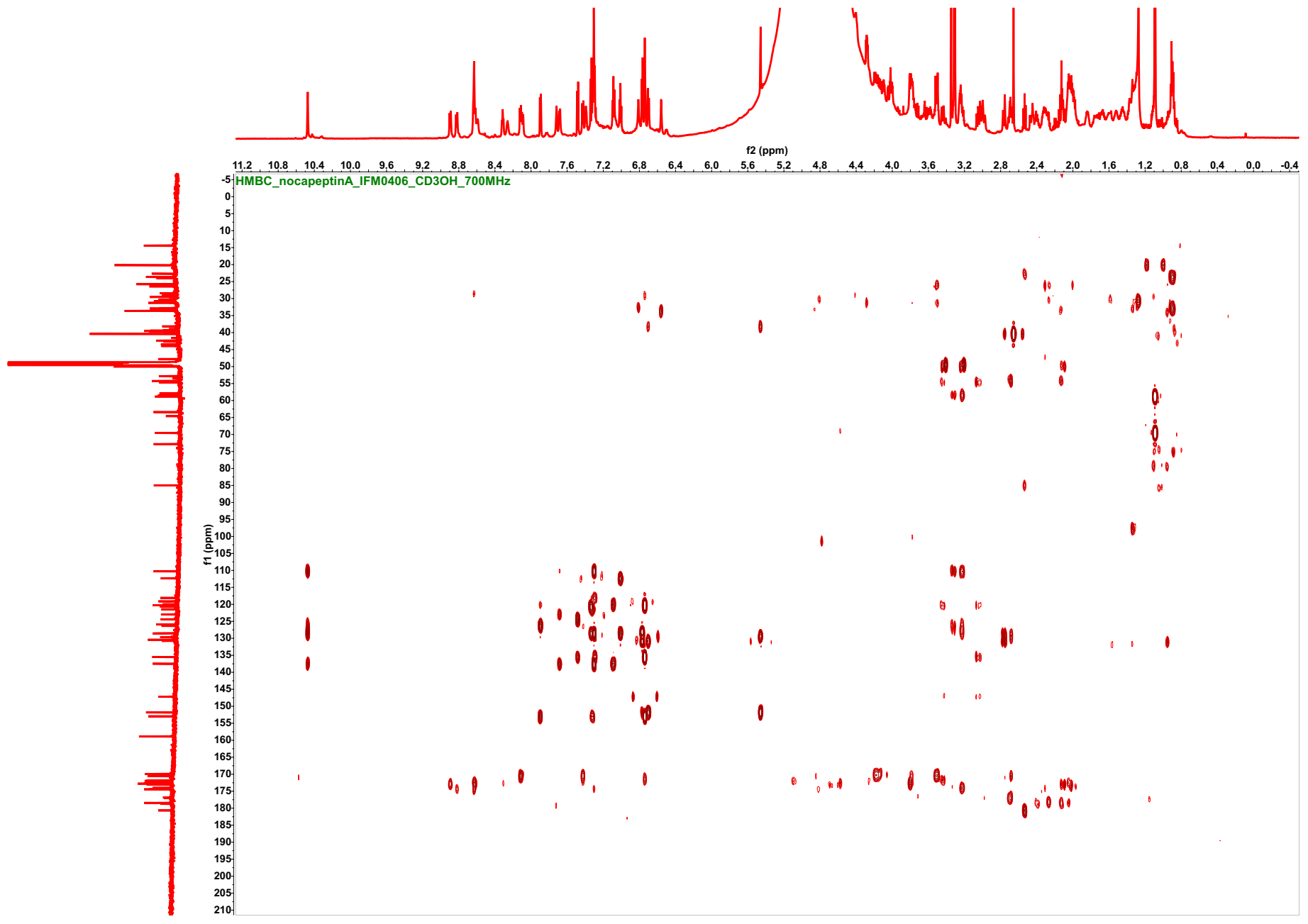

**Figure S30**. ^1^H-^13^C HMBC spectrum of nocapeptin A (**1**) (*d*_3_-CH_3_OH/H_2_O (96:4), 700/176 MHz)

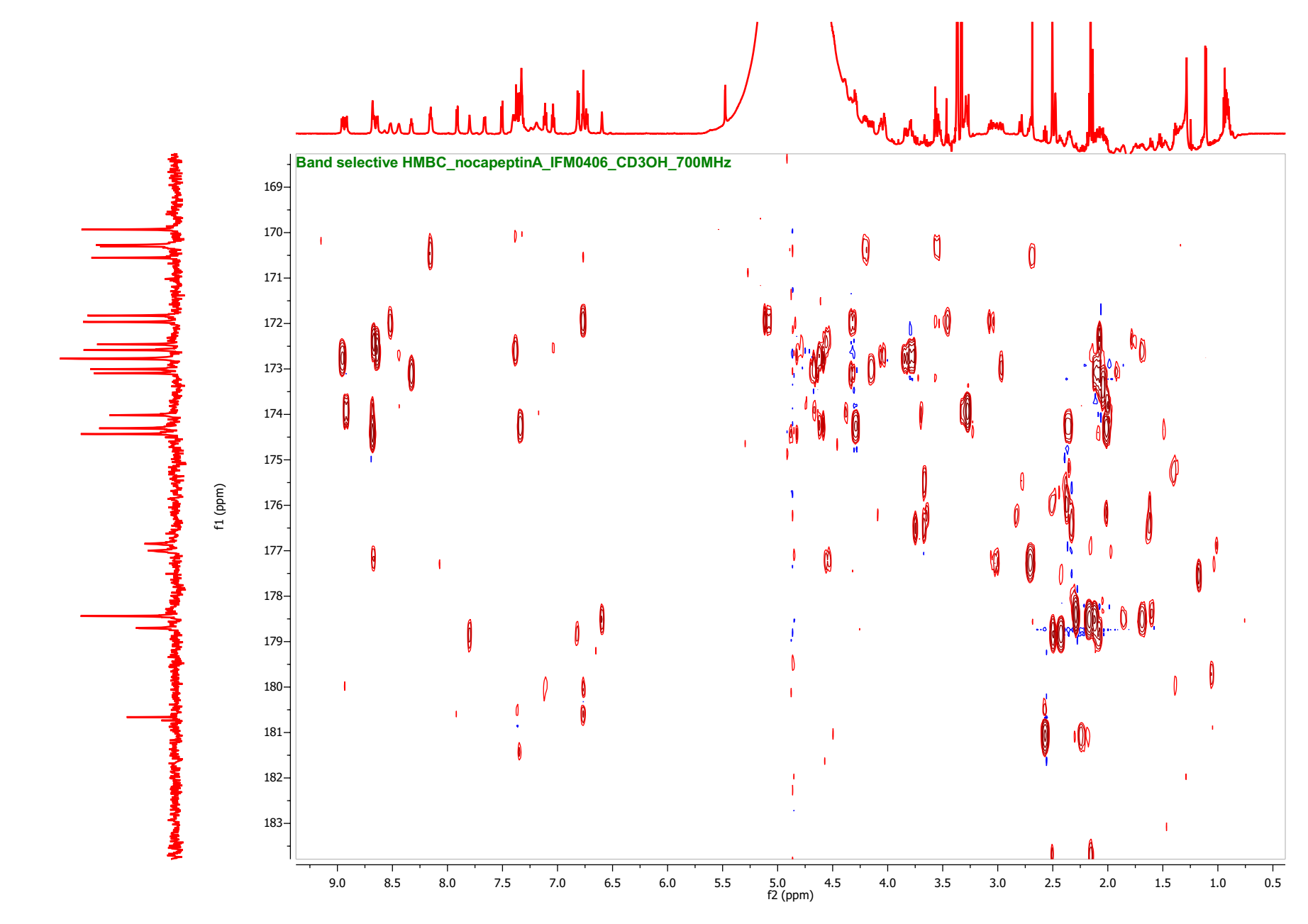

**Figure S30A**. ^1^H-^13^C Band-selective HMBC spectrum of nocapeptin A (**1**) (*d*_3_-CH_3_OH/H_2_O (96:4), 700/176 MHz)

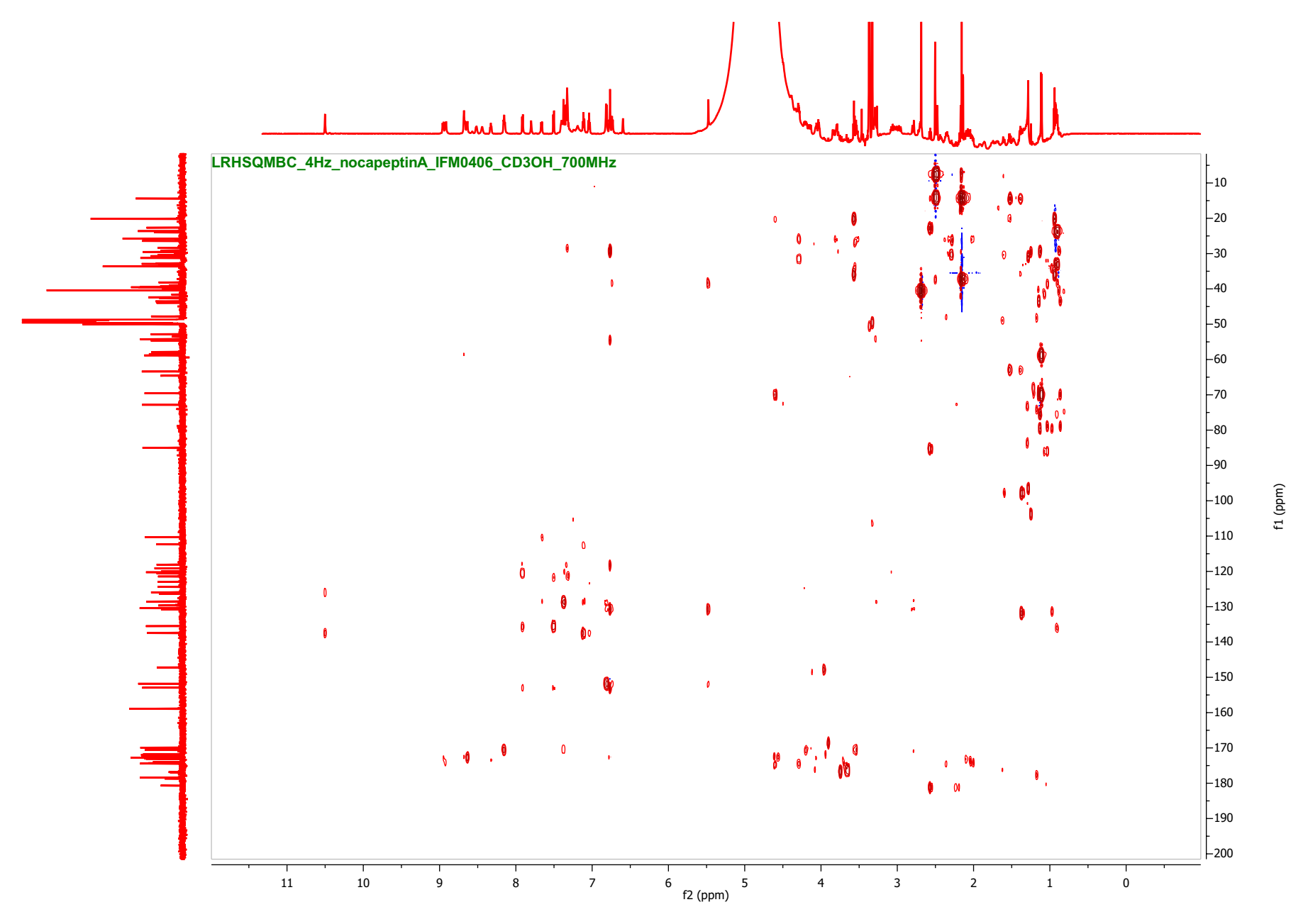

**Figure S31**. ^1^H-^13^C LR-HSQMBC spectrum of nocapeptin A (**1**) (*d*_3_-CH_3_OH/H_2_O (96:4), 700/176 MHz)

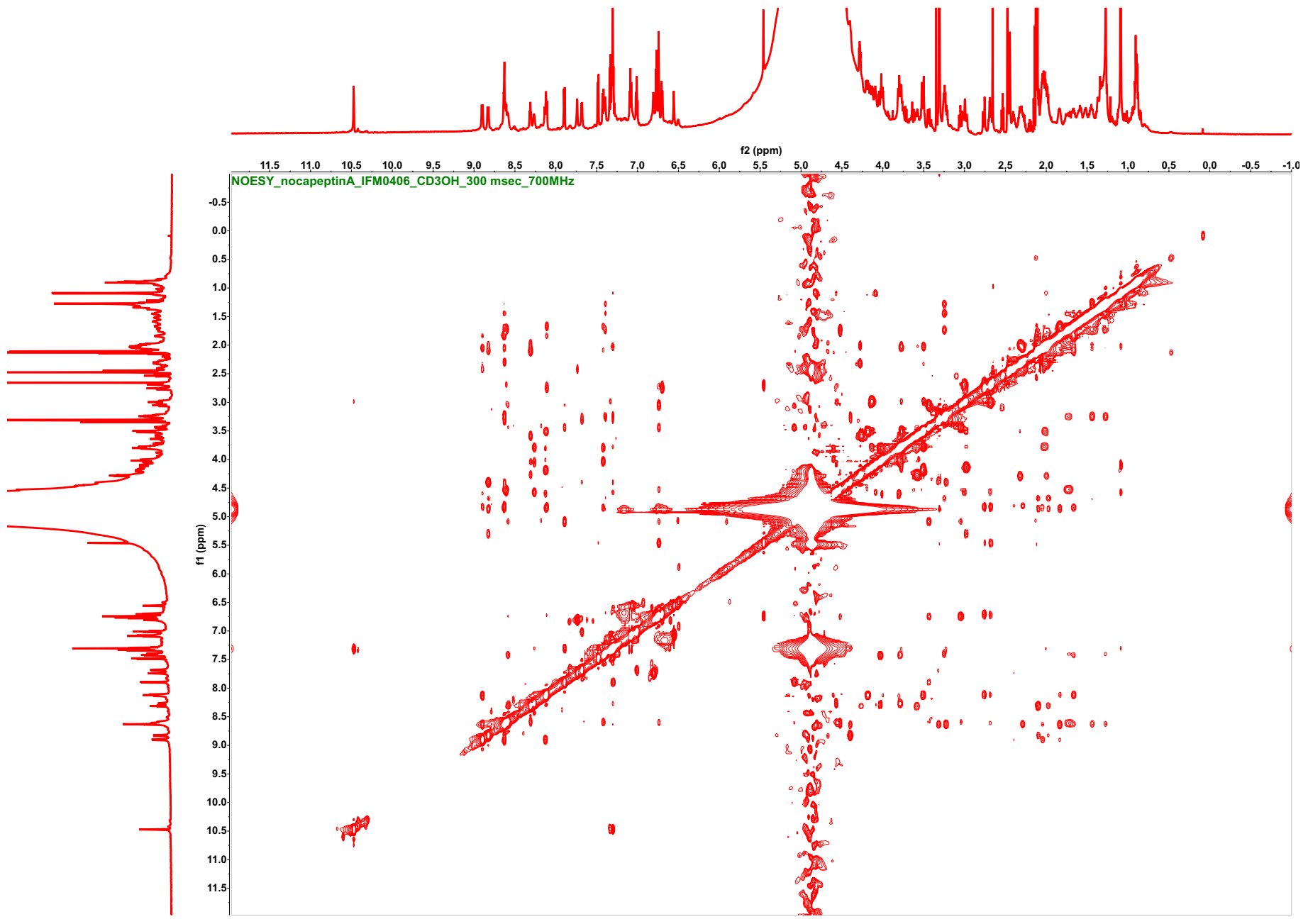

**Figure S32**. ^1^H-^1^H NOESY spectrum of nocapeptin A (**1**) (*d*_3_-CH_3_OH/H_2_O (96:4), 300 msec, 700 MHz)

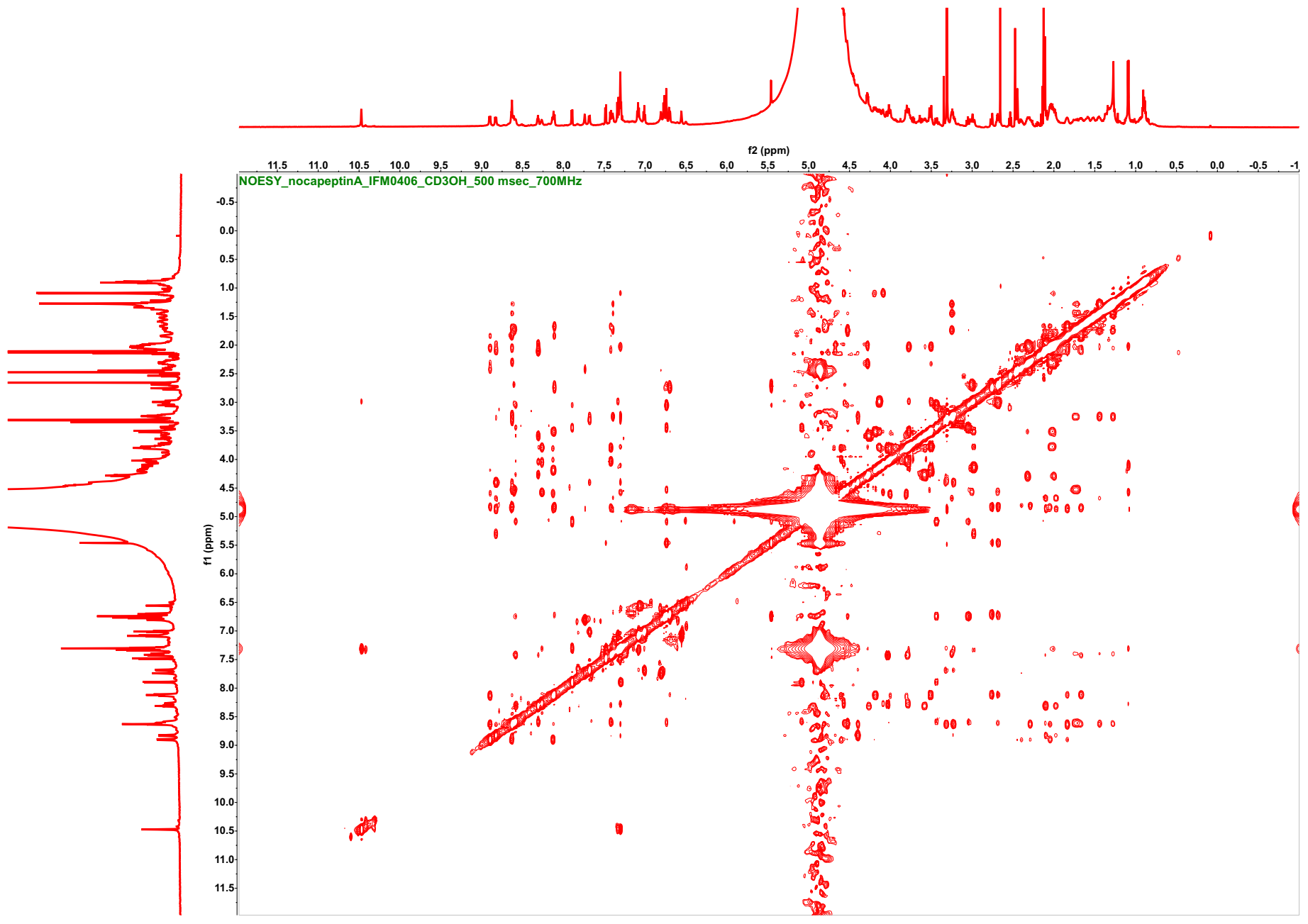

**Figure S33**. ^1^H-^1^H NOESY spectrum of nocapeptin A (**1**) (*d*_3_-CH_3_OH/H_2_O (96:4), 500 msec, 700 MHz)

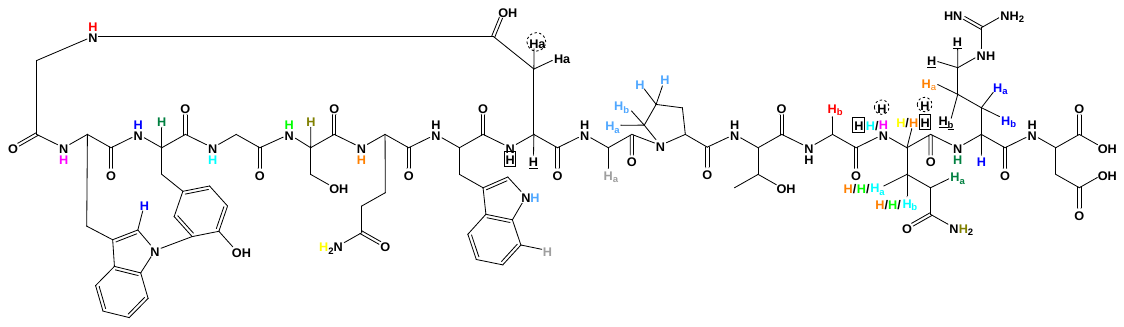

**Figure S33A**. NOE correlations, highlighting the threading property of nocapeptin A (**1**). Protons with similar patterns are in close proximity to each other

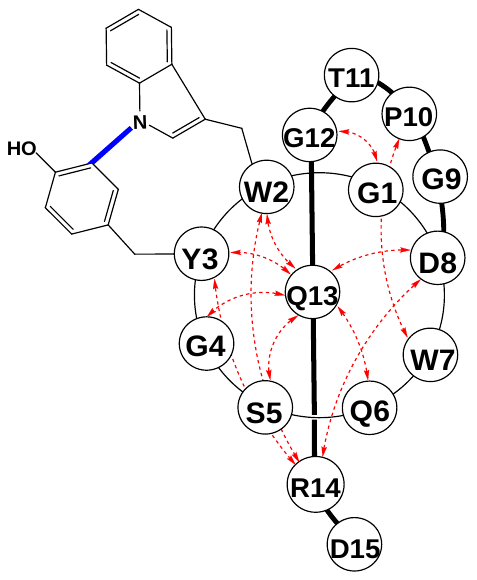

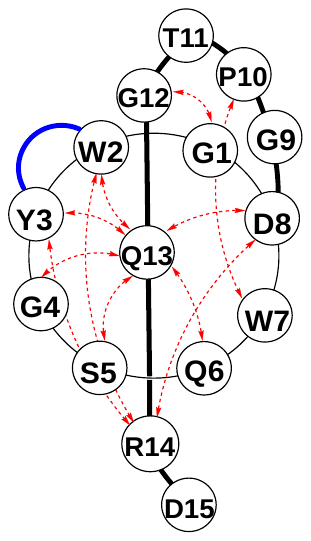

**Figure S33B**. Schematic representation of the lasso conformation of nocapeptin A (**1**) based on NOE correlations (red arrows)

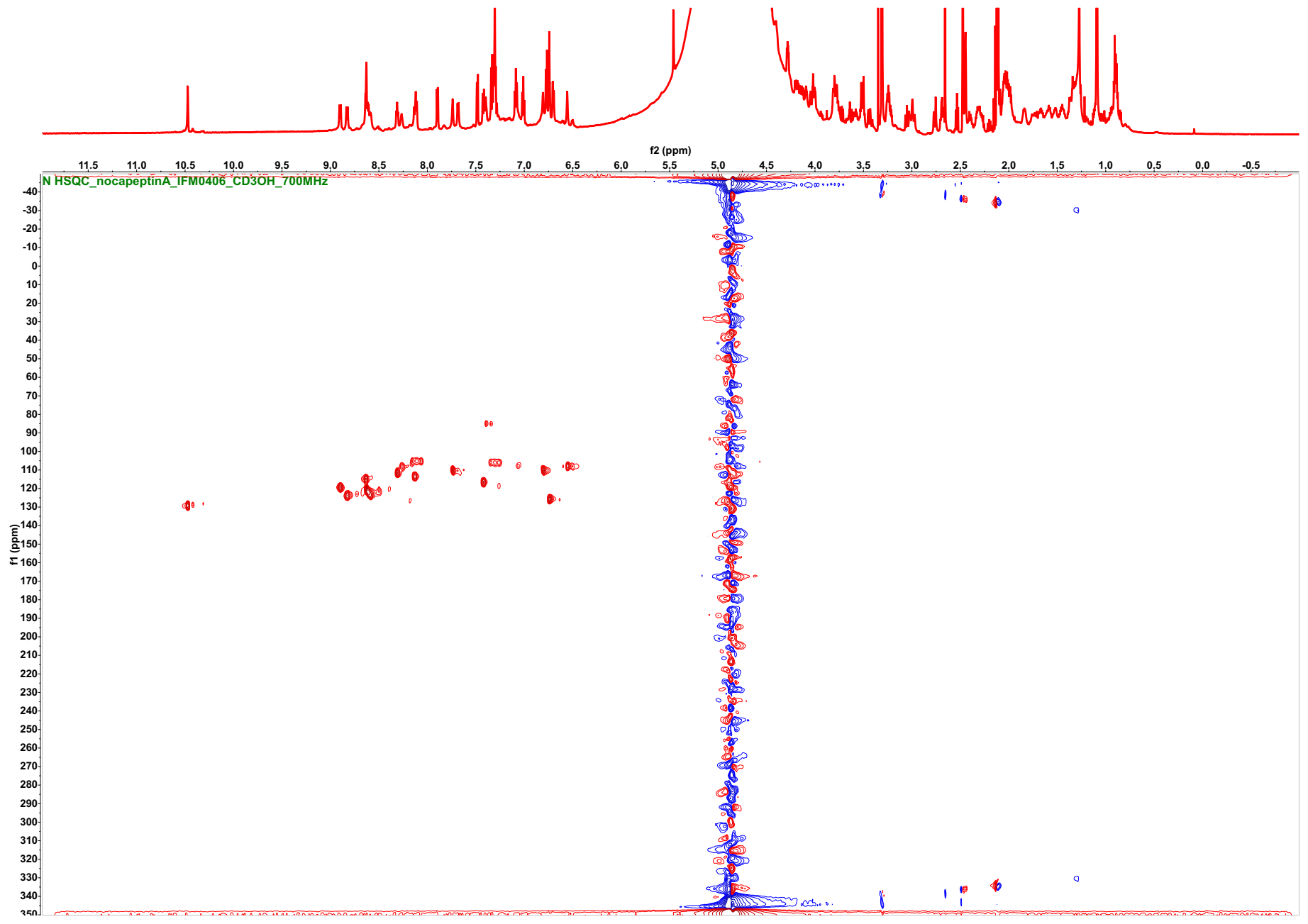

**Figure S34**. ^1^H-^15^N HSQC spectrum of nocapeptin A (**1**) (*d*_3_-CH_3_OH/H_2_O (96:4), 700/71 MHz)

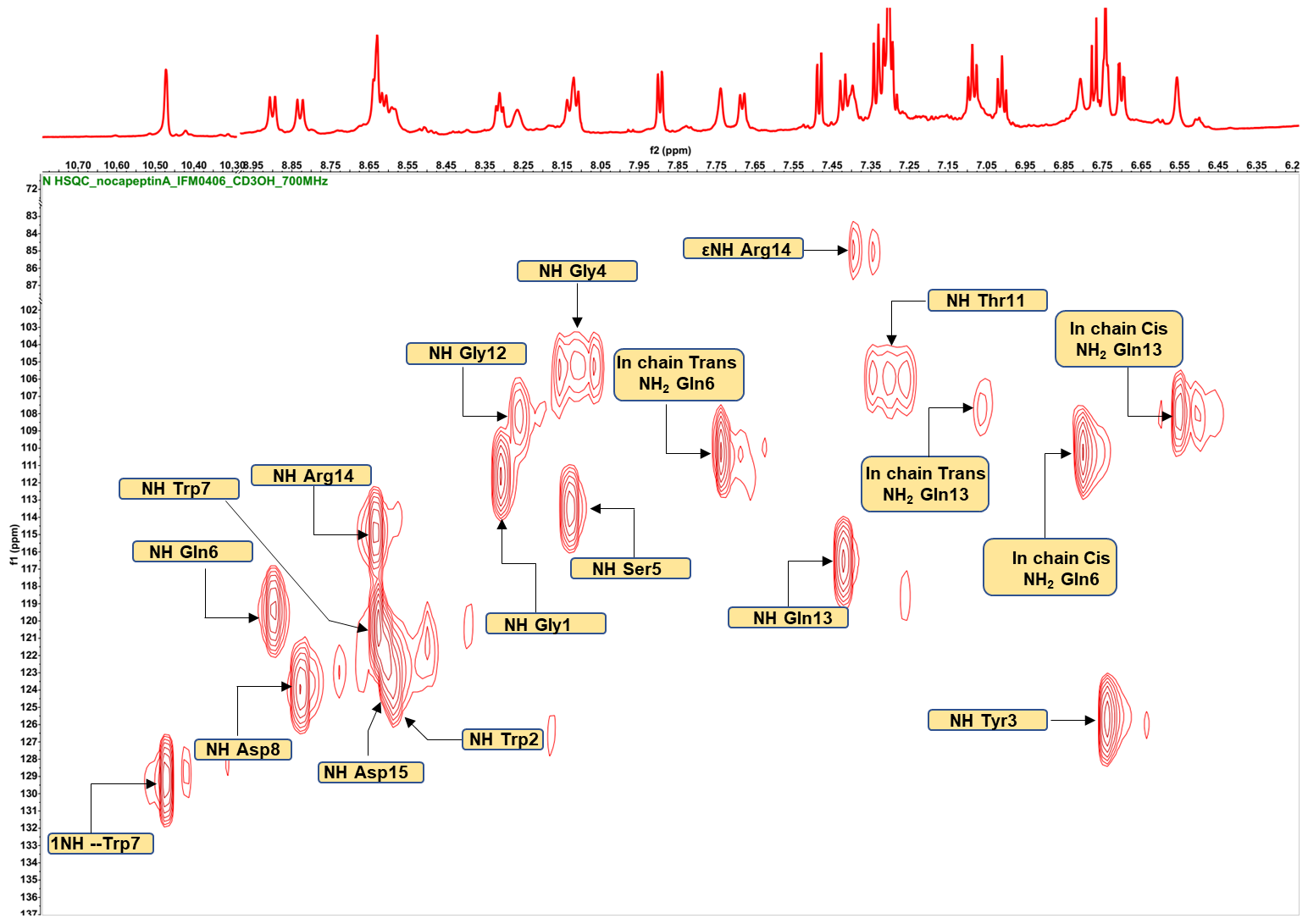

**Figure S34A**. Annotated ^1^H-^15^N HSQC spectrum of nocapeptin A (**1**)

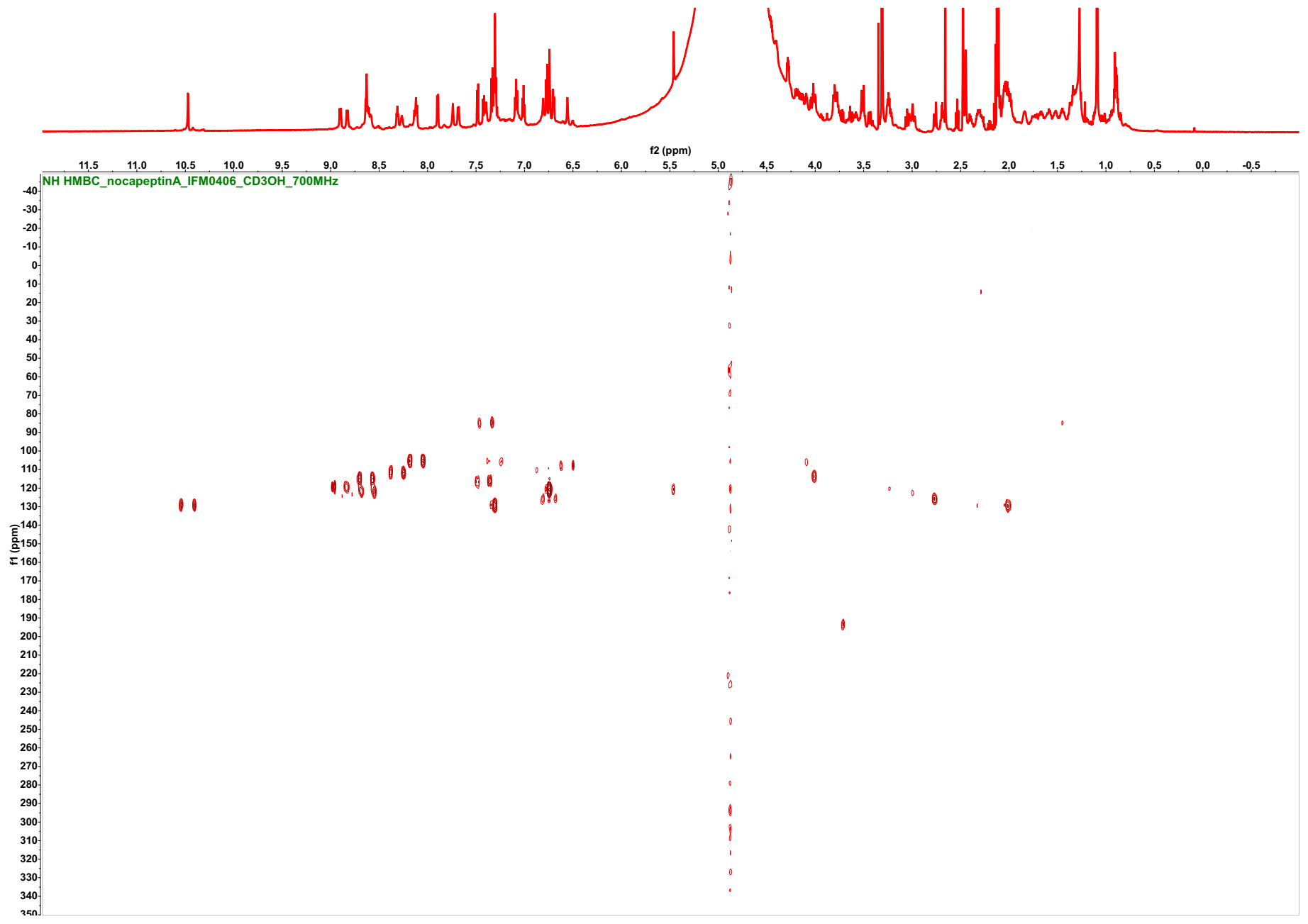

**Figure S35**. ^1^H-^15^N HMBC spectrum of nocapeptin A (**1**) (*d*_3_-CH_3_OH/H_2_O (96:4), 700/71 MHz)

**Table S8**. ^1^H, ^13^C, and ^15^N-NMR data of nocapeptin A (**1**) [*d*_3_-CH_3_OH/H_2_O (96:4); 700/176/71 MHz]. All chemical shifts are given in ppm.

| **Residue** | **Position** | **δ_C_/δ_N_** | **δ_H_, mult (*J* in Hz)** | **Residue** | **Position** | **δ_C_/δ_N_** | **δ_H_, mult (*J* in Hz)** |
| --- | --- | --- | --- | --- | --- | --- | --- |
| **Gly1** | CO | 171.82/171.96, C | --------- | **Trp7** | CO | 174.01, C | --------- |
|  | α | 43.53, CH_2_ | 3.59, dd (11.6, 5.0) + 4.27, m |  | α | 58.45, CH | 4.41, m |
|  | -NH- | 111.70 | 8.31, t (6.3) |  | β | 28.42, CH_2_ | 3.24, m + 3.32, m |
| **Trp2*** | CO | 171.82/171.96, C | --------- |  | 1(NH) | 129.20 | 10.47, s |
|  | α | 54.55/54.64, CH | 5.09, m |  | 2 | 125.93, CH | 7.30, overlapped |
|  | β | 29.14, CH_2_ | 3.05, m + 3.44, dm (13.4) |  | 3 | 110.28, C | --------- |
|  | 1(N) | 120.80 | --------- |  | 3a | 128.51/128.56, C | --------- |
|  | 2 | 147.20, CH | 6.74, overlapped |  | 4 | 119.97, CH | 7.68, d (7.7) |
|  | 3 | 120.25, C | --------- |  | 5 | 120.61, CH | 7.01, t (7.5, 7.4) |
|  | 3a | 135.51, C | --------- |  | 6 | 122.95, CH | 7.09, dd (7.7, 7.6) |
|  | 4 | 121.44, CH | 7.90, d (7.7) |  | 7 | 112.36, CH | 7.34, d (8.3) |
|  | 5 | 124.39, CH | 7.30, overlapped |  | 7a | 137.49, C | --------- |
|  | 6 | 126.36, CH | 7.32, overlapped |  | -NH- | 120.40 | 8.64, overlapped |
|  | 7 | 118.10, CH | 7.48, d (7.6) | **Asp8** | CO | 173.00, C | --------- |
|  | 7a | 152.95, C | --------- |  | α | 49.68; CH | 4.67, masked |
|  | -NH- | 123.70 | 8.59, m |  | β | 41.60, CH_2_ | 1.99, m + 2.11, m |
| **Tyr3*** | CO | 170.55, C | --------- |  | γ | 173.09, C | --------- |
|  | α | 54.55/54.64, CH | 4.84, masked |  | -NH- | 123.99 | 8.82, d (9,8) |
|  | β | 38.16, CH_2_ | 2.69, m + 2.76, m | **Gly9** | CO | n.d. | --------- |
|  | 1 | 128.51/128.56, C | --------- |  | α | 42.66, CH_2_ | 2.99, m + 4.15, dd (16.3, 8.3) |
|  | 2 | 130.77, CH | 5.46, d (2) |  | -NH- | n.d. | 5.30, masked |
|  | 3 | 130.42, C | --------- | **Pro10** | CO | 174.30, C | --------- |
|  | 4 | 151.81, C | --------- |  | α | 63.40, CH | 4.28, m |
|  | 5 | 119.12, CH | 6.77, d (8.3) |  | β | 31.25, CH_2_ | 2.00, m + 2.32, m |
|  | 6 | 129.66, CH | 6.70, dd (8.3, 2) |  | γ | 25.76, CH_2_ | 2.03, m |
|  | OH | --------- | n.d. |  | δ | 47.78, CH_2_ | 3.51, m + 3.78, m |
|  | -NH- | 125.78 | 6.74, m |  | =N- | 129.72 | --------- |
| **Gly4** | CO | 170.27/170.30, C | --------- | **Thr11** | CO | 172.77, C | --------- |
|  | α | 43.96, CH_2_ | 3.51, m + 4.18, dd (17.4, 7.8) |  | α | 58.82, CH | 4.58, masked |
|  | -NH- | 105.43 | 8.12, d (7.8) |  | β | 69.56, CH | 4.09, m |
| **Ser5** | CO | 172.77, C | --------- |  | γ | 20.16, CH_3_ | 1.09, d (6.3) |
|  | α | 57.87, CH | 4.61, masked |  | OH | --------- | n.d. |
|  | β | 64.50/64.60, CH_2_ | 3.80, m + 4.01, dd (11.3, 4.1) |  | -NH- | 106.11 | 7.31, m |
|  | OH | --------- | n.d. | **Gly12** | CO | 170.27/170.30, C | --------- |
|  | -NH- | 113.60 | 8.09, d (9.3) |  | α | 43.80, CH_2_ | 3.80, m + 4.04, dd (17.6, 5.0) |
| **Gln6** | CO | 174.43, C | --------- |  | -NH- | 108.21 | 8.26, br s |
|  | α | 52.83, CH | 4.82, masked | **Gln13** | CO | 172.58, C | --------- |
|  | β | 30.36, CH_2_ | 2.04, m + 2.29, m |  | α | 54.24/54.26, CH | 4.87, m |
|  | γ | 32.80, CH_2_ | 2.40, m + 2.45, m |  | β | 33.45, CH_2_ | 1.66, m + 1.84, m |
|  | δ | 178.70, C | --------- |  | γ | 33.62, CH_2_ | 2.05, m + 2.12, m |
|  | -NH_2_ | 110.25 | 6.81, br s + 7.72, br s |  | δ | 178.44, C | --------- |
|  | -NH- | 119.40 | 8.89, d (9.5) |  | -NH_2_ | 107.92 | 6.56, br s+ 7.07, overlapped |
|  |  |  |  |  | -NH- | 116.58 | 7.43, d (9.1) |
| **Arg14** | CO | 172.46, C | --------- |  | | | |
|  | α | 54.24/54.26, CH | 4.53, masked |  | | | |
|  | β | 30.80, CH_2_ | 1.75, m |  |  |  |  |
|  | γ | 23.99, CH_2_ | 1.28, m + 1.44, m |  |  |  |  |
|  | δ | 42.37, CH_2_ | 3.25, m |  |  |  |  |
|  | εNH | 85.05 | 7.39, m |  |  |  |  |
|  | ζ | 158.90**, C | --------- |  |  |  |  |
|  | NHη1 | n.d. | n.d. |  |  |  |  |
|  | NH_2_η1 | n.d. | n.d. |  |  |  |  |
|  | -NH- | 114.90 | 8.64, overlapped |  |  |  |  |
| **Asp15** | CO_2_H | 176.85/177.00, C | --------- |  |  |  |  |
|  | α | 53.51, CH | 4.58, m |  |  |  |  |
|  | β | 38.98, CH_2_ | 2.70, m + 3.01, m |  |  |  |  |
|  | γ | 177.00/176.85, C | --------- |  |  |  |  |
|  | -NH- | 122.77 | 8.62, overlapped |  |  |  |  |
| * Residues involved in the C-N biaryl crosslink | | | |  |  |  |  |
| ** The chemical shift was assigned based on literature values.  masked => overlapped with the water signal | | | |  |  |  |  |

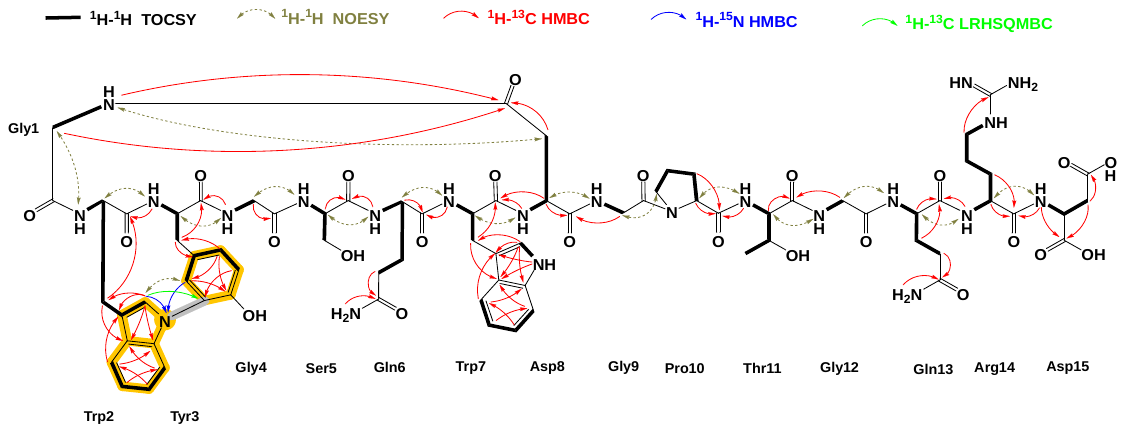

**Figure S36**. NMR key correlations of nocapeptin A (**1**)

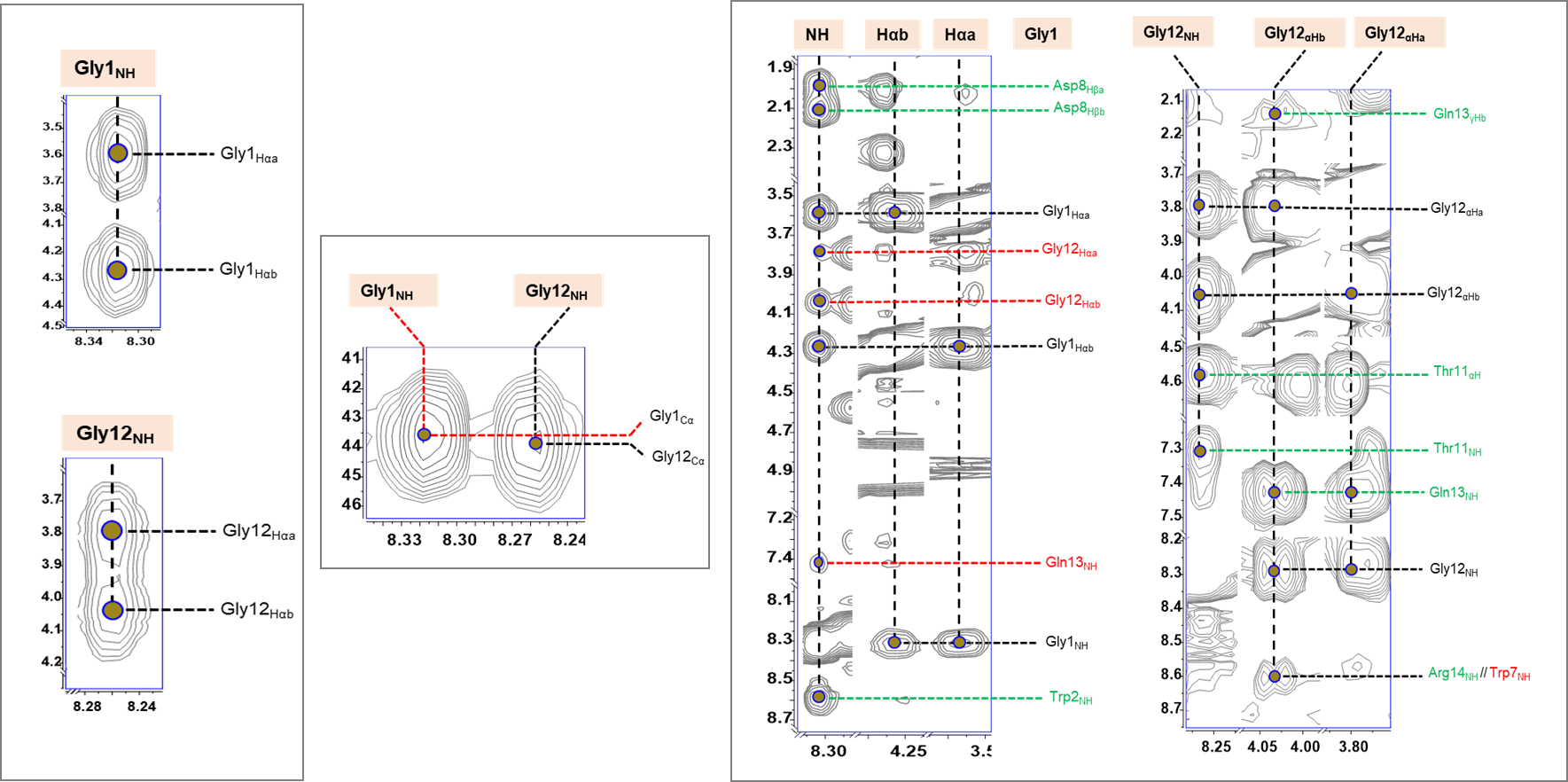

**Figure S37**. Annotated ^1^H-^1^H TOCSY (left), ^1^H-^13^C HSQC-TOCSY (middle) and ^1^H-^1^H NOESY (right) spectra of nocapeptin A (**1**) depicting residues **Gly1** and **Gly12**

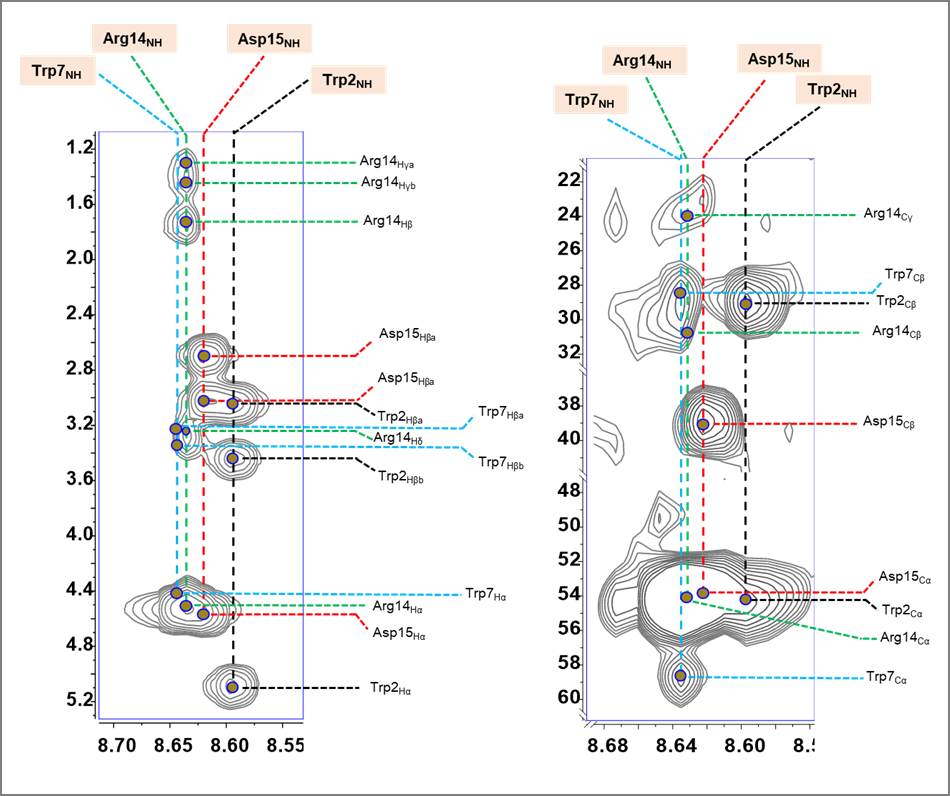

**Figure S38**. Annotated ^1^H-^1^H TOCSY (left), and ^1^H-^13^C HSQC-TOCSY (right) spectra of nocapeptin A (**1**) depicting residue **Trp2**

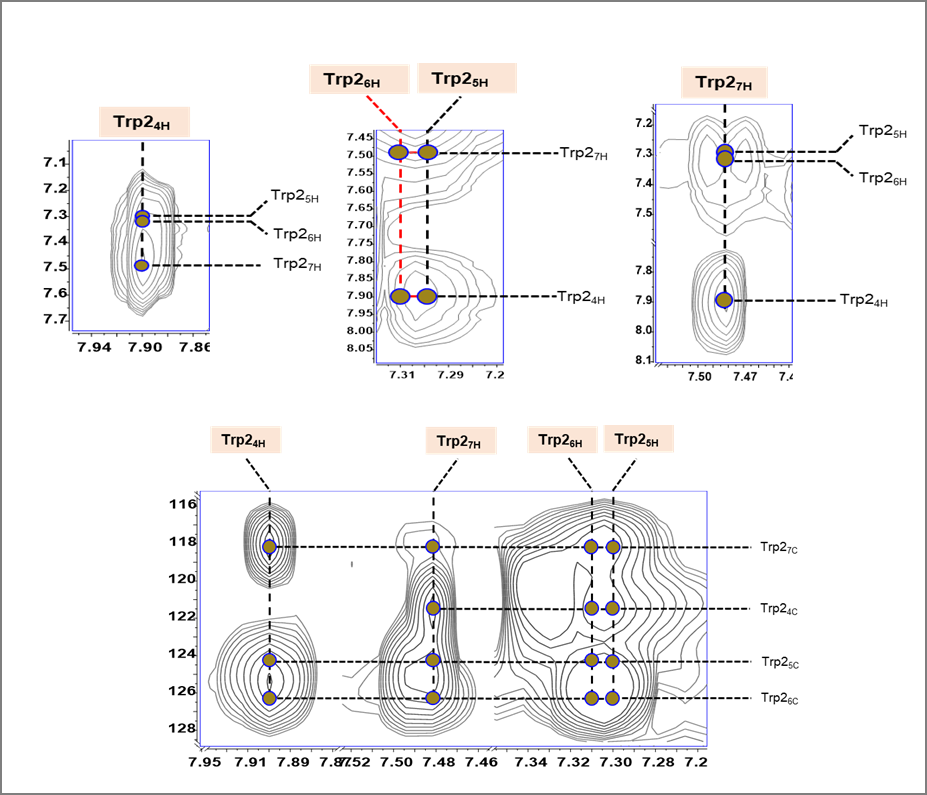

**Figure S38A**. Annotated ^1^H-^1^H TOCSY (upper), and ^1^H-^13^C HSQC-TOCSY (lower) spectra of nocapeptin A (**1**) depicting residue **Trp2**

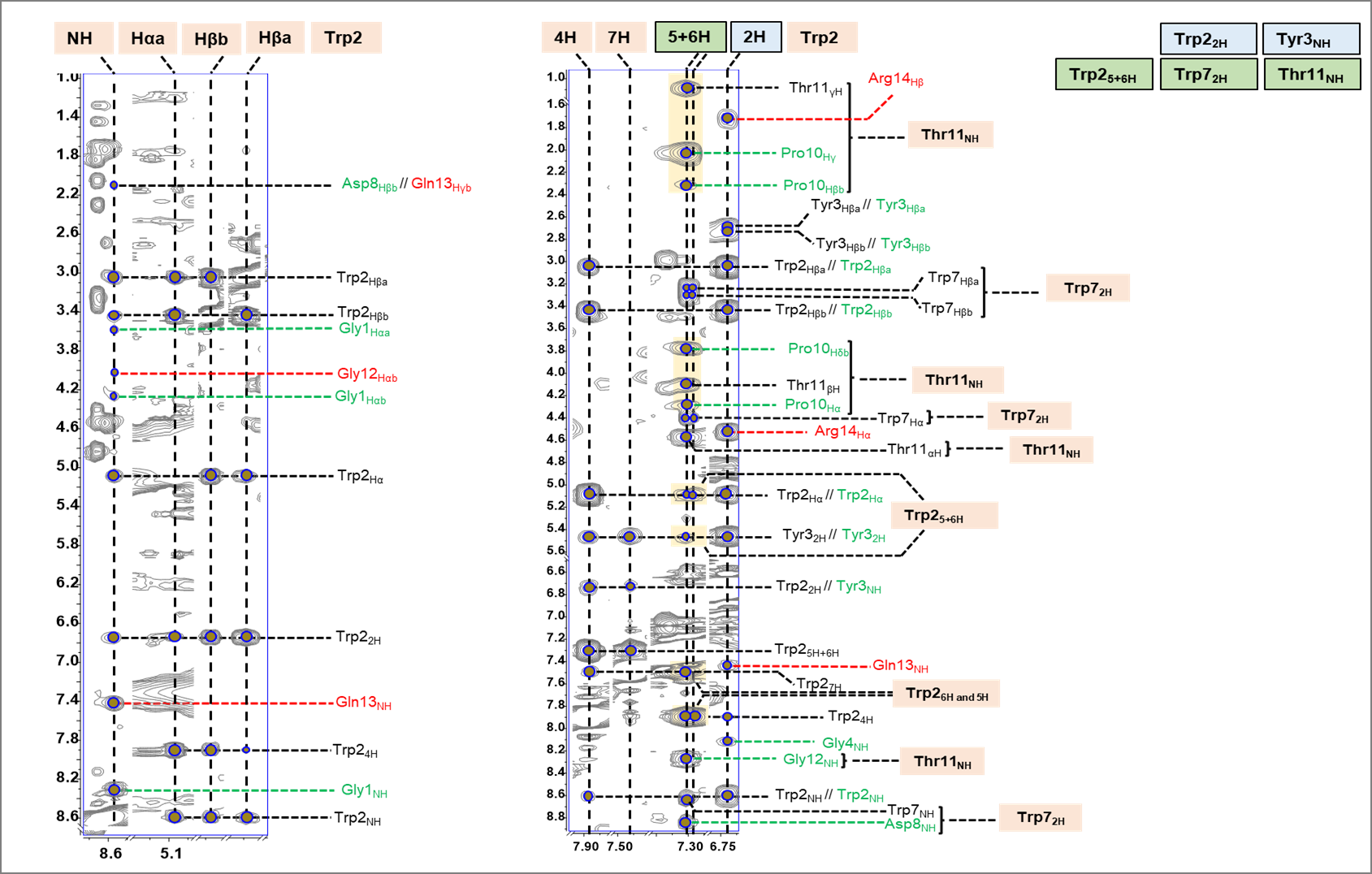

**Figure S38B**. Annotated ^1^H-^1^H NOESY spectrum of nocapeptin A (**1**) depicting residue **Trp2**

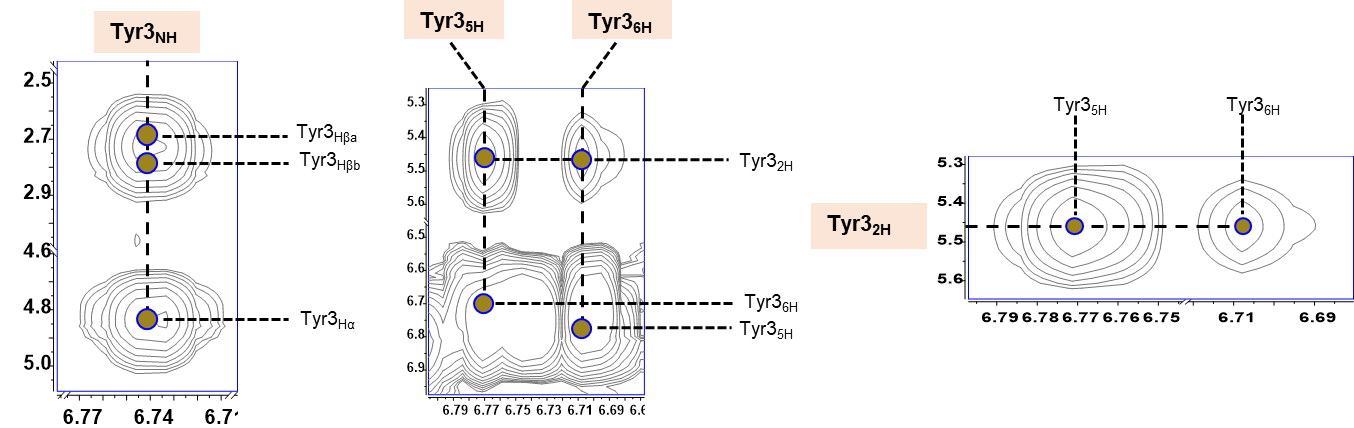

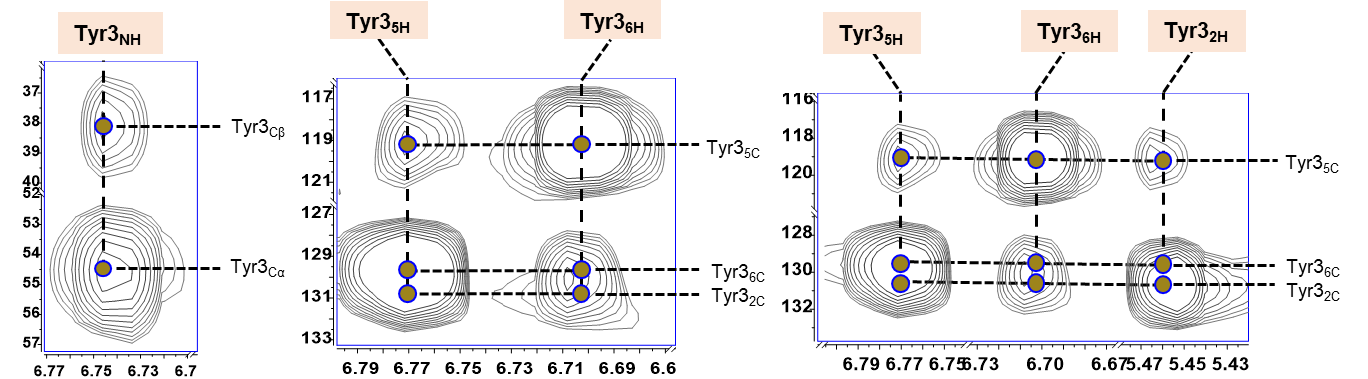

**Figure S39**. Annotated ^1^H-^1^H TOCSY (upper), and ^1^H-^13^C HSQC-TOCSY (lower) spectra of nocapeptin A (**1**) depicting residue **Tyr3**

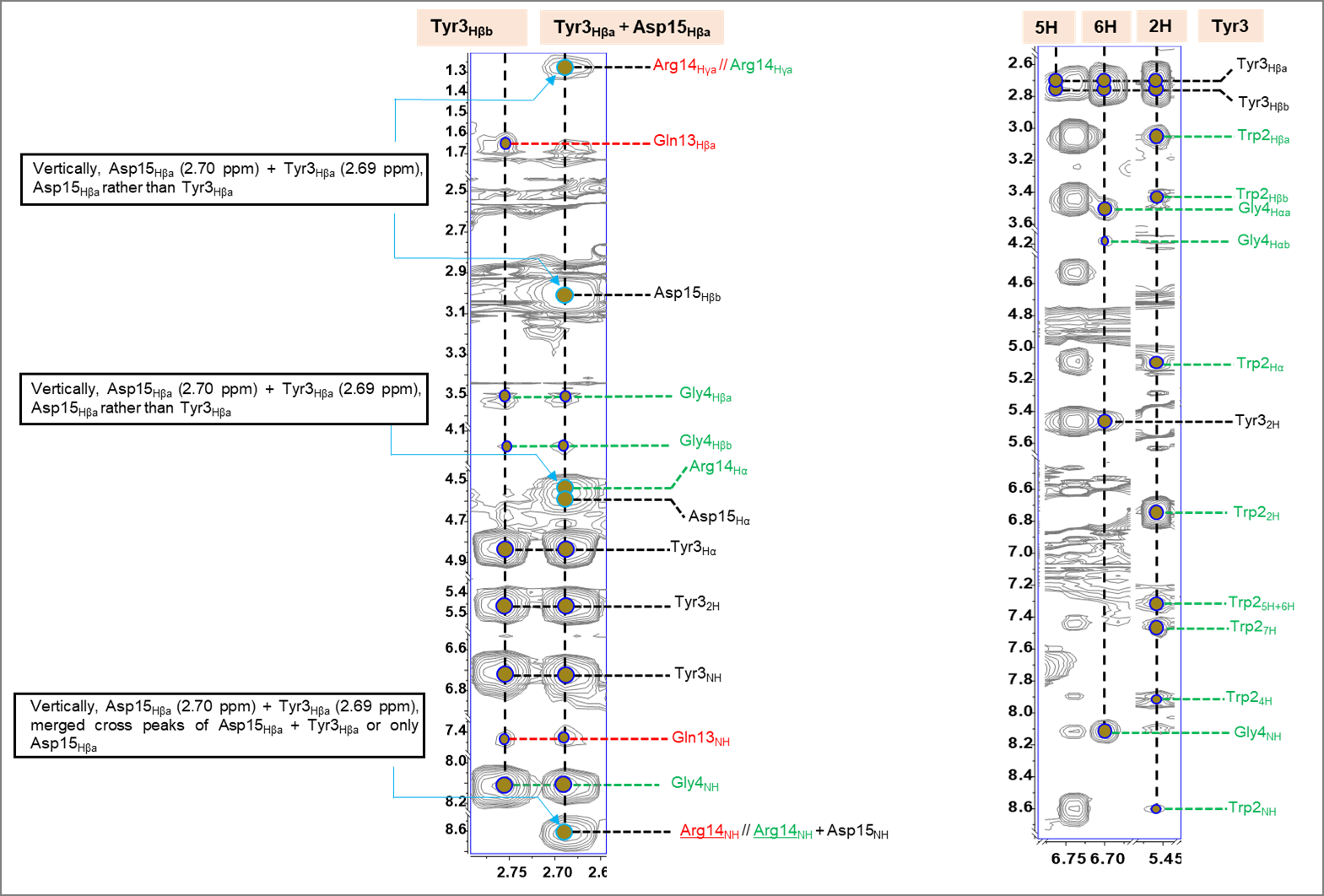

**Figure S39A**. Annotated ^1^H-^1^H NOESY spectrum of nocapeptin A (**1**) depicting residue **Tyr3**

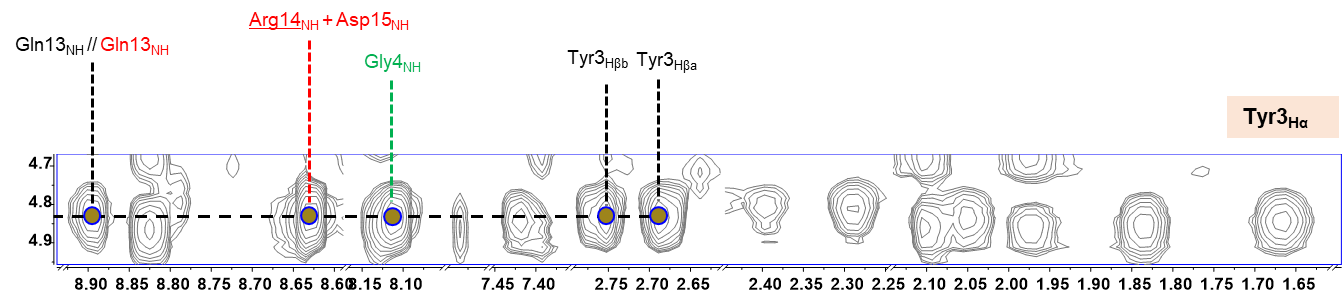

**Figure S39B**. Annotated ^1^H-^1^H NOESY spectrum of nocapeptin A (**1**) depicting residue **Tyr3**

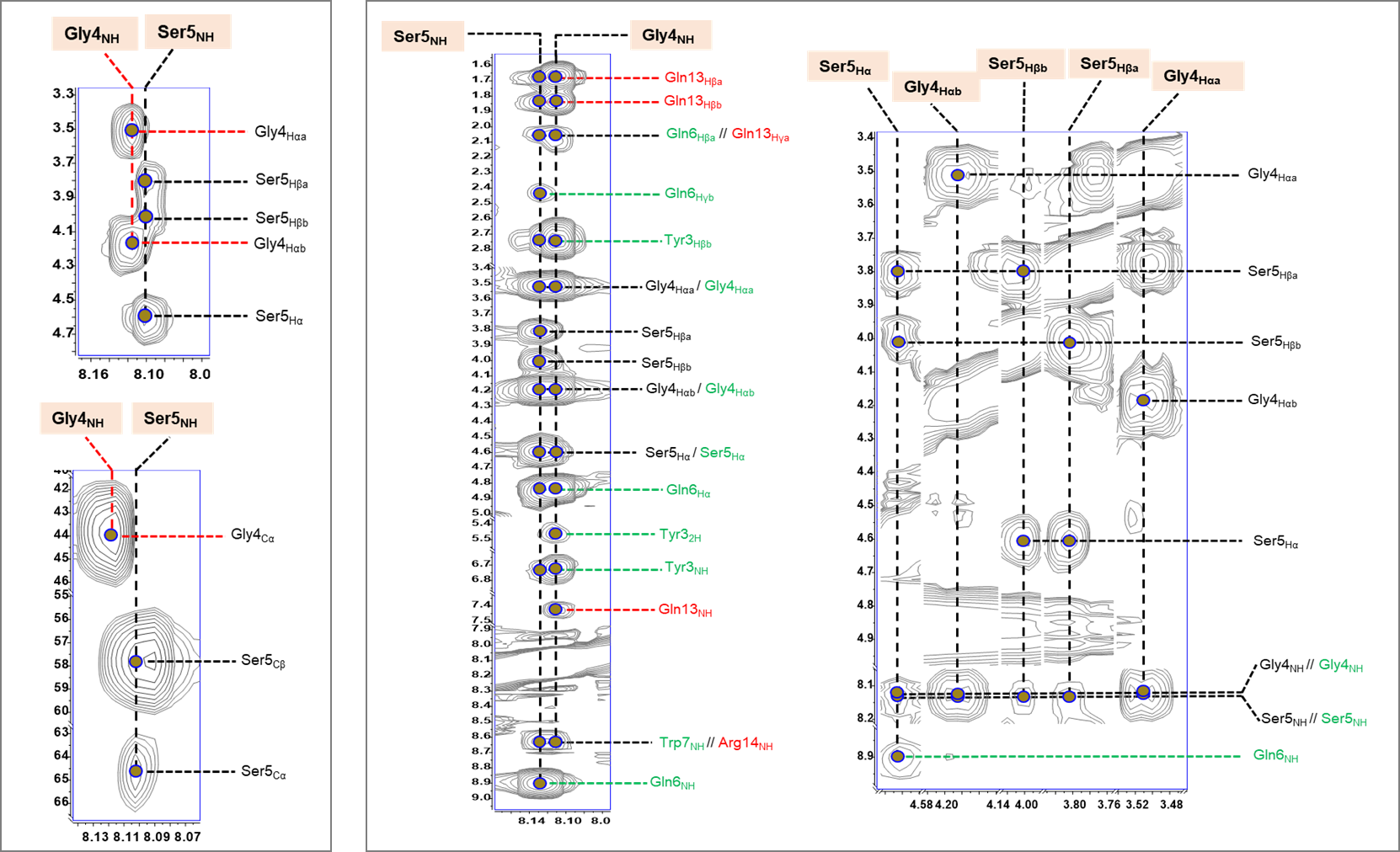

**Figure S40**. Annotated ^1^H-^1^H TOCSY, ^1^H-^13^C HSQC-TOCSY (left), and ^1^H-^1^H NOESY (right) spectra of nocapeptin A (**1**) depicting residues **Gly4** and **Ser5**

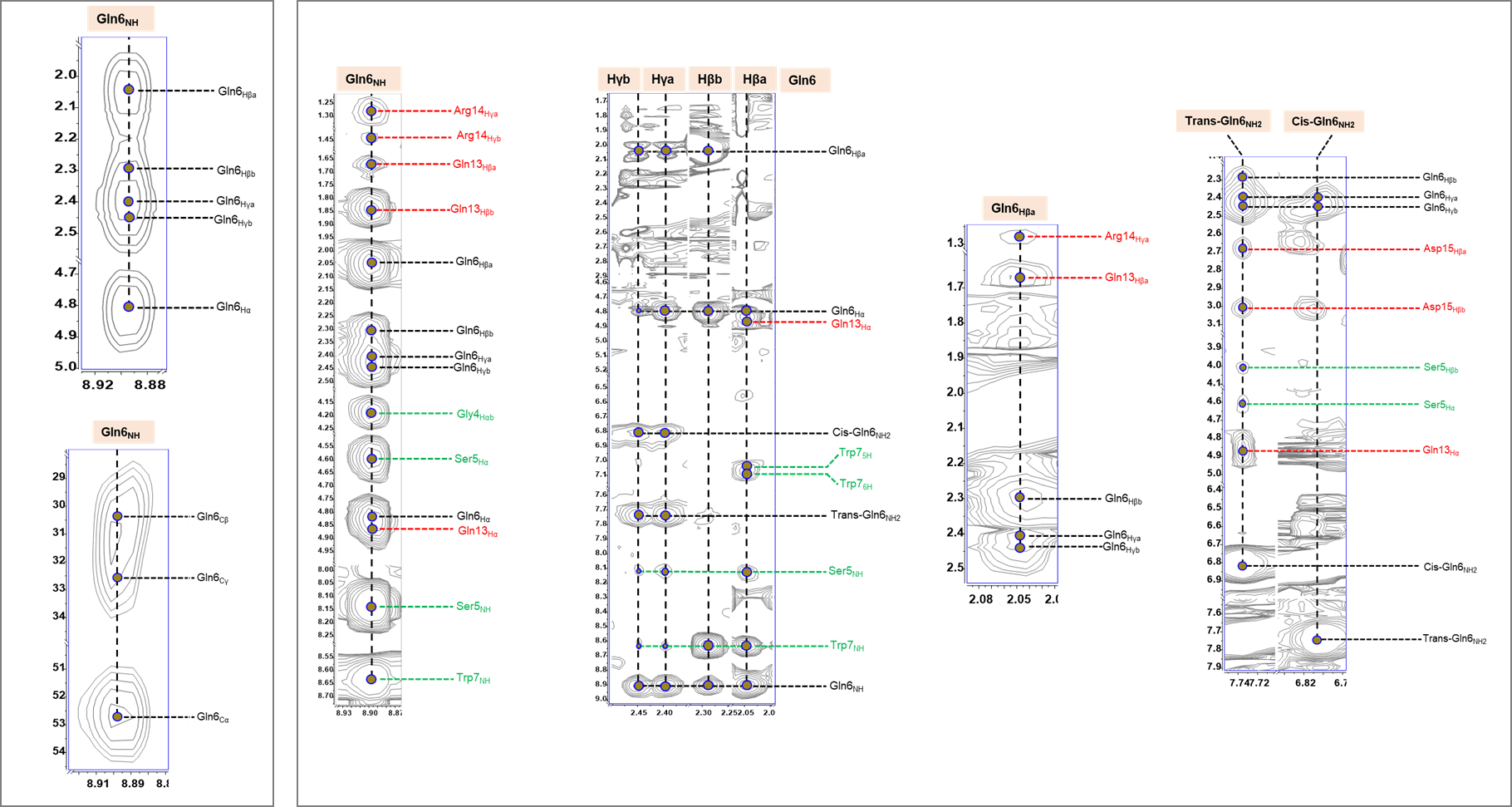

**Figure S41**. Annotated ^1^H-^1^H TOCSY, ^1^H-^13^C HSQC-TOCSY (left), and ^1^H-^1^H NOESY (right) spectra of nocapeptin A (**1**) depicting residue **Gln6**

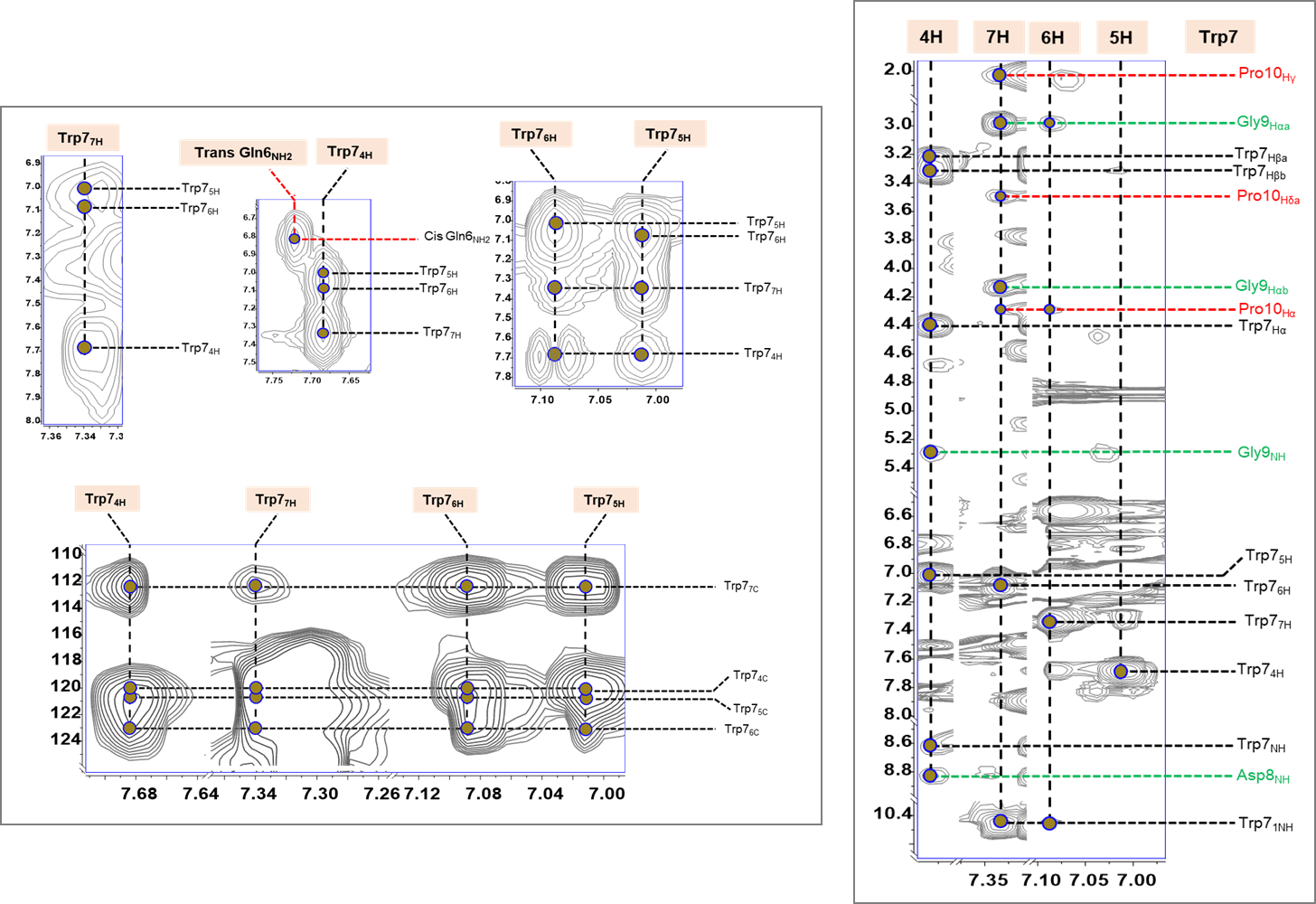

**Figure S42**. Annotated ^1^H-^1^H TOCSY, ^1^H-^13^C HSQC-TOCSY (left), and ^1^H-^1^H NOESY (right) spectra of nocapeptin A (**1**) depicting residue **Trp7**

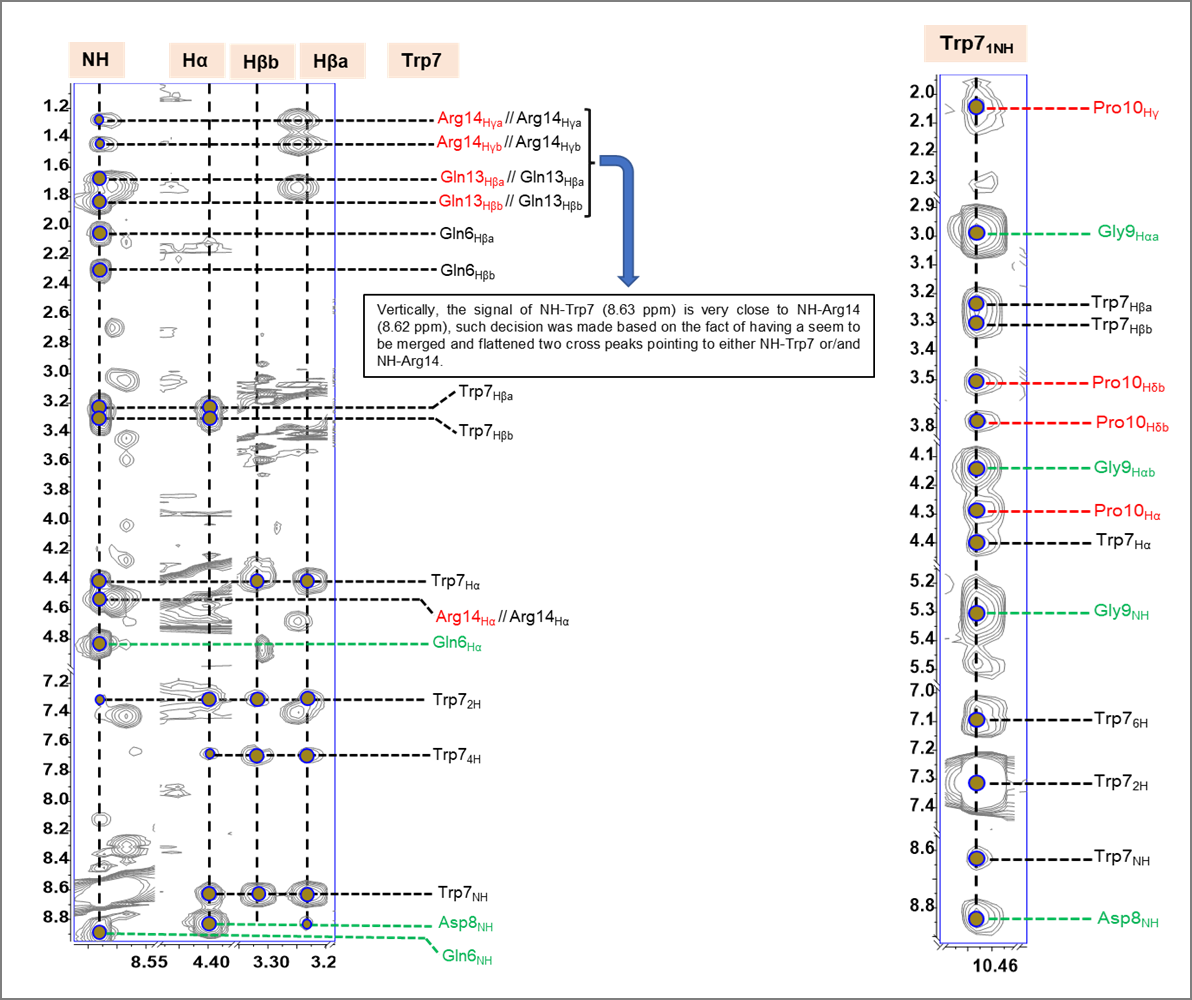

**Figure S42A**. Annotated ^1^H-^1^H NOESY spectra of nocapeptin A (**1**) depicting residue **Trp7**

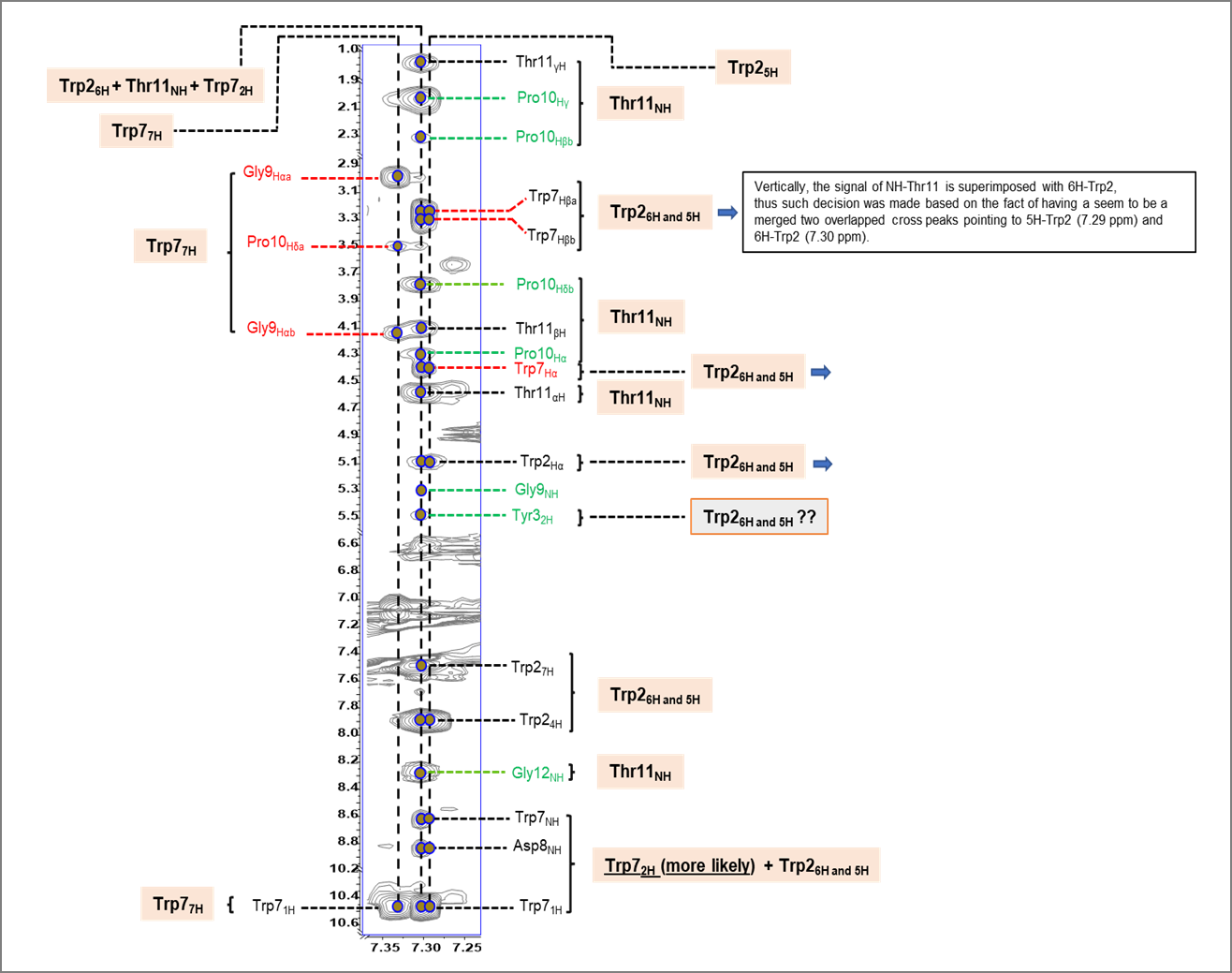

**Figure S42B**. Annotated ^1^H-^1^H NOESY spectrum of nocapeptin A (**1**) depicting residue **Trp7**

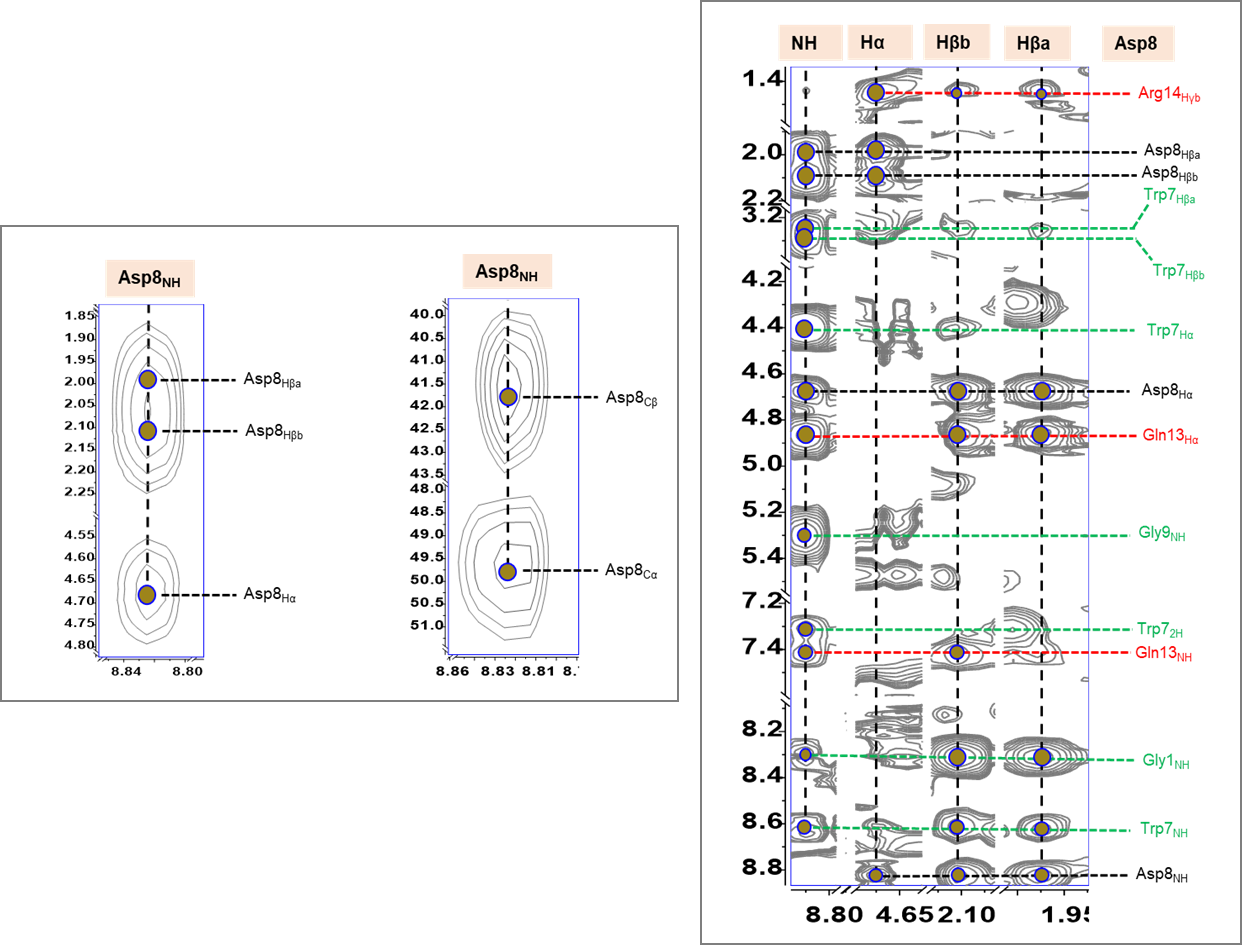

**Figure S43**. Annotated ^1^H-^1^H TOCSY, ^1^H-^13^C HSQC-TOCSY (left), and ^1^H-^1^H NOESY (right) spectra of nocapeptin A (**1**) depicting residue **Asp8**

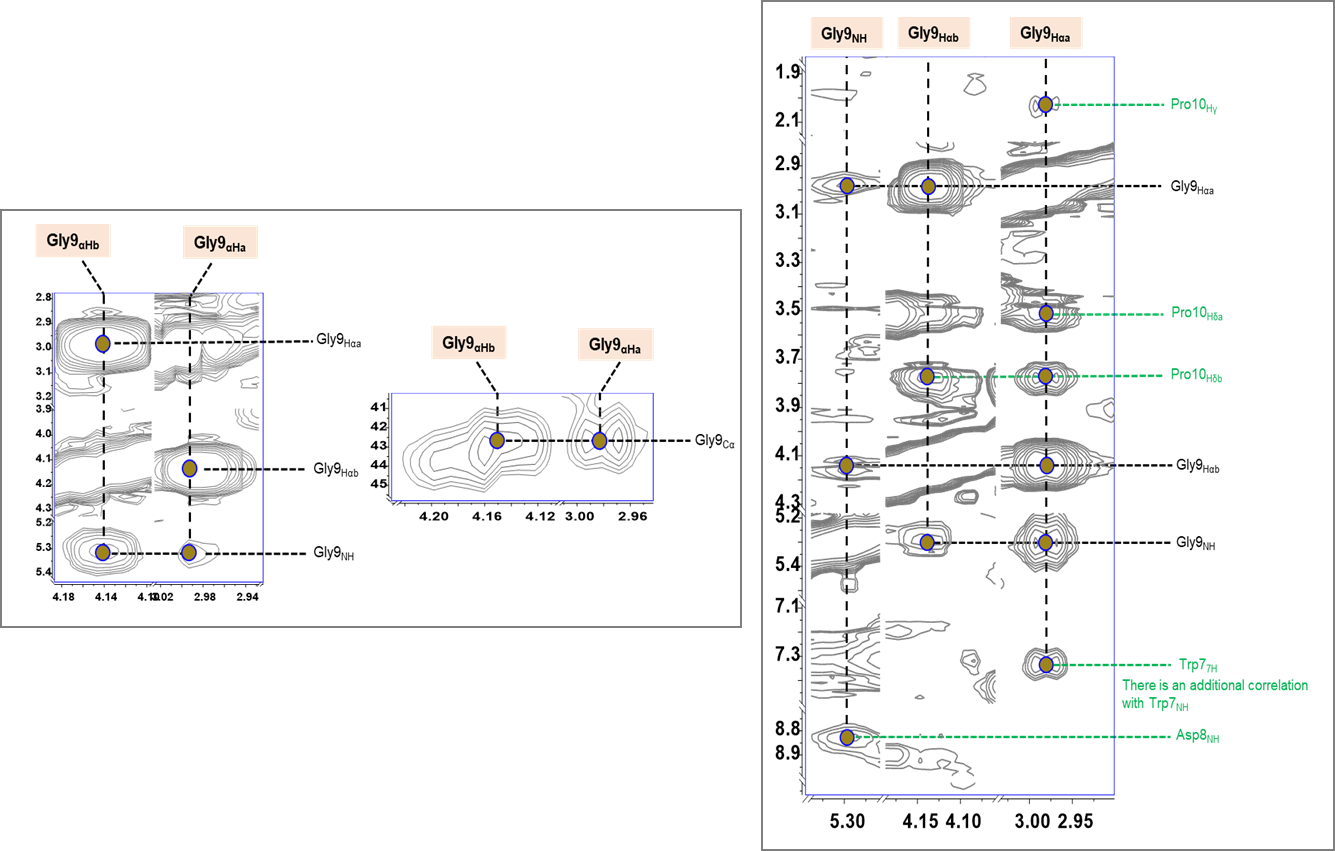

**Figure S44**. Annotated ^1^H-^1^H TOCSY, ^1^H-^13^C HSQC-TOCSY (left), and ^1^H-^1^H NOESY (right) spectra of nocapeptin A (**1**) depicting residue **Gly9**

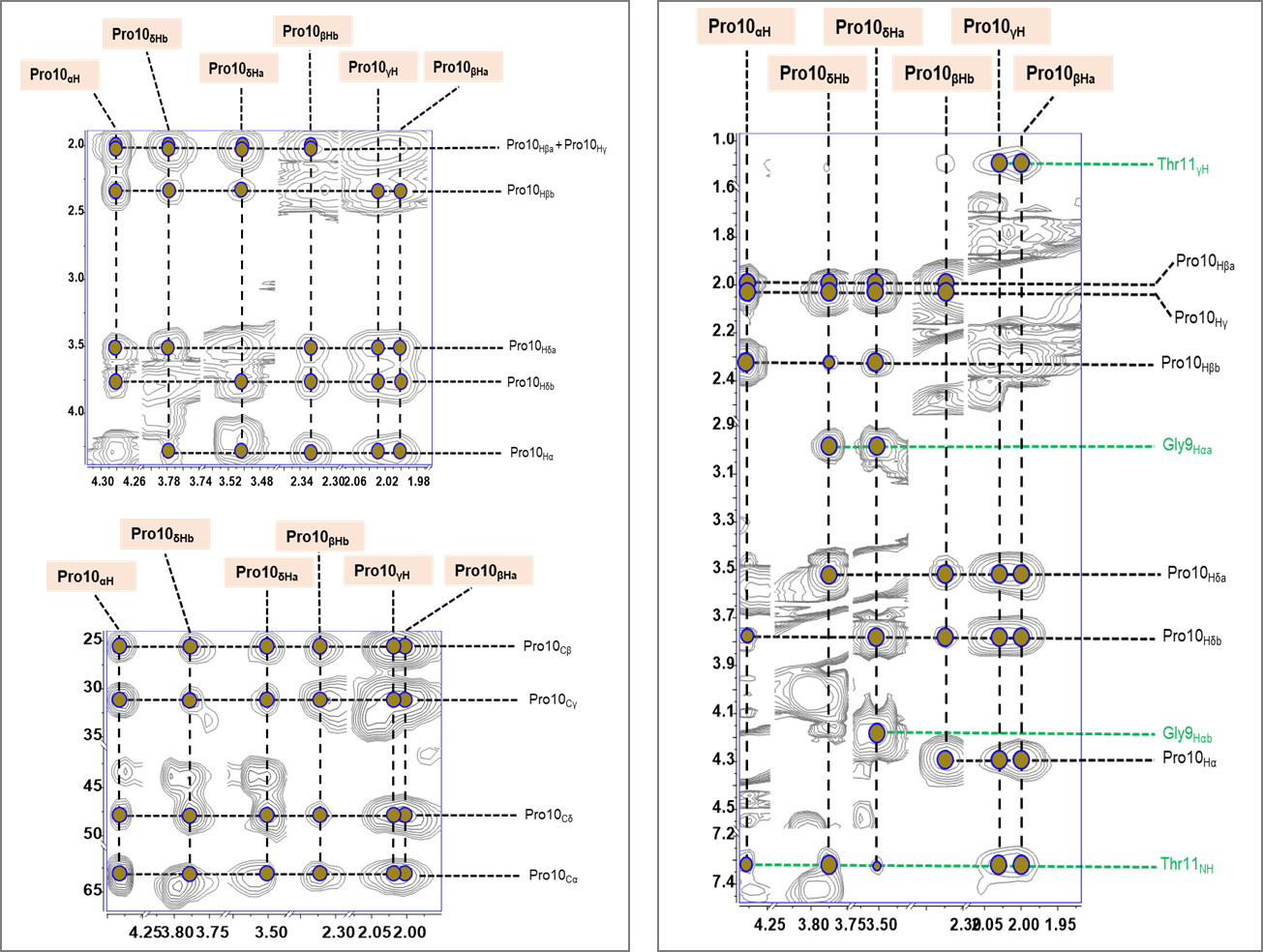

**Figure S45**. Annotated ^1^H-^1^H TOCSY, ^1^H-^13^C HSQC-TOCSY (left), and ^1^H-^1^H NOESY (right) spectra of nocapeptin A (**1**) depicting residue **Pro10**

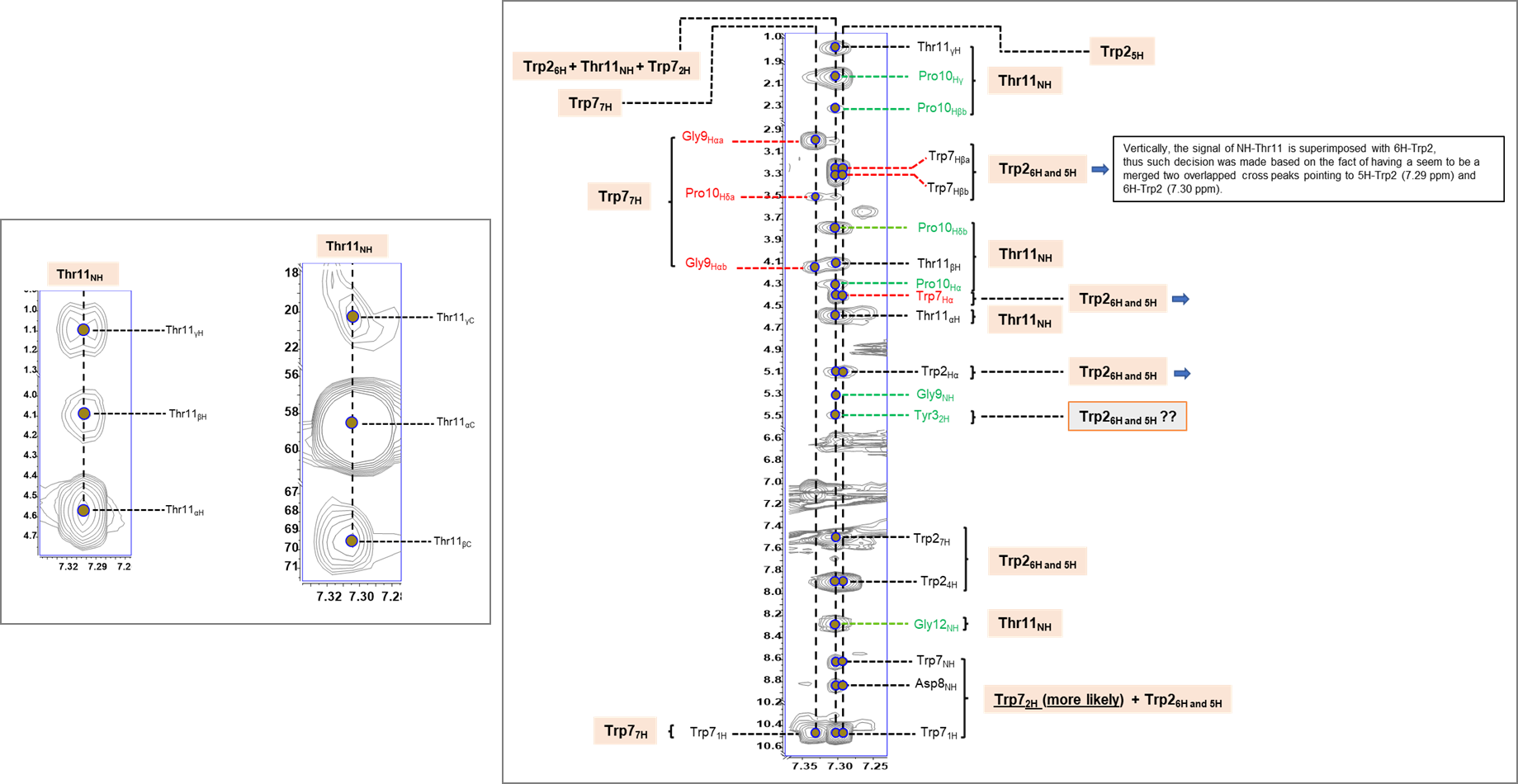

**Figure S46**. Annotated ^1^H-^1^H TOCSY, ^1^H-^13^C HSQC-TOCSY (left), and ^1^H-^1^H NOESY (right) spectra of nocapeptin A (**1**) depicting residue **Thr11**

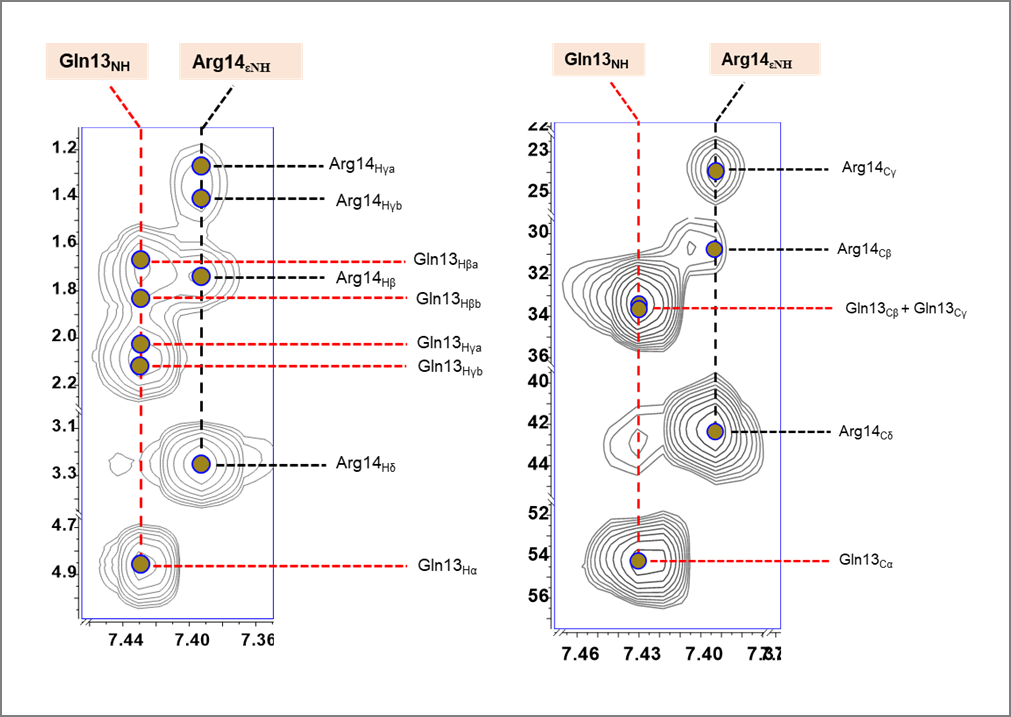

**Figure S47.** Annotated ^1^H-^1^H TOCSY (left), and ^1^H-^13^C HSQC-TOCSY (right) spectra of nocapeptin A (1) depicting residue **Gln13**

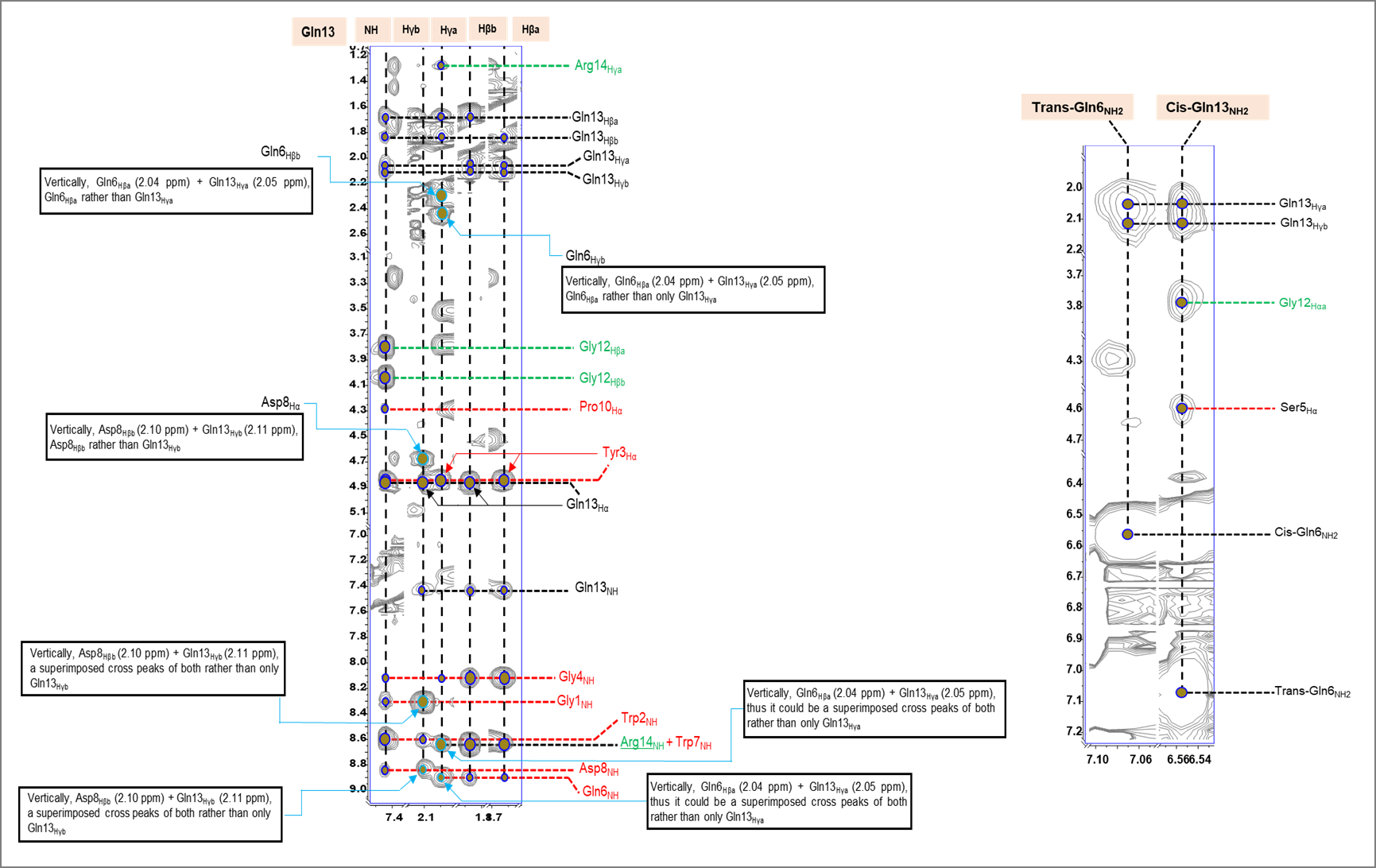

**Figure S48**. Annotated ^1^H-^1^H NOESY spectrum of nocapeptin A (**1**) depicting residue **Gln13**

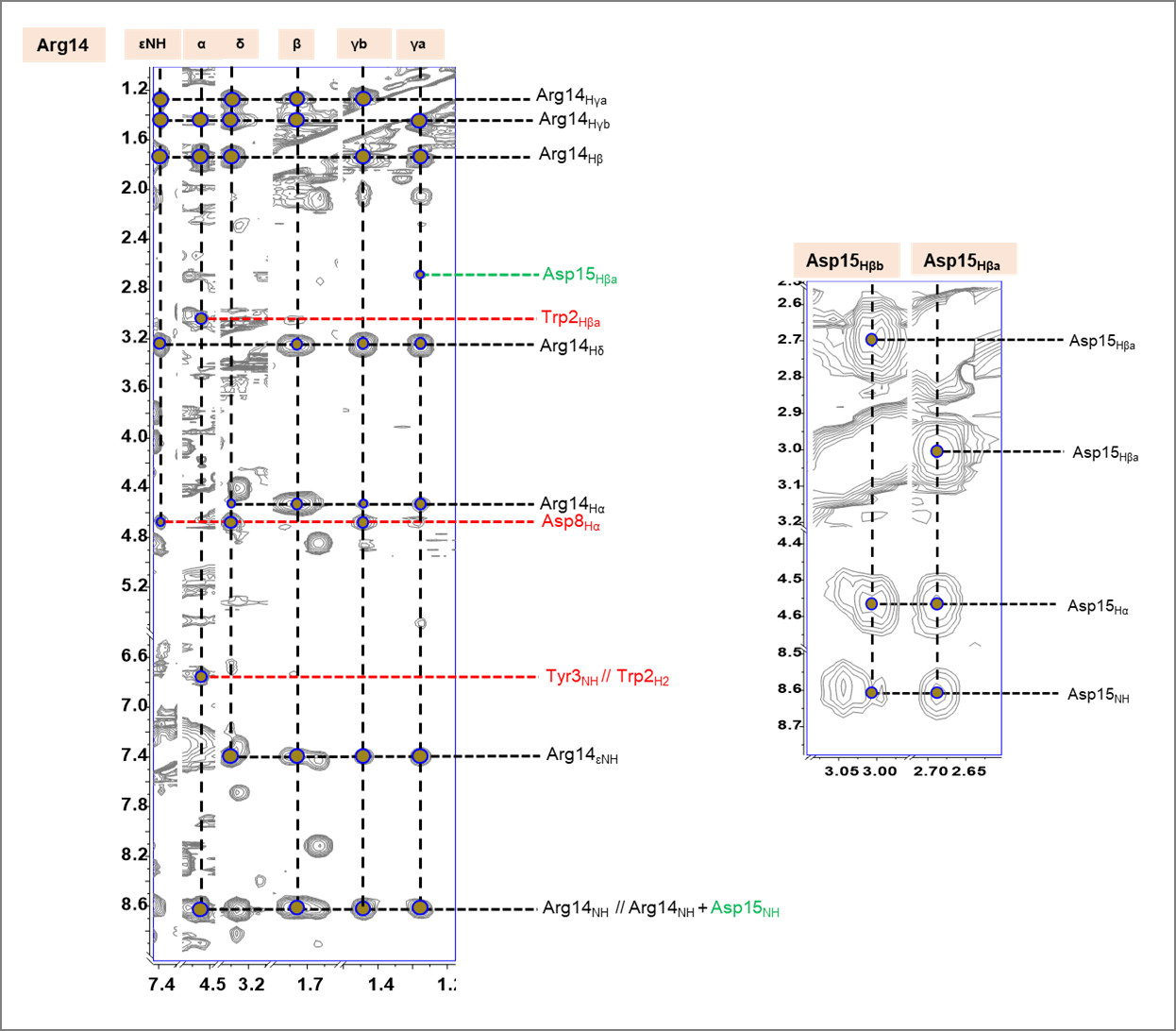

**Figure S49**. Annotated ^1^H-^1^H NOESY spectra of nocapeptin A (**1**) depicting residues **Arg14** and **Asp15**

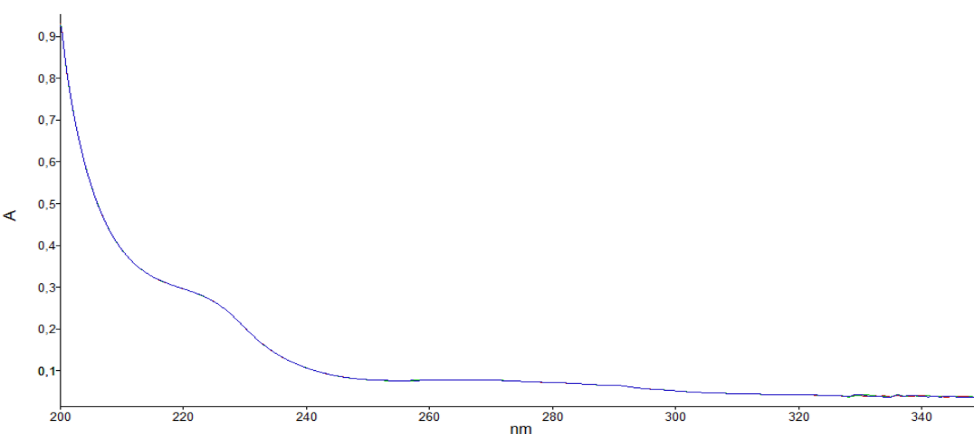

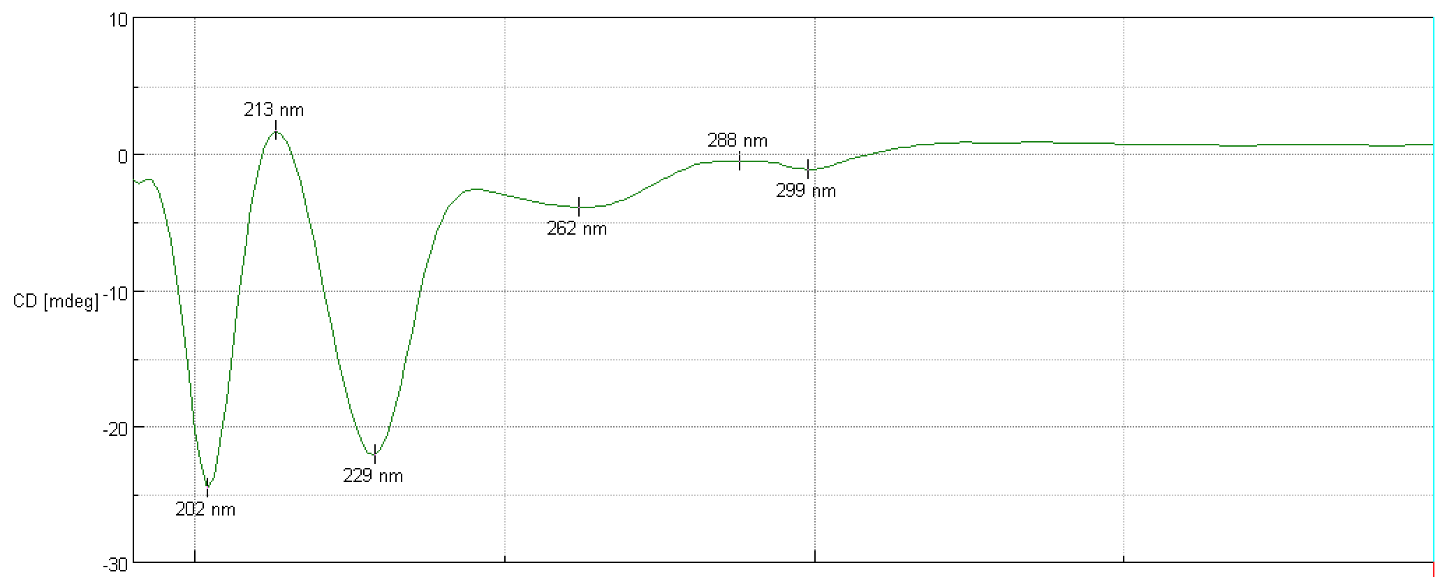

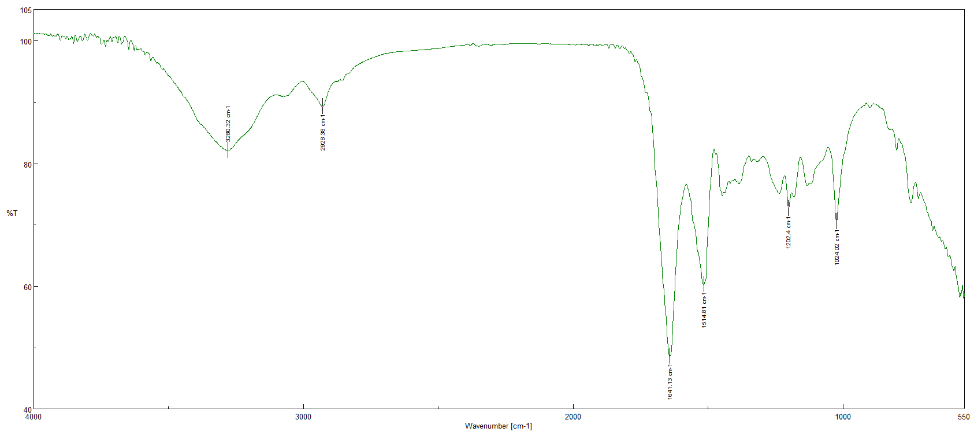

**B**

**A**

**C**)

**Figure S50.** UV (**A**), FT-IR (**B**) and CD (**C**) spectra of nocapeptin A (**1**)

Nocapeptin A (**1**): C_75_H_96_N_22_O_24_. White amorphous powder, ${[a]}_{D}^{24}$ -15.6 (*c* 0.64, MeOH) see Figure S50C; ^1^H-NMR (400 MHz in *d*_3_-CH_3_OH, and 700 MHz in 96:4 *d*_3_-CH_3_OH/H_2_O): see Tables S7 and S8, Figures S14 and S24 respectively; ^13^C-NMR (100 MHz in *d*_3_-CH_3_OH, and 176 MHz in 96:4 *d*_3_-CH_3_OH/H_2_O): see Tables S7 and S8, Figures S15 and S25 respectively; ^15^N-NMR (71 MHz, in 96:4 *d*_3_-CH_3_OH/H_2_O): see Table S8, Figure S34; FT-IR (ATR) ν_max_ (cm^-1^): 3280, 2928, 1641, 1515, 1202, 1024: see Figure S50B; UV (MeOH) λ_max_ (log ε) 220 sh (4.27), 272 (3.57) nm: see Figure S50A. HR-ESIMS *m/z* 845.3563 [M+2H]^2+^ (calcd for C_75_H_98_N_22_O_24_, 845.355667); 1689.7050 [M+H]+ (calcd for C_75_H_97_N_22_O_24_, 1689.7041). HRMSMS: see Figure S4A and Table S2

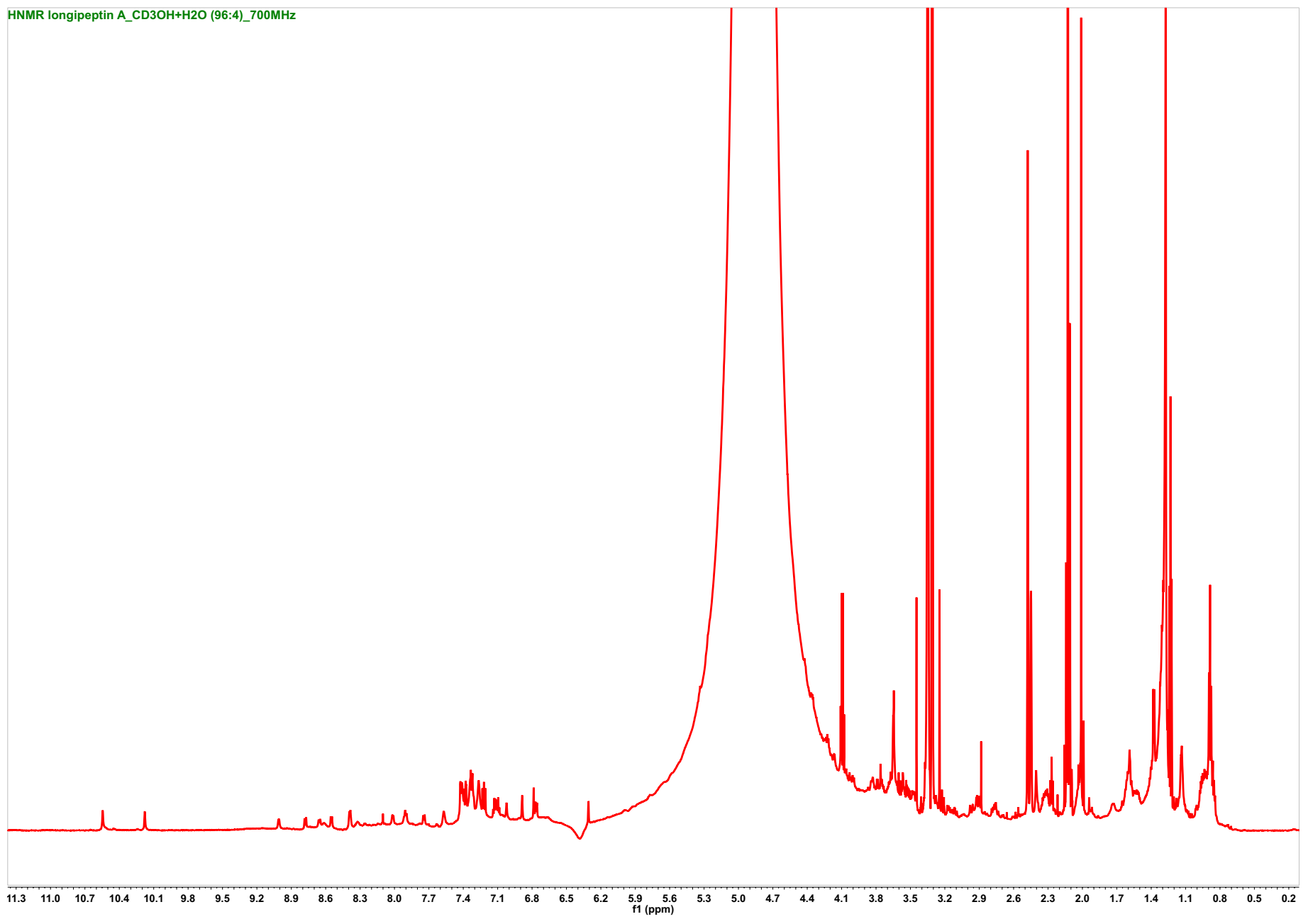

**Figure S51**. ^1^H- NMR spectrum of longipeptin A (**3**) (*d*_3_-CH_3_OH/H_2_O (96:4), 700 MHz)

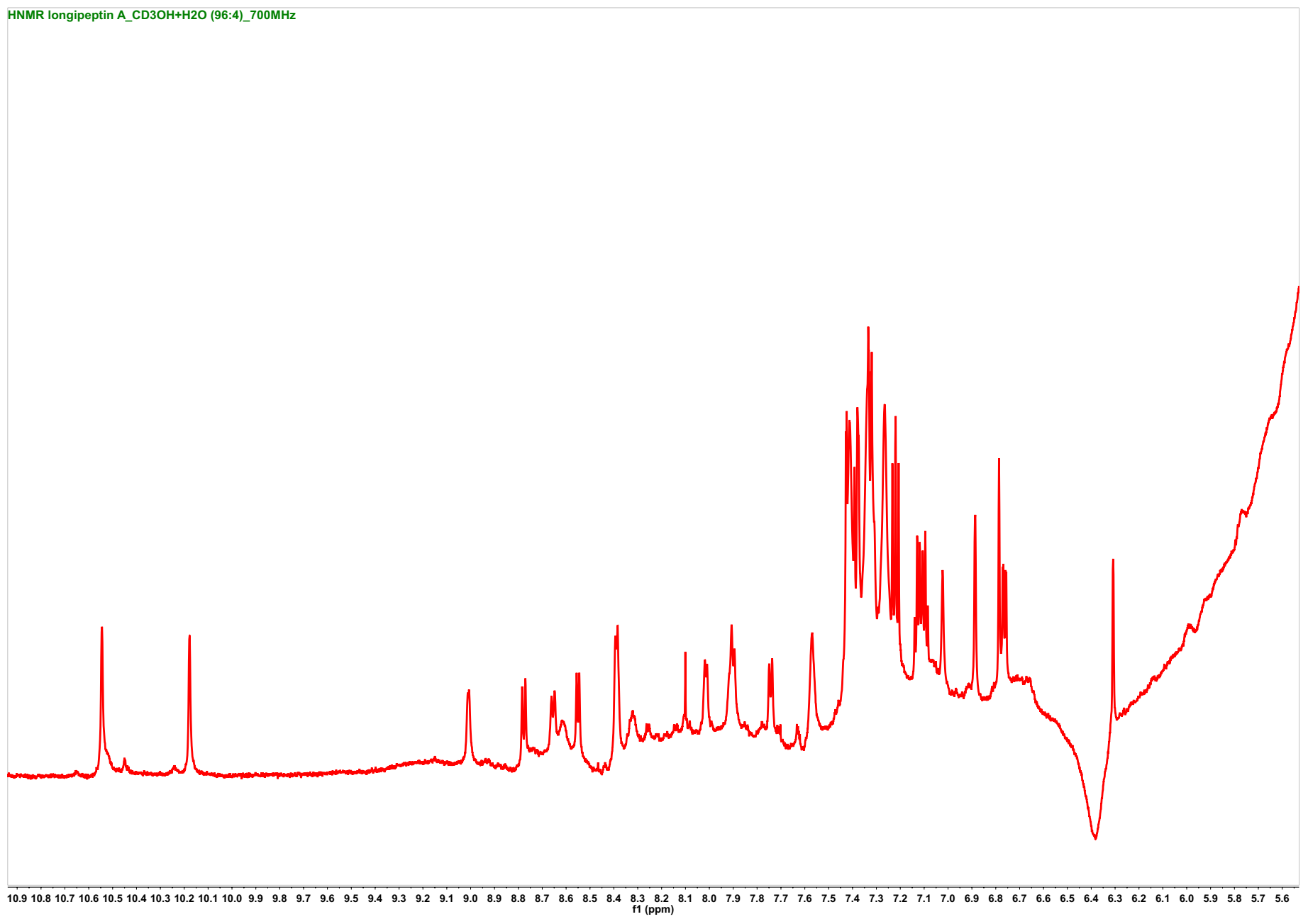

**Figure S51A**. Expanded ^1^H- NMR spectrum of longipeptin A (**3**) detailing the aromatic region (*d*_3_-CH_3_OH/H_2_O (96:4), 700 MHz)

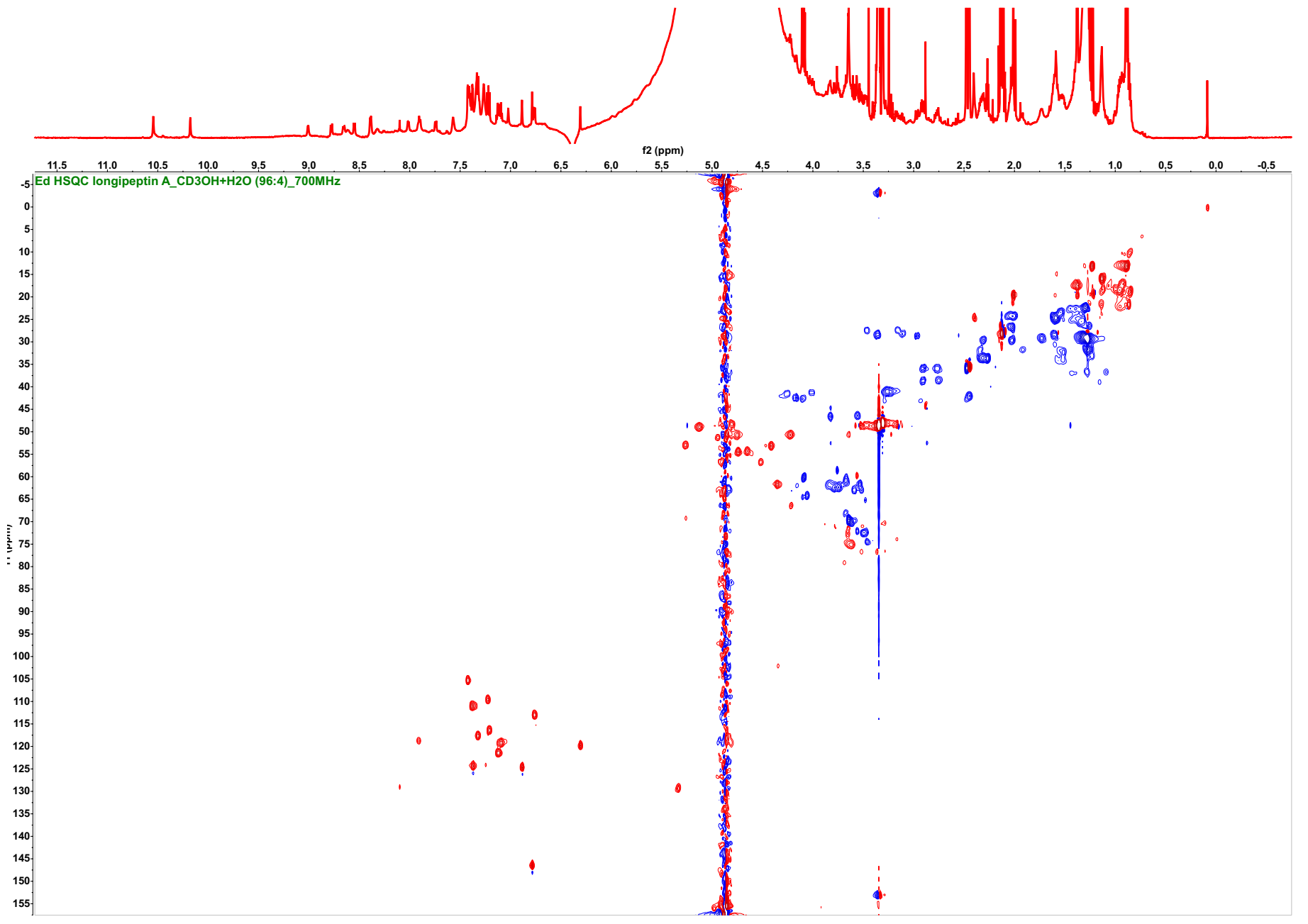

**Figure S52**. ^1^H-^13^C Edited HSQC spectrum of longipeptin A (**3**) (CH/CH_3_ are in red and CH_2_ are in blue, *d*_3_-CH_3_OH/H_2_O (96:4), 700/176 MHz)

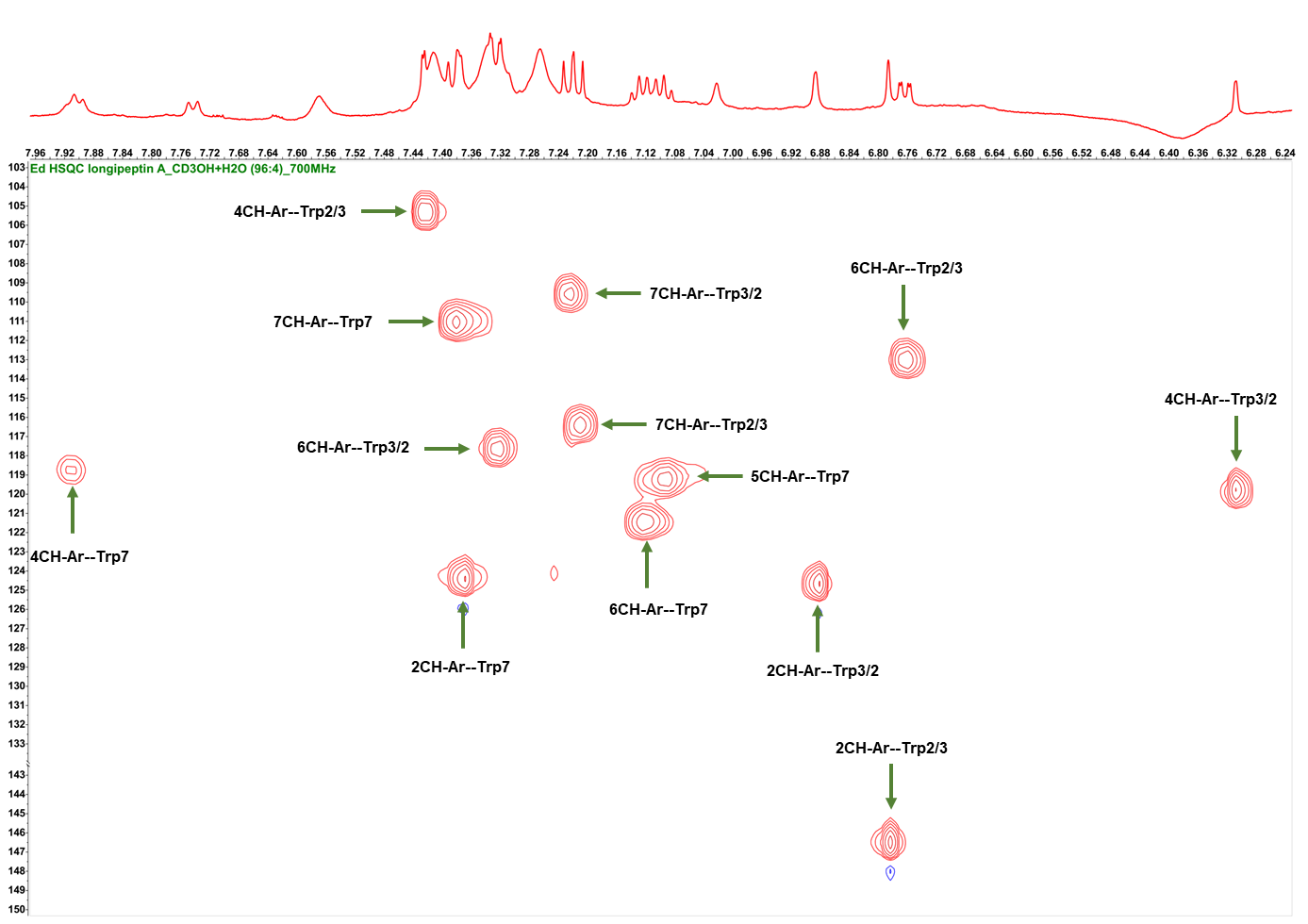

**Figure S52A**. Annotated ^1^H-^13^C edited HSQC spectrum of the assembled indolic systems (**Trp2/3**, **Trp3/2** and **Trp7**) of longipeptin A (**3**)

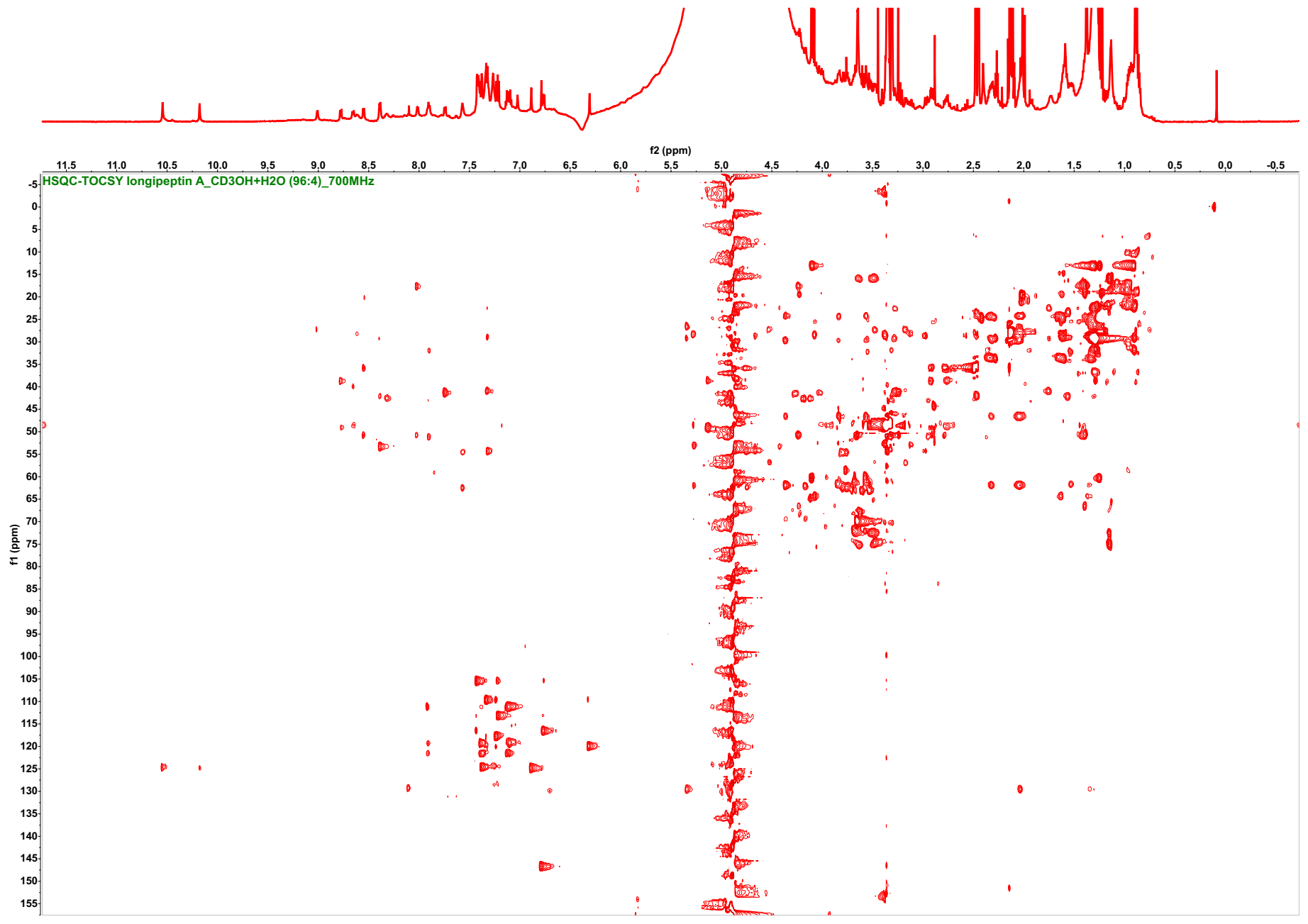
 **Figure S53**. ^1^H-^13^C HSQC-TOCSY spectrum of longipeptin A (**3**) (*d*_3_-CH_3_OH/H_2_O (96:4), 700/176 MHz)

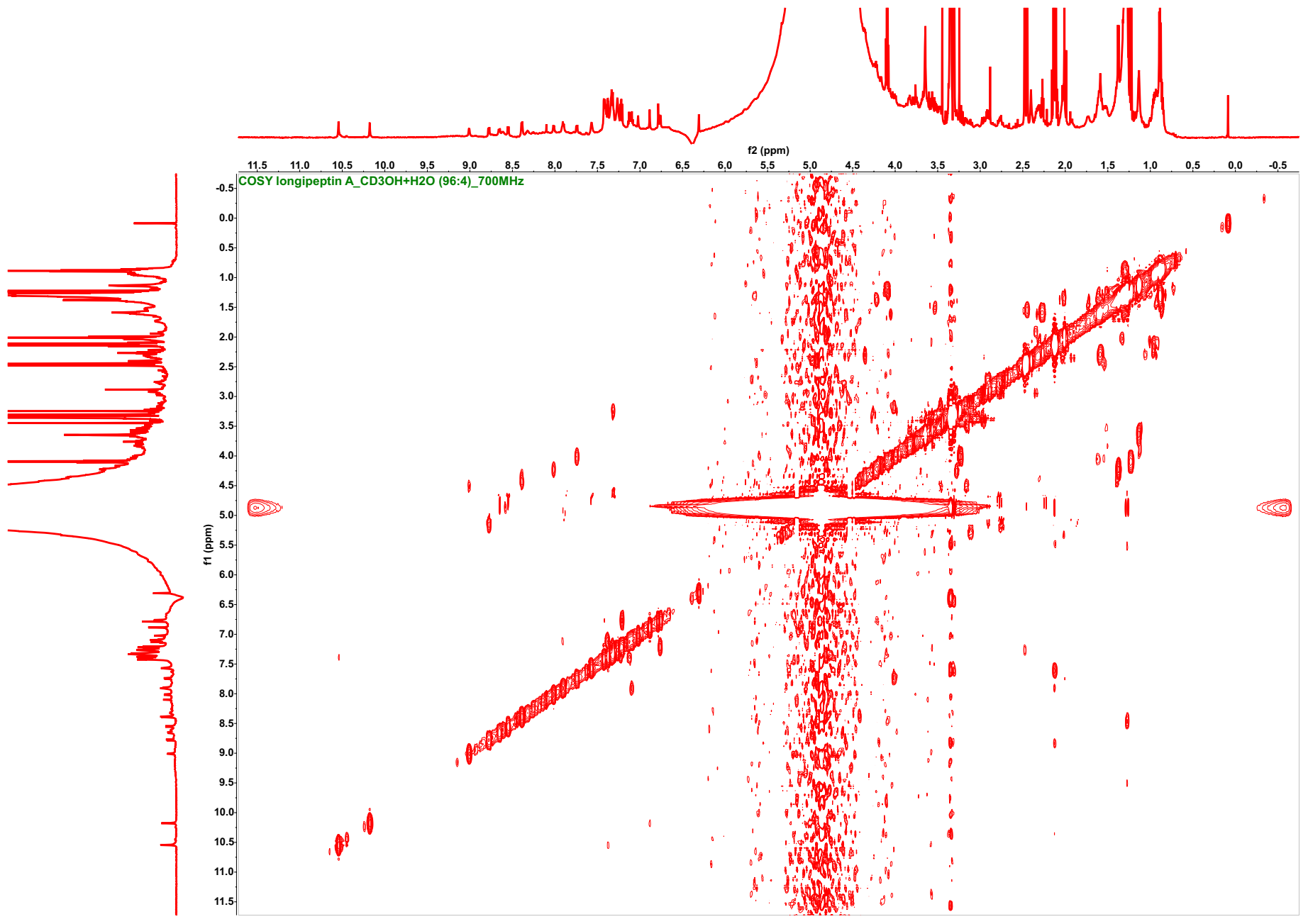

**Figure S54**. ^1^H-^1^H COSY spectrum of longipeptin A (**3**) (*d*_3_-CH_3_OH/H_2_O (96:4), 700 MHz)

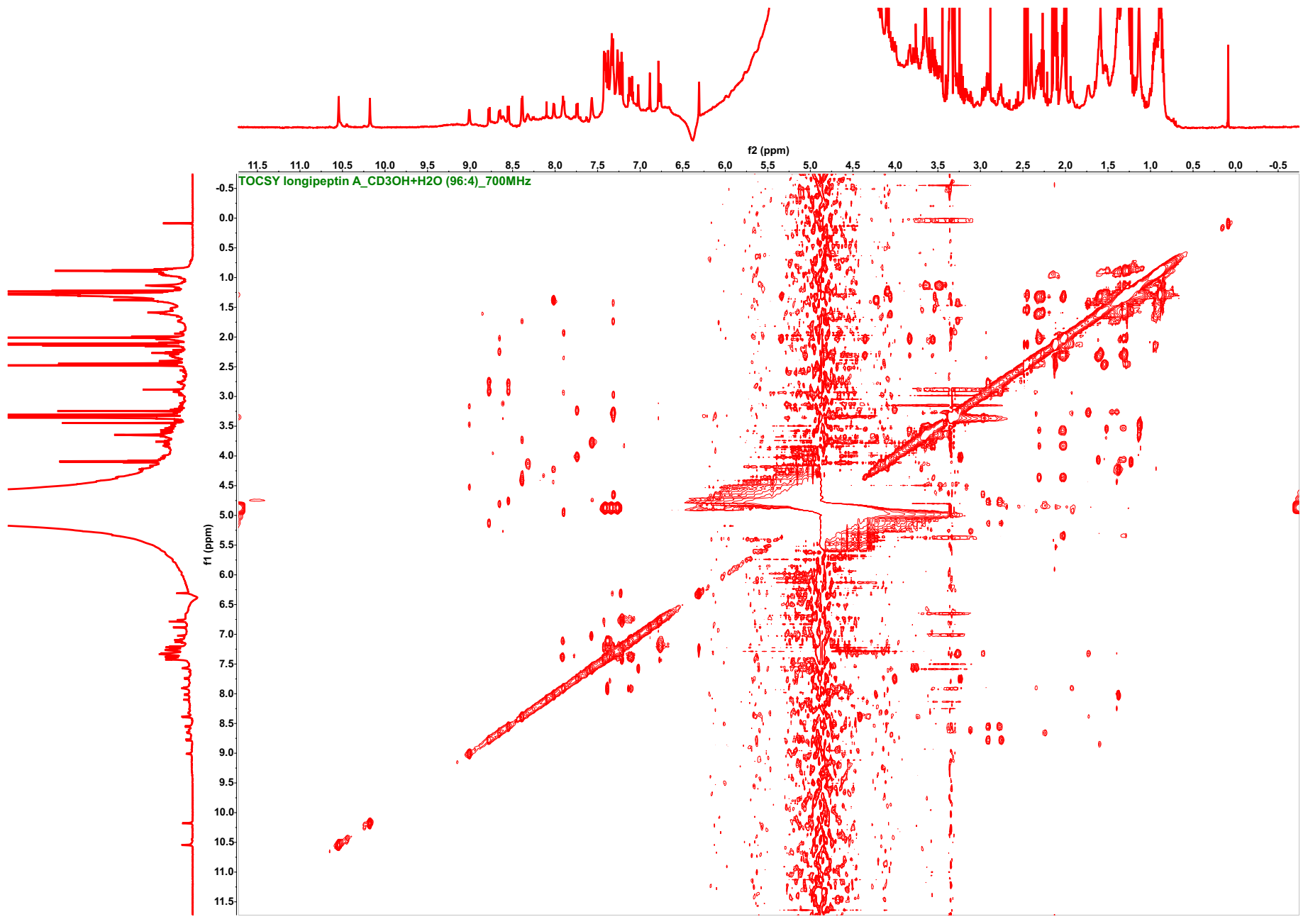
 **Figure S55**. ^1^H-^1^H TOCSY spectrum of longipeptin A (**3**) (*d*_3_-CH_3_OH/H_2_O (96:4), 700 MHz)

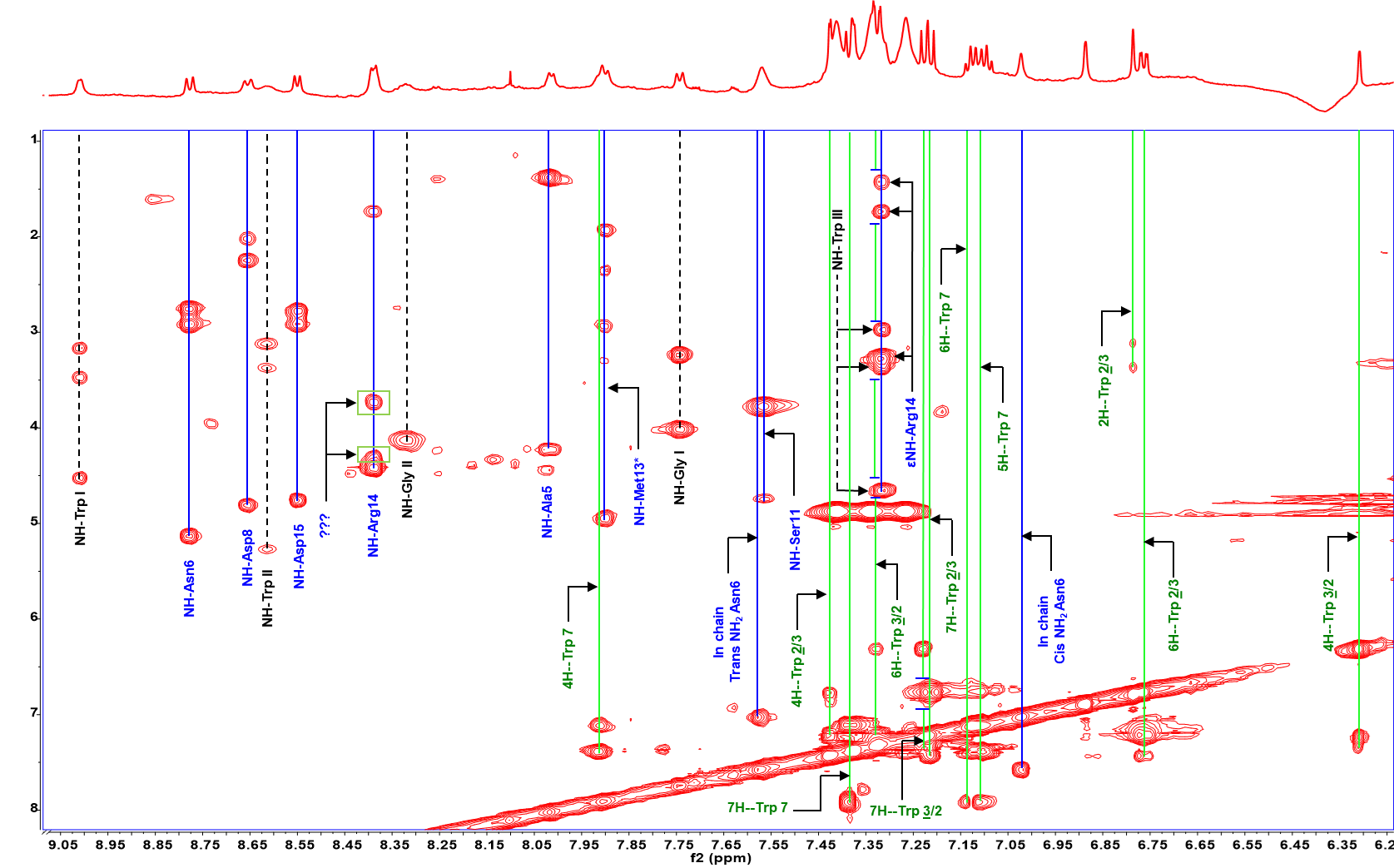

**Figure S55A**. Annotated ^1^H-^1^H TOCSY spectrum, highlighting the assembled spin systems of longipeptin A (**3**)

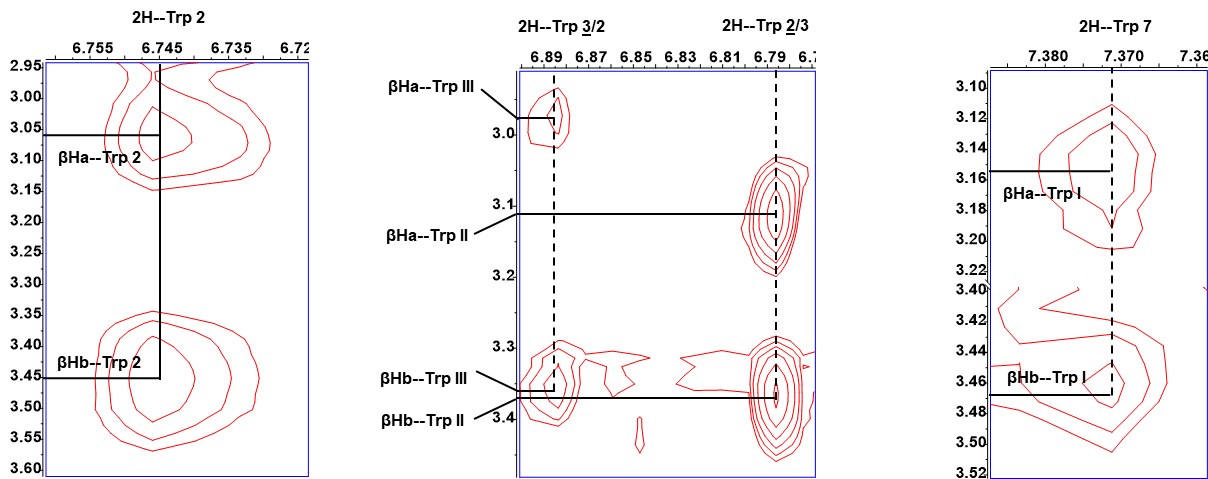

**Figure S55B**. Annotated ^1^H-^1^H TOCSY spectra, highlighting the observation of a W coupling between the 2H and βHa+b spin systems of the Trp residues

in nocapeptin A (**1**) (left) vs longipeptin A (**3**) (middle and right)

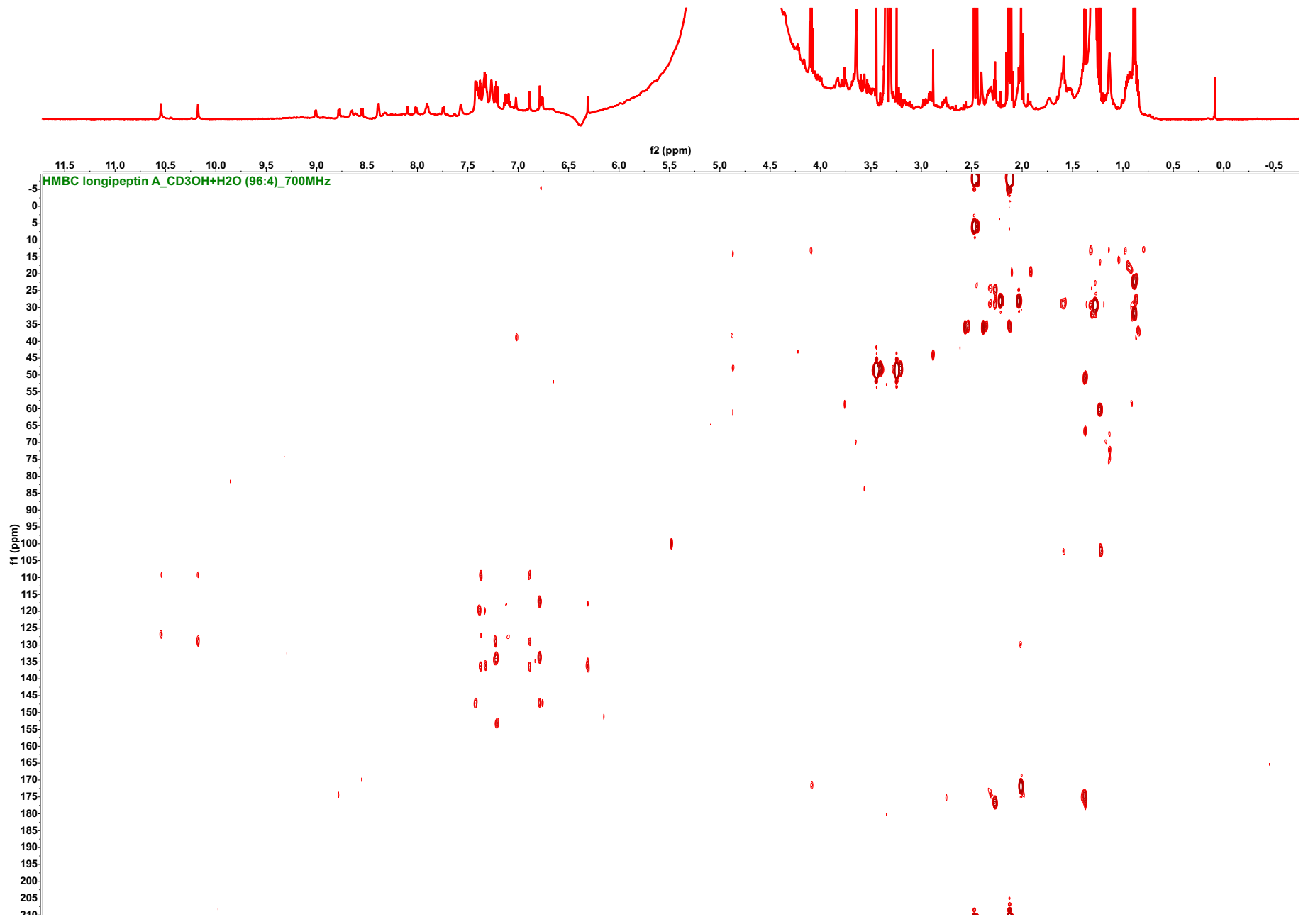

**Figure S56**. ^1^H-^13^C HMBC spectrum of longipeptin A (**3**) (d_3_-CH_3_OH/H_2_O (96:4), 700/176 MHz)

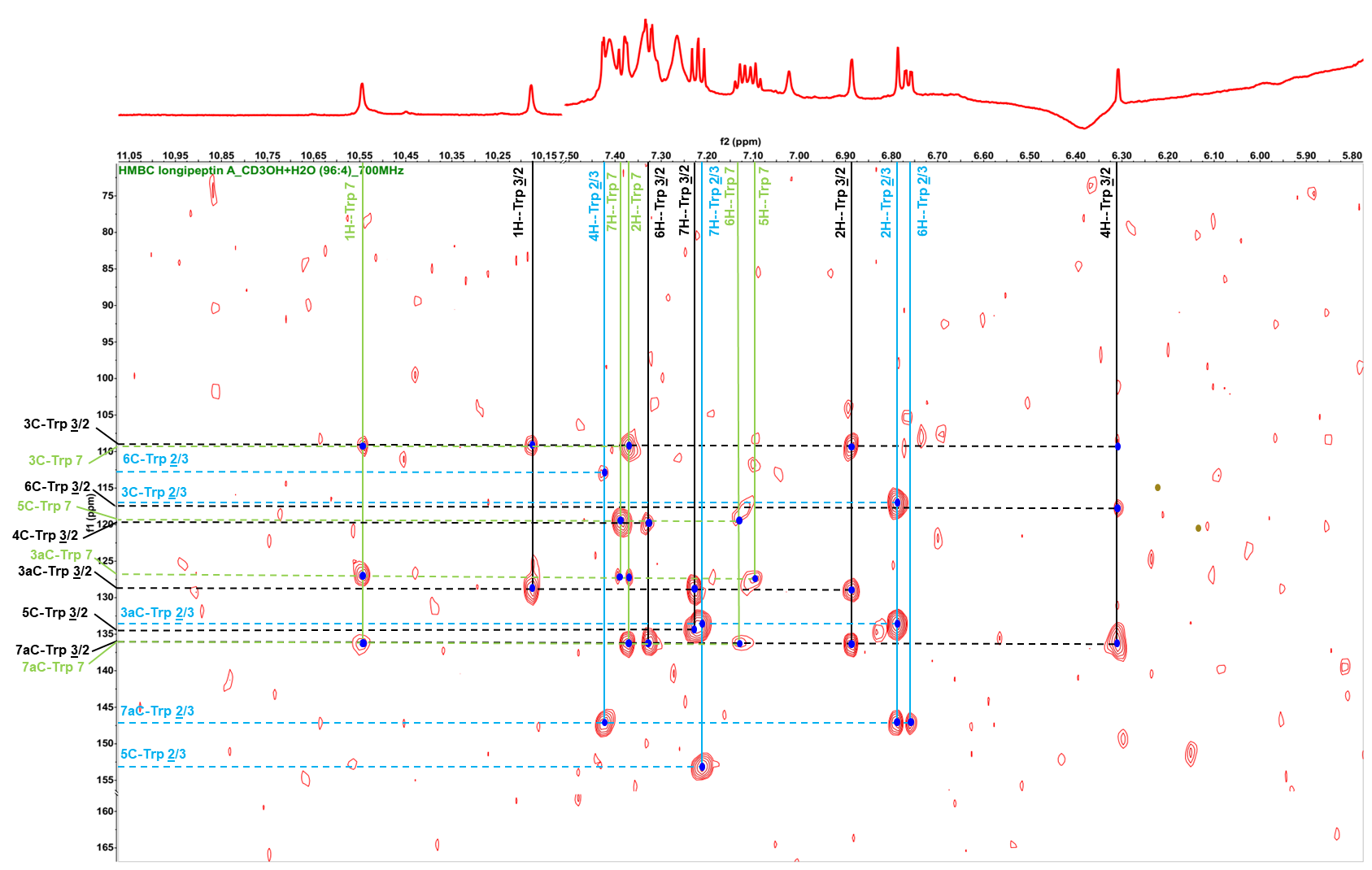

**Figure S56A**. Annotated ^1^H-^13^C HMBC spectrum highlighting the assembled indolic systems (**Trp2/3**, **Trp3/2** and **Trp7)** of longipeptin A (**3**)

**Table S9**. ^1^H, and ^13^C-NMR data of longipeptin A (**3**) [*d*_3_-CH_3_OH/H_2_O (96:4); 700/176 MHz]

| **Residue** | **Position** | **δ_C_** | **δ_H_, mult (*J* in Hz)** | **Residue** | **Position** | **δ_C_** | **δ_H_, mult (*J* in Hz)** |
| --- | --- | --- | --- | --- | --- | --- | --- |
| **Gly1** | CO | n.d. | -------- | **Trp7** | CO | n.d. | -------- |
|  | α | *See Figure S57* | *See Figure S57* |  | α | 56.78, CH | 4.52 |
|  | -NH- | -------- | *See Figure S57* |  | β | 27.44, CH_2_ | 3.16 + 3.47 |
| **Trp2/3*** | CO | n.d. | -------- |  | 1 (NH) | -------- | 10.55 s |
|  | α | 52.97, CH | 5.27 |  | 2 | 124.40, CH | 7.37 |
|  | β | 28.41, CH_2_ | 3.12 + 3.37 |  | 3 | 109.44, C | -------- |
|  | 1(N) | -------- | -------- |  | 3a | 127.26, C | -------- |
|  | 2 | 146.46, CH | 6.79 br s |  | 4 | 118.74, CH | 7.91 |
|  | 3 | 117.16, C | -------- |  | 5 | 119.22, CH | 7.09 dd (7.6, 7.2) |
|  | 3a | 133.60, C | -------- |  | 6 | 121.46, CH | 7.12 dd (7.6, 7.2) |
|  | 4 | 105.33, CH | 7.42 d (2.2) |  | 7 | 111.07, CH | 7.38 |
|  | 5 | 153.20, C | -------- |  | 7a | 136.25/136.34, C | -------- |
|  | OH | -------- | n.d. |  | -NH- | -------- | 9.01 |
|  | 6 | 113.03, CH | 6.76 dd (8.7, 2.2) | **Asp8** | CO | n.d. | -------- |
|  | 7 | 116.41, CH | 7.21 d (8.7) |  | α | 48.29, CH | 4.81 |
|  | 7a | 147.31, C | -------- |  | β | 39.91, CH_2_ | 2.02 + 2.25 |
|  | -NH- | -------- | 8.61 |  | γ | n.d. | -------- |
| **Trp3/2*** | CO | n.d. | -------- |  | -NH- | -------- | 8.65 d (8.6) |
|  | α | 54.33, CH | 4.65 | **Gly9** | CO | n.d. | -------- |
|  | β | 28.69, CH_2_ | 2.97 + 3.36 |  | α | *See Figure S57* | *See Figure S57* |
|  | 1 (NH) | -------- | 10.18 s |  | -NH- | -------- | *See Figure S57* |
|  | 2 | 124.70, CH | 6.89 br s | **Pro10** | CO | n.d. | -------- |
|  | 3 | 109.25, C | -------- |  | α | 61.76, CH | 4.36 |
|  | 3a | 128.95, C | -------- |  | β | 29.63, CH_2_ | 2.01 + 2.31 |
|  | 4 | 119.80, CH | 6.31 br s |  | γ | 24.24, CH_2_ | 2.02 |
|  | 5 | 134.45, C | -------- |  | δ | 46.44, CH_2_ | 3.56 + 3.83 |
|  | 6 | 117.66, CH | 7.32 dd (8.4, 1.8) | **Ser11** | CO | n.d. | -------- |
|  | 7 | 109.59, CH | 7.23 dd (8.4) |  | α | 54.41, CH | 4.74 |
|  | 7a | 136.25/136.34, C | -------- |  | β | 62.38, CH_2_ | 3.75 + 3.79 |
|  | -NH- | -------- | 7.31 |  | OH | -------- | n.d. |
| **Gly4** | CO | n.d. | -------- |  | -NH- | -------- | 7.57 |
|  | α | *See Figure S57* | *See Figure S57* | **Gly12** | CO | n.d. | -------- |
|  | -NH- | -------- | *See Figure S57* |  | α | *See Figure S57* | *See Figure S57* |
| **Ala5** | CO | n.d. | -------- |  | -NH- | -------- | *See Figure S57* |
|  | α | 50.70, CH | 4.22 | **Met13**** | CO | n.d. | -------- |
|  | β | 17.57, CH_2_ | 1.38 d (7.2) |  | α | 51.31, CH | 4.95 |
|  | -NH- | -------- | 8.01 d (7.0) |  | β | 31.78, CH_2_ | 1.93 + 2.35 |
| **Asn6** | CO | n.d. | -------- |  | γ | 43.13, CH_2_ | 2.93 + 3.29 |
|  | α | 48.96, CH | 5.13 |  | S-Me_1_+ Me_2_ | n.d. | n.d. |
|  | β | 38.70, CH_2_ | 2.75 + 2.91 |  | -NH- | -------- | 7.90 d (8.3) |
|  | γ | n.d. | -------- |  |  |  |  |
|  | -NH_2_ | -------- | 7.02 br s + 7.57 br s |  |  |  |  |
|  | -NH- | -------- | 8.78 d (9.4) |  |  |  |  |
| **Arg14** | CO | n.d. | -------- |  |  |  |  |
|  | α | 53.17, CH | 4.41 |  | | | |
|  | β | 29.25, CH_2_ | 1.73 |  |  |  |  |
|  | γ | 22.70, CH_2_ | 1.43 |  |  |  |  |
|  | δ | 41.09, CH_2_ | 3.27 |  |  |  |  |
|  | εNH | -------- | 7.32 |  |  |  |  |
|  | ζ | n.d. | -------- |  |  |  |  |
|  | NHη1 | -------- | n.d. |  |  |  |  |
|  | NH_2_η1 | -------- | n.d. |  |  |  |  |
|  | -NH- | -------- | 8.39 d (6.7) |  |  |  |  |
| **Asp15** | CO_2_H | n.d. | -------- |  |  |  |  |
|  | α | 50.79, CH | 4.76 |  |  |  |  |
|  | β | 35.97, CH_2_ | 2.77 + 2.91 |  |  |  |  |
|  | γ | n.d. | -------- |  |  |  |  |
|  | -NH- | -------- | 8.55 d (7.7) |  |  |  |  |

* Residues involved in the C-N biaryl crosslink in an alternative manner.

** Residues involved in the S-methylation.

- CH_3_, CH_2_, and CH chemical shift values were extracted indirectly from ^1^H-^13^C HSQC.
- C chemical shift values were extracted indirectly from ^1^H-^13^C HMBC.

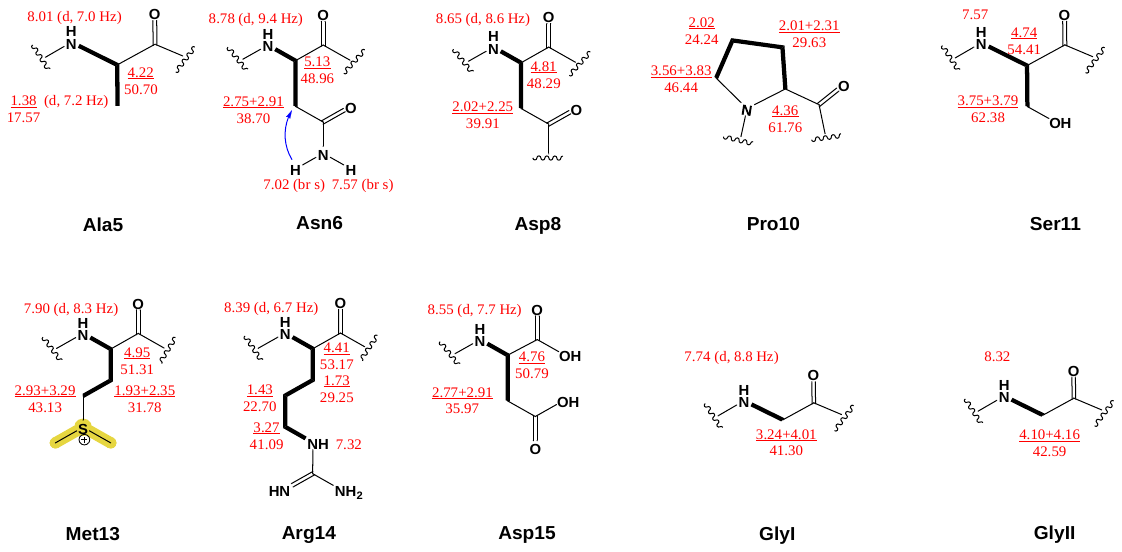

**Figure S57**. Schematic representation of the assembled spin systems of longipeptin A (**3**) using ^1^H-^1^H COSY, ^1^H-^1^H TOCSY (bold lines) and ^1^H-^13^C HMBC (blue arrows) correlations

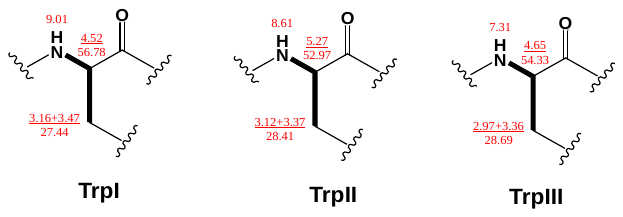

**Figure S57A**. Schematic representation of the assembled spin systems constituting the Trp residues of longipeptin A (**3**) using ^1^H-^1^H COSY and ^1^H-^1^H TOCSY correlations (bold lines)

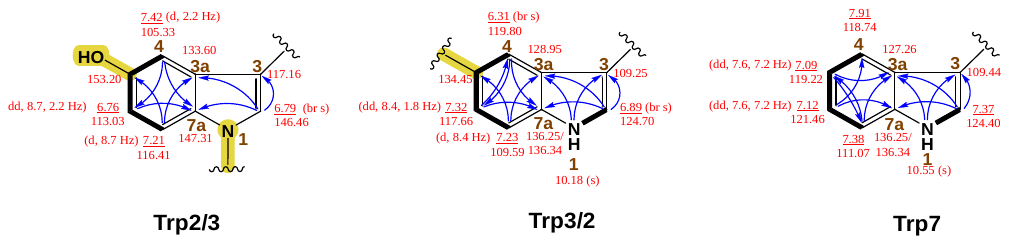

**Figure S57B**. Schematic representation of the assembled spin systems constituting the Trp residues of longipeptin A (**3**) using ^1^H-^1^H COSY, ^1^H-^1^H TOCSY, ^1^H-^13^C HSQC-TOCSY (bold lines) and ^1^H-^13^C HMBC (blue arrows) correlations

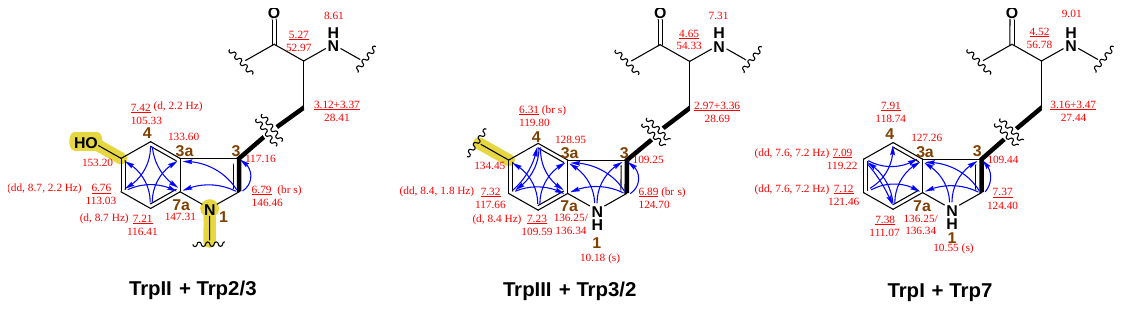

**Figure S57C**. *Schematic representation of the connected structural units,* **Trp I--Trp7**, **Trp II--Trp2/3** and **Trp III--Trp3/2** *constituting the W residues of longipeptin A (****3****) based on the ^1^H-^1^H TOCSY W-coupling (bold lines) and ^1^H-^13^C HMBC (blue arrows) correlations,*

**Partial Structure Elucidation of longipeptin A (3)**

Considering the exhaustive analysis of the MS^1^ and MS^2^ longipeptin series (A-C, **3**-**5**), a reasonable assumption was formulated in terms of their morphed skeletons with the differently appended PTMs. The highly modified major variant, longipeptin A (**3**) was hypothesized to be the final product of the *lop* BGC loaded with three tailoring events (oxidation ‘+O’, methylation ‘+CH_2_’ and crosslink ‘-2H’) aligning with the unusual genetic elements featured in the BGC that consists of three ancillary processing enzymes (Figure S1).

The usage of tandem MS enabled not only sequencing **3** but also defining the nature and the positions of the multiple structural modifications. The first tailoring, the methylation event, was initially envisioned to occur at the C-terminus residue, D15 as previously reported^[22]^ however, such an assumption was readily ruled out after studying the MS^2^ fragments. The observation of the B ion series (**b_3_-b_12_**) and the surprising inability to spot the typical intense Y fragments, normally arising from the tail, indicated an unusual fragmentation behavior upon the methylation reaction. Such an observation was further supported by the spectral similarity with variant B (**4**) which shares the same PTM of interest (methylation) (Figure S8). An orthogonal piece of evidence was also extracted from the extensive annotation of the highly informative MS^2^ spectrum of the biosynthetic intermediate longipeptin C (**5**) that lacked such a structural modification (Figure S11).

As a result, and having in hand a dereplicated set of B ions from the MS^2^ spectrum of **3**, several possible methylation centers were postulated, principally at the side chain heteroatoms of the tail (O => **Ser11**, N => **Arg14**, S => **Met13** or N amidic backbone). Interestingly, the S-methylation at the **Met13** residue was found to be the most convincing hypothesis which was validated by the perfect alignment with the newly observed series of Y ions, arising upon the neutral loss of a characteristic 62 Da in the form of dimethylsulfide (Me_2_S) (Figures S9-9A). Similarly, the MS^2^ fragments of **4**, a methylated shunt product, were also found to fit with such a structural suggestion confirming this rare C-S bond formation (Figures S10-10A).

The remaining PTMs of **3**, oxidation and crosslink, were also found to be introduced at the **G1-W2-W3** sequence, based on the MS/MS annotation in a similar pattern to nocapeptins (Figures S4A-4C). To corroborate the MS-deduced structural findings of **3**, ^1^H-^1^H COSY, ^1^H-^1^H TOCSY and ^1^H-^13^C HSQC spectra were utilized to construct the majority of the constituting residues of longipeptin A (**3**) scaffold (Figures S57-57B). Although the predicted CP and MS analysis highlighted the presence of 4x Gly residues, we were only limited to assembling a pair of them from the homonuclear 2D-NMR (Figure S55A). Interestingly, the tandem usage of ^1^H-^1^H TOCSY, ^1^H-^13^C HSQC-TOCSY and ^1^H-^13^C HSQC supported indirectly the S-catalyzed methylation event of M13 residue via the downfield chemical shift of its γCH_2_ to be resonating at around 43.20 ppm in contrast to the typical values, 29-31 ppm (Figures S52 and S57).

In addition, three candidate spin systems, **TrpI**, **TrpII** and **TrpIII** were elucidated in concert with three indolic substructures (**Trp2/3**, **Trp3/2** and **Trp7**), mainly with the aid of ^1^H-^1^H TOCSY, ^1^H-^13^C HSQC, ^1^H-^13^C HSQC-TOCSY and ^1^H-^13^C HMBC spectra, to define the three W residues featured in **3**. Despite the complete absence of a suitable ^1^H-^1^H NOESY and the inability to uncover any ^1^H-^13^C HMBC correlations to connect these structural fragments together, weaker couplings (W-coupling, ^4^*J*_2H,βHa+b_) from ^1^H-^1^H TOCSY suggested the following possible connectivities, **TrpI--Trp7, TrpII--Trp2/3** and **TrpIII--Trp3/2**. Analogously, the phenomenon of having ^4^*J*_2H,βH_ couplings in W residues was similarly witnessed in the ^1^H-^1^H TOCSY spectrum of nocapeptin A (**1**) (Figures S55B and S57C).

As expected, the ^1^H-NMR of **3** displayed two downfield singlets, *δ*_H_ = 10.18, 10.55, indicative for only two indolic NHs of three Trp moieties, proposing a possible substitution at one indolic NH of these three W residues (Figure S51A). Tracking such downfield singlets through ^1^H-^1^H TOCSY, ^1^H-^13^C HSQC and ^1^H-^13^C HMBC enabled the structural elucidation of three (un)substituted aromatic systems, **Trp2/3**, **Trp3/2** and **Trp7** (Figure S56A). **Trp7** as an unsubstituted indolic fragment was readily assigned to be the aromatic part of W7 considering the CP sequence and the formerly annotated MS fragments. However, the monosubstituted (**Trp3/2**) and the disubstituted (**Trp2/3**) systems were envisioned to represent W2 and W3 residues, alternatively. In addition, the hydroxylation event was figured out to be introduced into **Trp2/3** substructure at position 5 as suggested by the ^1^H-^13^C HMBC correlations.

Although the structural finalization of the biaryl fragment of **3** was crippled due to the absence of a proper ^1^H-^1^H NOESY spectrum, the C-N crosslink was indirectly NMR deduced using the comparable shifts of nocapeptin A (**1**). The first evidence was counting on the delineated **TrpII--Trp2/3** as a disubstituted system lacking the indolic proton proposing 1N as an interlinkage atom. This was additionally supported by the characteristic *δ*_2C_ 146.46 which shares a comparable downfield shift to **1**. The second evidence was derived from **TrpIII--Trp3/2** as a monosubstituted system having a diagnostic *δ*_4H_ 6.31 with a comparable upfield drift to **1** (Figures S52A and S57C). As a result, two positional structural possibilities were laid out featuring the biaryl (C-N) installation between the structural units **1N-Trp2/3** and **5C-Trp3/2** in an alternating manner. Considering the current genomic and biosynthetic setting, the final product likely favors the substructure in which N1-W2 couples with C5-W3. (Figure S58)

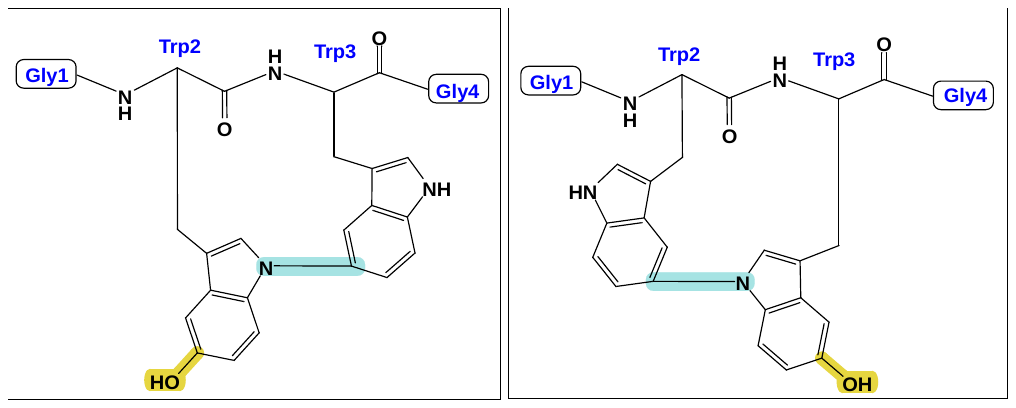

**Figure S58**. Schematic representation of the two possible biaryl substructures of longipeptin A (**3**) regarding the C-N linkage between **W2** and **W3** residues

**Table S10.** Results of the antimicrobial assays for **1**.

| **Bacterial strain** | **MIC (µg/ml)** | **Assay medium** |
| --- | --- | --- |
| *Bacillus subtilis 168* | *>64* |  |
| *Staphylococcus aureus ATCC 29213* | *>64* |  |
| *Enterococcus faecium BM 4147-1* | *>64* |  |
| *Enterococcus faecalis ATCC 29212* | *>64* |  |
| *Escherichia coli ATCC 25922* | *>64* |  |
| *Escherichia coli HN 818* | *>64* |  |
| *Escherichia coli HN 818 (+ 15 µg/ml PMBN)* | *>64* |  |
| *Klebsiella pneumonia ATCC 12657* | *>64* |  |
| *Enterobacter aerogenes ATCC 13048* | *>64* | MH II broth |
| *Pseudomonas aeruginosa ATCC 27853* | *>64* |  |
| *Acinetobacter baumannii 09987* | *>64* |  |
| *Micrococcus luteus ATCC 4698* | *16* |  |
| *Neisseria gonorrhoeae ATCC 19424* | *>64* | MH II broth |
| *Neisseria gonorrhoeae S 1441* | *>64* | + 2,5 % FBS |
| *Mycobacterium smegmatis mc2 155* | *>64* | 7H9 broth |

**Table S11.** Results of the cytotoxicity assays for **1**. Developmental Therapeutics Program (DTP)-One dose Mean Graph NCI-60 data.

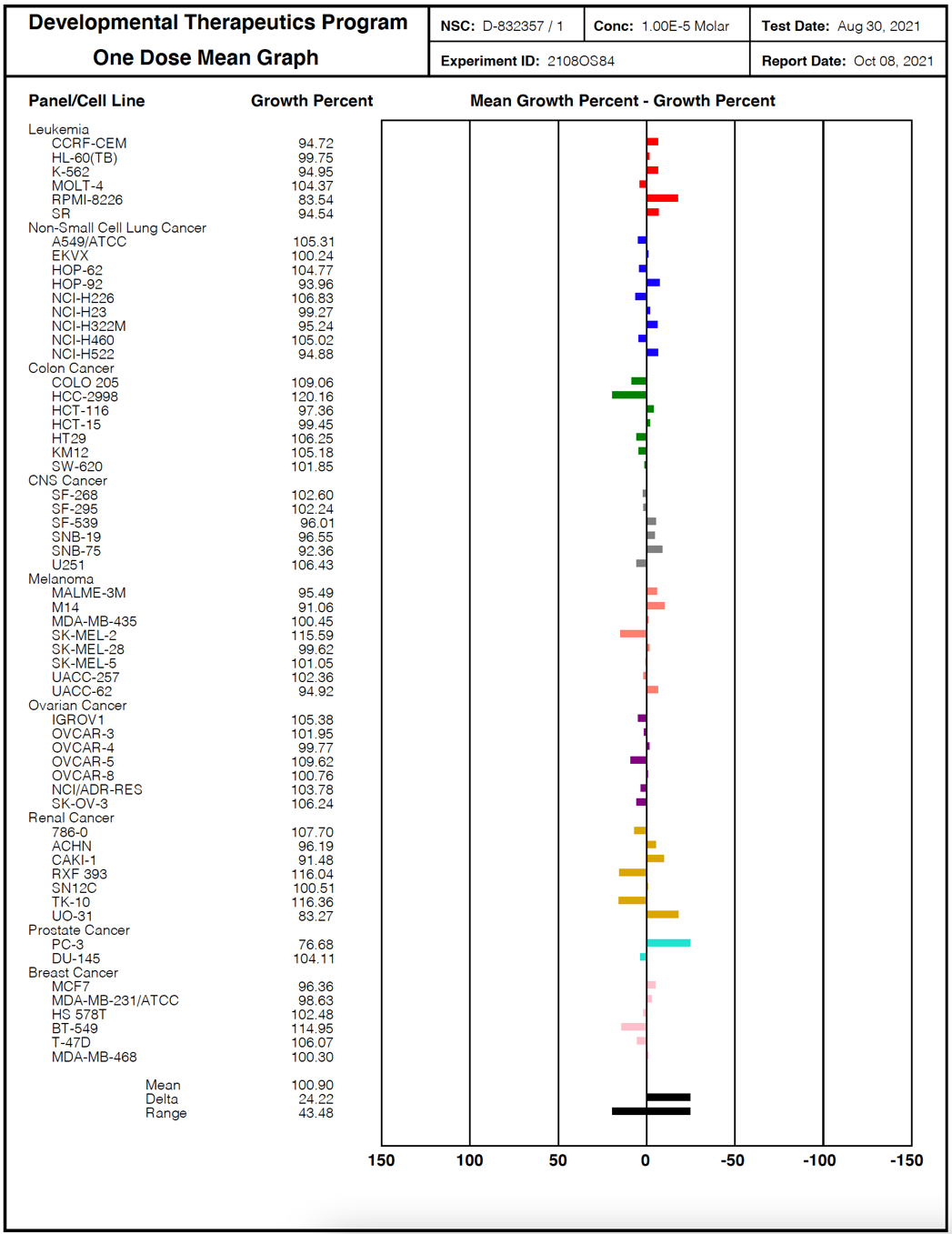

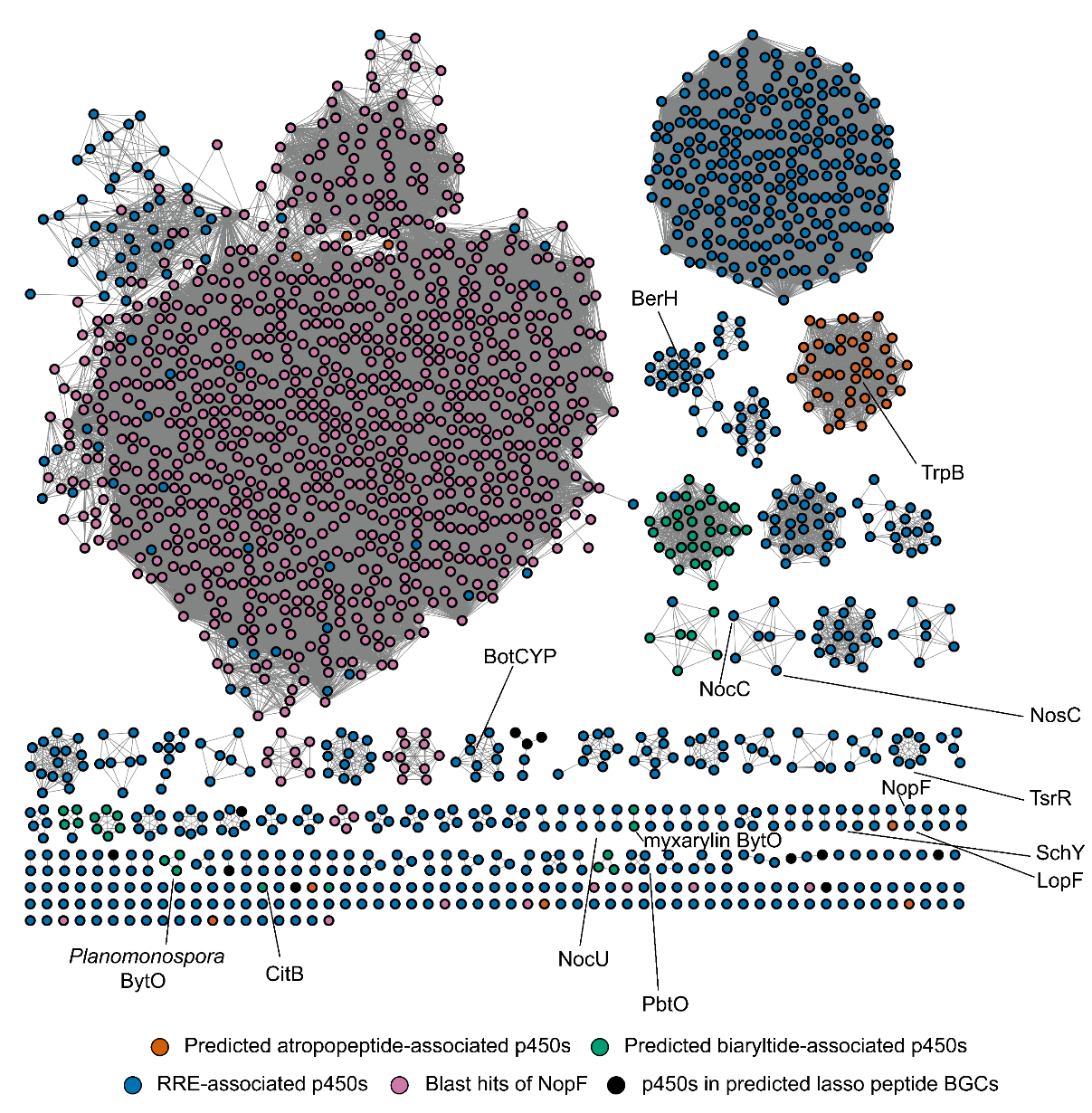

**Figure S59.** A sequence similarity network (SSN, RepNode 100) (alignment score = 99) of 1000 top BLAST-P hits to NopF, all cytochrome P450 proteins (883 sequences) within 10 open reading frames (ORFs) of a detected RRE (RRE-Finder, precision mode), predicted atropopeptide- and biaryltide-associated cytochrome P450 proteins,^[14]^ and cytochrome P450 proteins in predicted lasso peptide BGCs (Figure S67). Experimentally characterized cytochrome P450 proteins (NosC, TsrR, SchY, myxarylin and *Planomospora* BytO, CitB, NocU, PbtO, BotCYP, TrpB) and P450 proteins encoded in the BGCs of RiPPs with known structures (BerH; berninamycin, and NocC; nocathiacin) are annotated. A total of 1995 proteins were represented in this network (Supplementary Dataset 1).

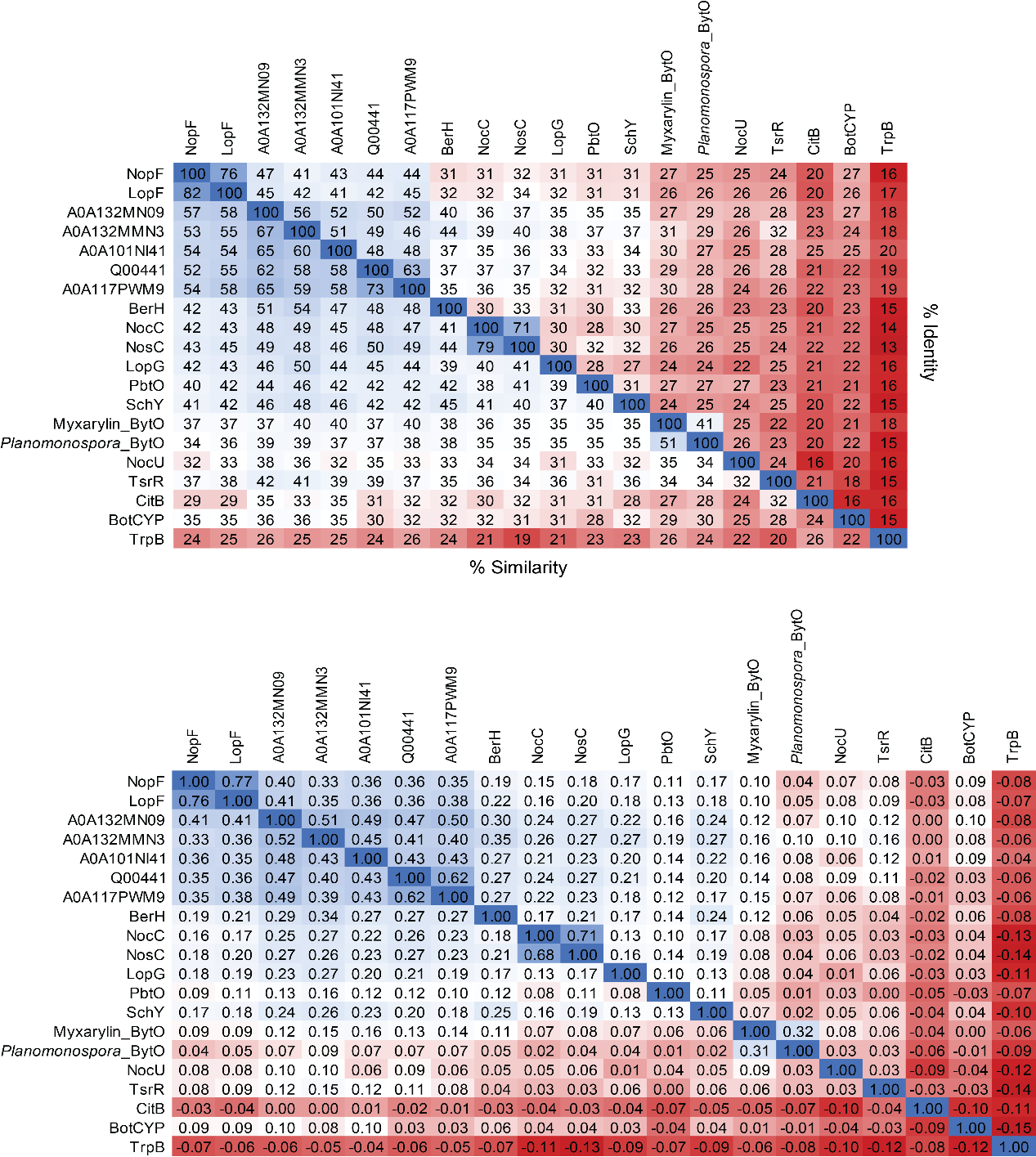

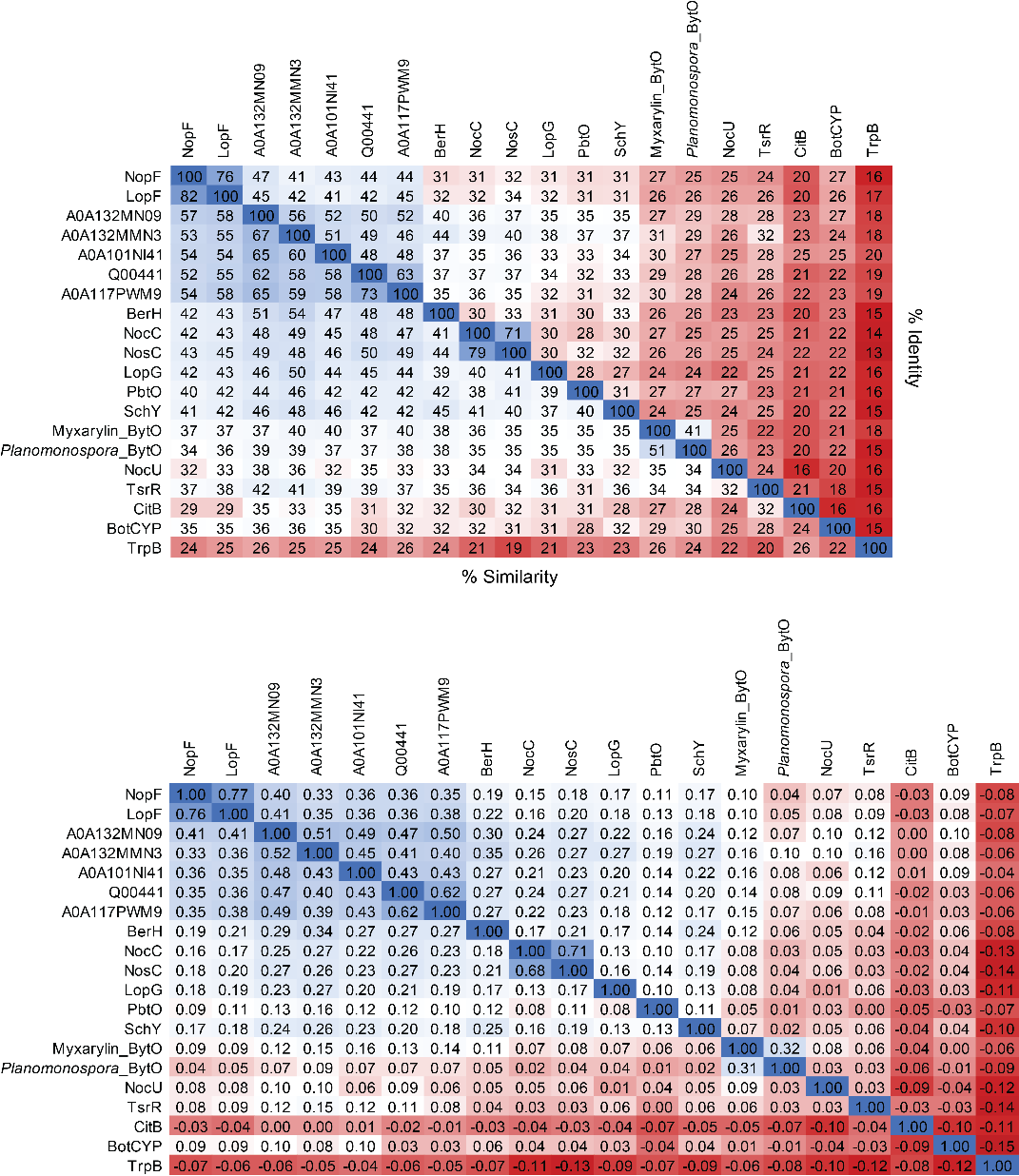

**B**

**A**

**Figure S60.** (**A**) Percent similarity and identity matrix and (**B**) global similarity matrix (BLOSUM62) of three cytochrome P450 proteins uncovered in this study (NopF, LopF, and LopG), experimentally characterized cytochrome p450s annotated in the SSN (Figure S59), and five top BLAST-P hits to NopF (depicted by UniProt accession codes).

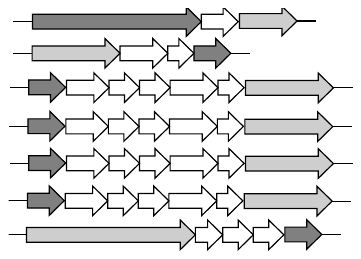

*Asanoa siamensis*

*Streptomyces* sp. KS 21

*Streptomyces* sp. A1136

*Streptomyces* sp. NRRL S-241

*Streptomyces* sp. ISL-21

*Kitasatospora* sp. RG8

*Kitasatospora purpeofusca*

**Figure S61**. Representative genomic neighborhoods of selected BLAST-P hits to LopH (dark grey; WP_051757134.1, WP_209415770.1, WP_051779479.1, WP_214943134.1, WP_136214693.1, WP_133900279.1, WP_203716497.1, respectively) show a strong co-occurrence with DNA polymerase III, beta subunit (light grey, PF00712/TIGR00663).

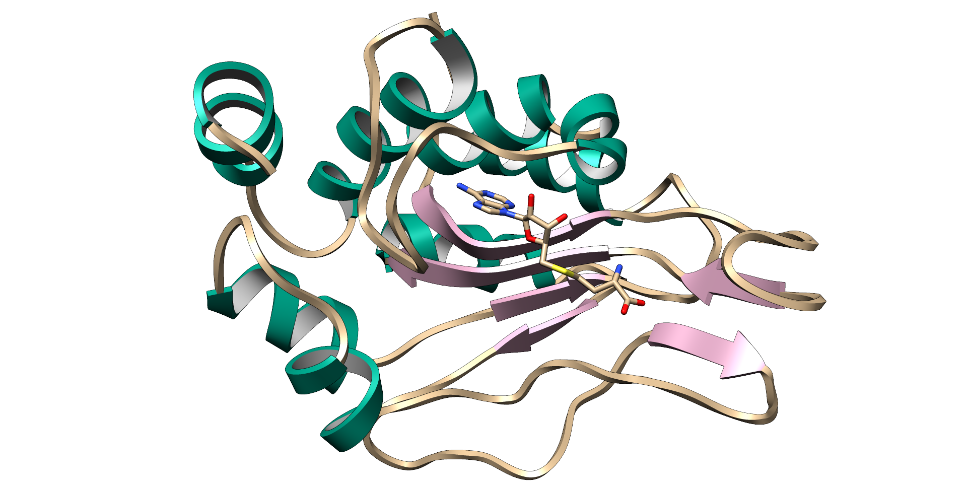

**A**

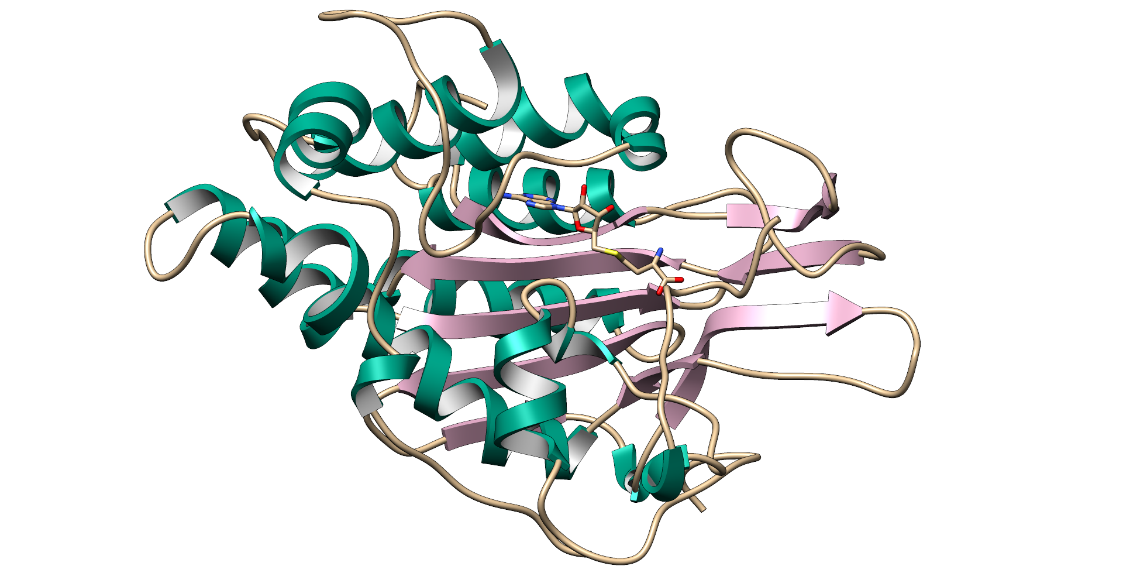

**B**

**Figure S62.** Tertiary structure comparison between (**A**) AlphaFold-predicted LopH with S-Adenosyl-L-homocysteine (SAH) docked through energy minimization, and (**B**) the closest LopH match upon performing a DALI search: the SAH-binding region of human 5,10-methylenetetrahydrofolate reductase (PDB code: 6fcx, residue 344-646, chain A). Parameters generated from DALI: Z-score = 11, RMSD = 3.1, number of aligned C-alpha atoms = 150, number of residues in target structure = 252. The coordinates for panel A are given as supplementary file 1. UCSF Chimera was used to generate these images.

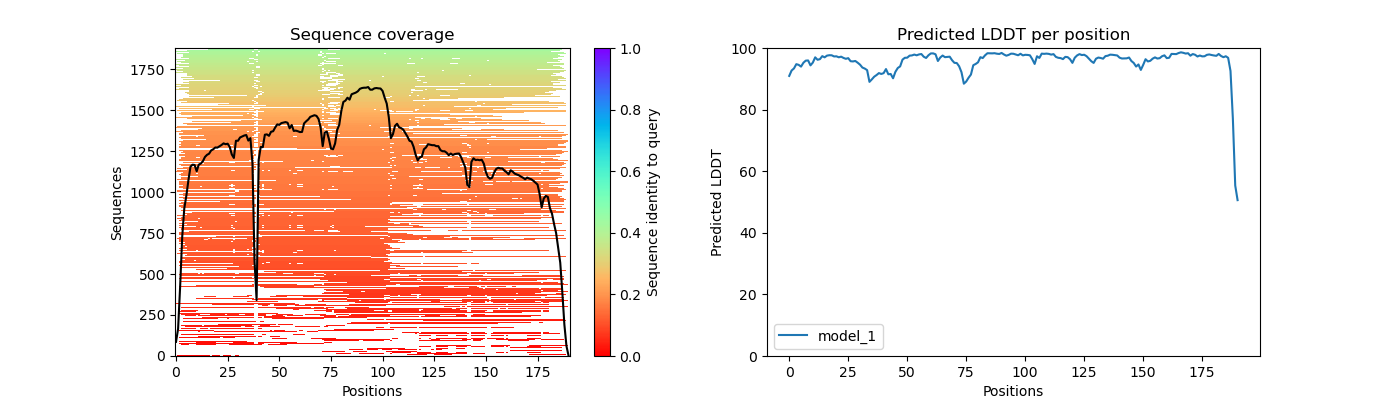

**C**

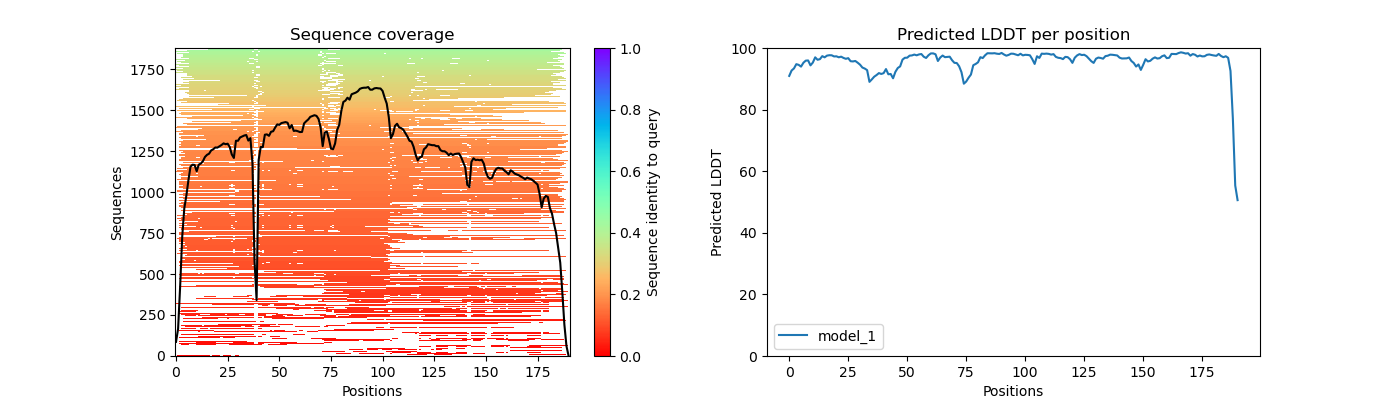

**B**

**A**

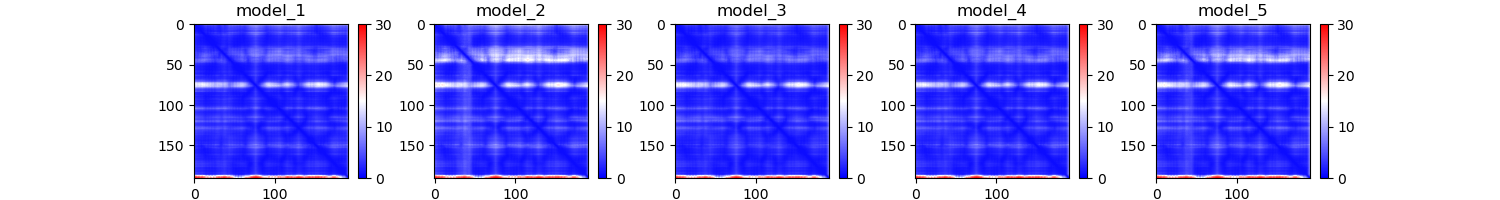

**Figure S63**. Confidence analysis of the AlphaFold-predicted LopH structure. (**A**) The predicted aligned error (pAE) plot, (**B**) the multiple sequence alignment summarized as a heatmap, and (**C**) the predicted LDDT per position (pLDDT) plot. The pAE and pLDDT plots correspond to the highest-ranked LopH structure generated by AlphaFold.

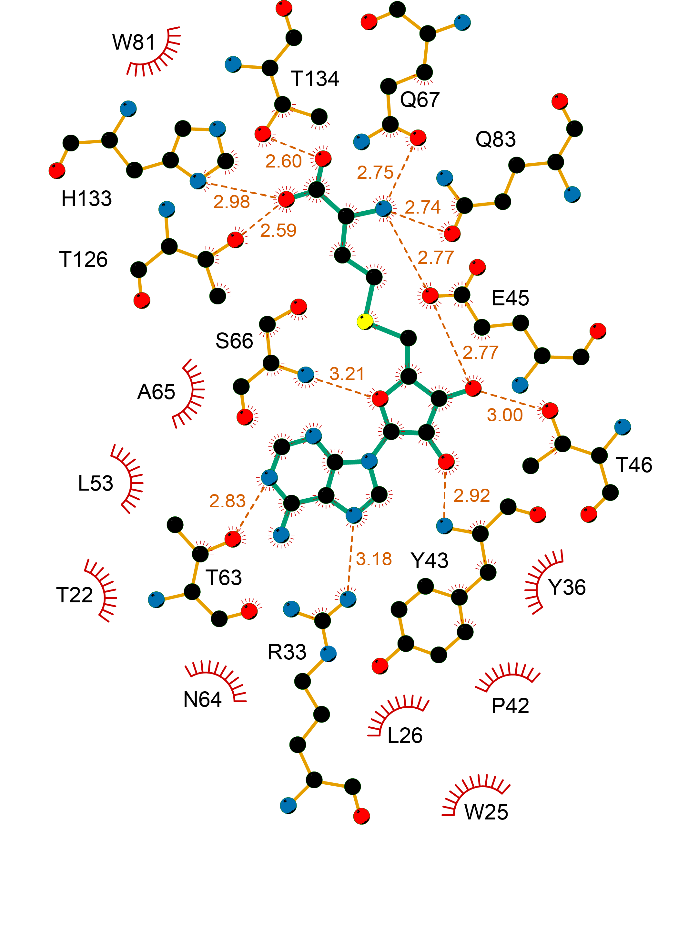

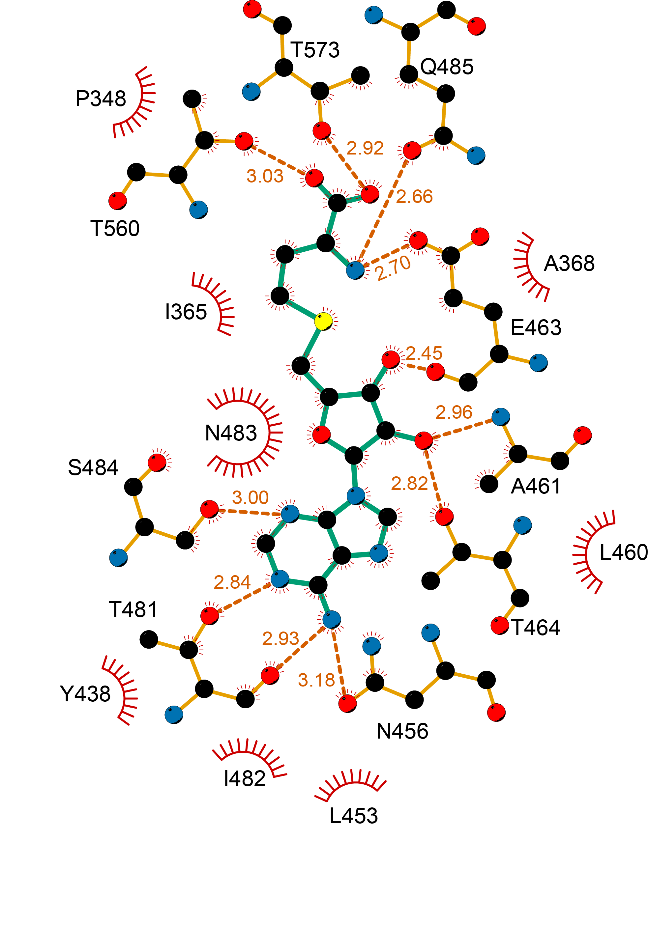

**B**

**A**

**Figure S64.** (**A**) A ligand interaction diagram for SAH docked with LopH through energy minimization, (**B**) A ligand interaction diagram for the crystal structure of SAH bound to human 5,10-methylenetetrahydrofolate reductase (PDB code: 6fcx, chain A). Predicted hydrogen bonds are described in orange with distances indicated, while red arcs denote hydrophobic interactions. LigPlot^+^ was utilized to generate these ligand interaction diagrams.

**Figure S65**. A secondary-structure alignment generated by DALI between LopH and human 5,10-methylenetetrahydrofolate reductase (Z-score = 11, RMSD = 3.1, number of aligned C-alpha atoms = 150). The three-state secondary structure definitions by DSSP (reduced to h= helix, e=sheet, l=coil) are utilized. Residues directly contacting the ligand are blue (for LopH) and orange (for 6fcx).

**B**

**A**

**Figure S66.** Structures depicting SAH interaction with amino acid residues in the (**A**) LopH structure predicted by AlphaFold, and (**B**) the crystallized human 5,10 -methylenetetrahydrofolate reductase structure (PDB code: 6fcx). The coordinates for panel A are given as supplementary file 1. UCSF Chimera was used to generate these images.

*Embleya hyalina*

*Streptomyces* sp. L-9-10

*Streptomyces* sp. 2323.1

*Nocardia nova Nocardia* sp. 852002-20019_SCH5090214

*Jannaschia* sp. EhC01

*Rhizobium lusitanum*

*Leptolyngbya* sp. Heron Island J

*Rivularia* sp. PCC 7116

*Saccharopolyspora phatthalungensis*

*Tistrella mobilis* KA081020-065

bacterium Bacteria

*Adonisia turfae* CCMR0081

*Niveispirillum* sp. SYP-B3756

*Tistrella* sp.

*Streptomyces* sp. 2314.4

*Streptomyces* sp. 2333.5

*Mycolicibacter engbaekii*

*Nocardia transvalensis*

*Streptomyces alkaliterrae*

*Streptomyces huiliensis*

*Streptomyces lavendofoliae*

*Streptomyces piniterrae*

*Nonomuraea endophytica*

*Amycolatopsis vastitatis*

*Kribbella catacumbae* DSM 19601

**Figure S67**. Expanded list of predicted lasso peptides BGCs associated with cytochrome P450 proteins.

**Table S12.** The prediction of the corresponding core peptide of each lasso peptide gene cluster

| **Precursor identifier** | **CYP450 identifier** | **Species** | **Predicted Core Peptide Sequence** |
| --- | --- | --- | --- |
| GCD92712.1 | GCD92710.1 | *Embleya hyalina* | RKPGPWWEWIGPNYLD |
| WP_020387001.1 | WP_020387006.1 | *Kribbella catacumbae* DSM 19601 | GSGGHNWEWIDAWGW |
|  | WP_020387008.1 |  |  |
|  | WP_020387009.1 |  |  |
| unannotated | OXM64042.1 | *Amycolatopsis vastitatis* | SNCGRTWEWVFCGESRC |
| unannotated |  |  | GPCGNQWEWIACGTEC |
| unannotated |  |  | GGQGCNWEWIGCGWC |
| MBB5084694.1 | MBB5084699.1 | *Nonomuraea endophytica* | AGCGCVWEWLTDRLDW |
| MBB5084695.1 |  |  | GGCGGPHWEWIYPNYCR |
| TJZ59061.1 | TJZ59062.1 | *Streptomyces piniterrae* | LLAKHGNDRLIFSKN |
| GGU49781.1 | GGU49789.1 | *Streptomyces lavendofoliae* | LAGQGSPDLLGGHSLL |
| WP_223766706.1 | WP_223766708.1 | *Streptomyces huiliensis* | ANKQGMGFDWYLTRK |
| MQS00863.1 | MQS00861.1 | *Streptomyces alkaliterrae* | ANRRGMGFDWYLTRK |
| MBB5915993.1 | MBB5915992.1 | *Nocardia transvalensis* | GSSYVIIEGYPSAWGSNY |
| WP_109560825.1 | ORV51843.1 | *Mycolicibacter engbaekii* | GSGNYLSDSSTGYGYMGWYNRHCDTPETAAPLPPRA |
| WP_146098681.1 | PPJ28635.1 | *Nocardia nova* | GVFTTESDLLVGRRGMI |
| WP_146098681.1 | OBA54897.1 | *Nocardia* sp. 852002-20019_SCH5090214 | GVFTTESDLLVGRRGMI |
| SEE71607.1 | SEE71688.1 | *Streptomyces* sp. 2314.4 | GTNSFDTADDFSVKSCVLELHVAAR...STCDWALASVQAPEMHGAERCGGLE |
| PJJ05250.1 | PJJ05254.1 | *Streptomyces* sp. 2333.5 | GTNSFDTADDFSVKSCVLELHVAAR...STCDWALASVQAPEMHGAERCGGLE |
| WP_159394969.1 | SOE10351.1 | *Streptomyces* sp. 2323.1 | GTNNFDTADDTQYKNA |
| unannotated | MAM73040.1 | *Tistrella* sp. | GTFSGSGSDSSYS |
| RYJ28010.1 | RYJ28015.1 | *Streptomyces* sp. L-9-10 | FFRNGANEAYFFFQNQND |
| RYJ28011.1 |  |  | VFGIRNGDEITWFFDTWQ |
| RYX82715.1 | RYX82717.1 | bacterium Bacteria | ASRVGSKLDRAINAGPGTPVGPLLNEISNSLS |
| unannotated | NEZ55279.1 | *Adonisia turfae* CCMR0081 | TSTTFVGSDGGSGIFQYAS |
| MQP68237.1 | MQP68242.1 | *Niveispirillum* sp. SYP-B3756 | SNGNDDGSDSMYS |
| unannotated | MBB6488871.1 | *Rhizobium lusitanum* | GSAGPLAFDFHLSDRNT |
| MBB5158765.1 | MBB5158767.1 | *Saccharopolyspora phatthalungensis* | GNQYLYVEGFFSYLGTI |
| AFK55229.1 | AFK55230.1 | *Tistrella mobilis* KA081020-065 | SGGSGPGSDNNYS |
| WP_107073100.1 | KJS59601.1 | *Streptomyces rubellomurinus* | ALGLHGAEPFFPTLHTSWW |
| WP_015121441.1 | AFY57880.1 | *Rivularia* sp. PCC 7116 | ANAPNFPFNGFDGGSSPNNYAS |
| unannotated | ESA35183.1 | *Leptolyngbya* sp. Heron Island J | TSTTFVGSDGGSGIFQYAS |
| WP_161489756.1 | OAN82783.1 | *Jannaschia* sp. EhC01 | NNSGSGSDAGIYSS |

**Relevant Known Scaffolds Containing the PTMs under Investigation:**

**Peptides-Based Crosslinks**

A) **YYH** (R= OH), and **YFH** (R= H), Cell Chem. Bio. 2021, 28, 733-739, **RiPPs, CYP450**
B) **MeYLH**, Molecules 2021, 26(24), 7483, **RiPPs, CYP450**
C) **Cittilin A** (R= OMe) and **B** (R= OH) **RiPPs, CYP450**

A) **Tryptorubin A**, JACS 2017, 139, 12899-12902, **RiPPs, CYP450 B) Darobactin A**, Nature 2019, 576, 459-464, **RiPPs, rSAM**

A) **Chloropeptin** I; JNP 2001, 64, 874-882, **NRPS** **B)** **Aspergilazine A**, Tetrahedron Lett. 2012, 53, 2615-2617, **NRPS**

A) **TMC-95A**, J. Org. Chem. 2000, 65, 990–995 **B)** **Diazonamide A,** JACS 1991, 113, 2303–2304

**Known S-methylated RiPPs**

A) **Sch 40832**, J. Antibiot. 1998, 51, 221−224 **B)** **Thioxamycin** and **Thioactin,** J. Antibiot. 1989, 42, 1465−1469

J. Antibiot. 1994, 47, 1541-1545

**Known S-methylated non-RiPPs**

**A) Echinomycin,** Nat. Chem. Biol. 2006, 2, 423−428, **NRPS** **B) Thiocoraline,** JACS 2014, 136, 17350−17354, **NRPS**

**A) Maremycin A/B, NRPS	B) FR900452, NRPS	C) Maremycin G,** ACS Chem. Biol. 2018, 13, 2387–2391, **NRPS**

**A) Bismethylgliotoxin,** Chem. Biol. 2014, 21, 999−1012 **B) Collismycin,** J. Antibiot. 1994, 47, 1072−1074

**C) Lincomycin,** Adv. Appl. Microbiol. 2004, 56, 121−154 **D) Brassinin,** Nat. Prod. Rep. 2011, 28, 1381−1405

**DMSP and Related Compounds**
